# Descent of Bacteria and Eukarya from an archaeal root of life

**DOI:** 10.1101/745372

**Authors:** Xi Long, Hong Xue, J. Tze-Fei Wong

## Abstract

The three biological domains delineated based on SSU rRNAs are confronted by uncertainties regarding the relationship between Archaea and Bacteria, and the origin of Eukarya. Herein the homologies between the paralogous valyl-tRNA and isoleucyl-tRNA synthetases in a wide spectrum of species revealed vertical gene transmission from an archaeal root of life through a Primitive Archaea Cluster to an Ancestral Bacteria Cluster of species. The higher homologies of the ribosomal proteins (rProts) of eukaryotic *Giardia* toward archaeal relative to bacterial rProts established that an archaeal-parent rather than a bacterial-parent underwent genome merger with an alphaproteobacterium to generate Eukarya. Moreover, based on the top-ranked homology of the proteins of *Aciduliprofundum* among archaea toward the *Giardia* and *Trichomonas* proteomes and the pyruvate phosphate dikinase of *Giardia*, together with their active acquisition of exogenous bacterial genes plausibly through *foodchain gene adoption*, the *Aciduliprofundum* archaea were identified as leading candidates for the archaeal-parent of Eukarya.

---

Molecular evolution analysis of small subunit ribosomal RNA (SSU rRNA) yielded a universal but unrooted tree of life (ToL) that comprises the three biological domains of Archaea, Bacteria and Eukarya^1^. A ToL of transfer RNAs based on the genetic distances between the 20 classes of tRNA acceptors for different amino acids located its root near the hyperthermophilic archaeal methanogen *Methanopyrus* (Mka)^2^. Although this rooting is supported by a wide range of evidence^3–9^, and the age of ~2.7 Gya for the *Methanopyrus* lineage as the oldest among living organisms^10^, the phylogenies of the three biological domains are beset by two fundamental problems: viz. the uncertain relationship between Archaea and Bacteria, and the identity of the prokaryotic-parent that underwent genome merger with an alphaproteobacterium to give rise to Eukarya. As long as these two problems remain unresolved, the nature of the root of life is open to diverse formulations^11–15^. Accordingly, the objective of the present study is to examine the pathways of descent of Bacteria and Eukarya from an archaeal root of life, and the nature of the archaeal-parent of Eukarya.

The antiquity of proteins could be assessed based on the increasing divergence of paralogous proteins in time^16^. Applying this approach, BLASTP was performed between the intraspecies valyl-tRNA synthetase (VARS) and isoleucyl-tRNA synthetase (IARS) in the genomic sequences for 5,398 species in NCBI Genbank. Arrangement of the BLASTP bitscores obtained in descending order (Supplementary Table S1 and partly in Fig. 1) showed that the 119 highest scoring species were all archaeons, topped by Mka and including Mfe, Afu, Mnt and Mja with bitscores of 473, 436, 387, 387 and 387 respectively. The top scoring bacterium was the Clostridium Mau with a bit score of 378, and the top scoring eukaryote was the filamentous brown alga Esi with a bit score of 240. These results established the foremost antiquity of Mka among extant organisms.

**Figure 1.**
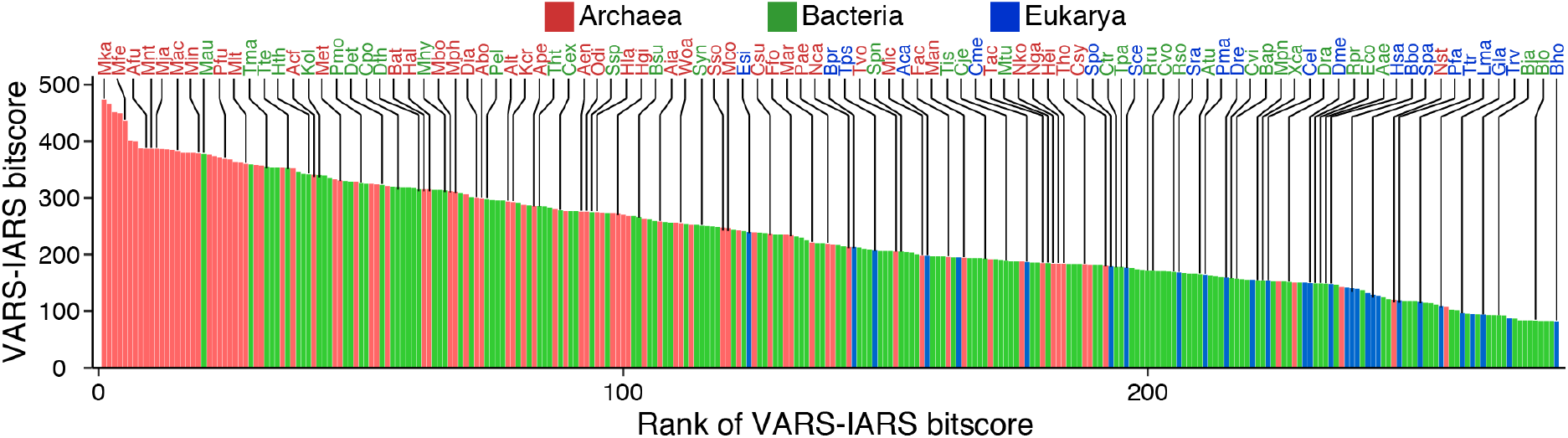
Ranking of bitcores for the VARS-IARS homology of species of organisms in descending order (from left to right). Bitscores were obtained using BLASTP for a selection of archaeal (red), bacterial (green) and eukaryotic (blue) species. The complete list of 1,185 archaeal, 3,621 bacterial and 592 eukaryotic species from NCBI Genbank release 231 and their bitscores are given in Supplementary Table S1. The full names and three-letter abbreviations for a number of the species are indicated in Table 1.

The positions of some of the species analyzed in Fig. 1 were indicated on the SSU rRNA tree, with their intraspecies VARS-IARS bitscores expressed in circles colored according to the thermal scale (Fig. 2a). There was a concentration of euryarchaeons with high VARS-IARS homology in a ‘Primitive Archaea Cluster’ spread between Hal and Mfe. In the Bacteria domain, there was likewise an ‘Ancestral Bacteria Cluster’ with high VARS-IARS homology spread between Det and Mau. The deepest branching species in the Bacteria domain were two members of the Aquificae phylum, viz. the anaerobic Det with high VARS-IARS homology, and the microaerobic, low-homology Aae. Since mutations could cause loss of homology more easily than gain of homology, this suggests that Aae has evolved far from the ancestral Aquificiae species possibly as part of the wave of tumultuous changes undergone by former anaerobes in response to the appearance of atmospheric oxygen^17^, thereby sustaining extensive evolutionary erosion of its VARS-IARS homology. The enhanced resistance of paralogue homology to perturbation by horizontal gene transfer (HGT), due to the difficulty of transfer of a pair of proteins compared to transfer of a single protein, was illustrated by the preservation of low VARS-IARS bitscores in the proteobacterial region of the tree against any shift toward elevated VARS-IARS homology on account of HGT events, despite the high HGT-susceptibility of for example Eco, which acquired 18% of its genes through HGT subsequent to its departure from *Salmonella enterica* about 100 million years ago^18^. Previously, based on the intraspecies alloacceptor tRNA-distances of various species on the tRNA tree, LUCA was positioned between the branches leading to Mka and Ape at a distance ratio of 1.00 from the Mka branch versus 1.14 from the Ape branch^2^, and this position was adopted in the SSU rRNA tree in Fig. 2.

**Figure 2.**
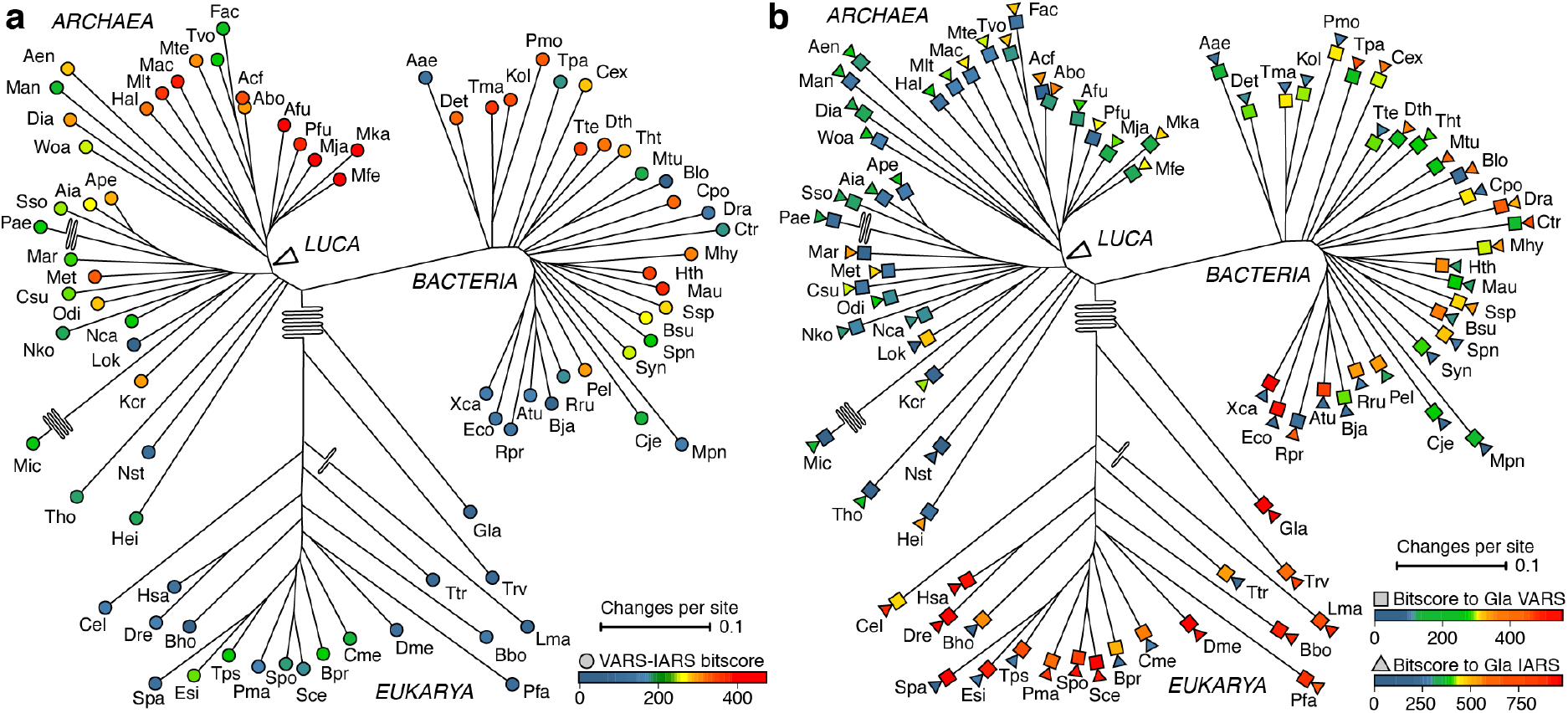
The rooted universal SSU rRNA tree. The tree was constructed using the Fitch-Margoliash method from the alignment of SSU rRNAs from 33 archaeal, 31 bacterial, and 19 eukaryotic species. Pairwise distances between the aligned sequences were estimated using DNADIST in PHYLIP with Jukes and Cantor model. The thermal scale in (a) expresses the bitscore for the VARS-IARS pair of each species (Supplementary Table S1), and that in expresses the genetic distances of VARS (squares), or IARS (triangles) between Gla and other organisms on the tree.

Given the relative paucity of HGT effects on VARS-IARS homology in a majority if not all of the species on the SSU rRNA tree, the parallel prominence of the high VARS-IARS homology species in the Primitive Archaea Cluster and the Ancestral Bacteria Cluster was explicable by vertical genetic transmission of the VARS and IARS genes from an Mka-proximal root of life to the archaeal cluster, and in turn to the bacterial cluster. Since the top ranked bacterial bitscore of Mau at 378 was between that of Mac at 382 and Pfu at 369, the results suggest that the Ancestral Bacteria Cluster branched off from the Primitive Archaea Cluster close to the Mka-proximal root of Archaea. The medium VARS-IARS bitscores of Esi, Tps, Bpr and Cme among the Eukarya (Fig. 2a) were indicative of the extension of the intraspecies VARS-IARS homology into this domain. The much higher VARS homologies (colored squares) and IARS homologies (colored triangles) between bacterial species and the eukaryote Gla compared to those between archaeal species and Gla indicated that Eukarya received the VARS and IARS genes from the Bacteria instead of the Archaea domain (Fig. 2b).

The aligned segments of VARS and IARS from the archaeon Mka, the bacterium Mau and the eukaryote Esi in Fig. 3 were portions of the six complete sequences (Supplementary Fig. S1). These segments showed 42/207 columns where all six sequences carried the same amino acid, thereby providing strong evidence for the vertical transmission of VARS and IARS genes from Archaea to Bacteria and Eukarya.

**Figure 3.**
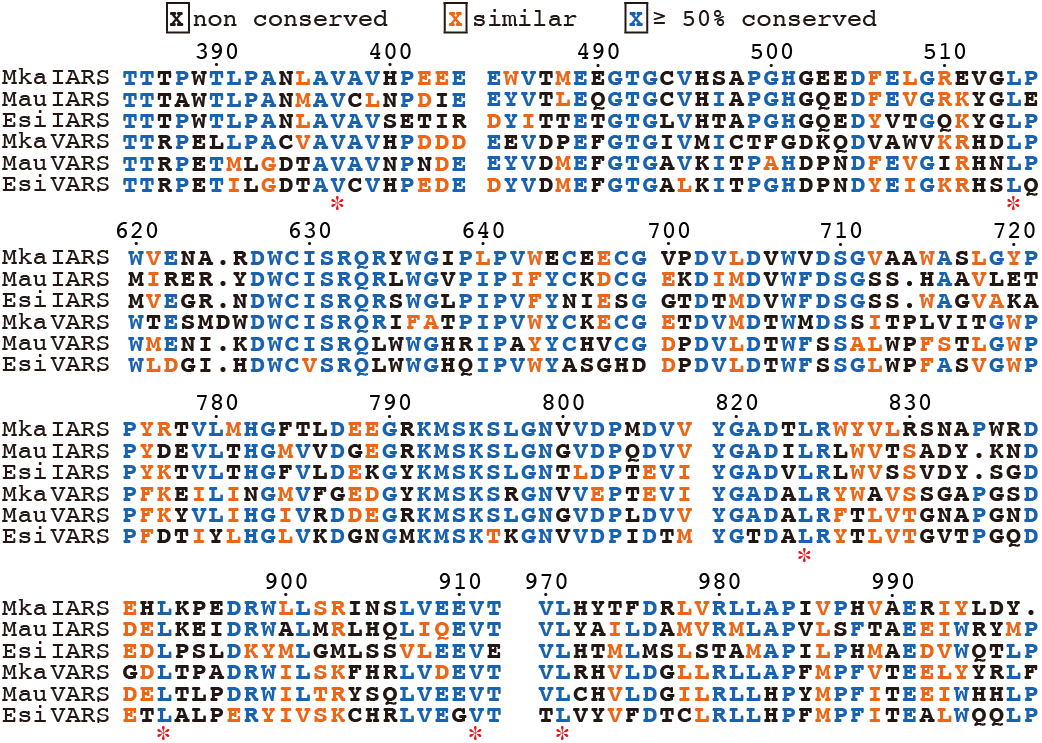
Segments of the aligned VARS and IARS sequences of Mka, Mau and Esi. The six sequences were aligned using ClustalOmega^50^. Tick marks indicate the positions of the sequence segments on the complete alignment (Supplementary Fig. S1). Similar amino acids are colored in orange, and ≥ 50% conserved ones in blue. Asterisks indicate the six positions displaying conservation of the same V or L amino acid in all six sequences.

It has been suggested that an endosymbiotic event between an archaeal-parent and an alphaproteobacterium acting as mitochondrion-parent led to the formation of the Last Eukaryotic Common Ancestor (LECA) and ushered in the Eukarya domain^19–21^. Proposals regarding the identity of the archaeal-parent have focused on a range of prokaryotes including Thermoplasmatales where the lack of a rigid cell wall could facilitate its engulfment of the mitochondrion-parent to bring about endosymbiosis^22–24^; and various archaeons, especially the Asgard archaeons Lok and Tho^25,26^, that are enriched in eukaryotic signature proteins (ESPs)^27^. There is no consensus regarding a choice between these two groups of organisms^28^.

Upon BLASTP comparisons of the fifty-six ribosomal proteins (rProts) of Gla, the lowest branching eukaryote on the SSU rRNA tree, with corresponding prokaryotic rProts, fifty-three of them yielded higher bitscores with archaeons relative to bacteria (Fig. 4a), indicating that eukaryogenesis began with an archaeal-parent instead of a bacterial-parent. S21e and L36e yielded no bitscore with any archeal or bacterial rProt, suggesting that they were derived from a prokaryote not surveyed in the present study, altered beyond recognition by BLASTP, or invented by the eukaryogenesis system. The S7e of eleven eukaryotes from Table 1 although not that of Gla showed detectable homology toward the archaeon Alt. The Sce, Esi and Hsa rProts differed from those of Gla in two aspects: about one-sixth of the rProts that showed higher homology toward archaea relative to bacteria in Gla switched to lower homology toward archaea relative to bacteria in Sce, Esi and Hsa; and there were also some additional rProts in Sce, Esi and Hsa, mainly bacteria-derived ones, not found in Gla (Supplementary Fig. S2). These findings pointed to a significant influx of bacterial rProts into Sce, Esi and Hsa after their divergence from Gla, resulting in the replacement of some of the archaea-derived rProts found in Gla by bacteria-derived ones. Acf, Abo and Mac displayed the highest BLASTP bitscores among archaeons toward Gla pertaining to pyruvate phosphate dikinase (EC2.7.9.1), which would be consistent with a possible role of these three archaeons as archaeal-parent. Interestingly, the bitscores were high for Hei and Tho but low for Odi and Lok among the Asgard archaea, and high for Tac and Tvo but low for Fac, Min and Mte among the Thermoplasmatales (Fig. 4b).

**Figure 4.**
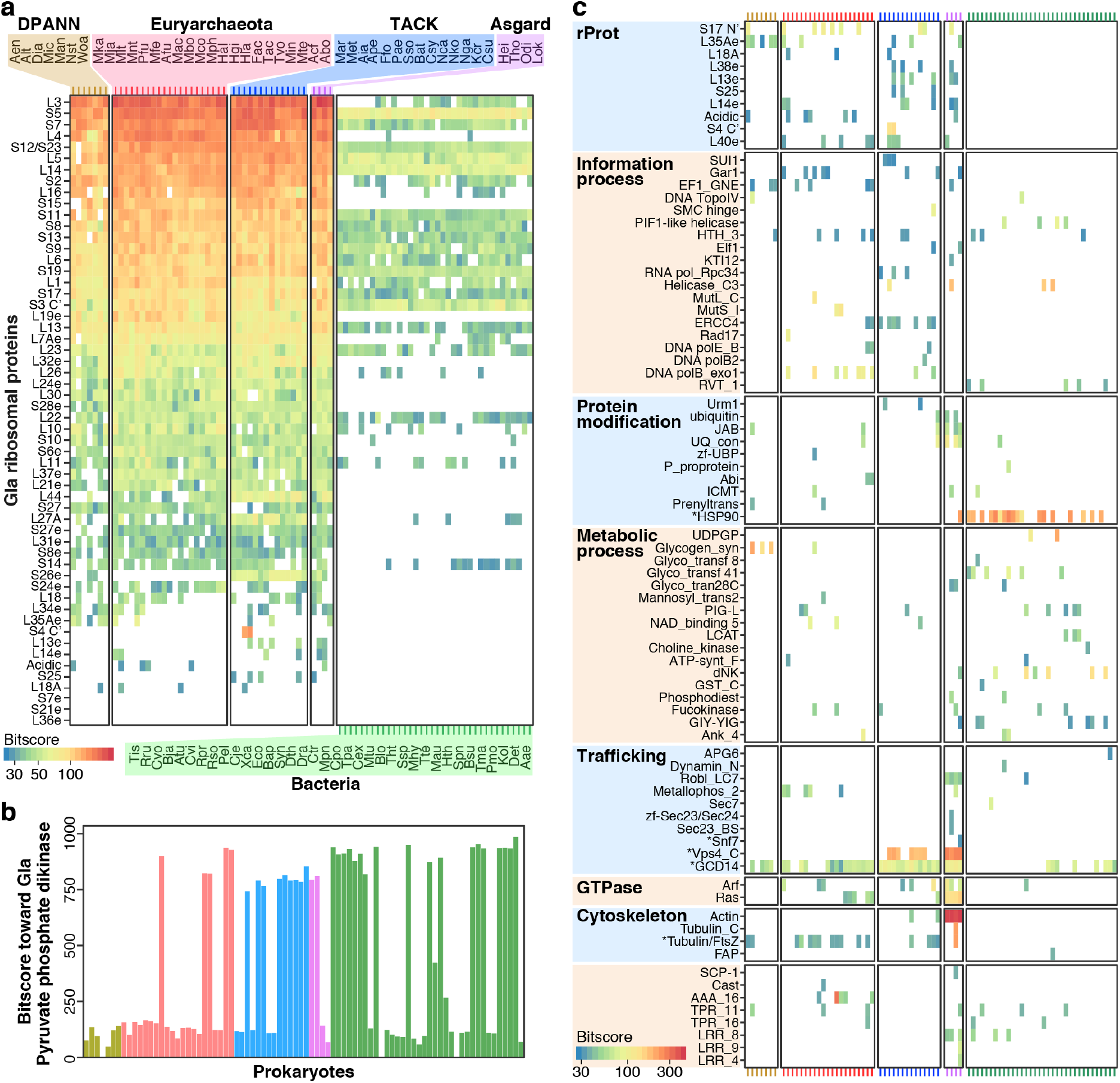
Proteins sequence homology between eukaryotic and prokaryotic species. (a) BLASTP bitscores between Gla rProts and prokaryotic rProts. (b) Bitscores between Gla pyruvate phosphate dikinase and prokaryotic proteins. Among all Gla proteins, this protein elicited the highest total BLASTP bitscore from all the prokaryotes. (c) Bitscores displayed by some of the 162-Gla proteins from Supplementary Table S2 towards prokaryotes. The color coding and order of different prokaryotic species on the x-axis in (b) and (c) are the same as those in (a). Bitscores with E-value < 0.05 are shown, except for asterisked entries in where E-value < 0.1.

**Table 1.**
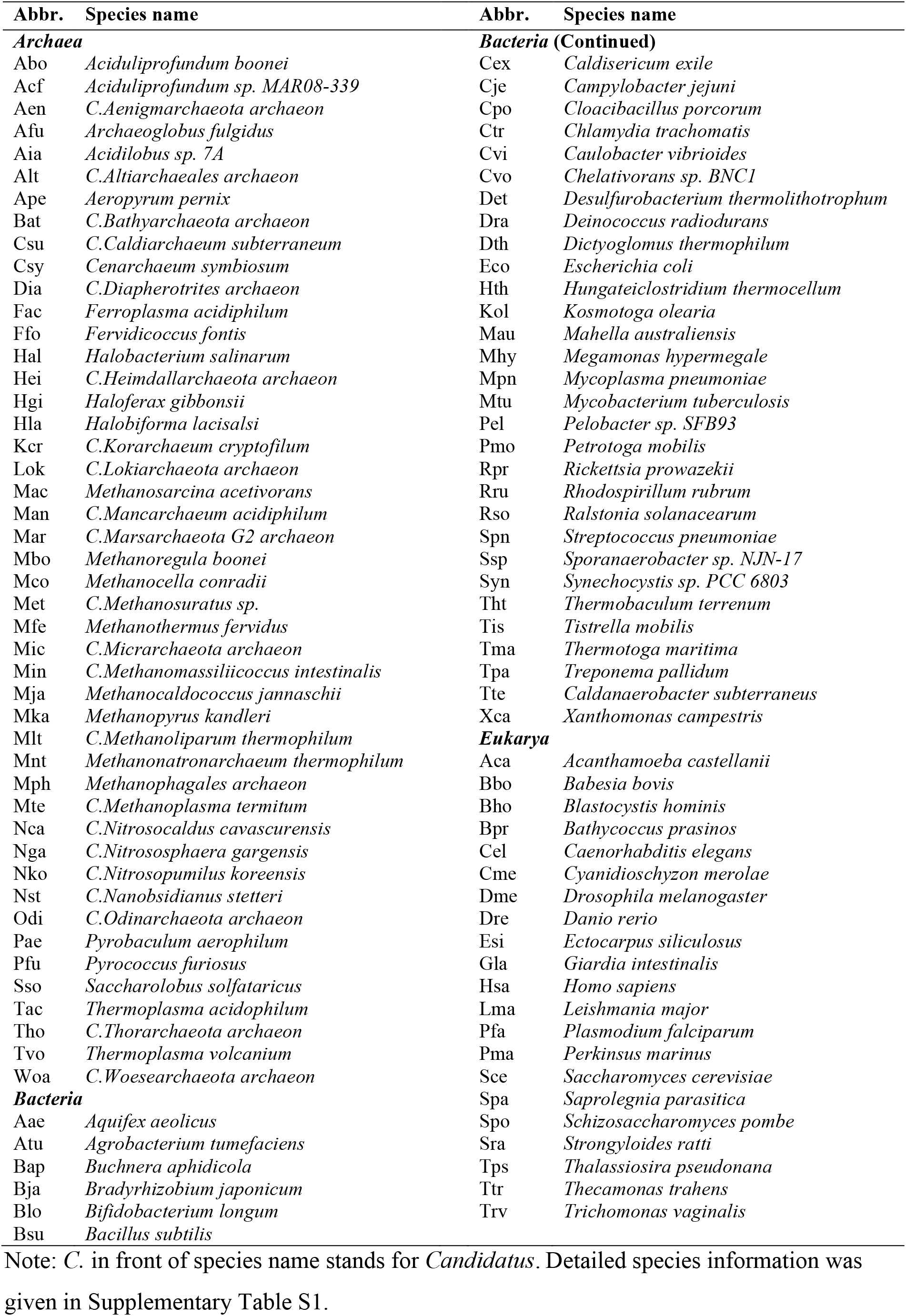
Partial list of species analyzed.

Figure 4c shows the prokaryotic distribution of homologues of some the 162 Gla proteins that were ESPs or homologous toward a limited number of prokaryotes (Supplementary Table S2). Tho, Odi, Xca and Lok, the four prokaryotes endowed with the largest numbers of the Gla-homologous proteins, harbored only 26, 19, 17 and 16 of these Gla-homologous proteins respectively (Supplementary Table S3), and the four Asgard archaeons also did not fully share their Gla-like proteins with one another via HGT, thus underlining the difficulty for any one archaeon or bacterium to accumulate a sufficient number of eukaryote-type proteins to launch the Eukarya domain relying only on their own inventiveness and HGT. On the other hand, one or more potential prokaryotic sources were found for each of the Gla-homologous proteins despite the survey of only a small spectrum of prokaryotes, indicating that the obstacle to eukaryogenesis posed by gene deficiency may be overcome if some dependable mechanism were available to assemble the requisite genes from a broad spectrum of prokaryotes. Addressing the inadequacy of ESP coverage by single archaeal species^29,30^, it was suggested that HGTs, or development of phagocytosis by an ESP-rich archaeon might provide a solution^26,31^. However, the frequencies of HGTs might be a limiting factor^10,32^, and rProts could be particularly resistant to HGT^33^.

During eukaryogenesis, the archaeal-parent might join up with the mitochondrion-parent to develop directly into LECA in a *mitochondria-early* scenario, or it might serve as First Eukaryotic Common Ancestor (FECA), and go through successive generations of genomic expansion prior to merger with the mitochondrion-parent to form LECA in a *mitochondria-late* scenario^34^. By measuring the phylogenetic distances between different components of LECA and their closest prokaryotic relatives, evidence has been obtained in support of a mitochondria-late time table, with the appearance of nucleolus preceding that of nucleus, endomembrane system and finally mitochondria^35^. Previously, the proteins of the eukaryote Sce were observed to contain a substantial variety of bacterial proteins and also some archaeal ones, and it was pointed out that the influx or bacterial genes into Sce could not be explained by a merger between archaeal-parent with another bacterium besides the mitochondrion-parent, or by Sce uptake of bacterial genes through ingestion of bacteria as food. Instead, the mitochondrion-parent was a major source of the exogenous bacterial proteins in Sce^20^. When the eukaryotic Gla and Trv proteomes were employed as homology probes for BLASTP query against various prokaryotic proteomes, it gave rise to so many hits with both archaea and bacteria (Supplementary Table S4) that the influx of both archaeal and bacterial genes into the eukaryogenesis system had to be mediated by some highly efficient mechanism; and the similar prokaryotic-homology spectra for Gla and Trv (Fig. 5a) suggest that a majority of the prokaryotic genes in these two eukaryotic genomes entered the eukaryogenic lineage prior to the divergence between Gla and Trv. In fact, archaea have long relied on bacteria as a source of genetic diversity, and there was precedent of influx of bacterial genes being a determinant of archaeal phylogenies: a large number of bacterial genes entered into the methanogen that begot the haloarchael archaeons^36,37^. In view of this, an influx of beneficial prokaryotic genes into the archaeal-parent lineage likely began prior to the FECA stage and continued through LECA to the early eukaryotes as illustrated by the entry of bacterial rProt genes into Sce, Esi and Hsa (Supplementary Fig. S2).

**Figure 5.**
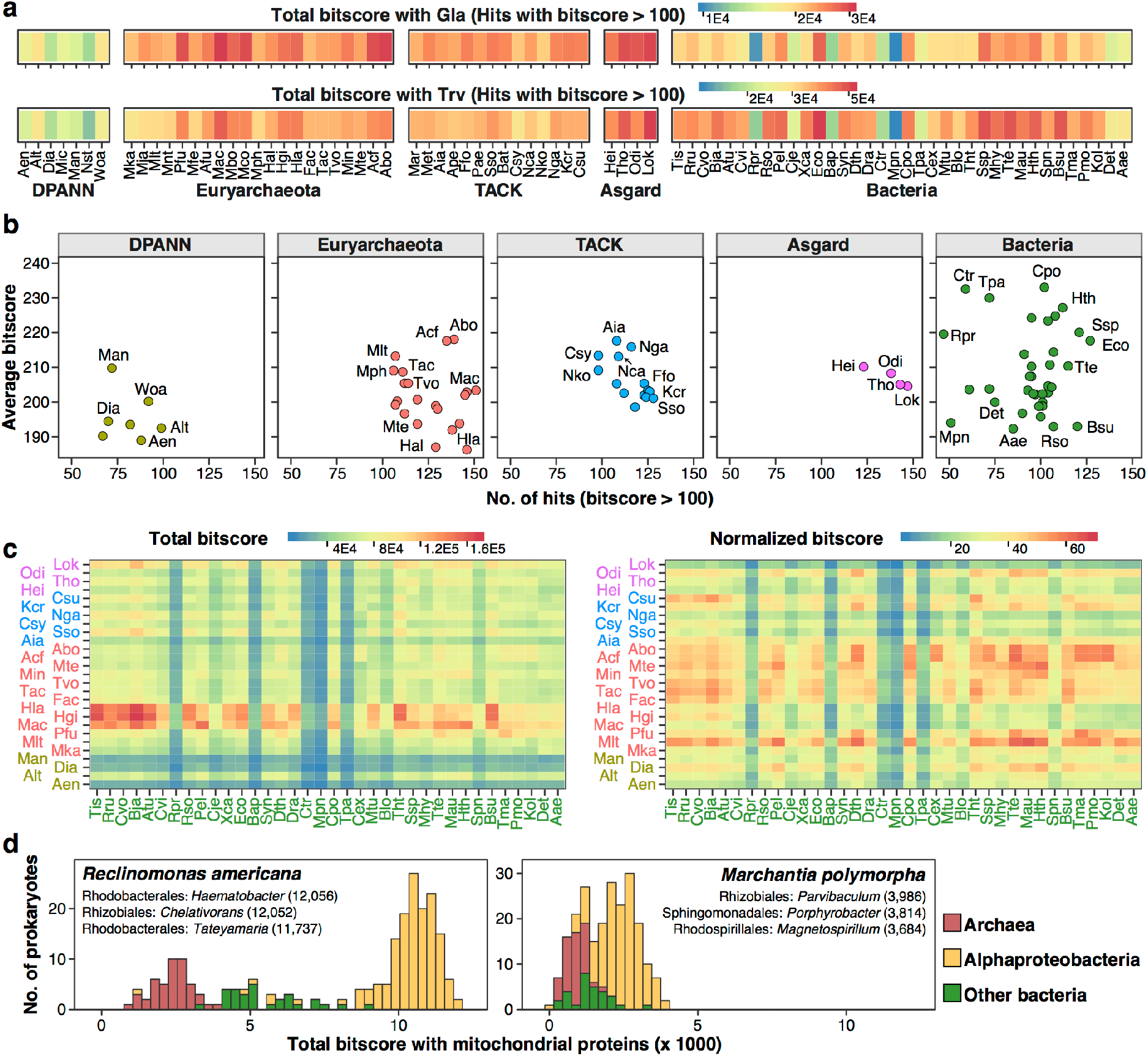
Interspecies protein homologies. (a) Total homology bitscores of Gla and Trv proteomes towards prokaryotic proteomes. (b) Relationships between the number of homologous hits (x-axis) and average bitscore per hit (y-axis) between prokaryotic and Gla proteomes. (c) Homologous hits between archaeal proteomes (y-axis) and bacterial proteomes (x-axis) with (left) or without (right) normalization based on the number of proteins in the archaeon. Complete heat map is given in Supplementary Fig. S4. (d) Bitscores of homologous hits of two eukaryotic species of mitochondrial gene-encoded proteins towards prokaryotes. Total bitscores in each instance are shown in parentheses.

Among 46 archael proteomes analyzed, the Abo and Acf proteomes displayed the highest average homology bitscores toward the eukaryotic proteomes of both Gla and Trv (Fig. 5b, Supplementary Fig. S3 and Supplementary Table S4), which suggests that these archaeons could be candidate archaeal-parents. Once a bacterial protein entered into the FECA-eukaryote lineage, its bacterial and eukaryotic versions became segregated locationwise, and evolved independently. The divergence between the two versions would thus increase with time, as in the case of functional paralogues such as VARS and IARS. Accordingly, the bitscores of Ctr and Tpa proteins were high toward Gla and Trv likely because they were taken up late by the eukaryogenic lineage, whereas the bitscores of Mpn and Aae were low likely because they were taken up early (Fig. 5b and Supplementary Fig. S3). Moreover, when the 46 archaeons were compared regarding their ability to import bacterial genes into their own genomes, several archaeons with relatively large proteomes, viz. Hla, Hgi and Mac (with 3,704 to 4,469 protein genes), as well as Lok (5,378 protein genes) and Pfu (2,053 protein genes), displayed distinctive homologies toward bacteria (Fig. 5c left panel). However, when the individual archaeal bitscores were normalized with respect to the protein gene numbers of the archaeons, the normalized bitscores of the small-proteome Abo-group of seven euryarchaeons comprising Abo, Acf, Mte, Tvo and Tac (each with <1,600 protein genes), Min and Fac (both with <1,900 protein genes), along with Odi, Csu, Pfu, Mlt and Dia, toward bacteria became more evident (Fig. 5c right panel, Supplementary Fig. S4). The results suggest that these latter archaeons had access to gene-import not only via HGT, which would be open to most or all archaeons, but also via some additional mechanism. Only the small parasitic bacteria Rpr, Bap, Ctr, Mpn and Tpa furnished relatively few genes to these archaeons.

Abo, the first cultivatable archaeon from the ‘Deep-sea hydrothermal vent euryarchaeota 2’ (DHVE2) group, and its facultatively anaerobic companion species Acf were closely related (84-87% 16S rRNA similarity) to the Thermoplasmatales archaeons Tac, Tvo Mte, Min and Fac^38,39^. The heterotrophic scavenger lifestyle of Abo and Acf based on their peptide fermentation, and also that of Tac^40^, suggests that the additional gene-import mechanism available to them consisted of *foodchain gene adoption* (FGA), which was proposed previously for the replacement of nuclear genes of early eukaryotes by genes in the bacteria they took as food^41^. To perform FGA, Abo would employ its array of thirty proteases to digest away the proteins of dead prokaryotes to prepare their naked DNA, import it through the Beveridge bridal S-layer of its cell surface (S-layer lattices are known to house regular channels of 2-6 nm diameter^42^) for implantation into its own genome. In using proteases to purify DNA for cloning, Abo predates by eons the same usage by modern genetic engineering. The S-layer of Abo is highly flexible and can be bent to form small blebbing vesicles with sharp curvature, indicating that the bonding forces between the S-layer subunits are unusually weak or transient. These vesicles can bud off and anneal with other cells^43^. *Pseudomonas aeruginosa* also releases comparable membrane vesicles that contain the pseudomonas quinoline signal for cell-cell communication and group behavior^44^; and *Sulfolobus islandicus* produces cell-derived S-layer coated spherical membrane vesicles of 90-180 nm diameter^45^. Importantly, such flexibility of the Abo cell surface could facilitate the formation of eukaryotic endomembrane and the prerequisite phagocytosis machinery for eukaryogenesis^31,46,47^. Furthermore, while all prokaryotic cells evolve on the basis of *nucleotidyl mutations* through the replacement, addition and subtraction of nucleotides, FGA gene uptake would enable the cells to evolve also on the basis of *gene-content mutations* through the replacement, addition and subtraction of genes, or gene clusters, expediting eukaryogenesis by orders of magnitude. In the example of Tac, it has succeeded in acquiring gene clusters from other organisms for rProts, NADH dehydrogenase, precorrin biosynthesis, flagellar proteins and a protein degradation pathway, amounting to 32% of its total open reading frames plausibly via FGA^40^. Besides, the blebbing vesicles of Abo and Acf could mediate gene exchanges between cells engaged in eukaryogenesis, thus further facilitating the process. Therefore, based on the highest archaeal BLASTP bitscores of Abo and Acf toward Gla pertaining to pyruvate phosphate dikinase (Fig. 4b), their highest average archaeal bitscores toward Gla and Trv proteomes (Fig. 5b and Supplementary Fig. S3) and highest archaeal normalized bitscores toward bacteria (Fig. 5c), blebbing membrane vesicles, and possession of nine out of ten glycolytic enzymes needed for metabolic cooperation with mitochondrial respiration, these Aciduliprofundum archaea represent exceptionally advantaged candidates for the archaeal-parent role. They even share with the deep branching eukaryote Gla, the rProts of which have remained more archaeal than those of Sce, Esi and Hsa, the scavenger life style. Between Acf and Abo, the catalase/peroxidase HP1-encoding facultatively anaerobic Acf might be more resistant to oxidative damage than Abo during merger with an alphaproteobacterium, and could hunt a wide range of ecological niches for beneficial food species. The bitscore 936 of Acf toward Gla with respect to pyruvate phosphate dikinase was also slightly higher than the 928 bitscore of Abo. The Asgard archaeons have been highly regarded as candidate archaeal-parents on the strength of their important ESP genes, which could render any one of them a valuable food species as well for the archaeal-parent. Moreover, the new cultivatable Asgard *Candidatus Prometheoarchaeum syntrophicum MK-D1*^48,49^ can degrade amino acids through syntrophy, and display wisp-like membrane protrusions indicative of flexible cell surface and possible FGA activity in keeping with the significant albeit modest bitscores between Lok and the bacterial species Bja and Tht, and between Odi and several bacterial species, in Fig. 5c left and right panels respectively.

Based on the BLASTP bitscores between the proteins of various prokaryotes and the mitochondrial gene-encoded proteins in four eukaryotes (Supplementary Table S5), the alphaproteobacteria *Haematobacter*, *Chelativorans* and *Tateyamaria* were closest to the lineage of the mitochondrion-parent (Fig. 5d).

In conclusion, in the present study, *Methanopyrus kandleri* was found to be the top-ranked organism in VARS-IARS homology among 5,398 species from all three biological domains, and therefore close to the root of life. Moreover, the high VARS-IARS homologies in the Primitive Archaea Cluster and in the Ancestral Bacteria Cluster delineated a pathway of descent of Bacteria from Archaea that diverged early from Archaea to form the Bacteria domain. The preeminent homology between the Gla rProts and archaeal rProts established that the prokaryote-parent of Eukarya that entered into genome merger with an alphaproteobacterial mitochondrion-parent was an archaeal-parent. The archaeal-parent was suggested to a scavenger archaeon such as the *Aciduliprofundum* archaea, capable of generating a chimeric eukaryote through large scale foodchain gene adoption. Notwithstanding such elaborate phylogenetic developments, the asterisked columns in Fig. 3, where all six aligned protein sequences showed the same Val or Leu residue despite the ease with which Val, Leu and Ile can be interchanged in evolution, represented a level of protein sequence conservation across two different proteins, three biological domains, and two giga year-plus time span that required the vertical genetic descent of Bacteria and Eukarya from an archaeal root of life.

**Supplementary Information** is available online.

## Author Contribution

J.T.W. and H.X. conceived the study; X.L. collected the data and performed computational analysis; and J.T.W., H.X. and X.L. wrote the paper. All authors read and approved the final manuscript.

## Acknowledgments

This study was supported by University Grants Council of Hong Kong SAR (ITS/113/15FP), and X.L. was recipient of a Hong Kong Government Ph. D. Fellowship.

## Author information

The authors declare that they have no competing interests. Correspondence and material requests should be addressed to T.F.W.

## Data availability

All data supporting the findings of this study are available within the paper and its Supplementary Information files.

## Methods

### Source of data and materials

The protein and SSU rRNA sequences analyzed in the present study were retrieved from the NCBI GenBank release 231 (ftp://ftp.ncbi.nlm.nih.gov/genomes/)^51,52^. For species without available SSU rRNA information in NCBI, quality checked SSU rRNA sequences were downloaded from the SILVA database release 132 (https://www.arb-silva.de/)^53^. For species with multiple SSU rRNA sequences, the one yielding the highest total bitscore (using BLASTN^54^ with ‘-word_size’ flag set to 4) with SSU rRNAs of other species from the same domain was employed for analysis. The alignment of the 83 SSU rRNA sequences used to build the SSU rRNA tree was given in Supplementary File S1. Eukaryotic mitochondrial gene-encoded protein sequences were retrieved from NCBI Protein database (https://www.ncbi.nlm.nih.gov/protein) by searching species name of the eukaryote of interest and setting the ‘Genetic compartments’ filter as ‘Mitochondrion’.

### Estimation of nuclear or mitochondrial proteome homology

When the proteomes of two species were compared using BLASTP^54^ (with ‘-evalue’ flag setting to 0.05), query and subject sequences that were the only best match of each other, viz. query ***n*** has the highest bitscore with subject ***m***, and subject ***m*** has the highest bitscore with query ***n*** at the same time, were considered in calculating proteome similarity.

### Estimation of protein family homology

Protein sequence similarity was estimated by the maximum bitscore of each pair of sequences yielded by BLASTP^54^ with all default parameters. To estimate ribosomal protein similarities among eukaryotes and prokaryotes, all seed sequences of 80 ribosomal protein families (Supplementary Table S6) retrieved from Pfam database^55^ were blasted (with ‘-evalue’ flag set to 0.05) against the proteomes of 21 eukaryotes, 46 archaea and 36 bacteria respectively. For every ribosomal protein family, only one protein sequence of each species yielding the highest bitscore with the seed sequences were selected for further analysis. To remove false-positive sequences, the selected sequences were submitted to NCBI Batch CD-search Tool^56^ to search against the Pfam database, and sequences that failed to map to the same ribosomal protein family were removed. Finally, the eukaryotic ribosomal protein sequences were blasted to all the prokaryotic ribosomal protein sequences, and the maximum bitscores are shown in Fig. 4a.

To determine non-rProt families with sequence homology between Gla and prokaryotes, the Gla protein sequence in each instance was blasted against each of the 82 prokaryotic proteomes, and the best matches were submitted to NCBI Batch CD-search Tool to search against the Pfam database. Only cases where both query and subject sequences belonged to the same protein family were kept, and 162 of these cases were shown in Supplementary Table S2, and in part in Fig. 4c.

