## Supplementary Information for "Descent of Bacteria and Eukarya from an archaeal root of life"

### Supplementary Figures

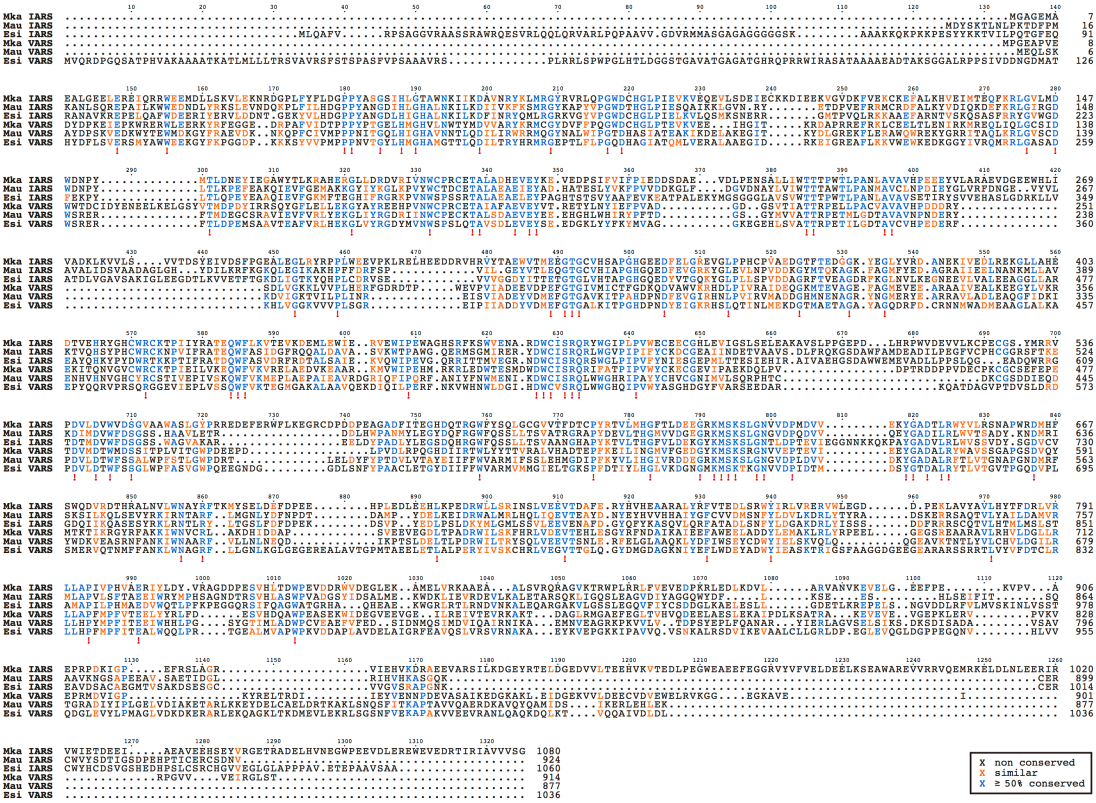

**Supplementary Figure S1. Amino acid sequence alignment of VARS and IARS of Mka, Mau and Esi.** The six sequences were aligned using ClustalOmega. Similar amino acid positions are colored in orange, and ≥ 50% conserved ones in blue. Exclamation marks indicate the positions displaying conserved amino acid in all six sequences.

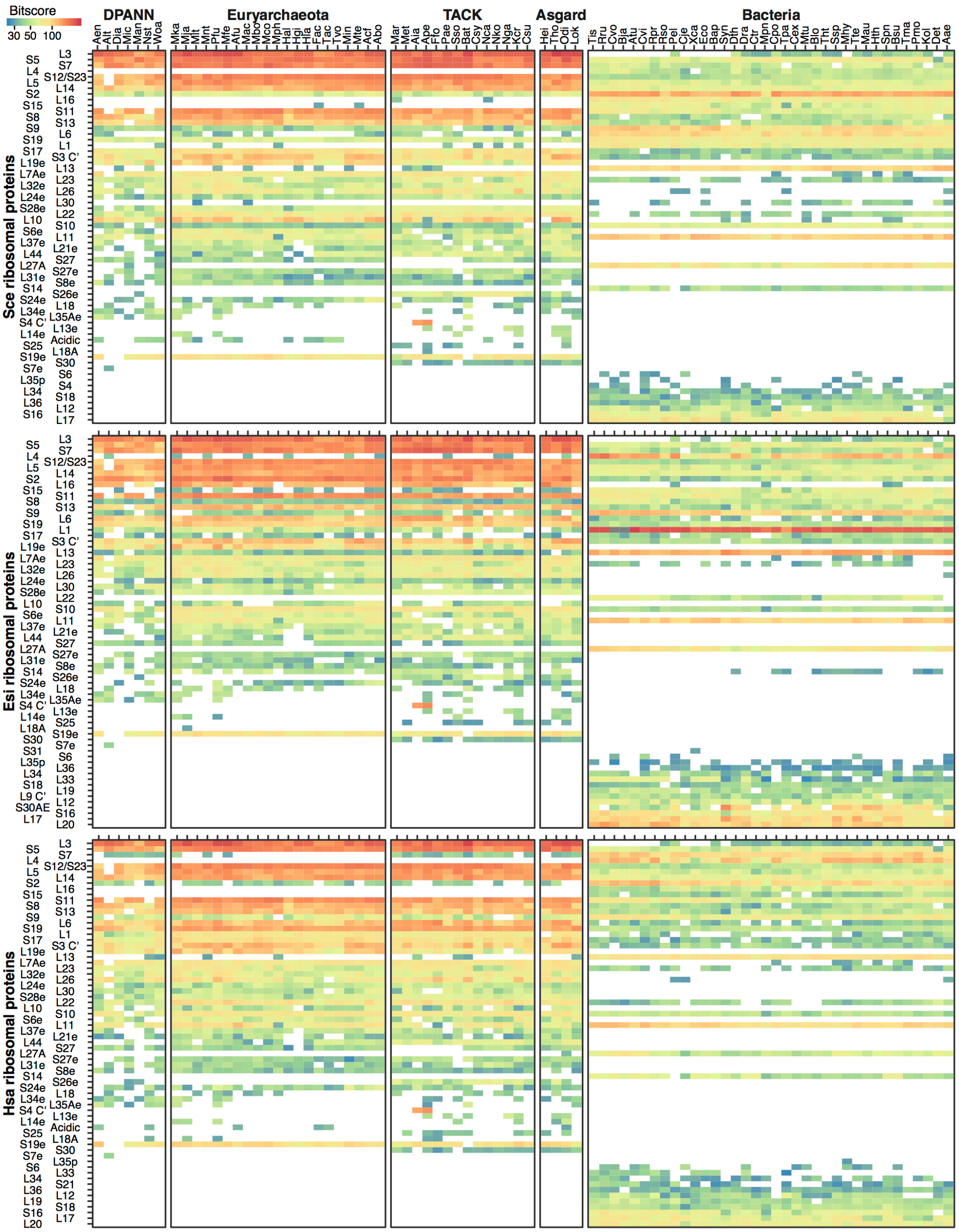

**Supplementary Figure S2. Sequence homologies of rProt families between eukaryotic and prokaryotic species yielding homologous hits with E-value < 0.05 in BLASTP.** Homologies exhibited by rProt families found in Saccharomyces cerevisiae (Sce), Ectocarpus siliculosus (Esi) and Homo sapiens (Hsa) toward 46 archaeal species of the DPANN, Euryarchaeota, TACK and Asgard superkindoms, as well as 36 bacterial species were determined. The pairwise maximum bitscores for each rProt are expressed by the thermal scale with blue for low and red for high homology.

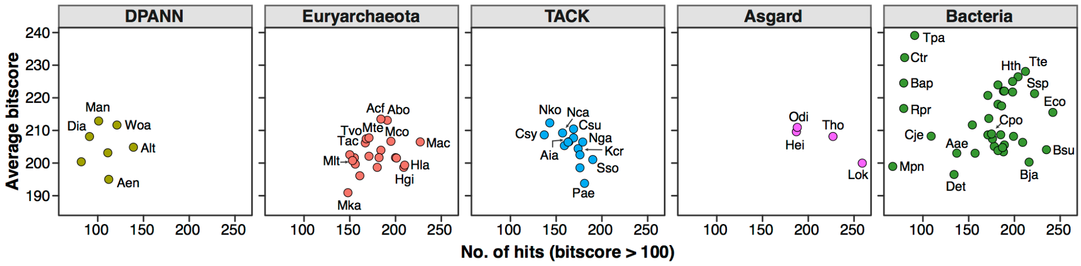

**Supplementary Figure S3. Relationship between the number of homologous hits (x-axis) and average bitscore (y-axis) of each prokaryote toward *Trichomonas vaginalis*.**

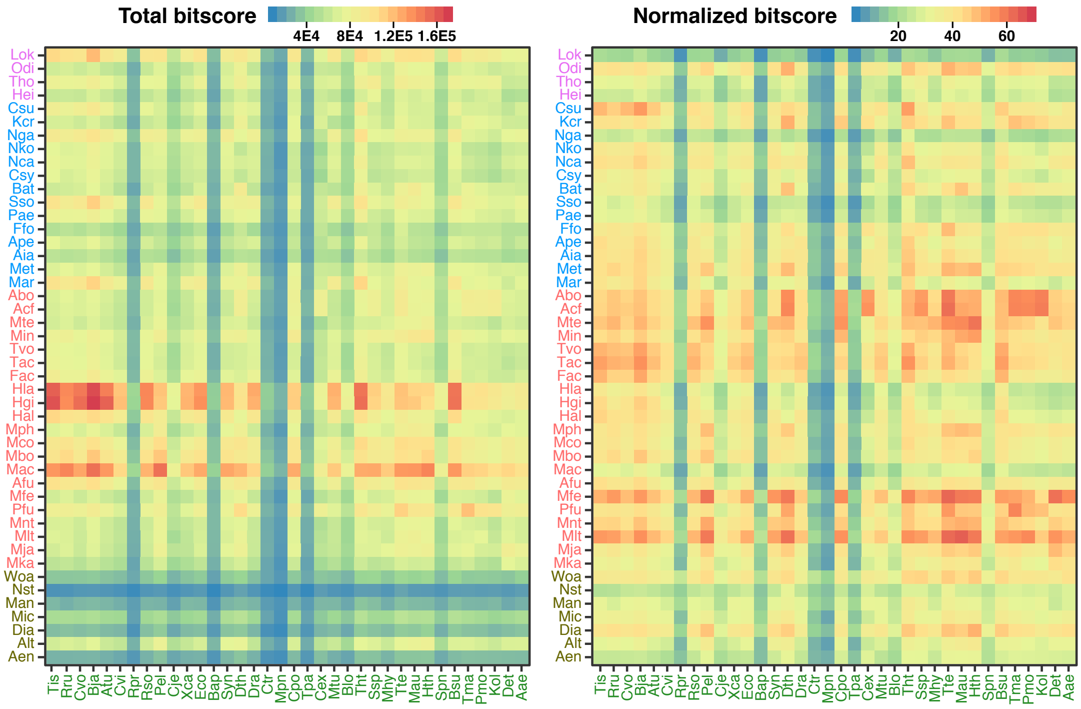

**Supplementary Figure S4. Proteome homology between archaeal (y-axis) and bacterial (x-axis) proteomes.** Left panel: total bitscore of homologous proteins. Right panel: normalized bitscore calculated as total bitscore divided by the number of proteins in the query archaeal species.

**Captions for Supplementary Tables**

**Supplementary Table S1.** VARS-IARS homology of 5,398 species.

**Supplementary Table S2.** Protein families of Gla with homologous sequences in ≤ 10/82 prokaryotes.

**Supplementary Table S3.** Count of Gla ESPs in prokaryotes.

**Supplementary Table S4.1** Homologous hits with ≥ 100 bitscores between Gla and prokaryotic proteomes.

**Supplementary Table S4.2** Homologous hits with ≥ 100 bitscores between Trv and prokaryotic proteomes.

**Supplementary Table S5.** Homology between prokaryotic proteins and eukaryotic mitochondrial gene-encoded proteins.

**Supplementary Table S6.** Homology of eighty eukaryotic ribosomal protein families toward archaea and bacteria.

**Description of Supplementary File S1**

**Supplementary File S1.** ClustalOmega alignment of the 83 SSU rRNA sequences used to build the SSU rRNA tree. The three-lettered species name was added at the end of each sequence name.
