## Supplementary File S1 for "Descent of Bacteria and Eukarya from an archaeal root of life"

>lcl|AE000657.1_rrna_15 [locus_tag=aq_r03] [product=16S ribosomal RNA] [location=complement(571199..572785)] [gbkey=rRNA] Aae
--------------------------------------------------------------------------------
--------------------------------------------------------------------------------
--------------------------------------------------------------------------------
--------------------------------------------------------------------------------
--------------------------------------------------------------------------------
--------------------------------------------------------------------------------
--------------------------------------------------------------------------------
--------------------------------------------------------------------------------
-------------------------------------------TTTCC---CTGAAGAGTTTGATCCTGGCTCAG-CGCG
AACGCTGGCGGCGTGCCTAACACATGCAAGTCGTGCGCAGGGTCGGCCCCT--------T------T-TGGGGCCGGCCG
CTGAGCGGCAAACGGGTGAGTAACA-CGTGGGTAACC-------------------------------------------
-----TACCCCCAGG--AGGGGGATAACCCCGGGAAA-CCGGGGCTAATACCCCAT-AATGCCACCCGCCACTAAGGCGC
GGTG---G-------------------------------CCAAAGGGGGCCTCTGGCCTTCGGCCA---CGCTCCCGCCT
GGGGATGGGCCCGCGGCCCATCAGGTAGTTGGTGGGGT-AACGGCCCACCAAGCCTATGACGGGTAGCCGGCCTGAGAGG
GTGGCCGGCCACAGCGGGACTGAGACACGGCCCGCACCCCTACGGG--GGGCAGCAGTGGGGAATCGTGGGCAATGGGCG
AAAGCCTGAC---CCCG---C-----------GACGCCGCGTGGGGGATGAAGCCCTGCGG-GGTGTAAACCCCTGTCGG
GGGGGACGATGCGGACACGGG-TTAATAGCCCGTGTCCGTGACGGTACCCCCAGAGGAAGGGACGGCTAACTACGTGCCA
GCAGCCGCGGTAATACGTAGGTCCCAAGCGTTGCGCGAAGTCACTGGGCGTAAAGCGTCCGCAGCCGGTCGGGTAAGCGG
GATGTCAAAGCCCACGGCTCAACCGTGG-ACCG----------------------------------GCATCCCGAACT-
-----------------------------------GCCCGACTTGAGGCACGCCCGGGCAGGCGGAATTCCCGGGGTAGC
GGTGAAATGCGTAGATCTCGGGAGGA------------------------------------------------------
--------------------------------------------------------------------------------
-----------------------------------ACACCGAAGGGGAAGCCAGCCTGCTGGGGCTGTCCTGACGGTCAG
GGACGAAAGCCGGGGGAGCAAACCGGATTAGATACCCGGGTAGTCCCGGCCGTAAAC-----------------------
--------------------------------------------------------------------------------
---------CATGGGCGCTAGGGCTTGTCCC---T------C---TGGGGCAGGCTCGCAGCTAACGCGTTAAGCGCCCC
GCCTGGGGAGTACGGGCGCAAGCCTGAAACTCAA-AGGAATTGGCGGGGGCCCGCACA-ACCGGTGGAGCG-TCTGGTTC
AATTCGATGCTAACCGAAGAACCTTACCCGGGCTTGACATGCCGGGGAGACTCCGCGAAAGCGGAGTT------------
-----------------------GTGGAAGTCCCTAGGACTTTCCCCCGGCACAGGTGGTGCATGGCCGTCGTCAGCTCG
TGTCG-TGAGATG--TTGGGTTAAGTCCCGCAAC----------------------------------------------
--------------------------------------------------------------------------------
----------G------AGCGCAACCCCTGCCCCCAGTTGCTAC---CCCGTT--TG----------GGGAGCACTCTGG
GGGGACCGC----CGGCGATA-AGCCGGAGGAAGGGGGGGATGACGTCAGGTCA-GTATGC-CCTTTATGCCCGG-GGCC
ACACAGGCGCTACAGTGGCCGGGACAATGGGATG-CGACCCCG-------------------------------------
----------------------------------------TAAGGGGGAGCTA--ATCCCTAAACCCGGTCATGGTGCGG
ATTGGGGGCTGAAACTCGCCCCCATGAAG--CCGGAATCGGTAGTAACGGGGTATCAGCGATGTCCCCGTGAATACGTTC
TCGGGCCTTGCACACACCGCCCGTCACGCCACGGAAGTCGGTCCGGCCGGAAGTCCCCGAGCTAACCGGCTC--------
-------------CTTCGGGAGCCGGAGGCAGGGGCCGATGGCCGGGCCGGCGACTGGGGCGAAGTCGTAACAAGGTAGC
CGTAGGGGAACCTGCGGCTGGATCACCTCCTTTA----------------------------------------------
------------------------------------------------------
>lcl|CP002543.1_rrna_41 [locus_tag=Dester_R0042] [product=16S ribosomal RNA] [location=complement(1425680..1427250)] [gbkey=rRNA] Det
--------------------------------------------------------------------------------
--------------------------------------------------------------------------------
--------------------------------------------------------------------------------
--------------------------------------------------------------------------------
--------------------------------------------------------------------------------
--------------------------------------------------------------------------------
--------------------------------------------------------------------------------
--------------------------------------------------------------------------------
-------------------------------------------------------AGAGTTTGATCCTGGCTCAG-CGCG
AACGCTGTCGGCGTGCCTAACACATGCAAGTCGTGGGGCAGCAGGCGGGTTCCTT---TT------G-GAACCCGTGCTG
GCGACCGGCGAACGGGTGAGTAACG-CGTGAGCAACC-------------------------------------------
-----TACCCCGAGG--AGGGGGATAACCCGGGGAAA-CTCGGGCTAATACCCCAT-AAGCGCACGACCTGCATGGGCTG
TGTG---A-------------------------------AAAGGGT-----GGCCTCTGCTTTGCA---AGCTGCCGCCT
CGGGATGGGCTCGCGTCCCATCAGGTAGTTGGTGAGGT-AACGGCTCACCAAGCCTACGACGGGTAGCCGGCCTGAGAGG
GTGGTCGGCCACACCGGGACTGAGACACGGCCCGGACTCCTACGGG--AGGCAGCAGTGGGGAATCTTGGGCAATGGGCG
CAAGCCTGAC---CCAG---C-----------GACGCCGAGTGGAGGAAGAAGCCCTTCGG-GGTGTAAACTCCTGTCGC
CGGGGACGATGCGGGTGTGGG-TTAATAGCTCACACTCGTGACGGTACCCGGAGAGGAAGGGACGGCTAACTACGTGCCA
GCAGCCGCGGTAATACGTAGGTCCCGAGCGTTGCGCGAAATTACTGGGCGTAAAGAGCGCGTAGGCGGGCCAGTAAGTCA
GAGGTGAAAGCCCCCGGCTCAACCGGGG-AATT----------------------------------GCCTCTGATACT-
-----------------------------------GCTGGTCTAGGGTCCGGAAGAGGCTGGCGGAATTCCCGGTGTAGC
GGTGAAATGCGTAGATATCGGGAGGA------------------------------------------------------
--------------------------------------------------------------------------------
-----------------------------------ACACCGGTGGCGAAGGCGGCCAGCCAGGACGGTACCGACGCTGAG
GCGCGAAAGCGTGGGGAGCAAACCGGATTAGATACCCGGGTAGTCCACGCTGTAAAC-----------------------
--------------------------------------------------------------------------------
---------GATGGGTGCTGGGCGTTGGCAC---C---GATA---GGTGTCAGTGCCGAAGCTAACGCGTTAAGCACCCC
GCCTGGGGAGTACGGCCGCAAGGCTGAAACTCAA-AGGAATAGACGGGGGCCCGCACA-AGCGGTGGAGCA-TGTGGTTT
AATTCAATGCTACGCGAAGAACCTTACCAGGGCTTGACATGCGGTGTGTCGTAGCCGAAAGGCTACTA------------
-----------------------GCTACC-CTTTGTGGGGTAGCGCGCCGCACAGGTGGTACATGGCCGTCGTCAGCTCG
TGCCG-TGAGGTG--TTGGGTTAAGTCCCGCAAC----------------------------------------------
--------------------------------------------------------------------------------
----------G------AGCGCAACCCCTGTCCCTAGTTGCCAG---CGGGTA--AA---------GCCGGGCACACTAG
GGAGACTGC----CGGCGATA-AGCCGGAGGAAGGAGGGGATGACGTCAGGTCA-GTATGC-CCCTTACGCCCTG-GGCT
ACACACGTGCTACAATGGTCGGGACAGAGGGAAG-CGAAGCCG-------------------------------------
----------------------------------------CGAGGTGGAGCAAAACCCACCAAACCCGGTCCCAGTTCGG
ATTGCAGGCTGCAACTCGCCTGCATGAAG--CTGGAATCGCTAGTAACGGCCGGTCAGCTACACGGCCGTGAATACGTTC
CCGGGCCTTGTACTCACCGCCCGTCACACCCCGAAAGCTGGCTCTACCAGAAGTCCCCGGGCTAACCCTCTT--------
-------------ATGAGGG------AGGCAGGGGCCGAAGGTAGGGTTGGTGATTGGGGTGAAGTCGTAACAAGGTAGC
CGTAGGAGAACCTGCGGCTGGATCACCT----------------------------------------------------
------------------------------------------------------
>lcl|CP001146.1_rrna_2 [gene=rrs_1] [locus_tag=DICTH_0617] [product=16S ribosomal RNA] [location=624121..625660] [gbkey=rRNA] Dth
--------------------------------------------------------------------------------

--------------------------------------------------------------------------------
--------------------------------------------------------------------------------
--------------------------------------------------------------------------------
--------------------------------------------------------------------------------
--------------------------------------------------------------------------------
--------------------------------------------------------------------------------
--------------------------------------------------------------------------------
-------------------------------------------------------AGGGTTTGATCCTGGCTCAG-GACG
AACGCTGGCGGCGTGCCTAAGGCATGCAAGTCGAGCGGTAGCTCCTATTGGTTT-----A------T-GCCGATAGGAGT
GAGAGCGGCGCACGGGTGAGTAACA-CGTAGGTAACC-------------------------------------------
-----TACCCCAGAG--AGGGGGATAACACCTCGAAA-GGGGTGCTAAACCCCCAT-ATACTTACCGAGCGATATGCTCA
GGTA---A-------------------------------GGAAAGAG--TATAGAGGGGTAACCCT---CTATGCTGCTC
TGGGATGGGCCTGCGGCCTATCAGCTAGTTGGTGGGGT-AATGGCCCACCAAGGCTAAGACGGGTAGCCGGCCTGAGAGG
GTGGTCGGCCACACTGGGACTGAGACACGGCCCAGACTCCTACGGG--AGGCAGCAGCCGGGAATCTTCCGCAATGGGGG
AAACCCTGAC---GGAG---C-----------GACGCCGCGTGAGGGAAGAAGCCCTTCGG-GGTGTAAACCTCTTTTCT
CGGGGAAGAATA--------------------------CTGACGGTACCCGAGGAAAAAGCCCCGGCTAACTACGTGCCA
GCAGCCGCGGTAAGACGTAGGGGGCGAGCGTTGTCCGGATTTACTGGGCGTAAAGGGCGTGTAGGCGGCTTAGCAAGTCA
GATGTGGAAGCCCTGAGCTCAACTCAGG-GAGG----------------------------------TCATCTGATACT-
-----------------------------------GCTAAGCTAGAGGGCAGGAGAGGAGAGCGGAACTTCCGGTGTAGC
GGTGAAATGCGTAGATATCGGAAGGA------------------------------------------------------
--------------------------------------------------------------------------------
-----------------------------------ACGCCGGTGGCGAAGGCGGCTCTCTGGACTGACCCTGACGCTGAG
GCGCGAAAGCTAGGGGAGCGAACGGGATTAGATACCCCGGTAGTCCTAGCCGTAAAC-----------------------
--------------------------------------------------------------------------------
---------GATGGGCACTAGGTGTGGGGAG-------TTAG---ACTCTCCGTGCTGCAGCTAACGCGATAAGTGCCCC
GCCTGGGGAGTACGGCCGCAAGGCTAAAACTCAA-AGGAATTGACGGGGGCCCGCACA-AGCGGTGGAGCG-CGTGGTTT
AATTCGATGCTAAGCGAAGAACCTTACCAGGGCTTGACATGCAGGCTGTAAGCTCACTCGAAAGGGTGAGT---------
-----------------GCCAAATAGGGGGTTTCCCCTATTTGGAGCCTGCACAGGTGGTGCATGGCTGTCGTCAGCTCG
TGTCG-TGAGATG--TTGGGTTAAGTCCCGCAAC----------------------------------------------
--------------------------------------------------------------------------------
----------G------AGCGCAACCCCTGCCCTTAGTTGCCAG---CGGGTA--AA---------GCCGGGCACTCTAA
GGGGACTGC----CGGCGAAG-AGCCGGAGGAAGGTGGGGATGACGTCAAGTCA-GTATGC-CCCTTATGCCCTG-GGCT
ACACACGCGCTACAATGGGTGGTACAGAGGGGAG-CGAAGCCG-------------------------------------
----------------------------------------CGAGGCGGAGCGAATC-C-CTAAAGCCACCCCCAGTTCAG
ATCGCAGGCTGCAACTCGCCTGCGTGAAG--GCGGAATCGCTAGTAACCGCAGATCAGCCACGCTGCGGTGAATACGTTC
TCGGGCCTTGTACACACCGCCCGTCACACCACGAGAGTCCGCAACACCCGAAGTCAGGCGAAGA----------------
------------------------------GCCTGCCGAAGGTGGGGCGGATGATTGGGGTGAAGTCGTAACAAGGTAGC
CGTACCGGAAGGTGCGGCTGGATCACCTCCTTT-----------------------------------------------
------------------------------------------------------
>lcl|CP000879.1_rrna_52 [locus_tag=Pmob_R0052] [product=16S ribosomal RNA] [location=2041444..2042943]
[gbkey=rRNA] Pmo
--------------------------------------------------------------------------------
--------------------------------------------------------------------------------
--------------------------------------------------------------------------------
--------------------------------------------------------------------------------
--------------------------------------------------------------------------------
--------------------------------------------------------------------------------
--------------------------------------------------------------------------------
--------------------------------------------------------------------------------
-------------------------------------------------------AGGGTTTGATCCTGGCTCAG-GGTT
AACGCTGGCGGCGTGCGTAACACATGCAAGTCGGACGGGTTATAGATCTTCGG-----------------AGATA---TA
ACCAGTGGCGAACGGGTGAGTAAAA-GGTAGGGACC--------------------------------------------
-----TGCCCTAAGG--ACAGAGATAGCTACTGGAAA-CGGTAGGTAAACTCTGAT-AAGCCCGAGAGGG----------
-----------------------------------------------------------GAAAGTG---GTAGACAGCCT
TAGGATGGACCTACTATCCATCAGGTAGTTGGTGAGGT-AAAGGCTTACCAAGCCGATGACGGATAACCGGTGTGAGAGC
ATGGACGGTCACAAGGGCACTGAGACACGGGCCCTACTCCTACGGG--AGGCAGCAGTGGGGAATCTTGGACAATGGGCG
AAAGCCTGAT---CCAG---C-----------GACGCCGCGTGAAGGAAGAAGTCCCTCGG-GATGTAAACTTCTGAACT
AATCGAATAAAAGGGTAGTGGA------CACACTAC-AGAAGAAGGTAGGTTAGGAAAAGTCCCGGCTAACTACGTGCCA
GCAGCCGCGGTAAGACGTAGGGGGCGAGCGTTGCCCGGAATTACTGGGTGTAAAGGGGACGTAGGCGGGTGATCAAGTCA
TCTGTGAAAAG--ATTGCTCAACGATCG-GCTT----------------------------------GCGGATGAAACT-
-----------------------------------GATCATCTTGGGCGTAGCAGAGGTAGACGGAATTACCTGAGTAGG
GGTGAAATCCGCAGATACAGGTAAGA------------------------------------------------------
--------------------------------------------------------------------------------
-----------------------------------ACGCCGGTGAAGAAGTTGGTCTACTGGGCTACAGCTGACGCTGAG
GTCCGAAAGCCAGGGGAGCAAACCGGATTAGATACCCGGGTAGTCCTGGCCCTAAAC-----------------------
--------------------------------------------------------------------------------
---------GATGCTCACTAGGTGTAGGGAG---CG--AAAG---ACTCTCTGTGCTGAAGCGAACGCGCTAAGTGAGCC
ACCTGGGGAGTACGTCCGCAAGGATGAAACTCAA-AGGAATTGACGGGGGTCCGCACA-AGCGGTGGAGCA-TGTGGTTT
AATTCGAAGCTAACCGAAGAACCTTACCAGGGATTGACATGTAACTGAAGGTAGAGAAAT--------------------
---------------------CTACTGGCCTACCGTAAGGTAGGAGGTTACACAGGTGGTGCACGGTCGTCGTCAGCTCG
TGCCG-TGAGGTG--TTGGGTTAAGTCCCACAAC----------------------------------------------
--------------------------------------------------------------------------------
----------G------AGCGCAACCCCTGCAATTAGTTACCAG---CAAGTA--AA---------GTTGGGGACACTAA
TTGGACAGC----CGCCGACG-AGGCGGAGGAAGGAGGGGATGACGTCAGATAA-GCGTGC-CCCTTATACTCTG-GGCG
ACACACGTGCTACAATGGGGAGGACAAAGGGAAG-CGAAGCCG-------------------------------------
----------------------------------------GAAGGTGGAGCGGATC-CGGAAAAACTCTCCGTAATATGG
ATTGTAGGCTGAAACCCGCCTACATGAAG--CTGGAATCGCTAGTAATCGCAGGTCAGCCAAACTGCGGTGAATACGTTC
CCGGGCCTTGTACACACCGCCCGTCACGCCACCCGAGTTGGGAACACCTGAAGGCAGTACGGTA----------------
----------------------------GGTACTGTTGAAGGTGGGCTTAGCGAGGGGGGCGAAGTCGTAACAAGGTAGG
TGTACCGGAAGGTGCGCCTGGATCACCT----------------------------------------------------
------------------------------------------------------
>lcl|CP001634.1_rrna_40 [locus_tag=Kole_R0040] [product=16S ribosomal RNA] [location=1770534..1772060]
[gbkey=rRNA] Kol
--------------------------------------------------------------------------------
--------------------------------------------------------------------------------
--------------------------------------------------------------------------------
--------------------------------------------------------------------------------

```
--------------------------------------------------------------------------------
--------------------------------------------------------------------------------
--------------------------------------------------------------------------------
--------------------------------------------------------------------------------
-------------------------------------------------------AGGGTTTGATCCTGGCTCAG-GGTG
AACGCTGGCGGCGTGCCTGACACATGCAAGTCGTACGGGCGGACCCCCTTCGG-----------------GGGGCGAG--
-CCAGTGGCGAACGGGTGAGTAACA-CGTGGGAACA--------------------------------------------
-----TATCCGCTGG--TGGGGATAACTACCCGAAA-GGGTAGCTAATACCGCAT-AAGCCCGA----------------
--------------------------------------------------AAGGGGAAAGGTGCAA---CAGCATCGCCA
GTGGAGTGGCCCGCGGCCCATCAGCTAGTTGGTGGGGT-AAGAGCCCACCAAGGCGACGACGGGTAGCCGGCGTGAGAGC
GCGACCGGCCACAAGGACACTGAGACACGGGTCCTACTCCTACGGG--AGGCAGCAGTGCGGAATATTGGACAATGGGCG
AAAGCCTGAT---CCAG---C-----------GACGCCGCGTGCGGGACGAAGGCCTTCGG-GTCGTAAACCGCTGTGGT
AAGGGACGAATAGCCGGTGGAGTGGAAAGCCACCGGAGGAGACGGTACCTTACTAGGAAGCCCCGGCTAACTACGTGCCA
GCAGCCGCGGTAATACGTAGGGGGCGAGCGTTACCCGGAATCACTGGGCGTAAAGGGAGCGTAGGCGGCCCAGCAAGTCG
ACTGTGAAAAACCGGAGCTCAACTTCGG-TCCT----------------------------------GCGGTCGAAACT-
-----------------------------------GCTGGGCTTGAGGACAGTAGAGGGAGACGGAACTGCCGGTGTAGG
GGTAAAATCCTTAGATATCGGCAGGA------------------------------------------------------
--------------------------------------------------------------------------------
-----------------------------------ACGCCGGTGGCGAAGGCGGTCTCCTGGGCTGGACCTGACGCTGAG
GCTCGAAAGCCAGGGGAGCGAACGGGATTAGATACCCCGGTAGTCCTGGCCCTAAAC-----------------------
--------------------------------------------------------------------------------
---------GATGCTCACTAGGTGTGGGGTG---CG--TAGA---GCACTCCGTGCTGAAGCAAACGCGATAAGTGAGCC
GCCTGGGGAGTACGCCCGCAAGGGTGAAACTCAA-AGGAATTGACGGGGGCCCGCACA-AGCGGTGGAGCG-TGTGGTTT
AATTCGATGCTAAGCGAAGGACCTTACCAGGGCTTGACATGCTGGTGGTACTGAACCGAAAGGGGAAG------------
---------------------GACCC-----ACCGTTAGGTGGGAGCCAGCACAGGTGGTGCACGGCCGTCGTCAGCTCG
TGCCG-TGAGGTG--TTGGGTTAAGTCCCGCAAC----------------------------------------------
--------------------------------------------------------------------------------
----------G------AGCGCAACCCCTGCCCCTAGTTGCCAG---CGGTTA--G----------GCCGGGCACTCTAG
GGGGACCGC----CGCCGACG-AGGCGGAGGAAGGGGGGGATGACGTCAGGTA--TCGTGC-CCCTTATGTCCTG-GGCG
ACACACGCGCTACAATGGGACGGACAGCGGGGAG-CGAGGCAG-------------------------------------
----------------------------------------CGATGCTGAGCAAATC-CCTGAAACCGTTCCTCAGTTCGG
ATTGCAGGCTGCAACCCGCCTGCATGAAG--CCGGAATCGCTAGTAATCGCGGATCAGCCATGCCGCGGTGAATACGTTC
CCGGGCCTTGTACACACCGCCCGTCAAGCCACCCGAGTTGGCAACACCTGAAGACGGGTGTCCCAACCGAAA--------
-------------G--------GAGG--GAGCCCGTTTAAGGTGGGGCTGGCGAGGGGGGCTAAGTCGTAACAAGGTAGC
CCTACCGGAAGGTGGGGCTGGATCACCT----------------------------------------------------
------------------------------------------------------
>lcl|AE000512.1_rrna_2 [locus_tag=TM_r01] [product=16S ribosomal RNA] [location=188968..190526] [gbkey=rRNA] Tma
--------------------------------------------------------------------------------
--------------------------------------------------------------------------------
--------------------------------------------------------------------------------
--------------------------------------------------------------------------------
--------------------------------------------------------------------------------
--------------------------------------------------------------------------------
--------------------------------------------------------------------------------
--------------------------------------------------------------------------------
---------------------------------------------TAT---ATGGAGGGTTTGATCCTGGCTCAG-GGTG
AACGCTGGCGGCGTGCCTAACACATGCAAGTCGAGCGGGGAAACTCCCTTCG------------------GGGAGGAGTA
CCCAGCGGCGGACGGGTGAGTAACA-CGTGGGTAACC-------------------------------------------
-----TGCCCTCCGG--AGGGGGATAACCAGGGGAAA-CCCTGGTTAATACCCCAT-ACGCTCCATCAACGCAA------
-------------------------------------------GTTG--GTGGAGGAAAGGGGCGT---TTGCCCCGCCG
GAGGAGGGGCCCGCGGCCCATCAGGTAGTTGGTGGGGT-AACGGCCCACCAAGCCGACGACGGGTAGCCGGCCTGAGAGG
GTGGTCGGCCACAGGGGCACTGAGACACGGGCCCCAC-TCTACGGG--AGGCAGCAGTGGGGAATCTTGGACAATGGGGG
AAACCCTGAT---CCAG---C-----------GACGCCGCGTGCGGGACGAAGCCTTCGGG-GTG-TAAACCGCTGTGGC
GGGGGAAGAATA--AGGTAGGGAGGAAATGCCCTACCGATGACGGTACCCCGCTAGAAAGCCCCGGCTAACTACGTGCCA
GCAGCCGCGGTAATACGTAGGGGGCAAGCGTTACCCGGATTTACTGGGCGTAAAGGGGGCGTAGGCGGCCTGGTGTGTCG
GATGTGAAATCCCACGGCTCAACCGTGG-GGCT----------------------------------GCATCCGAAACT-
-----------------------------------ACCAGGCTTGGGGGCGGTAGAGGGAGACGGAACTGCCGGTGTAGG
GGTGAAATCCGTAGATATCGGCAGGA------------------------------------------------------
--------------------------------------------------------------------------------
-----------------------------------ACGCCGGTGGGGAAGCCGGTCTCCTGGGCCGACCCCGACGCTGAG
GCCCGAAAGCCAGGGGAGCAAACCGGATTAGATACCCGGGTAGTCCTGGCTGTAAAC-----------------------
--------------------------------------------------------------------------------
---------GATGCCCACTAGGTGTGGGGGG-------GTAA---TCCCTCCGTGCTGAAGCTAACGCGTTAAGTGGGCC
GCCTGGGGAGTACGCCCGCAAGGGTGAAACTCAA-AGGAATTGACGGGGGCCCGCACA-AGCGGTGGAGCG-TGTGGTTT
AATTGGATGCTAAGCCAAGAACCTTACCAGGGCTTGACATGCCGGTGGTACCTCCCCGAAAGGGGTAG------------
---------------------GGACCCAGTCCTTCGGGACTGGGAGCCGGCACAGGTGGTGCACGGCCGTCGTCAGCTCG
TGCCG-TGAGGTG--TTGGGTTAAGTCCCGCAAC----------------------------------------------
--------------------------------------------------------------------------------
----------G------AGCGCAACCCCTGCCCCTAGTTGCCAG---CGGTTC--G----------GCCGGGCACTCTAG
GGGGACTGC----CGGCGACG-AGCCGGAGGAAGGAGGGGATGACGTCAGGTAC-TCGTGC-CCCTTATGCCCTG-GGCG
ACACACGCGCTACAATGGGCGGTACAATGGGTTG-CGACCCCG-------------------------------------
----------------------------------------CGAGGGGGAGCCAATC-C-CCAAAGCCGCCCTCAGTTCGG
ATCGCAGGCTGCAACCCGCCTGCGTGAAG--CCGGAATCGCTAGTAATCGCGGATCAGCCATGCCGCGGTGAATACGTTC
CCGGGCCTTGTACACACCGCCCGTCACGCCACCCGAGTCGGGGGCTCCCGAAGACACCTACCCCAACCCGAA--------
-------------A--------GGGAGGGGGGGTGTCGAGGGAGAACCTGGCGAGGGGGGCGAAGTCGTAACAAGGTAGC
CGTACCGGAAGGTGCGGCTGGATCACCTCCTTTC----------------------------------------------
------------------------------------------------------
>lcl|U00089.2_rrna_4 [locus_tag=MPN_r01] [product=16S ribosomal RNA] [location=118312..119824] [gbkey=rRNA] Mpn
--------------------------------------------------------------------------------
--------------------------------------------------------------------------------
--------------------------------------------------------------------------------
--------------------------------------------------------------------------------
--------------------------------------------------------------------------------
--------------------------------------------------------------------------------
--------------------------------------------------------------------------------
--------------------------------------------------------------------------------
--------------------------------------------------------------------------------
-------------------------------------------ATTTT---TCTGAGAGTTTGATCCTGGCTCAG-GATT
```

```
AACGCTGGCGGCATGCCTAATACATGCAAGTCGATCGAAAGTAGTAAT------------------------------AC
TTTAGAGGCGAACGGGTGAGTAACA-CGTATCCAATC-------------------------------------------
-----TACCTTATAA--TGGGGGATAACTAGTTGAAA-GACTAGCTAATACCGCAT-AAGAACTTTGGTTCGCATGAATC
A-----------------------------------------------AAGTTGAAAGGACCTGCA---AGGGTTCGTTA
TTTGATGAGGGTGCGCCATATCAGCTAGTTGGTGGGGT-AACGGCCTACCAAGGCAATGACGTGTAGCTATGCTGAGAAG
TAGAATAGCCACAATGGGACTGAGACACGGCCCATACTCCTACGGG--AGGCAGCAGTAGGGAATTTTCACAATGAGCG
AAAGCTTGAT---GGAG---C-----------AATGCCGCGTGAACGATGAAGGTCTTTAAGATTGTAAAGTTCTTTTAT
TTGGGAAGAATG-ACTTTAGCAGGTAATGGCTAGAGTTTGACTGTACCATTTTGAATAAGTGACGACTAACTATGTGCCA
GCAGTCGCGGTAATACATAGGTCGCAAGCGTTATCCGGATTTATTGGGCGTAAAGCAAGCGCAGGCGGATTGAAAAGTCT
GGTGTTAAAGGCAGCTGCTTAACAG-TT-GTAT----------------------------------GCATTGGAAACT-
-----------------------------------ATTAATCTAGAGTGTGGTAGGGAGTTTTGGAATTTCATGTGGAGC
GGTGAAATGCGTAGATATATGAAGGA------------------------------------------------------
--------------------------------------------------------------------------------
-----------------------------------ACACCAGTGGCGAAGGCGAAAACTTAGGCCATTACTGACGCTTAG
GCTTGAAAGTGTGGGGAGCAAATAGGATTAGATACCCTAGTAGTCCACACCGTAAAC-----------------------
--------------------------------------------------------------------------------
---------GATAGATACTAGCTGTCGGGGC---GA-----T---CCCCTCGGTAGTGAAGTTAACACATTAAGTATCTC
GCCTGGGTAGTACATTCGCAAGAATGAAACTCAAACGGAATTGACGGGGACCCGCACA-AGTGGTGGAGCA-TGTTGCTT
AATTCGACGGTACACGAAAAACCTTACCTAGACTTGACATCCTTGGCAAAGTTATGGAAACA------------------
-------------------------------TAATGGAGGTTAACCGAGTGACAGGTGGTGCATGGTTGTCGTCAGCTCG
TGTCG-TGAGATG--TTGGGTTAAGTCCCGCAAC-----------------------------------------------
--------------------------------------------------------------------------------
----------G------AGCGCAACCCTTATCGTTAGTTACATT---GTCTAGCGA------------------------
-----GACT----GCTAATGCAAATTGGAGGAAGGAAGGGATGACGTCAAATCA-TCATGC-CCCTTATGTCTAG-GGCT
GCAAACGTGCTACAATGGCCAATACAAACAGTCG-CCAGCTTG-------------------------------------
----------------------------------------TAAAAGTGAGCAAATCT--GTAAAGTTGGTCTCAGTTCGG
ATTGAGGGCTGCAATTCGTCCTCATGAAG--TCGGAATCACTAGTAATCGCGAATCAGCTATGTCGCGGTGAATACGTTC
TCGGGTCTTGTACACACCGCCCGTCAAACTATGAAAGCTGGTAATATTTAAAAACGTGTTGCTAACCATTAG--------
-------------------------GAAGCGCATGTCAAGGATAGCACCGGTGATTGGAGTTAAGTCGTAACAAGGTACC
CCTACGAGAACGTGGGGGTGGATCACCT----------------------------------------------------
------------------------------------------------------
>lcl|AE000513.1_rrna_6 [locus_tag=DR_r01] [product=16S ribosomal RNA] [location=84836..86337] [gbkey=rRNA] Dra
--------------------------------------------------------------------------------
--------------------------------------------------------------------------------
--------------------------------------------------------------------------------
--------------------------------------------------------------------------------
--------------------------------------------------------------------------------
--------------------------------------------------------------------------------
--------------------------------------------------------------------------------
--------------------------------------------------------------------------------
---------------------------------------------TTT---ATGGAGAGTTTGATCCTGGCTCAG-GGTG
AACGCTGGCGGCGTGCTTAAGACATGCAAGTCGAACGCGGTCTTCGGAC-------------------------------
-CGAGTGGCGCACGGGTGAGTAACA-CGTAACTGACC-------------------------------------------
-----TACCCAGAAG--TCACGAATAACTGGCCGAAA-GGTCCGCTAATACGTGAT-GTGGTGATGC-------------
----------------------------------------AC------CGTGGTGCATCACTAAAG---ATTTATCGCTT
CTGGATGGGGTTGCGTTCCATCAGCTGGTTGGTGGGGT-AAAGGCCTACCAAGGCGACGACGGATAGCCGGCCTGAGAGG
GTGGCCGGCCACAGGGGCACTGAGACACGGTCCCACTCCTACGGG--AGGCAGCAGTTAGGAATCTTCCACAATGGGCG
CAAGCCTGAT---GGAG---C-----------GACGCCGCGTGAGGGATGAAGGTTTTCGG-ATCGTAAACCTCTGAATC
TGGGACGAAAGA-GCCT--------------TAGGGCAGATGACGGTACCAGAGTAATAGCACCGGCTAACTCCGTGCCA
GCAGCCGCGGTAATACGGAGGGTGCAAGCGTTACCCGGAATCACTGGGCGTAAAGGGCGTGTAGGCGGAAATTTAAGTCT
GGTTTTAAAGACCGGGGCTCAACCTCGG-GGAT----------------------------------GGACTGGATACT-
-----------------------------------GGATTTCTTGACCTCTGGAGAGGTAACTGGAATTCCTGGTGTAGC
GGTGGAATGCGTAGATACCAGGAGGA------------------------------------------------------
--------------------------------------------------------------------------------
-----------------------------------ACACCAATGGCGAAGGCAAGTTACTGGACAGAAGGTGACGCTGAG
GCGCGAAAGTGTGGGGAGCAAACCGGATTAGATACCCGGGTAGTCCACACCCTAAAC-----------------------
--------------------------------------------------------------------------------
---------GATGTACGTTGGCTAAGCGCAG---G-----AT---GCTGTGCTTGGCGAAGCTAACGCGATAAACGTACC
GCCTGGGAAGTACGGCCGCAAGGTTGAAACTCAA-AGGAATTGACGGGGGCCCGCACA-AGCGGTGGAGCA-TGTGGTTT
AATTCGAAGCAACGCGAAGAACCTTACCAGGTCTTGACATGCTAGGAACTTTGCAGAGATGCAG----------------
---------------------------AGGTGCCCTTCGGGGAACCTAGACACAGGTGCTGCATGGCTGTCGTCAGCTCG
TGTCG-TGAGATG--TTGGGTTAAGTCCCGCAAC-----------------------------------------------
--------------------------------------------------------------------------------
----------G------AGCGCAACCCTTGCCTTTAGTTGTCAG---CATTCA-------------GTTGGACACTCTAG
AGGGACTG------CCTATGAAAGTAGGAGGAAGGCGGGGATGACGTCTAGTCA-GCATGG-TCCTTACGTCCTG-GGCG
ACACACGTGCTACAATGGGTAGGACAACGCGCAG-CAAACCCG-------------------------------------
----------------------------------------CGAGGGTAAGCGAATCGC-TAAAACCTATCCCCAGTTCAG
ATCGGAGTCTGCAACTCGACTCCGTGAAG--TTGGAATCGCTAGTAATCGCGGGTCAGCA-TACCGCGGTGAATACGTTC
CCGGGCCTTGTACACACCGCCCGTCACACCATGGGAGTAGATTGCAGTTGAAACCGCCGGGAGCTT--------------
------------------------AACGGCAGGCGTCTAGACTGTGGTTTATGACTGGGGTGAAGTCGTAACAAGGTAAC
TGTACCGGAAGGTGCGGTTGGATCACCTCCTTT-----------------------------------------------
------------------------------------------------------
>lcl|BA000022.2_rrna_33 [gene=rrn16Sa] [product=16S ribosomal RNA] [location=complement(2452187..2453675)]
[gbkey=rRNA] Syn
--------------------------------------------------------------------------------
--------------------------------------------------------------------------------
--------------------------------------------------------------------------------
--------------------------------------------------------------------------------
--------------------------------------------------------------------------------
--------------------------------------------------------------------------------
--------------------------------------------------------------------------------
--------------------------------------------------------------------------------
---------------------------------------------ACA---ATGGAGAGTTTGATCCTGGCTCAG-GATG
AACGCTGGCGGTATGCCTAACACATGCAAGTCGAACGGAGTTC---------------T---------------TCGGAA
CTTAGTGGCGGACGGGTGAGTAACA-CGTGAGA-ACC--------------------------------------------
-----TACCTTCAGA--ATGGGGACAACAGTTGGAAA-CGACTGCTAATACCCAAT-GTG--------------------
---------------------------------------------------C-CGAAAGGTGAAAG---ATTTATCGTCT
```

GAAGATGGGCTCGCGTCTGATTAGCTAGATGGTGGGGT-AAGAGCCTACCATGGCAACGATCAGTAGCTGGTCTGAGAGG
ATGAGCAGCCACACTGGGACTGAGACACGGCCCAGACTCCTACGGG--AGGCAGCAGTGGGGAATTTTCCGCAATGGGCG
AAAGCCTGAC---GGAG---C-----------AATACCGCGTGAGGGAGGAAGGTCCTTGG-ATTGTAAACCTCTTTTAT
CAGGGAAGAAG-------------------------TTCTGACGGTACCTGATGAATAAGCATCGGCTAACTCCGTGCCA
GCAGCCGCGGTAATACGGAGGATGCAAGCGTTATCCGGAATTATTGGGCGTAAAGCGTCCGTAGGTGGTTATGCAAGTCT
GCCGTTAAAGAATGGAGCTTAACTCCAT-AGGA----------------------------------GCGGTGGAAACT-
-----------------------------------GCAAGACTAGAGTACAGTAGGGGTAGCAGGAATTCCCAGTGTAGC
GGTGAAATGCGTAGATATTGGGAAGA------------------------------------------------------
--------------------------------------------------------------------------------
-----------------------------------ACATCGGTGGCGAAAGCGTGCTACTGGGCTGAAACTGACACTGAG
GGACGAAAGCTAGGGTAGCGAAAGGGATTAGATACCCCTGTAGTCCTAGCCGTAAAC-----------------------
--------------------------------------------------------------------------------
---------GATGGATACTAGGCGTGGCTTG---TAT-CGAC---CCGAGCCGTGCCGAAGCTAACGCGTTAAGTATCCC
GCCTGGGGAGTACGCACGCAAGTGTGAAACTCAA-AGGAATTGACGGGGGCCCGCACA-AGCGGTGGAGTA-TGTGGTTT
AATTCGATGCAACGCGAAGAACCTTACCAAGGCTTGACATCCCTGGAATCCTGCGGAAA----CG---------------
--------------------------TGGGAGTGCCTTAGGGAGCCAGGAGACAGGTGGTGCATGGCTGTCGTCAGCTCG
TGTCG-TGAGATG--TTGGGTTAAGTCCCGCAAC----------------------------------------------
--------------------------------------------------------------------------------
----------G------AGCGCAACCCTCGTTTTAGTTGCCAT---CATTAA--GT----------TG--GGCACTCTAG
AGAGACTGC----CGGTGACA-AACCGGAGGAAGGTGGGGATGACGTCAAGTCA-TCATGC-CCCTTACGCCTTG-GGCT
ACACACGTACTACAATGGTCGGGACAACGGGCAG-CGAGCTCG-------------------------------------
----------------------------------------CGAGAGTAAGCGAATCCCATCAAACCCGGCCTCAGTTCAG
ATTGCAGGCTGCAACTCGCCTGCATGAAG--GAGGAATCGCTAGTAATCGCAGGTCAGAA-TACTGCGGTGAATTCGTTC
CCGGGCCTTGTACACACCGCCCGTCACACCATGGGAGCTGGTCACGCCCGAAGTCGTTACTCTAACCTT--A--------
-----------------------GGGAGGAGGGCGCCGAAGGCAGGGCTAGTGACTGGGGTGAAGTCGTAACAAGGTAGC
CGTACCGGAAGGTGTGGCTGGATCACCTCCTTT-----------------------------------------------
------------------------------------------------------
>lcl|AJ235269.1_rrna_26 [location=complement(772263..773769)] [gbkey=rRNA] Rpr
--------------------------------------------------------------------------------
--------------------------------------------------------------------------------
--------------------------------------------------------------------------------
--------------------------------------------------------------------------------
--------------------------------------------------------------------------------
--------------------------------------------------------------------------------
--------------------------------------------------------------------------------
--------------------------------------------------------------------------------
------------------------------------GACAGAATCAAA---CTTGAGAGTTTGATCCTGGCTCAG-AACG
AACGCTATCGGTATGCTTAACACATGCAAGTCGAACGGATTAACTAGAGCTCGCT---T---------------TAGTTA
ATTAGTGGCAGACGGGTGAGTAACA-CGTGGGA-ATC-------------------------------------------
-----TACCCATCAG--TACGGAATAACTTTTAGAAA-TAAAAGCTAATACCGTAT-ATT--------------------
---------------------------------------------------CTCTACGGAGGAAAG---ATTTATCGCTG
ATGGATGGGCCCGCGTCAGATTAGGTAGTTGGTGAGGT-AATGGCTCACCAAGCCGACGATCTGTAGCTGGTCTGAGAGG
ATGATCAGCCACACTGGGACTGAGACACGGCCCAGACTCCTACGGG--AGGCAGCAGTGGGGAATATTGGACAATGGGCG
AAAGCCTGAT---CCAG---C-----------AATACCGAGTGAGTGATGAAGGCCTTAGG-GTTGTAAAGCTCTTTTAG
CAAGGAAG--A-------------------------TAATGACGTTACTTGCAGAAAAAGCCCCGGCTAACTCCGTGCCA
GCAGCCGCGGTAAGACGGAGGGGGCTAGCGTTGTTCGGAATTACTGGGCGTAAAGAGTGCGTAGGCGGTTTAGTAAGTTG
GAAGTGAAAGCCCGGGGCTTAACCTCGG-AATT----------------------------------GCTTTCAAAACT-
-----------------------------------ACTAATCTAGAGTGTAGTAGGGGATGATGGAATTCCTAGTGTAGA
GGTGAAATTCTTAGATATTAGGAGGA------------------------------------------------------
--------------------------------------------------------------------------------
-----------------------------------ACACCGGTGGCGAAGGCGGTCATCTGGGCTACAACTGACGCTGAT
GCACGAAAGCGTGGGGAGCAAACAGGATTAGATACCCTGGTAGTCCACGCCGTAAAC-----------------------
--------------------------------------------------------------------------------
---------GATGAGTGCTAGATATCGGAGG---ATT----C---TCTTTCGGTTTCGCAGCTAACGCATTAAGCACTCC
GCCTGGGGAGTACGGTCGCAAGATTAAAACTCAA-AGGAATTGACGGGGGCTCGCACA-AGCGGTGGAGCA-TGCGGTTT
AATTCGATGTTACGCGAAAAACCTTACCAACCCTTGACATGGTGGTTACGGATTGCAGAGATGCT---------------
--------------------------TTCCTTCAGTTCGGCTGGGCCACACACAGGTGTTGCATGGCTGTCGTCAGCTCG
TGTCG-TGAGATG--TTGGGTTAAGTCCCGCAAC----------------------------------------------
--------------------------------------------------------------------------------
----------G------AGCGCAACCCTTATTCTTATTTGCCAG---TGGGTA--AT---------GCCGGGAACTATAA
GAAAACTGC----CGGTGATA-AGCCGGAGGAAGGTGGGGACGACGTCAAGTCA-TCATGG-CCCTTACGGGTTG-GGCT
ACACGCGTGCTACAATGGTGTTTACAGAGGGAAG-CAATACGG-------------------------------------
----------------------------------------TGACGTGGAGCAAATCC--CTAAAAGACATCTCAGTTCGG
ATTGTTCTCTGCAACTCGAGAGCATGAAG--TTGGAATCGCTAGTAATCGCGGATCAGCA-TGCCGCGGTGAATACGTTC
TCGGGCCTTGTACACACTGCCCGTCACGCCATGGGAGTTGGTTTTACCTGAAGGTGGTGAGCTAACGCA-----------
------------------------AGAGGCAGCCAACCACGGTAAAATTAGCGACTGGGGTGAAGTCGTAACAAGGTAGC
CGTAGGGGAACTGCGGCTGGA-TTACCTCCTTA-----------------------------------------------
------------------------------------------------------
>lcl|CP000230.1_rrna_64 [locus_tag=Rru_AR0064] [product=16S ribosomal RNA] [location=complement(3810698..3812174)] [gbkey=rRNA] Rru
--------------------------------------------------------------------------------
--------------------------------------------------------------------------------
--------------------------------------------------------------------------------
--------------------------------------------------------------------------------
--------------------------------------------------------------------------------
--------------------------------------------------------------------------------
--------------------------------------------------------------------------------
--------------------------------------------------------------------------------
-------------------------------------------------------AGAGTTTGATCCTGGCTCAG-GACG
AACGCTGGCGGCAGGCCTAACACATGCAAGTCGAACGCATCCT---------------T-----------------CGGG
ATGAGTGGCGCACGGGTGAGTAACA-CGTGGGA-ACG-------------------------------------------
-----TACCTTGGAG--TGCGGAATAATCTTTGGAAA-CGAGGACTAATACCGCAT-ACG--------------------
---------------------------------------------------CCCTTAGGGGGAAAG---ATTTATCGCTC
CAAGATCGGCCCGCGTCCGATTAGCTAGTTGGCGGGGT-AATGGCCCACCAAGGCGACGATCGGTAGCTGGTCTGAGAGG
ATGGCCAGCCACACTGGGACTGAGACACGGCCCAGACTCCTACGGG--AGGCAGCAGTGGGGAATATTGCGCAATGGGGG
CAACCCTGAC---GCAG---C-----------CATGCCGCGTGAGTGAAGAAGGCCTTCGG-GTTGTAAAGCTCTTTCGG
GTGTGAAG--A-------------------------TGATGACGGTAACACCAGAAGAAGCCCCGGCTAACTTCGTGCCA

GCAGCCGCGGTAATACGAAGGGGGCAAGCGTTGTTCGGAATTACTGGGCGTAAAGAGCGCGTAGGCGGTCTGATTAGTCA
GAGGTGAAATCCCAGAGCTCAACTTTGG-AACT----------------------------------GCCTTTGATACT-
-----------------------------------GTTAGACTAGAATCCGTGAGAGGGTGGTGGAATTCCCAGTGTAGA
GGTGAAATTCGTAGATATTGGGAGGA------------------------------------------------------
--------------------------------------------------------------------------------
-----------------------------------ACACCAGTGGCGAAGGCGGCCACCTGGCGCGGTATTGACGCTGAG
GCGCGAAAGCGTGGGGAGCAAACAGGATTAGATACCCTGGTAGTCCACGCCGTAAAC-----------------------
--------------------------------------------------------------------------------
---------GATGAGTGCTAGATGTCGGGGT---ACA---TG---TACCTCGGTGTCGCAGCTAACGCATTAAGCACTCC
GCCTGGGGAGTACGGCCGCAAGGTTAAAACTCAA-AGGAATTGACGGGGGCCCGCACA-AGCGGTGGAGCA-TGTGGTTT
AATTCGAAGCAACGCGCAGAACCTTACCAGCCCTTGACATCCCGTGACACTTCCAGA----GATG---------------
--------------------------GAAGGTTCCCTTCGGGGACACGGTGACAGGTGCTGCATGGCTGTCGTCAGCTCG
TGTCG-TGAGATG--TTGGGTTAAGTCCCGCAAC-----------------------------------------------
--------------------------------------------------------------------------------
----------G------AGCGCAACCCTCATCTTCAGTTGCCAG---CAAGTA--AC---------GTTGGGCACTCTGA
AGAGACTGC----CGGTGACA-AGCCGGAGGAAGGTGGGGATGACGTCAAGTCC-TCATGG-CCCTTACGGGCTG-GGCT
ACACACGTGCTACAATGGCGCCTACAATGGGCAG-CGACCTCG-------------------------------------
----------------------------------------CGAGGGGAAGCTAATCT--CCAAAAGGCGTCTCAGTTCGG
ATTGCACTCTGCAACTCGGGTGCATGAAG--TCGGAATCGCTAGTAATCGTGGATCAGCA-TGCCACGGTGAATACGTTC
CCGGGCCTTGTACACACCGCCCGTCACACCATGGGAGTTGGTTCTACCCGAAGACGGTACGCTAACCGC--A--------
-----------------------AGGAGGCAGCCGGCCACGGTAGGGTCAGCGACTGGGGTGAAGTCGTAACAAGGTAGC
CGTAGGGGAACCTGCGGCTGGATCACCTCCTTT-----------------------------------------------
------------------------------------------------------
>lcl|CP011246.1_rrna_1 [locus_tag=Ach5_r010] [product=16S ribosomal RNA] [location=57233..58705] [gbkey=rRNA] Atu
--------------------------------------------------------------------------------
--------------------------------------------------------------------------------
--------------------------------------------------------------------------------
--------------------------------------------------------------------------------
--------------------------------------------------------------------------------
--------------------------------------------------------------------------------
--------------------------------------------------------------------------------
--------------------------------------------------------------------------------
-------------------------------------------------------AGAGTTTGATCCTGGCTCAG-AACG
AACGCTGGCGGCAGGCTTAACACATGCAAGTCGAACGCCCCG----------------------------------CAAG
GGGAGTGGCAGACGGGTGAGTAACG-CGTGGGA-ATC-------------------------------------------
-----TACCGTGCCC--TGCGGAATAGCTCCGGGAAA-CTGGAATTAATACCGCAT-ACG--------------------
---------------------------------------------------CCCTACGGGGGAAAG---ATTTATCGGGG
TATGATGAGCCCGCGTTGGATTAGCTAGTTGGTGGGGT-AAAGGCCTACCAAGGCGACGATCCATAGCTGGTCTGAGAGG
ATGATCAGCCACATTGGGACTGAGACACGGCCCAAACTCCTACGGG--AGGCAGCAGTGGGGAATATTGGACAATGGGCG
CAAGCCTGAT---CCAG---C-----------CATGCCGCGTGAGTGATGAAGGCCTTAGG-GTTGTAAAGCTCTTTCAC
CGGAGAAG--A-------------------------TAATGACGGTATCCGGAGAAGAAGCCCCGGCTAACTTCGTGCCA
GCAGCCGCGGTAATACGAAGGGGGCTAGCGTTGTTCGGAATTACTGGGCGTAAAGCGCACGTAGGCGGATATTTAAGTCA
GGGGTGAAATCCCAGAGCTCAACTCTGG-AACT----------------------------------GCCTTTGATACT-
-----------------------------------GGGTATCTTGAGTATGGAAGAGGTAAGTGGAATTCCGAGTGTAGA
GGTGAAATTCGTAGATATTCGGAGGA------------------------------------------------------
--------------------------------------------------------------------------------
-----------------------------------ACACCAGTGGCGAAGGCGGCTTACTGGTCCATTACTGACGCTGAG
GTGCGAAAGCGTGGGGAGCAAACAGGATTAGATACCCTGGTAGTCCACGCCGTAAAC-----------------------
--------------------------------------------------------------------------------
---------GATGAATGTTAGCCGTCGGGCA---GTA---TA---CTGTTCGGTGGCGCAGCTAACGCATTAAACATTCC
GCCTGGGGAGTACGGTCGCAAGATTAAAACTCAA-AGGAATTGACGGGGGCCCGCACA-AGCGGTGGAGCA-TGTGGTTT
AATTCGAAGCAACGCGCAGAACCTTACCAGCTCTTGACATTCGGGGTTTGGGCAGTGGAGACATT---------------
--------------------------GTCCTTCAGTTAGGCTGGCCCCAGAACAGGTGCTGCATGGCTGTCGTCAGCTCG
TGTCG-TGAGATG--TTGGGTTAAGTCCCGCAAC-----------------------------------------------
--------------------------------------------------------------------------------
----------G------AGCGCAACCCTCGCCCTTAGTTGCCAG---CATTTA--GT---------TG--GGCACTCTAA
GGGGACTGC----CGGTGATAAGCCGAGAGGAAGGTGGGGATGACGTCAAGTCC-TCATGG-CCCTTACGGGCTG-GGCT
ACACACGTGCTACAATGGTGGTGACAGTGGGCAG-CGAGACAG-------------------------------------
----------------------------------------CGATGTCGAGCTAATCT--CCAAAAGCCATCTCAGTTCGG
ATTGCACTCTGCAACTCGAGTGCATGAAG--TTGGAATCGCTAGTAATCGCAGATCAGCA-TGCTGCGGTGAATACGTTC
CCGGGCCTTGTACACACCGCCCGTCACACCATGGGAGTTGGTTTTACCCGAAGGTAGTGCGCTAACCGC--A--------
-----------------------AGGAGGCAGCTAACCACGGTAGGGTCAGCGACTGGGGTGAAGTCGTAACAAGGTAGC
CGTAGGGGAACCTGCGGCTGGATCACCT----------------------------------------------------
------------------------------------------------------
>lcl|AP012206.1_rrna_52 [locus_tag=BJ6T_61860] [product=16S ribosomal RNA] [location=complement(6255003..6256492)] [gbkey=rRNA] Bja
--------------------------------------------------------------------------------
--------------------------------------------------------------------------------
--------------------------------------------------------------------------------
--------------------------------------------------------------------------------
--------------------------------------------------------------------------------
--------------------------------------------------------------------------------
--------------------------------------------------------------------------------
--------------------------------------------------------------------------------
--------------------------------------------CCAA---CTTGAGAGTTTGATCCTGGCTCAG-AGCG
AACGCTGGCGGCAGGCTTAACACATGCAAGTCGAGCGGGCGTA---------------G---------------CAATAC
GTCAGCGGCAGACGGGTGAGTAACG-CGTGGGA-ACG-------------------------------------------
-----TACCTTTTGG--TTCGGAACAACACAGGGAAA-CTTGTGCTAATACCGGAT-AAG--------------------
---------------------------------------------------CCCTTACGGGGAAAG---ATTTATCGCCG
AAAGATCGGCCCGCGTCTGATTAGCTAGTTGGTGAGGT-AATGGCTCACCAAGGCGACGATCAGTAGCTGGTCTGAGAGG
ATGATCAGCCACATTGGGACTGAGACACGGCCCAAACTCCTACGGG--AGGCAGCAGTGGGGAATATTGGACAATGGGGG
CAACCCTGAT---CCAG---C-----------CATGCCGCGTGAGTGATGAAGGCCCTAGG-GTTGTAAAGCTCTTTTGT
GCGGGAAG--A-------------------------TAATGACGGTACCGCAAGAATAAGCCCCGGCTAACTTCGTGCCA
GCAGCCGCGGTAATACGAAGGGGGCTAGCGTTGCTCGGAATCACTGGGCGTAAAGGGTGCGTAGGCGGGTCTTTAAGTCA
GGGGTGAAATCCTGGAGCTCAACTCCAG-AACT----------------------------------GCCTTTGATACT-
-----------------------------------GAGGATCTTGAGTTCGGGAGAGGTGAGTGGAACTGCGAGTGTAGA

```
GGTGAAATTCGTAGATATTCGCAAGA------------------------------------------------------
--------------------------------------------------------------------------------
-----------------------------------ACACCAGTGGCGAAGGCGGCTCACTGGCCCGATACTGACGCTGAG
GCACGAAAGCGTGGGGAGCAAACAGGATTAGATACCCTGGTAGTCCACGCCGTAAAC-----------------------
--------------------------------------------------------------------------------
---------GATGAATGCCAGCCGTTAGTGG---GTT---TA---CTCACTAGTGGCGCAGCTAACGCTTTAAGCATTCC
GCCTGGGGAGTACGGTCGCAAGATTAAAACTCAA-AGGAATTGACGGGGGCCCGCACA-AGCGGTGGAGCA-TGTGGTTT
AATTCGACGCAACGCGCAGAACCTTACCAGCCCTTGACATGTCCAGGACCGGTCGCAGAGATGT----------------
---------------------------GACCTTCTCTTCGGAGCCTGGAGCACAGGTGCTGCATGGCTGTCGTCAGCTCG
TGTCG-TGAGATG--TTGGGTTAAGTCCCGCAAC----------------------------------------------
--------------------------------------------------------------------------------
----------G------AGCGCAACCCCCGTCCTTAGTTGCTAC---CATTTA--GT---------TG--AGCACTCTAA
GGAGACTGC----CGGTGATAAGCCGCGAGGAAGGTGGGGATGACGTCAAGTCC-TCATGG-CCCTTACGGGCTG-GGCT
ACACACGTGCTACAATGGCGGTGACAATGGGATG-CTAAGGGG-------------------------------------
----------------------------------------CGACCCTTCGCAAATCT--CAAAAAGCCGTCTCAGTTCGG
ATTGGGCTCTGCAACTCGAGCCCATGAAG--TTGGAATCGCTAGTAATCGTGGATCAGCA-CGCCACGGTGAATACGTTC
CCGGGCCTTGTACACACCGCCCGTCACACCATGGGAGTTGGTTTTACCTGAAGACGGTGCGCTAACCCGCAA--------
-----------------------GGGAGGCAGCCGGCCACGGTAGGGTCAGCGACTGGGGTGAAGTCGTAACAAGGTAGC
CGTAGGGGAACCTGCGGCTGGATCACCTCCTTT-----------------------------------------------
------------------------------------------------------
>lcl|AP012051.1_rrna_12 [locus_tag=CSE_r0010] [product=16S ribosomal RNA] [location=541386..542944] [gbkey=rRNA]
Cex
--------------------------------------------------------------------------------
--------------------------------------------------------------------------------
--------------------------------------------------------------------------------
--------------------------------------------------------------------------------
--------------------------------------------------------------------------------
--------------------------------------------------------------------------------
--------------------------------------------------------------------------------
--------------------------------------------------------------------------------
------------------------------------------ATTTTT---TCGGAGAGTTTGATCCTGGCTCAG-GATG
AACGCTGGCGGCGTGCCTTACACATGCAAGTCGAGCGGAGAAGATTTCCTGTGGC---AA--------CACAGGAGTCTT
CTTAGCGGCAAACGGGTGAGTAATG-GATGAGTAACC-------------------------------------------
-----TGCCCTGTAG--ACGGGGATAGCCATCCGAAA-GGATGGGTAATACCCGAT-AATGTTGGAGGACTTCGGTT---
----------------------------------------CT------CCAAAGAAAGGTGGCGCA---AGCTACCGCTT
CAGGATGGGCTCATCTCCTATCAGCTAGTTGGTGAGGT-AATGGCTCACCAAGGCTATGACGGGTAGCTGGCCTGAGAGG
GTGGTCAGCCACATTGGGACTGAGACACGGCCCAGACTCCTACGGG--AGGCAGCAGTGGGGAATCTTGGTCAATGGGCG
AAAGCCTGAA---CCAG---C-----------GACGCCGCGTGGAGGACGAAGGCCGAAAG-GTCGTAAACTCCTTTTGC
AAGGGAAAAAGG-GAGGGGTTCGAATAGGGCCCTATCCTTGATGGTACCTTGCGAATAAGCCCCAGCCAACTACGTGCCA
GCAGCTGCGGTAAAACGTAGGGGGCGAGCGTTATCCGGAATCACTGGGTGTAAAGGGTACGCAGGCGGACTTGTAAGTTG
GATGTAAAAACTCAGGGCTCAACCTTGA-GCTT----------------------------------GCATTCAATACT-
-----------------------------------ACAAGTCTAGAGAGCAGGAGAGGGGAGTGGAATTCCCGGTGTAGG
GGTAAAATCCGTAGATATCGGGAGGA------------------------------------------------------
--------------------------------------------------------------------------------
-----------------------------------ACACCAGTGGCGAAGGCGGCTCCCTGGCCTGTCTCTGACGCTCAG
GTACGAAAGCGTGGGGAGCAAACAGGATTAGATACCCTGGTAGTCCACGCCCTAAAC-----------------------
--------------------------------------------------------------------------------
---------GATGGGTGCTTGGTGTTGGGGG---G---TAAT---TCCCTCAGTGCCGTAGCTAACGCGTTAAGCACCCC
ACCTGGGGAGTACGGCCGCAAGGTTGAAACTCAA-AGGAATTGACGGGGACCCGCACA-AGCAGCGAGGCA-TGTGGTTT
AATTCGATGCTACACGAAGAACCTTACCTGGGCTTGACATGCATGTGAAAGCCTGAAGAAACTCGGGCC-----------
---------------------CCCATAGGGGTTACACCCTATGGCACATGCACAGGTGCTGCATGGTTGTCGTCAGCTCG
TGTCG-TGAGATG--TACGGTTAAGTCCGTGAAC----------------------------------------------
--------------------------------------------------------------------------------
----------G------AGCGCAACCCCTGCCCTTAGTTGCTAA---TAGGCTTCGG---------CCTATGCACTCTAA
GGGGACTGC----CAGCGATA-AGCTGGAGGAAGGTGGGGATGACGTCAAATCC-TCATGG-CCCTTATGCCCAG-GGCT
ACACACATGCGACAATGGTCGGGACAATGCGTTG-CAAACCAG-------------------------------------
----------------------------------------TAATGGGGAGCTAATCGC-AAAAAACCGACCCCAGTACGG
ATTGAGGGTTGCAACTCACCCTCATGAAG--CTGGAGTTGCTAGTAACCGCCGGTCAGCTATACGGCGATGAATACGTTC
CCGGGTCTTGTACACACCGCCCGTCACACCACCCGAGTTGCGTGCACCCGAAGTGGCTCGGTGAGTCACG----------
--------------------------------------AAGGTGTGCGTGATGAGGAGGGTGAAGTCGTAACAAGGTAGG
TGTACGAGAACGTGCGCCTGGATCACCTCCTTT-----------------------------------------------
------------------------------------------------------
>lcl|AM884176.1_rrna_12 [gene=16s rRNA] [product=16S ribosomal RNA] [location=156015..157564] [gbkey=rRNA] Ctr
--------------------------------------------------------------------------------
--------------------------------------------------------------------------------
--------------------------------------------------------------------------------
--------------------------------------------------------------------------------
--------------------------------------------------------------------------------
--------------------------------------------------------------------------------
--------------------------------------------------------------------------------
--------------------------------------------------------------------------------
----------------------------------------------------CTGAGAATTTGATCTTGGTTCAG-ATTG
AACGCTGGCGGCGTGGATGAGGCATGCAAGTCGAACGGAGCAATTGTTTCGG----------------------CAATTG
TTTAGTGGCGGAAGGGTTAGTAATG-CATAGATAATT-------------------------------------------
-----TGTCCTTAAC--TTGGGAATAACGGTTGGAAA-CGGCCGCTAATACCGAAT-GTGGCGATATTTGGGCATCCGAG
TAAC---G--------------------------------TT------AAAGAAGGGGATCTTAGG---ACCTTTCGGTT
AAGGGAGAGTCTATGTGATATCAGCTAGTTGGTGGGGT-AAAGGCCTACCAAGGCTATGACGTCTAGGCGGATTGAGAGA
TTGGCCGCCAACACTGGGACTGAGACACTGCCCAGACTCCTACGGG--AGGCTGCAGTCGAGAATCTTTCGCAATGGACG
GAAGTCTGAC---GAAG---C-----------GACGCCGCGTGTGTGATGAAGGCTCTAGG-GTTGTAAAGCACTTTCGC
TTGGGAATAAGA-GAA-GGCGGTTAATACCCGCTGGATTTGAGCGTACCAGGTAAAGAAGCACCGGCTAACTCCGTGCCA
GCAGCTGCGGTAATACGGAGGGTGCTAGCGTTAATCGGATTTATTGGGCGTAAAGGGCGTGTAGGCGGAAAGGTAAGTTA
GTTGTCAAAGATCGGGGCTCAACCCCGA-GTCG----------------------------------GCATCTAATACT-
-----------------------------------ATTTTTCTAGAGGGTAGATGGAGAAAAGGGAATTTCACGTGTAGC
GGTGAAATGCGTAGATATGTGGAAGA------------------------------------------------------
--------------------------------------------------------------------------------
-----------------------------------ACACCAGTGGCGAAGGCGCTTTTCTAATTTATACCTGACGCTAAG
GCGCGAAAGCAAGGGGAGCAAACAGGATTAGATACCCTGGTAGTCCTTGCCGTAAAC-----------------------
```

```
--------------------------------------------------------------------------------
---------GATGCATACTTGATGTGGATGG---TCT-CAAC---CCCATCCGTGTCGGAGCTAACGCGTTAAGTATGCC
GCCTGAGGAGTACACTCGCAAGGGTGAAACTCAA-AAGAATTGACGGGGGCCCGCACA-AGCAGTGGAGCA-TGTGGTTT
AATTCGATGCAACGCGAAGGACCTTACCTGGGTTTGACATGTATATGACCGCGGCAGAAATG------------------
----------------------------TCGTTTTCCGCAAGGACATATACACAGGTGCTGCATGGCTGTCGTCAGCTCG
TGCCG-TGAGGTG--TTGGGTTAAGTCCCGCAAC----------------------------------------------
--------------------------------------------------------------------------------
----------G------AGCGCAACCCTTATCGTTAGTTGCCAG---CACTT---AG---------GGTGGGAACTCTAA
CGAGACTGC----CTGGGTTA-ACCAGGAGGAAGGCGAGGATGACGTCAAGTCA-GCATGG-CCCTTATGCCCAG-GGCG
ACACACGTGCTACAATGGCCAGTACAGAAGGTAG-CAAGATCG-------------------------------------
----------------------------------------TGAGATGGAGCAAATC-C-TCAAAGCTGGCCCCAGTTCGG
ATTGTAGTCTGCAACTCGACTACATGAAG--TCGGAATTGCTAGTAATGGCGTGTCAGCCATAACGCCGTGAATACGTTC
CCGGGCCTTGTACACACCGCCCGTCACATCATGGGAGTTGGTTTTACCTTAAGTCGTTGACTCAACCCGCAA--------
-----------------------GGGAGAGAGGCGCCCAAGGTGAGGCTGATGACTAGGATGAAGTCGTAACAAGGTAGC
CCTACCGGAAGGTGGGGCTGGATCACCTCCTTTT----------------------------------------------
------------------------------------------------------
>lcl|AL111168.1_rrna_8 [product=16S ribosomal RNA] [location=394130..395642] [gbkey=rRNA] Cje
--------------------------------------------------------------------------------
--------------------------------------------------------------------------------
--------------------------------------------------------------------------------
--------------------------------------------------------------------------------
--------------------------------------------------------------------------------
--------------------------------------------------------------------------------
--------------------------------------------------------------------------------
--------------------------------------------------------------------------------
-------------------------------------------TTTTT---ATGGAGAGTTTGATCCTGGCTCAG-AGTG
AACGCTGGCGGCGTGCCTAATACATGCAAGTCGAACGATGAAGCTTTTAGCTTGC---T-------------AGAAGTGG
ATTAGTGGCGCACGGGTGAGTAAGG-TATAGTTAATC-------------------------------------------
-----TGCCCTACAC--AAGAGGACAACAGTTGGAAA-CGACTGCTAATACTCTAT-ACTCCTGCTTAACAC--------
--------------------------------------------------AAGTTGAGTAGGGAAA---GTTTTTCGGTG
TAGGATGAGACTATATAGTATCAGCTAGTTGGTAAGGT-AATGGCTTACCAAGGCTATGACGCTTAACTGGTCTGAGAGG
ATGATCAGTCACACTGGAACTGAGACACGGTCCAGACTCCTACGGG--AGGCAGCAGTAGGGAATATTGCGCAATGGGGG
AAACCCTGAC---GCAG---C-----------AACGCCGCGTGGAGGATGACACTTTTCGG-AGCGTAAACTCCTTTTCT
TAGGGAAGAA--------------------------TTCTGACGGTACCTAAGGAATAAGCACCGGCTAACTCCGTGCCA
GCAGCCGCGGTAATACGGAGGGTGCAAGCGTTACTCGGAATCACTGGGCGTAAAGGGCGCGTAGGCGGATTATCAAGTCT
CTTGTGAAATCTAATGGCTTAACCATTA-AACT----------------------------------GCTTGGGAAACT-
-----------------------------------GATAGTCTAGAGTGAGGGAGAGGCAGATGGAATTGGTGGTGTAGG
GGTAAAATCCGTAGATATCACCAAGA------------------------------------------------------
--------------------------------------------------------------------------------
-----------------------------------ATACCCATTGCGAAGGCGATCTGCTGGAACTCAACTGACGCTAAG
GCGCGAAAGCGTGGGGAGCAAACAGGATTAGATACCCTGGTAGTCCACGCCCTAAAC-----------------------
--------------------------------------------------------------------------------
---------GATGTACACTAGTTGTTGGGGT---GCT--AGT---CATCTCAGTAATGCAGCTAACGCATTAAGTGTACC
GCCTGGGGAGTACGGTCGCAAGATTAAAACTCAA-AGGAATAGACGGGGACCCGCACA-AGCGGTGGAGCA-TGTGGTTT
AATTCGAAGATACGCGAAGAACCTTACCTGGGCTTGATATCCTAAGAACCTTATAGAGATATGA----------------
---------------------G----GGTGCTAGCTTGCTAGAACTTAGAGACAGGTGCTGCACGGCTGTCGTCAGCTCG
TGTCG-TGAGATG--TTGGGTTAAGTCCCGCAAC----------------------------------------------
--------------------------------------------------------------------------------
----------G------AGCGCAACCCACGTATTTAGTTGCTAA---CGGTTC--GG---------CCG-AGCACTCTAA
ATA-GACTG----CCTTCGTA-AGGAGGAGGAAGGTGTGGACGACGTCAAGTCA-TCATGG-CCCTTATGCCCAG-GGCG
ACACACGTGCTACAATGGCATATACAATGAGACG-CAATACCG-------------------------------------
----------------------------------------CGAGGTGGAGCAAATC-T-ATAAAATATGTCCCAGTTCGG
ATTGTTCTCTGCAACTCGAGAGCATGAAG--CCGGAATCGCTAGTAATCGTAGATCAGCCATGCTACGGTGAATACGTTC
CCGGGTCTTGTACTCACCGCCCGTCACACCATGGGAGTTGATTTCACTCGAAGCCGGAATACTAAACTAGT---------
------------------------------TACCGTCCACAGTGGAATCAGCGACTGGGGTGAAGTCGTAACAAGGTAAC
CGTAGGAGAACCTGCGGTTGGATCACCTCCT-------------------------------------------------
------------------------------------------------------
>lcl|CP003679.1_rrna_12 [locus_tag=TPSea814_00r05411] [product=16S ribosomal RNA] [location=231299..232859]
[gbkey=rRNA] Tpa
--------------------------------------------------------------------------------
--------------------------------------------------------------------------------
--------------------------------------------------------------------------------
--------------------------------------------------------------------------------
--------------------------------------------------------------------------------
--------------------------------------------------------------------------------
--------------------------------------------------------------------------------
--------------------------------------------------------------------------------
------------------------------------------ATAATG---ATGGAGAGTTTGATCCTGGCTCAG-AACG
AACGCTGGCGGTGCGTTTTAAGCATGCAAGTCGAACGGCAAGGAAGCGAATTTTC------------------GTTTCTC
TAGAGTGGCGGACTGGTGAGTAACG-CGTGGGTAATC-------------------------------------------
-----TGCCTTTGAG--ATGGGGATAGCCTCTAGAAA-TAGGGGGTAATACCGAAT-ACGCTCTTTTGGACGTAGGTCTT
T-------------------------------------------------GAGAGGAAAGGGGCTG---CGGCCTCGCTC
AGAGATGAGCCTGCGACCCATTAGCTTGTTGGTGGGGT-AATGGCCTACCAAGGCGTCGATGGGTATCCGACCTGAGAGG
GTGACCGGACACACTGGGACTGAGATACGGCCCAGACTCCTACGGG--AGGCAGCAGCTAAGAATATTCCGCAATGGGCG
AAAGCCTGAC---GGAG---C-----------GACACCGCGTGGATGAGGAAGGTCGAAAG-ATTGTAAAGTTCTTTTGC
CGACGAAGAATG-AGGACGGGAGGGAATGCCCGTTTGATGACGGTAGTCGTGCGAATAAGCCCCGGCTAATTACGTGCCA
GCAGCCGCGGTAACACGTAAGGGGCGAGCGTTGTTCGGAATTATTGGGCGTAAAGGGCATGCAGGCGGACTGGTAAGCCT
GGTGTGAAATCCCCGAGCTCAACTTGGG-AACT----------------------------------GCACTGGGTACT-
-----------------------------------GCTGGTCTAGAATCACGGAGGGGAAACCGGAATTCCAAGTGTAGG
GGTGGAATCTGTAGATATTTGGAAGA------------------------------------------------------
--------------------------------------------------------------------------------
-----------------------------------ACACCGGTGGCGAAGGCGGGTTTCTGGCCGATGATTGACGCTGAG
GTGCGAAGGTGTGGGGAGCGAACAGGATTAGATACCCTGGTAGTCCACACAGTAAAC-----------------------
--------------------------------------------------------------------------------
---------GATGTACACTAGGTGTTGGGGC---AT-----G---AGTCTCGGCGCCGACGCGAACGCATTAAGTGTACC
GCCTGGGGAGTATGCTCGCAAGAGTGAAACTCAA-AGGAATTGACGGGGGCCCGCACA-AGCGGTGGAGCA-TGTGGTTT
AATTCGATGATACGCGAGGAACCTTACCCGGGTTTGACATCAAGAGGAGCGCCGTAGAAATGCG----------------
```

```
----------------------------GTGGCGTAGCGATACGCCTCTTGACAGGTGCTGCATGGCTGTCGTCAGCTCG
TGCCG-TGAGGTG--TTGGGTTAAGTCCCGCAAC----------------------------------------------
--------------------------------------------------------------------------------
----------G------AGCGCAACCCCTACTGCCAGTTGCCAG---CAAGTG--GT---------GTTGGGGACTCTGG
CGGAACTGC----CGGTGACA-AACCGGAGGAAGGTGGGGATGACGTCAAGTCA-TCATGG-CCCTTATGTCCGG-GGCT
ACACACGTGCTACAATGGTTGCTACAGAGCGATG-CGAGGTTG-------------------------------------
----------------------------------------TGAAGTGGAGCAAACCGC-AAAAAGGCAATCGTAGTCCGG
ATTGAAGTCTGAAACTCGACTTCATGAAG--TTGGAATCGCTAGTAATCGCACATCAGCA-TGGTGCGGTGAATGTGTTC
CCGGGCCTTGTACACACCGCCCGTCACACCATCCGAGTTGGAGATACCCGAAGTCACTAGCCTAACCCGCAA--------
-----------------------GGGAGGGCGGTGCCGAAGGTATGTTTGGTAAGGAGGGTGAAGTCGTAACAAGGTAGC
CGTACCGGAAGGTGCGGCTGGATCACCTCCTTTCTAAGAGAA--------------------------------------
------------------------------------------------------
>lcl|U00096.3_rrna_88 [gene=rrsG] [locus_tag=b2591] [db_xref=ASAP:ABE-0008525,ECOCYC:EG30089,EcoGene:EG30089]
[product=16S ribosomal RNA] [location=complement(2729616..2731157)] [gbkey=rRNA] Eco
--------------------------------------------------------------------------------
--------------------------------------------------------------------------------
--------------------------------------------------------------------------------
--------------------------------------------------------------------------------
--------------------------------------------------------------------------------
--------------------------------------------------------------------------------
--------------------------------------------------------------------------------
--------------------------------------------------------------------------------
---------------------------------------------AAA---TTGAAGAGTTTGATCATGGCTCAG-ATTG
AACGCTGGCGGCAGGCCTAACACATGCAAGTCGAACGGTAACAGGAAGCAGC--T---TGCTG---------CTTCGCTG
ACGAGTGGCGGACGGGTGAGTAATG-TCTGGG-AAGC-------------------------------------------
-----TGCCTGATGG--AGGGGGATAACTACTGGAAA-CGGTAGCTAATACCGCAT-AATGTCGCAAGACC---------
------------------------------------------------AAAGAGGGGGACCTTCGG---GCCTCTTGCCA
TCGGATGTGCCCAGATGGGATTAGCTTGTTGGTGGGGT-AACGGCTCACCAAGGCGACGATCCCTAGCTGGTCTGAGAGG
ATGACCAGCCACACTGGAACTGAGACACGGTCCAGACTCCTACGGG--AGGCAGCAGTGGGGAATATTGCACAATGGGCG
CAAGCCTGAT---GCAG---C-----------CATGCCGCGTGTATGAAGAAGGCCTTCGG-GTTGTAAAGTACTTTCAG
CGGGGAGGAAGG-GAG-TAAAGTTAATACCTTTGCTCATTGACGTTACCCGCAGAAGAAGCACCGGCTAACTCCGTGCCA
GCAGCCGCGGTAATACGGAGGGTGCAAGCGTTAATCGGAATTACTGGGCGTAAAGCGCACGCAGGCGGTTTGTTAAGTCA
GATGTGAAATCCCCGGGCTCAACCTGGG-AACT----------------------------------GCATCTGATACT-
-----------------------------------GGCAAGCTTGAGTCTCGTAGAGGGGGGTAGAATTCCAGGTGTAGC
GGTGAAATGCGTAGAGATCTGGAGGA------------------------------------------------------
--------------------------------------------------------------------------------
-----------------------------------ATACCGGTGGCGAAGGCGGCCCCCTGGACGAAGACTGACGCTCAG
GTGCGAAAGCGTGGGGAGCAAACAGGATTAGATACCCTGGTAGTCCACGCCGTAAAC-----------------------
--------------------------------------------------------------------------------
---------GATGTCGACTTGGAGGTTGTGC---CCT--TGA---GGCGTGGCTTCCGGAGCTAACGCGTTAAGTCGACC
GCCTGGGGAGTACGGCCGCAAGGTTAAAACTCAA-ATGAATTGACGGGGGCCCGCACA-AGCGGTGGAGCA-TGTGGTTT
AATTCGATGCAACGCGAAGAACCTTACCTGGTCTTGACATCCACGGAAGTTTTCAGAGATGA------------------
---------------------------GAATGTGCCTTCGGGAACCGTGAGACAGGTGCTGCATGGCTGTCGTCAGCTCG
TGTTG-TGAAATG--TTGGGTTAAGTCCCGCAAC----------------------------------------------
--------------------------------------------------------------------------------
----------G------AGCGCAACCCTTATCCTTTGTTGCCAG---CGGT-C--CG---------GCCGGGAACTCAAA
GGAGACTGC----CAGTGATA-AACTGGAGGAAGGTGGGGATGACGTCAAGTCA-TCATGG-CCCTTACGACCAG-GGCT
ACACACGTGCTACAATGGCGCATACAAAGAGAAG-CGACCTCG-------------------------------------
----------------------------------------CGAGAGCAAGCGGACCTC-ATAAAGTGCGTCGTAGTCCGG
ATTGGAGTCTGCAACTCGACTCCATGAAG--TCGGAATCGCTAGTAATCGTGGATCAGAAT-GCCACGGTGAATACGTTC
CCGGGCCTTGTACACACCGCCCGTCACACCATGGGAGTGGGTTGCAAAAGAAGTAGGTAGCTTAACCTTCGG--------
-------------------------GAGGGCGCTTACCACTTTGTGATTCATGACTGGGGTGAAGTCGTAACAAGGTAAC
CGTAGGGGAACCTGCGGTTGGATCACCTCCTTA-----------------------------------------------
------------------------------------------------------
>lcl|AE008922.1_rrna_55 [locus_tag=XCC3840] [product=SSU 16S ribosomal RNA]
[location=complement(4561295..4562841)] [gbkey=rRNA] Xca
--------------------------------------------------------------------------------
--------------------------------------------------------------------------------
--------------------------------------------------------------------------------
--------------------------------------------------------------------------------
--------------------------------------------------------------------------------
--------------------------------------------------------------------------------
--------------------------------------------------------------------------------
--------------------------------------------------------------------------------
---------------------------------------------TAA---GTGAAGAGTTTGATCCTGGCTCAG-AGTG
AACGCTGGCGGCAGGCCTAACACATGCAAGTCGAACGGCAGCACAGTAAGAGCTT---GCTCT---------TATGGGTG
GCGAGTGGCGGACGGGTGAGGAATA-CATCGG-AATC-------------------------------------------
-----TACTCTTTCG--TGGGGGATAACGTAGGGAAA-CTTACGCTAATACCGCAT-ACGACCTACGGGTG---------
------------------------------------------------AAAGCGGAGGACCTTCGG---GCTTCGCGCGA
TTGAATGAGCCGATGTCGGATTAGCTAGTTGGCGGGGT-AAAGGCCCACCAAGGCGACGATCCGTAGCTGGTCTGAGAGG
ATGATCAGCCACACTGGAACTGAGACACGGTCCAGACTCCTACGGG--AGGCAGCAGTGGGGAATATTGGACAATGGGCG
CAAGCCTGAT---CCAG---C-----------CATGCCGCGTGGGTGAAGAAGGCCTTCGG-GTTGTAAAGCCCTTTTGT
TGGGAAAGAAAA-GCA-GTCGGTTAATACCCGATTGTTCTGACGGTACCCAAAGAATAAGCACCGGCTAACTTCGTGCCA
GCAGCCGCGGTAATACGAAGGGTGCAAGCGTTACTCGGAATTACTGGGCGTAAAGCGTGCGTAGGTGGTGGTTTAAGTCT
GTTGTGAAAGCCCTGGGCTCAACCTGGG-AATT----------------------------------GCAGTGGATACT-
-----------------------------------GGGTCACTAGAGTGTGGTAGAGGGTAGCGGAATTCCCGGTGTAGC
AGTGAAATGCGTAGAGATCGGGAGGA------------------------------------------------------
--------------------------------------------------------------------------------
-----------------------------------ACATCCGTGGCGAAGGCGGCTACCTGGACCAACACTGACACTGAG
GCACGAAAGCGTGGGGAGCAAACAGGATTAGATACCCTGGTAGTCCACGCCCTAAAC-----------------------
--------------------------------------------------------------------------------
---------GATGCGAACTGGATGTTGGGTG---CAA-TTTG---GCACGCAGTATCGAAGCTAACGCGTTAAGTTCGCC
GCCTGGGGAGTACGGTCGCAAGACTGAAACTCAA-AGGAATTGACGGGGGCCCGCACA-AGCGGTGGAGTA-TGTGGTTT
AATTCGATGCAACGCGAAGAACCTTACCTGGTCTTGACATCCACGGAACTTTCCAGAGATGG------------------
---------------------------ATTGGTGCCTTCGGGAACCGTGAGACAGGTGCTGCATGGCTGTCGTCAGCTCG
TGTCG-TGAGATG--TTGGGTTAAGTCCCGCAAC----------------------------------------------
--------------------------------------------------------------------------------
```

----------G------AGCGCAACCCTTGTCCTTAGTTGCCAG---CACGTA--AT---------GGTGGGAACTCTAA
GGAGACCGC----CGGTGACA-AACCGGAGGAAGGTGGGGATGACGTCAAGTCA-TCATGG-CCCTTACGACCAG-GGCT
ACACACGTACTACAATGGTAGGGACAGAGGGCTG-CAAACCCG-------------------------------------
----------------------------------------CGAGGGTAAGCCAATCCC-AGAAACCCTATCTCAGTCCGG
ATTGGAGTCTGCAACTCGACTCCATGAAG--TCGGAATCGCTAGTAATCGCAGATCAGCATTGCTGCGGTGAATACGTTC
CCGGGCCTTGTACACACCGCCCGTCACACCATGGGAGTTTGTTGCACCAGAAGCAGGTAGCTTAACCTTCGG--------
-------------------------GAGGGCGCTTGCCACGGTGTGGCCGATGACTGGGGTGAAGTCGTAACAAGGTAGC
CGTATCGGAAGGTGCGGCTGGATCACCTCCTTT-----------------------------------------------
------------------------------------------------------
>lcl|CP001825.1_rrna_39 [locus_tag=Tter_R0039] [product=16S ribosomal RNA]
[location=complement(1140248..1141740)] [gbkey=rRNA] Tht
--------------------------------------------------------------------------------
--------------------------------------------------------------------------------
--------------------------------------------------------------------------------
--------------------------------------------------------------------------------
--------------------------------------------------------------------------------
--------------------------------------------------------------------------------
--------------------------------------------------------------------------------
--------------------------------------------------------------------------------
-------------------------------------------------------AGAGTTTGATCCTGGCTCAG-GATG
AACGCTGGCGGCGTGCCTAACGCATGCAAGTCGTGCGGGTAGACCTTC--------------------------GGGTCT
GCCAGCGGCGAACGGGTGAGTAACA-CGTGGGTAACC-------------------------------------------
-----TGCCGTCTGG--TGGGGGATAACCCTCAGAAA-TGGGGGCTAATACCGCAT-GATGATCCTCGCTCGATG-----
------------------------------------------------GAGTGGGGATTGAAAGCC---GCAAGGCGCCA
GGCGAGGGGCCCGCGGCCCATCAGCTAGTTGGTGGGGT-AATGGCCCACCAAGGCTATGACGGGTAGCTGGTCTGAGAGG
ACGGCCAGCCACACTGGGACTGAGACACGGCCCAGACTCCTACGGG--AGGCAGCAGCGAGGAATCTTCCCCAATGGCCC
ATCTGGGGCTGAGGGAG---C-----------GACGCCGCGTGGAGGAAGAAGCCCTTCGG-GGTGTAAACTCCTTTTGG
GTGGGAAGA---------------------------TGATGACGGTACCACCCGAATAAGTCCCGGCTAACTACGTGCCA
GCAGCCGCGGTAACACGTAGGGGGCGAGCGTTATCCGGATTCACTGGGCGTAAAGAGCGCGTAGGCGGTTCCGCAAGTCA
CCCGTGAAAGCCCCCGGCTCAACCGGGG-AGGG----------------------------------CCGGGTGATACT-
-----------------------------------GTGGGGCTTGAGGGCGGAAGAGGGAAGCGGAATTCCCGGTGTAGC
GGTGAAATGCGTAGATACCGGGAGGA------------------------------------------------------
--------------------------------------------------------------------------------
-----------------------------------ACACCAGTGGCGAAGGCGGCTTCCTGGTCCGTACCTGACGCTGAG
GCGCGAAGGCTAGGGGAGCGAACGGGATTAGATACCCCGGTAGTCCTAGCAGTAAAC-----------------------
--------------------------------------------------------------------------------
---------GATGGACGCTCGGTGTCGGCGG---TAT-CCAC---TCCGTCGGTGCCCAAGCTAACGCATTAAGCGTCCC
GCCTGGGGAGTACGGCCGCAAGGCTAAAACTCAA-AGGAATTGACGGGGGCCCGCACA-AGCAGCGGAGCG-TGTGGTTT
AATTCGATGCAACGCGAAGAACCTTACCAGGGCTTGACATGCAGGTGAAAGCCTGGGGAAACTCA---------------
---------------------------GGCCCCCACTTCGGTGGCACCTGCACAGGTGTTGCATGGCTGTCGTCAGCTCG
TGCCG-TGAGGTG--TTGGGTTAAGTCCCGCAAC----------------------------------------------
--------------------------------------------------------------------------------
----------G------AGCGCAACCCTCGTCGCAAGTTACC------------------------------CTCTCTTG
CGAGACTGC----CGGGTGCA-ACCCGGAGGAAGGTGGGGATGACGTCAAGTCA-GCATGA-CCCTTATGCCCTG-GGCT
ACACACACGCTACAATGGCCGGTACAGAGGGTGG-CCAACCCG-------------------------------------
----------------------------------------CGAGGGGGAGCTAATCCC-ACAAAGCCGGTCTCAGTTGGA
ATTAGAGGCTGAAACCCGCCTCTATGAAC--GCGGAGTTGCTAGTAACCGCAGGTCAGCACTACTGCGGTGAATATGTTC
CCGGGCCTTGTACACACCGCCCGTCACGTCACGAAAGCCGGCAACACCTGAAGCCGGTGGGCCAACCCGCAA--------
-----------------------GGGAGGCAGCCGTCTAGGGTGGGGCTGGTGATTGGGACGAAGTCGTAACAAGGTAGC
CGTACCGGAAGGTGCGGCTGGATCACCT----------------------------------------------------
------------------------------------------------------
>lcl|CP015519.1_rrna_47 [locus_tag=A7E78_12805] [product=16S ribosomal RNA] [location=2832200..2833758]
[gbkey=rRNA] Pel
--------------------------------------------------------------------------------
--------------------------------------------------------------------------------
--------------------------------------------------------------------------------
--------------------------------------------------------------------------------
--------------------------------------------------------------------------------
--------------------------------------------------------------------------------
--------------------------------------------------------------------------------
--------------------------------------------------------------------------------
----------------------------------------------AA---CTGGAGAGTTTGATCCTGGCTCAG-AACG
AACGCTGGCGGCGTGCTTAACACATGCAAGTCGAACGAGAAAGGG---ACTTCGG---T---------------CCTGAG
TAGAGTGGCGCACGGGTGAGTAACA-CGTGGATAATC-------------------------------------------
-----TACCCAATGA--TCTGGGATAACACTTCGAAA-GGGGTGCTAATACCGGAT-AAGCCCACAGGCTCTTCGGAGCT
TGCG---G-------------------------------GAA------AA-GGTGGGGACCTTCGG---GCCTACCGTCA
TCGGATGAGTCCGCGGCCCATTAGCTAGTTGGTAGGGT-AACGGCCTACCAAGGCGACGATGGGTAGCTGGTCTGAGAGG
ATGATCAGCCACACTGGAACTGAGACACGGTCCAGACTCCTACGGG--AGGCAGCAGTGGGGAATTTTGCGCAATGGGCG
AAAGCCTGAC---GCAG---C-----------AACGCCGCGTGAGTGATGAAGGCTTTCGG-GTCGTAAAGCTCTGTCAG
TGGGGAAGAAAC-TCTTGTGGTCTAATATGCCGCAAGACTGACGGTACCCACAAAGGAAGCACCGGCTAACTCCGTGCCA
GCAGCCGCGGTAATACGGAGGGTGCAAGCGTTGTTCGGAATTATTGGGCGTAAAGCGCGTGTAGGCGGTCTATTAAGTCT
GATGTGAAAGCCCCGGGCTCAACCCGGG-AAGT----------------------------------GCATTGGATACT-
-----------------------------------GGTAGACTTGAGTACGGGAGAGGGAAGTGGAATTCCGAGTGTAGG
AGTGAAATCCGTAGATATTCGGAGGA------------------------------------------------------
--------------------------------------------------------------------------------
-----------------------------------ACACCGGTGGCGAAGGCGGCTTCCTGGACCGATACTGACGCTGAG
ACGCGAAAGCGTGGGGAGCAAACAGGATTAGATACCCTGGTAGTCCACGCCGTAAAC-----------------------
--------------------------------------------------------------------------------
---------GATGGGTACTAGGTGTCGCGGG---TAT-TGAC---CCCTGCGGTGCCGAAGCTAACGCATTAAGTACCCC
GCCTGGGGAGTACGGCCGCAAGGCTAAAACTCAA-AGGAATTGACGGGGGCCCGCACA-AGCGGTGGAGCA-TGTGGTTT
AATTCGACGCAACGCGTAGAACCTTACCTGGGTTTGACATCCCGATCGCACTTTCTGGAAACAGT---------------
---------------------------TAGGTCAGTTCGGCTGGATCGGTGACAGGTGCTGCATGGCTGTCGTCAGCTCG
TGTCG-TGAGATG--TTGGGTTAAGTCCCGCAAC----------------------------------------------
--------------------------------------------------------------------------------
----------G------AGCGCAACCCTTGTCCTTAGTTGCCAT---CAT--T--AA---------GTTGGGCACTCTAG
GGAGACTGC----CGGTGTTA-AACCGGAGGAAGGTGGGGATGACGTCAAGTCC-TCATGG-CCCTTATATCCAG-GGCT
ACACACGTGCTACAATGGCCGGTACAAAGGGCAG-CTATACCG-------------------------------------

----------------------------------------CAAGGTGGAGCGAATCCC-AGAAAGCCGGTCTCAGTTCGG
ATTGGAGTCTGCAACTCGACTCCATGAAG--TTGGAATCGCTAGTAATCGCGGATCAGCA-TGCCGCGGTGAATACGTTC
CCGGGCCTTGTACACACCGCCCGTCACACCACGGGAGTTGATTGTACCTGAAATCGGTGGGCTAACCTTC-G--------
------------------------GGAGGCAGCCGCTTATGGTATGATCGGTAACTGGGGTGAAGTCGTAACAAGGTAGC
CGTAGGGGAACCTGCGGCTGGATCACCTCCTTTCTAA-------------------------------------------
------------------------------------------------------
>lcl|AL009126.3_rrna_1 [gene=rrnO-16S] [locus_tag=BSU_rRNA_1] [product=ribosomal RNA-16S] [location=9810..11364]
[gbkey=rRNA] Bsu
--------------------------------------------------------------------------------
--------------------------------------------------------------------------------
--------------------------------------------------------------------------------
--------------------------------------------------------------------------------
--------------------------------------------------------------------------------
--------------------------------------------------------------------------------
--------------------------------------------------------------------------------
--------------------------------------------------------------------------------
-------------------------------------------ATTTA---TCGGAGAGTTTGATCCTGGCTCAG-GACG
AACGCTGGCGGCGTGCCTAATACATGCAAGTCGAGCGGACAGATGGGAGCTTGCT---C---------------CCTGAT
GTTAGCGGCGGACGGGTGAGTAACA-CGTGGGTAACC-------------------------------------------
-----TGCCTGTAAG--ACTGGGATAACTCCGGGAAA-CCGGGGCTAATACCGGAT-GGTTGTTTGAACCGCATGGTTCA
AACA---T-------------------------------AAA--------------AGGTGGCTTC---GGCTACCACTT
ACAGATGGACCCGCGGCGCATTAGCTAGTTGGTGAGGT-AACGGCTCACCAAGGCGACGATGCGTAGCCGACCTGAGAGG
GTGATCGGCCACACTGGGACTGAGACACGGCCCAGACTCCTACGGG--AGGCAGCAGTAGGGAATCTTCCGCAATGGACG
AAAGTCTGAC---GGAG---C-----------AACGCCGCGTGAGTGATGAAGGTTTTCGG-ATCGTAAAGCTCTGTTGT
TAGGGAAGAACA-AGTGCCGTTCGAATAGGGCGGTACCTTGACGGTACCTAACCAGAAAGCCACGGCTAACTACGTGCCA
GCAGCCGCGGTAATACGTAGGTGGCAAGCGTTGTCCGGAATTATTGGGCGTAAAGGGCTCGCAGGCGGTTTCTTAAGTCT
GATGTGAAAGCCCCCGGCTCAACCGGGG-AGGG---------------------------------TCATTGGAAACT-
-----------------------------------GGGGAACTTGAGTGCAGAAGAGGAGAGTGGAATTCCACGTGTAGC
GGTGAAATGCGTAGAGATGTGGAGGA------------------------------------------------------
--------------------------------------------------------------------------------
-----------------------------------ACACCAGTGGCGAAGGCGACTCTCTGGTCTGTAACTGACGCTGAG
GAGCGAAAGCGTGGGGAGCGAACAGGATTAGATACCCTGGTAGTCCACGCCGTAAAC-----------------------
--------------------------------------------------------------------------------
---------GATGAGTGCTAAGTGTTAGGGG---GTT-TCCG---CCCCTTAGTGCTGCAGCTAACGCATTAAGCACTCC
GCCTGGGGAGTACGGTCGCAAGACTGAAACTCAA-AGGAATTGACGGGGGCCCGCACA-AGCGGTGGAGCA-TGTGGTTT
AATTCGAAGCAACGCGAAGAACCTTACCAGGTCTTGACATCCTCTGACAATCCTAGAGA---TAG---------------
---------------------------GACGTCCCCTTCGGGGGCAGAGTGACAGGTGGTGCATGGTTGTCGTCAGCTCG
TGTCG-TGAGATG--TTGGGTTAAGTCCCGCAAC----------------------------------------------
--------------------------------------------------------------------------------
----------G------AGCGCAACCCTTGATCTTAGTTGCCAG---CAT--T--CA---------GTTGGGCACTCTAA
GGTGACTGC----CGGTGACA-AACCGGAGGAAGGTGGGGATGACGTCAAATCA-TCATGC-CCCTTATGACCTG-GGCT
ACACACGTGCTACAATGGACAGAACAAAGGGCAG-CGAAACCG-------------------------------------
----------------------------------------CGAGGTTAAGCCAATCCC-ACAAATCTGTTCTCAGTTCGG
ATCGCAGTCTGCAACTCGACTGCGTGAAG--CTGGAATCGCTAGTAATCGCGGATCAGCA-TGCCGCGGTGAATACGTTC
CCGGGCCTTGTACACACCGCCCGTCACACCACGAGAGTTTGTAACACCCGAAGTCGGTGAGGTAACCTTTTA--------
------------------------GGAGCCAGCCGCCGAAGGTGGGACAGATGATTGGGGTGAAGTCGTAACAAGGTAGC
CGTATCGGAAGGTGCGGCTGGATCACCTCCTTTCTA--------------------------------------------
------------------------------------------------------
>lcl|AE007317.1_rrna_53 [locus_tag=sprr09] [product=16S ribosomal RNA] [location=complement(1793205..1794718)]
[gbkey=rRNA] Spn
--------------------------------------------------------------------------------
--------------------------------------------------------------------------------
--------------------------------------------------------------------------------
--------------------------------------------------------------------------------
--------------------------------------------------------------------------------
--------------------------------------------------------------------------------
--------------------------------------------------------------------------------
--------------------------------------------------------------------------------
----------------------------------------------------------GTTTGATCCTGGCTCAG-GACG
AACGCTGGCGGCGTGCCTAATACATGCAAGTAGAACGCTGAAGGAGGAGCTTGCT---T---------------CTCTGG
ATGAGTTGCGAACGGGTGAGTAACG-CGTAGGTAACC-------------------------------------------
-----TGCCTGGTAG--CGGGGGATAACTATTGGAAA-CGATAGCTAATACCGCAT-AAGAGTGGATGTTGCATGACATT
TGCT---T-------------------------------AAA--------------AGGTGC--AC---TTGCATCACTA
CCAGATGGACCTGCGTTGTATTAGCTAGTTGGTGGGGT-AACGGCTCACCAAGGCGACGATACATAGCCGACCTGAGAGG
GTGATCGGCCACACTGGGACTGAGACACGGCCCAGACTCCTACGGG--AGGCAGCAGTAGGGAATCTTCGGCAATGGACG
GAAGTCTGAC---CGAG---C-----------AACGCCGCGTGAGTGAAGAAGGTTTTCGG-ATCGTAAAGCTCTGTTGT
AAGAGAAGAACG-AGTGTGAGAGTGGAAAGTTCACACTGTGACGGTATCTTACCAGAAAGGGACGGCTAACTACGTGCCA
GCAGCCGCGGTAATACGTAGGTCCCGAGCGTTGTCCGGATTTATTGGGCGTAAAGCGAGCGCAGGCGGTTAGATAAGTCT
GAAGTTAAAGGCTGTGGCTTAACCATAG--TAG---------------------------------GCTTTGGAAACT-
-----------------------------------GTTTAACTTGAGTGCAAGAGGGGAGAGTGGAATTCCATGTGTAGC
GGTGAAATGCGTAGATATATGGAGGA------------------------------------------------------
--------------------------------------------------------------------------------
-----------------------------------ACACCGGTGGCGAAAGCGGCTCTCTGGCTTGTAACTGACGCTGAG
GCTCGAAAGCGTGGGGAGCAAACAGGATTAGATACCCTGGTAGTCCACGCTGTAAAC-----------------------
--------------------------------------------------------------------------------
---------GATGAGTGCTAGGTGTTAGACC---CTT-TCCG---GGGTTTAGTGCCGTAGCTAACGCATTAAGCACTCC
GCCTGGGGAGTACGACCGCAAGGTTGAAACTCAA-AGGAATTGACGGGGGCCCGCACA-AGCGGTGGAGCA-TGTGGTTT
AATTCGAAGCAACGCGAAGAACCTTACCAGGTCTTGACATCCCTCTGACCGCTCTAGAG---ATA---------------
---------------------------GAGTTTTCCTTCGGGACAGAGGTGACAGGTGGTGCATGGTTGTCGTCAGCTCG
TGTCG-TGAGATG--TTGGGTTAAGTCCCGCAAC----------------------------------------------
--------------------------------------------------------------------------------
----------G------AGCGCAACCCCTATTGTTAGTTGCCAT---CAT--T--CA---------GTTGGGCACTCTAG
CGAGACTGC----CGGTAATA-AACCGGAGGAAGGTGGGGATGACGTCAAATCA-TCATGC-CCCTTATGACCTG-GGCT
ACACACGTGCTACAATGGCTGGTACAACGAGTCG-CAAGCCGG-------------------------------------
----------------------------------------TGACGGCAAGCTAATCTC-TTAAAGCCAGTCTCAGTTCGG
ATTGTAGGCTGCAACTCGCCTACATGAAG--TCGGAATCGCTAGTAATCGCGGATCAGCA-CGCCGCGGTGAATACGTTC
CCGGGCCTTGTACACACCGCCCGTCACACCACGAGAGTTTGTAACACCCGAAGTCGGTGAGGTAACCGTA-A--------

```
------------------------GGAGCCAGCCGCCTAAGGTGGATAGATGATTGGGGTGAAGTCGTAACAAGGTAGC
CGTATCGGAAG---------------------------------------------------------------------
------------------------------------------------------
>lcl|AE014295.3_rrna_2 [locus_tag=BLr01] [product=16S ribosomal RNA] [location=159243..160772] [gbkey=rRNA] Blo
--------------------------------------------------------------------------------
--------------------------------------------------------------------------------
--------------------------------------------------------------------------------
--------------------------------------------------------------------------------
--------------------------------------------------------------------------------
--------------------------------------------------------------------------------
--------------------------------------------------------------------------------
--------------------------------------------------------------------------------
-------------------------------------------TTTTT---GTGGAGGGTTCGATTCTGGCTCAG-GATG
AACGCTGGCGGCGTGCTTAACACATGCAAGTCGAACGGGATCCATCAGGCTTTGC---TT--------------GGTGGT
GAGAGTGGCGAACGGGTGAGTAATG-CGTGACCGACC-------------------------------------------
-----TGCCCCATAC--ACCGGAATAGCTCCTGGAAA-CGGGTGGTAATGCCGGAT-GCTCCAGTTGATCGC--------
----------------------------------------AT------GG---TCTTCTGGGAAAG---CTTTCGCGGTA
TGGGATGGGGTCGCGTCCTATCAGCTTGACGGCGGGT-AACGGCCCACCGTGGCTTCGACGGGTAGCCGGCCTGAGAGG
GCGACCGGCCACATTGGGACTGAGATACGGCCCAGACTCCTACGGG--AGGCAGCAGTGGGGAATATTGCACAATGGGCG
CAAGCCTGAT---GCAG---C-----------GACGCCGCGTGAGGATGGAGGCCTTCGG-GTTGTAAACCTCTTTTAT
CGGGGAGCAAGC-GAG---------------------AGTGAGTTTACCCGTTGAATAAGCACCGGCTAACTACGTGCCA
GCAGCCGCGGTAATACGTAGGGTGCAAGCGTTATCCGGAATTATTGGGCGTAAAGGGCTCGTAGGCGGTTCGTCGCGTCC
GGTGTGAAAGTCCATCGCTTAACGGTGG-ATCC----------------------------------GCGCCGGGTACG-
-----------------------------------GGCGGGCTTGAGTGCGGTAGGGGAGACTGGAATTCCCGGTGTAAC
GGTGGAATGTGTAGATATCGGGAAGA------------------------------------------------------
--------------------------------------------------------------------------------
-----------------------------------ACACCAATGGCGAAGGCAGGTCTCTGGGCCGTTACTGACGCTGAG
GAGCGAAAGCGTGGGGAGCGAACAGGATTAGATACCCTGGTAGTCCACGCCGTAAAC-----------------------
--------------------------------------------------------------------------------
---------GGTGGATGCTGGATGTGGGGCC---CGTTCCAC---GGGTTCCGTGTCGGAGCTAACGCGTTAAGCATCCC
GCCTGGGGAGTACGGCCGCAAGGCTAAAACTCAA-AGAAATTGACGGGGGCCCGCACA-AGCGGCGGAGCA-TGCGGATT
AATTCGATGCAACGCGAAGAACCTTACCTGGGCTTGACATGTTCCCGACGGTCGTAGAGATACGG---------------
-------------------------------CTTCCCTTCGGGGCGGGTTCACAGGTGGTGCATGGTCGTCGTCAGCTCG
TGTCG-TGAGATG--TTGGGTTAAGTCCCGCAAC----------------------------------------------
--------------------------------------------------------------------------------
----------G------AGCGCAACCCTCGCCCCGTGTTGCCAG---CGGATT--AT---------GCCGGGAACTCACG
GGGGACCGC----CGGGGTTA-ACTCGGAGGAAGGTGGGGATGACGTCAGATCA-TCATGC-CCCTTACGTCCAG-GGCT
TCACGCATGCTACAATGGCCGGTACAACGGGATG-CGACGCGG-------------------------------------
----------------------------------------CGACGCGGAGCGGATCCC-TGAAAACCGGTCTCAGTTCGG
ATCGCAGTCTGCAACTCGACTGCGTGAAG--GCGGAGTCGCTAGTAATCGCGAATCAGCAACGTCGCGGTGAATGCGTTC
CCGGGCCTTGTACACACCGCCCGTCAAGTCATGAAAGTGGGCAGCACCCGAAGCCGGTGGCCTAACCCCTTG--------
-------------T---------GGGATGGAGCCGTCTAAGGTGAGGCTCGTGATTGGGACTAAGTCGTAACAAGGTAGC
CGTACCGGAAGGTGCGGCTGGATCACCTCCTTTCT---------------------------------------------
------------------------------------------------------
>lcl|AL123456.3_rrna_30 [gene=rrs] [product=Ribosomal RNA 16S] [location=1471846..1473382] [gbkey=rRNA] Mtu
--------------------------------------------------------------------------------
--------------------------------------------------------------------------------
--------------------------------------------------------------------------------
--------------------------------------------------------------------------------
--------------------------------------------------------------------------------
--------------------------------------------------------------------------------
--------------------------------------------------------------------------------
--------------------------------------------------------------------------------
------------------------------------------TTTTGT---TTGGAGAGTTTGATCCTGGCTCAG-GACG
AACGCTGGCGGCGTGCTTAACACATGCAAGTCGAACGGAAAGGTCTCTTC--------GG--------------AGATAC
TCGAGTGGCGAACGGGTGAGTAACA-CGTGGGTGATC-------------------------------------------
-----TGCCCTGCAC--TTCGGGATAAGCCTGGGAAA-CTGGGTCTAATACCGGAT-AGGACCACGGGATGC--------
----------------------------------------AT------GTCTTGTGGTGGAAAGCG---CTTTAGCGGTG
TGGGATGAGCCCGCGGCCTATCAGCTTGTTGGTGGGGT-GACGGCCTACCAAGGCGACGACGGGTAGCCGGCCTGAGAGG
GTGTCCGGCCACACTGGGACTGAGATACGGCCCAGACTCCTACGGG--AGGCAGCAGTGGGGAATATTGCACAATGGGCG
CAAGCCTGAT---GCAG---C-----------GACGCCGCGTGGGGGATGACGGCCTTCGG-GTTGTAAACCTCTTTCAC
CATCGACGAAGG-TCCGGGTTCTCT---------CGGATTGACGGTAGGTGGAGAAGAAGCACCGGCCAACTACGTGCCA
GCAGCCGCGGTAATACGTAGGGTGCGAGCGTTGTCCGGAATTACTGGGCGTAAAGAGCTCGTAGGTGGTTTGTCGCGTTG
TTCGTGAAATCTCACGGCTTAACTGTGA-GCGT----------------------------------GCGGGCGATACG-
-----------------------------------GGCAGACTAGAGTACTGCAGGGGAGACTGGAATTCCTGGTGTAGC
GGTGGAATGCGCAGATATCAGGAGGA------------------------------------------------------
--------------------------------------------------------------------------------
-----------------------------------ACACCGGTGGCGAAGGCGGGTCTCTGGGCAGTAACTGACGCTGAG
GAGCGAAAGCGTGGGGAGCGAACAGGATTAGATACCCTGGTAGTCCACGCCGTAAAC-----------------------
--------------------------------------------------------------------------------
---------GGTGGGTACTAGGTGTGGGTTT---CCTTCCTT---GGGATCCGTGCCGTAGCTAACGCATTAAGTACCCC
GCCTGGGGAGTACGGCCGCAAGGCTAAAACTCAA-AGGAATTGACGGGGGCCCGCACA-AGCGGCGGAGCA-TGTGGATT
AATTCGATGCAACGCGAAGAACCTTACCTGGGTTTGACATGCACAGGACGCGTCTAGAGATAGGC---------------
-------------------------------GTTCCCTTGT-GGCCTGTGTGCAGGTGGTGCATGGCTGTCGTCAGCTCG
TGTCG-TGAGATG--TTGGGTTAAGTCCCGCAAC----------------------------------------------
--------------------------------------------------------------------------------
----------G------AGCGCAACCCTTGTCTCATGTTGCCAG---CACGTA--AT---------GGTGGGGACTCGTG
AGAGACTGC----CGGGGTCA-ACTCGGAGGAAGGTGGGGATGACGTCAAGTCA-TCATGC-CCCTTATGTCCAG-GGCT
TCACACATGCTACAATGGCCGGTACAAAGGGCTG-CGATGCCG-------------------------------------
----------------------------------------CGAGGTTAAGCGAATCCT-TAAAAGCCGGTCTCAGTTCGG
ATCGGGGTCTGCAACTCGACCCCGTGAAG--TCGGAGTCGCTAGTAATCGCAGATCAGCAACGCTGCGGTGAATACGTTC
CCGGGCCTTGTACACACCGCCCGTCACGTCATGAAAGTCGGTAACACCCGAAGCCAGTGGCCTAACCCTCGG--------
-------------G------------AGGGAGCTGTCGAAGGTGGGATCGGCGATTGGGACGAAGTCGTAACAAGGTAGC
CGTACCGGAAGGTGCGGCTGGATCACCTCCTTTCT---------------------------------------------
------------------------------------------------------
>lcl|CP016757.1_rrna_7 [locus_tag=BED41_02870] [product=16S ribosomal RNA] [location=633960..635480]
[gbkey=rRNA] Cpo
```

--------------------------------------------------------------------------------
--------------------------------------------------------------------------------
--------------------------------------------------------------------------------
--------------------------------------------------------------------------------
--------------------------------------------------------------------------------
--------------------------------------------------------------------------------
--------------------------------------------------------------------------------
--------------------------------------------------------------------------------
-----------------------------------------------A---ACGGAGAGTTTGATCCTGGCTCAG-GACG
AACGCTGGCGGCGTGCCTAACACATGCAAGTCGAACGAAATTACGAAGAAAGCTT---GCT---------TTAATCGTAA
TTTAGTGGCGGACGGGTGAGTAACG-CGTGAGGACTT-------------------------------------------
-----GTCG-GATAC--AGGGGGACAACAGATGGAAA-CGTCTGCTAATACCCCAT-AAGC-------------------
---------------------------------------------------CTTCGGGTAAAAGGA---GCAATCCGGTA
TCTGAGAGACTCGCGTTCTATCAGCTAGTAGGTGGGGT-AACGGCCCACCTAGGCCAAGACGGATAGCCGGACTGAGAGG
TCGACCGGCCACACTGGAACTGAGAGACGGTCCAGACTCCTACGGG--AGGCAGCAGTGGGGAATATTGGGCAATGGGGG
CAACCCTGAC---CCAG---C-----------GACGCCGCGTGGGTGAAGAAATCCTTCGG-GATGTAAAGCCCTGTTGT
GTGGGAAGAT---------------------------AATGACGGTACCACACGAGGAAGCCCCGGCAAACTACGTGCCA
GCAGCCGCGGTAATACGTAGGGGGCGAGCGTTGTTCGGAATTACTGGGCGTAAAGCGCACGCAGGCTGACCGTTAAGTCT
GTCGTCAAAGGCGGAGGCTCAACCTTCG-TTCT----------------------------------ACGATAGATACT-
-----------------------------------GGCGGTCTAGAGTATGTGAGAGGGAAGTGGAATTCCCGGTGTAGC
GGTGAAATGCGTAGATATCGGGAGGA------------------------------------------------------
--------------------------------------------------------------------------------
-----------------------------------ACACCAGTGGCGAAGGCGGCTTCCTGGCACACAACTGACGCTCAT
GTGCGAAAGCCAGGGCAGCGAACGGGATTAGATACCCCGGTAGTCCTGGCCGTAAAC-----------------------
--------------------------------------------------------------------------------
---------GATGGATACTGGGTGTGGGTGA---A---GCAG---TTCATCCGTGCCGTAGTTAACGCGTTAAGTATCCC
GCCTGGGGACTACGGTCGCAAGACTGAAACTCAA-AGGAATTGACGGGGGCCCGCACA-AGCGGTGGAGCA-CGTGGTTT
AATTCGATGCAAACCGAAGAACCTTACCTGGGTTTGACATACAAGTGGTACTGAGATGAAAGTTG--AG-----------
---------------------G----GACTGTAGCTTGCTACAGAGCTTGAACAGGTGCTGCATGGCTGTCGTCAGCTCG
TGTCG-TGAGATG--TTGGGTTAAGTCCCGCAAC----------------------------------------------
--------------------------------------------------------------------------------
----------G------AGCGCAACCCCTATGTCCAGTTGCTAA---CAAGTG--AA---------GTTGAGCACTCTGG
AGAGACTGC----CGCCGACA-AGGCGGAGGAAGGTGGGGATGACGTCAAGTCA-TCATGG-CCTTTATGCCCAG-GGCG
ACACACGTGCTACAATGGCCGGCACAGAAGGCAG-CTTGCTAG-------------------------------------
----------------------------------------TGATAGTTGGCGAATCCT-TAAAAGCCGGTCCCAGTTCGG
ATTGTAGTCTGCAACCCGACTACATGAAG--CCGGAATCGCTAGTAATCGCAGATCAGCCAAGCTGCGGTGAATACGTTC
CCGGGCCTTGTACACACCGCCCGTCACACCACCCGAGTTGGGGGCACCCGAAGCCGCAGGCTTAACCCGTAA--------
-----------------------GGGAAAGAAGCGTCTAAGGTGCGCCGAGTAAGGGGGGTGAAGTCGTAACAAGGTAGC
CGTACCGGAAGGTGCGGCTGGATCACCTCCTTTCTAAGGA----------------------------------------
------------------------------------------------------
>lcl|LT906446.1_rrna_53 [locus_tag=SAMEA4364220_01665] [product=16S ribosomal RNA] [location=complement(1679668..1681211)] [gbkey=rRNA] Mhy
--------------------------------------------------------------------------------
--------------------------------------------------------------------------------
--------------------------------------------------------------------------------
--------------------------------------------------------------------------------
--------------------------------------------------------------------------------
--------------------------------------------------------------------------------
--------------------------------------------------------------------------------
--------------------------------------------------------------------------------
-------------------------------------------------------AGAGTTTGATCCTGGCTCAG-GACG
AACGCTGGCGGCGTGCTTAACACATGCAAGTCGAACGGGGTGTTTATTTCGGTAA---AC-------------------A
CCAAGTGGCGAACGGGTGAGTAACG-CGTAAGCAATC-------------------------------------------
-----TACCTTCAAG--ATGGGGACAACACTTCGAAA-GGGGTGCTAATACCGAAT-GAATGAGAGATGACCGCATGGAT
ATTT---C-------------------------------TCT------AAAGGAGGCCTCTGAAAA---TGCTTCCGCTT
GAAGATGAGCTTGCGTCTGATTAGCTAGTTGGTGAGGGTAAAGGCCCACCAAGGCGACGATCAGTAGCCGGTCTGAGAGG
ATGAACGGCCACATTGGGACTGAGACACGGCCCAGACTCCTACGGG--AGGCAGCAGTGGGGAATCTTCCGCAATGGGCG
AAAGCCTGAC---GGAG---C-----------AACGCCGCGTGAACGAAGAAGGTCTTAGG-ATCGTAAAGTTCTGTTGT
TAGGGGCGAAGG-GTAGCATTTTGAATAAGGGTGCTATTTGACGGTACTTAACGAGGAAGCCACGGCTAACTACGTGCCA
GCAGCCGCGGTAATACGTAGGCGGCAAGCGTTGTCCGGAATTATTGGGCGTAAAGGGAGCGCAGGCGGGAAGTTAAGCGG
ACTTTAAAAGTGCGGGGCTCAACCC-CG-TGAG----------------------------------GGGGTCCGAACT-
-----------------------------------GACTTTCTTGAGTGCAGGAGAGGGAAGCGGAATTCCTAGTGTAGC
GGTGAAATGCGTAGATATTAGGAAGA------------------------------------------------------
--------------------------------------------------------------------------------
-----------------------------------ACACCAGTGGCGAAGGCGGCTTTCTGGACTGTAACTGACGCTGAG
GCTCGAAAGCTAGGGTAGCGAACGGGATTAGATACCCCGGTAGTCCTAGCCGTAAAC-----------------------
--------------------------------------------------------------------------------
---------GATGGATACTAGGTGTGGGAGG---TAT-CGAC---CCCTTCCGTGCCGGAGTTAACGCAATAAGTATCCC
GCCTGGGGAGTACGGCCGCAAGGTTGAAACTCAA-AGGAATTGACGGGGGCCCGCACA-AGCGGTGGAGTA-TGTGGTTT
AATTCGACGCAACGCGAAGAACCTTACCAAGACTTGACATCGACTGACGTACTTAGAGAT-------AA-----------
---------------------G----TATTTTTACTTCGGTAAACAGGAAGACAGGTGGTGCATGGCTGTCGTCAGCTCG
TGTCG-TGAGATG--TTGGGTTAAGTCCCGCAAC----------------------------------------------
--------------------------------------------------------------------------------
----------G------AGCGCAACCCCTATACTATGTTGCCAG---CACGTA--AA---------GGTGGGAACTCATA
GTAGACTGC----CGCGGACA-ACGCGGAGGAAGGCGGGGATGACGTCAAGTCA-TCATGC-CCCTTACGTCTTG-GGCT
ACACACGTACTACAATGGGATGAACAGAGGGAAG-CGAAGTCG-------------------------------------
----------------------------------------CGAGGCAGAGCGGAACCC-TAAAAGCATCTCTCAGTTCGG
ATTGCAGGCTGAAACTCGCCTGCATGAAG--TCGGAATCGCTAGTAATCGCAGGTCAGCA-TACTGCGGTGAATACGTTC
CCGGGCCTTGTACACACCGCCCGTCACACCACGAAAGTCATTCACACCCGAAGCCGGCTAAGGG----CCGA--------
-------------A---------AGGAACCGACCGTCGAAGGTGGGGGCGATGATTGGGGTGAAGTCGTAACAAGGTAGC
CGTATCGGAAGGTGCGGCTGGATCACCT----------------------------------------------------
------------------------------------------------------
>lcl|CP035282.1_rrna_1 [locus_tag=EQM13_00055] [product=16S ribosomal RNA] [location=10436..11969] [gbkey=rRNA] Ssp
--------------------------------------------------------------------------------
--------------------------------------------------------------------------------
--------------------------------------------------------------------------------

--------------------------------------------------------------------------------
--------------------------------------------------------------------------------
--------------------------------------------------------------------------------
--------------------------------------------------------------------------------
--------------------------------------------------------------------------------
-------------------------------------------TTTAA---ATTGAGAGTTTGATCCTGGCTCAG-GACG
AACGCTGGCGGCGTGCTTAACACATGCAAGTCGAGCGAGGGAATACTTAGGAAGC---CTTCGGGAA-GAATACGTATTC
CTTAGCGGCGGACGGGTGAGTAACG-CGTGAGGAATC-------------------------------------------
-----TGCCTCGCAC--AAGGGGATAGCCTCGGGAAA-CCGGGATTAATACCCTAT-GATACTTACACTGC---------
-------G-------------------------------CAT------GCAG---AGTAAGTTAAA---GAATATCGGTG
TGAGATGACCTCGCGTCCCATTAGCTGGTTGGCAGGGT-AAAAGCCTACCAAGGCGACGATGGGTAGCCGGCCTGAGAGG
GTGAACGGCCACATTGGAACTGAGACACGGTCCAGACTCCTACGGG--AGGCAGCAGTGGGGAATATTGCACAATGGAGG
GAACTCTGAT---GCAG---C-----------AACGCCGCGTGAGCGAAGAAGGTTTTAGG-ATCGTAAAGCTCTGTCCT
AGGGGAAGAT---------------------------AATGACGGTACCCGAGGAGGAAGCCCCGGCTAACTACGTGCCA
GCAGCCGCGGTAATACGTAGGGGGCGAGCGTTGTCCGGAATTATTGGGCGTAAAGGGTGCGTAGGCGGCTGAACAAGTCA
GATGTGAAAGGTTAAGCTTAACTT-TA-ATAA-----------------------------------GCATATGAAACT-
-----------------------------------GTTCAGCTTGAGTTAAGGAGAGGGGAGTGGAATTCCTAGTGTAGC
GGTGAAATGCGTAGATATTAGGAGGA------------------------------------------------------
--------------------------------------------------------------------------------
-----------------------------------ATACCAGTGGCGAAGGCGACTCTCTGGACTTAAACTGACGCTGAG
GCACGAAAGCGTGGGGAGCAAACAGGATTAGATACCCTGGTAGTCCACGCCGTAAAC-----------------------
--------------------------------------------------------------------------------
---------GATGAGTGCTAGGTGTTGGGTG---TC-----A---AAGCTCAGTGCCGCAGCTAACGCAATAAGCACTCC
GCCTGGGGAGTACGATGGCAACATTGAAACTCAA-AGGAATTGACGGGGACCCGCACA-AGCAGCGGAGCA-TGTGGTTT
AATTCGATGCAACGCGAAGAACCTTACCAAGGCTTGACATCCCTCTGAAGTATATAGAGATATATACGG-----------
---------------------C----T-------TTTAGGAGACAGAGGAGACAGGTGGTGCATGGTTGTCGTCAGCTCG
TGTCG-TGAGATG--TTGGGTTAAGTCCCGCAAC----------------------------------------------
--------------------------------------------------------------------------------
----------G------AGCGCAACCCTTACTTTTAGTTACTAA---CAGGTA--AA---------GCTGAGGACTCTAA
AAGGACTGC----CGGTGATA-AATCGGAGGAAGGTGGGGATGACGTCAAATCA-TCATGC-CCTTTATGTCTTG-GGCT
ACACACGTGCTACAATGGTCGGTACAGAGGGAAG-CGACGCCG-------------------------------------
----------------------------------------CAAGGTGAAGCAAATCCC-AAAAAGCCGATCCCAGTTCGG
ATTGCAGGCTGCAACCCGCCTGCATGAAG--CTGGAGTTGCTAGTAATCGCAGATCAGCA-TGCTGCGGTGAATGCGTTC
CCGGGTCTTGTACACACCGCCCGTCACACCATGGGAGTTGACAATACCCGAAGCCGGTAAGCGAACCCGAAA--------
-----------------------GGGGAGCAGCCGTCGAAGGTAGGGTTAATGACTGGGGTGAAGTCGTAACAAGGTAGC
CGTATCGGAAGGTGCGGCTGGATCACCTCCTTTCT---------------------------------------------
------------------------------------------------------
>lcl|AE008691.1_rrna_6 [locus_tag=TTEr01] [product=16S ribosomal RNA] [location=53858..55384] [gbkey=rRNA] Tte
--------------------------------------------------------------------------------
--------------------------------------------------------------------------------
--------------------------------------------------------------------------------
--------------------------------------------------------------------------------
--------------------------------------------------------------------------------
--------------------------------------------------------------------------------
--------------------------------------------------------------------------------
--------------------------------------------------------------------------------
------------------------------------------------------GAGAGTTTGATCCTGGCTCAG-GACG
AACGCTGGCGGCGTGCCTAACACATGCAAGTCGAGCGGGCCTGTCAAGGCGTTAT---GT------T-TT----GGCAGG
TTAAGCGGCGGACGGGTGAGTAACG-CGTGAGCAACC-------------------------------------------
-----TACCCTTGAG--ACAGGGATAACCCTGGGAAA-CCGGGGCTAATACCTGAT-ATACTCTATTAGGG---------
-------C-------------------------------ATC------CTGATGGAGGAAAGGGCG---GGAAGCCGCTC
AAGGATGGGCTCGCGTCCCATCAGCTAGTTGGTGAGGT-AACGGCTCACCAAGGCTACGACGGGTAGCCGGCCTGAGAGG
GTGGTCGGCCACACTGGGACTGAGACACGGCCCAGACTCCTACGGG--AGGCAGCAGTGGGGAATCTTCCGCAATGGGCG
AAAGCCTGAC---GGAG---C-----------GACGCCGCGTGAGCGAAGAAGGCCTTCGG-GTCGTAAAGCTCGATAGT
GTGGGAAGAAGG-G-----------------------ATGACG---GTACCACACGAAAGCCCCGGCTAACTACGTGCCA
GCAGCCGCGGTAAGACGTAGGGGGCGAGCGTTGTCCGGAGTTACTGGGCGTAAAGGGCGCGTAGGCGGTTTAGCAAGTCA
GGTGTAAAAGGCCACGGCTCAACCGTGG-ATAT----------------------------------GCATCTGAAACT-
-----------------------------------GCTGAGCTAGAGGGCAGGAGAGGGGAGTGGAATTCCCGGTGTAGC
GGTGAAATGCGTAGATATCGGGAGGA------------------------------------------------------
--------------------------------------------------------------------------------
-----------------------------------ATACCAGTGGCGAAGGCGACTCCCTGGACTGGCCCTGACGCTGAG
GCGCGAGAGCGTGGGGAGCAAACAGGATTAGATACCCTGGTAGTCCACGCTGTAAAC-----------------------
--------------------------------------------------------------------------------
---------GATGGGTGCTAGGTGTGGGGAG---C---GGAA---GCTTTCCGTGCCGTAGGAAACCCAATAAGCACCCC
GCCTGGGGAGTACGGCCGCAAGGCTGAAACTCAA-AGGAATTGACGGGGGCCCGCACA-AGCGGTGGAGCA-TGTGGTTT
AATTCGAAGCAACGCGAAGAACCTTACCAGGGCTTGACATGCAGGTAGTAGCGAGCCGAAAGGTGAGCG-----------
---------------------A----CCCTTTCTTATGGGAGGGAGCCTGCACAGGTGGTGCATGGTTGTCGTCAGCTCG
TGTCG-TGAGATG--TTGGGTTAAGTCCCGCAAC----------------------------------------------
--------------------------------------------------------------------------------
----------G------AGCGCAACCCCTGCCTCTAGTTGCCAT---CGGGTA--AA---------GCCGGGCACTCTAG
AGGGACTGC----CGTCGACA-AGACGGAGGAAGGTGGGGATGACGTCAAATCA-TCATGC-CCTATATGCCCTG-GGCC
ACACACGTGCTACAATGGCCGGTACAGAGGGAAG-CGAAGGAG-------------------------------------
----------------------------------------CGATCCGGAGCGAATCCC-AAAAAGCCGGTCTCAGTTCGG
ATTGCAGGCTGCAACTCGCCTGCATGAAG--GCGGAATCGCTAGTAATCGCGGATCAGCA-TGCCGCGGTGAATGCGTTC
CCGGGCCTTGTACACACCGCCCGTCACACCACGAGAGTCTGCAACACCCGAAGCCGGTGGCCCAACCCGAAG--------
-------------A--------GGGAGGGAGCCGTCGAAGGTGGGGCAGATGATTGGGGTGAAGTCGTAACAAGGTAGC
CGTACGGGAACGTGCGGCTGGATCACCTCCTT------------------------------------------------
------------------------------------------------------
>lcl|CP000568.1_rrna_54 [locus_tag=Cthe_R0054] [product=16S ribosomal RNA] [location=3427593..3429113] [gbkey=rRNA] Hth
--------------------------------------------------------------------------------
--------------------------------------------------------------------------------
--------------------------------------------------------------------------------
--------------------------------------------------------------------------------
--------------------------------------------------------------------------------
--------------------------------------------------------------------------------
--------------------------------------------------------------------------------

--------------------------------------------------------------------------------
-------------------------------------------------------AGAGTTTGATCCTGGCTCAG-GACG
AACGCTGGCGGCGTGCCTAACACATGCAAGTCGAGCGGGGATATACGGAAGGT-----TT------A-CCGGAAGTATAT
CCTAGCGGCGGACGGGTGAGTAACG-CGTGGGTAACC-------------------------------------------
-----TACCTCATAC--AGGGGGATAACACAGGGAAA-CCTGTGCTAATACCGCAT-AACATAACGGGGCG---------
-------G-------------------------------CAT------CGTCCTGTTATCAAAGGA---GAAATCCGGTA
TGAGATGGGCCCGCGTCCGATTAGCTAGTTGGTGAGGT-AACGGCTCACCAAGGCGACGATCGGTAGCCGAACTGAGAGG
TTGGTCGGCCACATTGGGACTGAGACACGGCCCAGACTCCTACGGG--AGGCAGCAGTGGGGAATATTGCGCAATGGGGG
AAACCCTGAC---GCAG---C-----------AACGCCGCGTGAAGGAAGAAGGCCTTCGG-GTTGTAAACTTCTTTGAT
TGGGGACGAAGG-A-----------------------AGTGACGGTACCCAAAGAACAAGCCACGGCTAACTACGTGCCA
GCAGCCGCGGTAATACGTAGGTGGCGAGCGTTGTCCGGAATTACTGGGTGTAAAGGGCGCGTAGGCGGGGATGCAAGTCA
GATGTGAAATTCCGGGGCTTAACCCCGG-CGCT----------------------------------GCATCTGAAACT-
-----------------------------------GTATCTCTTGAGTGCTGGAGAGGAAAGCGGAATTCCTAGTGTAGC
GGTGAAATGCGTAGATATTAGGAGGA------------------------------------------------------
--------------------------------------------------------------------------------
-----------------------------------ACACCAGTGGCGAAGGCGGCTTTCTGGACAGTAACTGACGCTGAG
GCGCGAAAGCGTGGGGAGCAAACAGGATTAGATACCCTGGTAGTCCACGCCGTAAAC-----------------------
--------------------------------------------------------------------------------
---------GATGGATACTAGGTGTAGGAGG---TAT-CGAC---CCCTTCTGTGCCGGAGTTAACACAATAAGTATCCC
ACCTGGGGAGTACGGCCGCAAGGTTGAAACTCAA-AGGAATTGACGGGGGCCCGCACA-AGCAGTGGAGTA-TGTGGTTT
AATTCGAAGCAACGCGAAGAACCTTACCAGGGCTTGACATCCCTCTGACAGCTCTAGAG--ATAGGGCT------------
---------------------T----CCC-------TTCGGGGCAGAGGAGACAGGTGGTGCATGGTTGTCGTCAGCTCG
TGTCG-TGAGATG--TTGGGTTAAGTCCCGCAAC----------------------------------------------
--------------------------------------------------------------------------------
----------G------AGCGCAACCCTTGTCGTTAGTTGCCAG---CACGTT--AA---------GGTGGGCACTCTAG
CGAGACTGC----CGGCGACA-AGTCGGAGGAAGGTGGGGACGACGTCAAATCA-TCATGC-CCCTTATGTCCTG-GGCT
ACACACGTACTACAATGGCTGCTACAAAGGGAAG-CGATACCG-------------------------------------
----------------------------------------CGAGGTGGAGCAAATC-C-CCAAAAGCAGTCCCAGTTCGG
ATTGCAGGCTGAAACTCGCCTGCATGAAG--TCGGAATTGCTAGTAATGGCAGGTCAGCA-TACTGCCGTGAATACGTTC
CCGGGCCTTGTACACACCGCCCGTCACACCATGAGAGTCTGCAACACCCGAAGTCAGTAGTCTAACCGCAAG--------
-------------------------GAGGGCGCTGCCGAAGGTGGGGCAGATGATTGGGGTGAAGTCGTAACAAGGTAGC
CGTATCGGAAGGTGCGGCTGGATCACCTCCTTT-----------------------------------------------
------------------------------------------------------
>lcl|CP002360.1_rrna_6 [locus_tag=Mahau_R0006] [product=16S ribosomal RNA] [location=333212..334738]
[gbkey=rRNA] Mau
--------------------------------------------------------------------------------
--------------------------------------------------------------------------------
--------------------------------------------------------------------------------
--------------------------------------------------------------------------------
--------------------------------------------------------------------------------
--------------------------------------------------------------------------------
--------------------------------------------------------------------------------
--------------------------------------------------------------------------------
-------------------------------------------------------AGAGTTTGATCCTGGCTCAG-GACG
AACGCTGGCGGCGTGCCTAAAACATGCAAGTCGAACGGATATACTAACATTAAGC---GA------T-TAATGGGAGTAT
ATTAGTGGCGGACGGGTGAGTAACA-CGTGAACAACC-------------------------------------------
-----TGCCCTGTAC--AGGGGGATAACAGACCGAAA-GGACTGCTAATACCCCAT-GAGACCACAGCATC---------
-------G-------------------------------CAT------GATGCGGGGGTCAAAGGA---GCAATCCGGTA
TAGGAGGGGTTCGCGGCCCATTAGCTAGTTGGTGAGGGTAGAAGCCCACCAAGGCGACGATGGGTAGCCGGCCTGAGAGG
GTGAACGGCCACACTGGGACTGAGACACGGCCCAGACTCCTACGGG--AGGCAGCAGTTGGGGATATTGCGCAATGGGCG
AAAGCCTGAC---GCAG---C-----------GACGCCGCGTGAGGGAAGAAGGTCTTCGG-ATTGTAAACCTCTGTCGA
GTGGGAAGAAAG-A-----------------------AATGACAGTACCACTGGAGGAAGCCCCGGCTAACTACGTGCCA
GCAGCCGCGGTAATACGTAGGGGGCTAGCGTTGTCCGGAATTACTGGGCGTAAAGGGCGTGTAGGCGGTAGAGCAAGTCA
GGTGTGAAACCCCCGGGCTCAACTCGGG-GCAT----------------------------------GCATCTGAAACT-
-----------------------------------GCGATACTTGAGTGCAGGAGAGGGAAGCGGAATTTCTGGTGTAGC
GGTGGAATGCGTAGATATCAGGAAGA------------------------------------------------------
--------------------------------------------------------------------------------
-----------------------------------ACACCGGTGGCGAAGGCGGCTTTCTGGCCTGCAACTGACGCTGAG
GCGCGAAAGCGTGGGGAGCGAACAGGATTAGATACCCTGGTAGTCCACGCTGTAAAC-----------------------
--------------------------------------------------------------------------------
---------GATGGATACTAGGTGTAGGTGG---TAT-CAAA---GCCATCTGTGCCGAAGTTAACACATTAAGTATCCC
GCCTGGGGAGTACGGCCGCAAGGTTGAAACTCAA-AGGAATTGACGGGGGCCCGCACA-AGCAGCGGAGCG-TGTGGTTT
AATTCGAAGCAACGCGAAGAACCTTACCAGGTTTTGACATGCATATGAATTATGCAGAA--ATGCATGA------------
---------------------G----GCTTATCGAAAGATAAGACATATGCACAGGTGGTGCATGGTTGTCGTCAGCTCG
TGTCG-TGAGATG--TTGGGTTAAGTCCCGCAAC----------------------------------------------
--------------------------------------------------------------------------------
----------G------AGCGCAACCCTTATTGTTAGTTGCCAG---CACGTA--GA---------GGTGGGCACTCTAA
CGAGACTGC----CGTGGATA-ACACGGAGGAAGGTGGGGATGACGTCAAATCA-TCATGC-CCCTTATGATCTG-GGCT
ACACACGCGCTACAATGGCCGGAACAAAGAGGAG-CGAAGCGG-------------------------------------
----------------------------------------TAACGCAGAGGGAATCTC-AAAAAACCGGTCTCAGTTCAG
ATTGTGGGCTGCAACCCGCCCACATGAAG--AAGGAGTTGCTAGTAATCGCGGATCAGCA-TGCCGCGGTGAATACGTTC
CCGGGCCTTGTACACACCGCCCGTCACACCATGAGAGTCTACAACACCTTAAGCCGGTGAGCTAACCGCAAG--------
-------------------------GAGGCAGCCGTCCAAGGTGGGGTAGATGATTGGGGTGAAGTCGTAACAAGGTAGC
CGTATCGGAAGGTGCGGCTGGATCACCT----------------------------------------------------
------------------------------------------------------
>lcl|NW_002477113.1_rrna_6133 [locus_tag=GL50803_r0019] [db_xref=GeneID:5696754] [product=18S ribosomal RNA]
[location=complement(4375..5823)] [gbkey=rRNA] Gla
-CATCCGGTCGATCCTGCCGGAGCGCGACGCTCTCCCCAAGG-ACGAAGCCATGCATGCCCGCTCACC--------CGGG
ACGCGGCGGACGGCTCAGGACAACGGTTGCACCCCCCGCGGCGGTCCCTGCTAGCCGGACACCGCTGGCAACCCGGCGCC
AAGACGTGCGCG--CAAGG------GCGGGCG-----CCCGCGGGCGAGCAGCGTGACGCAGCGACGGCCCGCCC-----
-GGGCTTCCGGGGCATCACCCGGTCGGCGCGGTCGCGGCGCGCCGAGGGCCCGACGCCTGGCGGAGAATCAGGGTTCGAC
TCCGGAGAGCGGGCCTGCGAGACGGCCCGCACATCCAAGGACGGCAGCAGGCGCGGAACTTGCCCAATGCGCGGCGCGCG
AGGCAG-CGACGGGGA----------------------------------------------------GCGCGCGAGCGA
GGCGGGCCCACAGCCCCCGCCGCGGAGCCGA-------------------------------------------------
--------------------------------------------------------------------------------
--------------------------------------------------------------------------------
--------------------------------------------------------------------------------

--------------------------------------------------------------------------------
--------------------------------------------------------------------------------
--------------------------------------------------------------------------------
--------------------------------------------------------------------------------
--------------------------------------------------------------------------------
--------------------------------------------------------------------------------
--------------------------------------------------------------GGGCAAGGTCTGGTGCCA
GCAGCCGCGGTAATTCCAGCTCGGCGAGCGTCGCGCGGCGCTGCTGCAGTTGAAACGCCCGTAGTTGGCCCCCCGCCGCC
ACGAGGAAACGGGAGCGCTCCAGGCAGGCCCGTTGGACCCGCCGCGTGGGACCGCGCAGCGGGCGCGGCGCGCCGCGGC-
-----------------------------------AGCCCCGAGGAGAGCGGGCGGGGGCACCGGTACCGGCCGGGGACG
GGTGAAACAGGATGATCCCGCCGAGACCGCCGGCC---------------------------------------------
---------------------------------------------------------------------GCGCAGGCGCC
TGCCAAGACCGCCTCT-------------------------------------------------------GTCAATCAA
GGGCGAAGGCCGGGGGCTAGAAGGCGATCAGACACCACCGTATTCCCGGCCGTAAAC-----------------------
--------------------------------------------------------------------------------
---------GGTGCCGCCCCGCGGCCGGC-----GCGCGCGTCCCGCCGGCCGCCCAGGGAAACCGGGAGGCTCCGGGCT
CTGGGGGGAGTATGGCCGCAAGGCTGAAACTTGAAGGCATTGACGGAGGGGTACCACC-AGACGTGGAGTC-TGCGGCTC
AATCTGACTCAACGCGCGCACCTCACCAGGCC-CGGACGCGCGGAGGACCGACAG-------------------------
----------------------CCGGGCGCGCTTTCGCGATC----GCGCGGGCGGTGGTGCATGGCCGCTCCCAGCCCG
TGGCG-CGAGCCG--TCTGCTCCATTGCGACAAC----------------------------------------------
--------------------------------------------------------------------------------
----------G------AGCGAGACCCCGGCCGCGGGCGCC---------------------------------------
GCGGGACGG----CCCGCGCG-AGCGGGAGGACGGCG-GGGCGATAGCAGGTCTGTGATGC-CCTCAGACGCCCTGGGCC
GCACGCGCGCTACACTGGCGGGGCCAGCCGGCGC-CC-------------------------------------------
-----------------------------------------------GCGAGGACGCGCGGAGCCCCCGCCGTGGCCGGG
ACCGCGGGCTGGAACGCCCCCGC--GCAC--CAGGAATGTCTTGTAGGCGCCCGCCCCCACCG-CGCGCCGGACGCGTCC
CTGCCCCTTGTACACACCGCCCGTCGCTCCTACCGACTGGGCGCGGCGGCGAGCGCCCCGGA------------------
----------------------CGCGCGAAGGGCCGCGAGCCCCCGCGCCTGGAGGAAGGAGAAGTCGTAACAAGGTATC
CGTAGGTGAACCTGCGGATGGATCC-------------------------------------------------------
------------------------------------------------------
>lcl|QMZX01000069.1_rrna_17 [locus_tag=DRP18_03030] [product=16S ribosomal RNA] [location=676..2505]
[gbkey=rRNA] Aen
--------------------------------------------------------------------------------
--------------------------------------------------------------------------------
--------------------------------------------------------------------------------
--------------------------------------------------------------------------------
--------------------------------------------------------------------------------
--------------------------------------------------------------------------------
--------------------------------------------------------------------------------
--------------------------------------------------------------------------------
-----------------------------------------------------ATTCCGGTTGATCCCGCCGGAGGTCAC
TGCTATCTGAATCTGACTAAGACATGCTAGTGCTGGGTAAAACCAGTG--------------------------------
--------CGTACAGCTCAGTATCA-GATTGACGTTCTGGAGGAACGT---CTTTCTGCTGGA-AACAC------GCAGT
TAACCTGCCCTTAGG--CCCGCCATAATCCCGGGAAA-CTGGGGCTAAAGGCGGAC-AGGTA--AGGGA-TGCTGGAAAC
CCTTTTTCTATATTAAACTTATTA----------AAAGCAGTTACGGAAAAGGGCTGGAATGCGCC---CTTATCGAACT
AAGGATGGGACTGCGGCCGATTAGCTTGTTGCTAGGGT-AAAATCCTGGCAAGGCGATAATCGGTACGGGCCTTGAAAGA
GGAAGCCCGGAGATGGATTCTGAGACAAGAATCCAGGTCCTACGGG--ACGCAGCAGTCGCGAAACTTTCACAATGCGCG
CAAGCGTGAT---GGAG---G-----------GATTCAGAGTGTCTTCTCTTTTTTTGA------GGAGACTTTT-----
-------------------------------GTGGAGTCTAAACCACTCCACGAATAAGTGGTGGGTAAGACGGGTGCCA
GCCGCCGCGGTAATACCCGCGCCACGAGTGGTGACCACGTTTATTGGGCTTAAAGCGTTCGTAGCCGGCCTGGTAAGTTT
CCAATGAAATTCCACAGATGACTGT-GGAGCTT----------------------------------GTTGGAAATACT-
-----------------------------------GCCGGGCTTGGGACCGGGAGAGGTCGAGGGTATTGTAGGGGTAGG
GGTAAAAACAGATGATACCCGAAGGGTATCTGTCCAGATACATTCCTATAATCCCTACAGGACCGCCTGTGGCGAAGCCT
TACTTTACGGAAGCTATTACTTTCTCAAAGAAAGCCTTGGGAAAGAAACACGCAAAAAAGAGGGCTCCGCATAAGGCGCT
CGACTAGAACGGATCC-------------------------------------------------------GACGGTGAG
GGACGAAAGCTAGGGGAGCGAACGGGATTAGATACCCCGGTAGTCCTAGCCGTAACAGATTGGACGGTT-----------
-------------------------------------------------------ACGTATTTAATTCACACTGTCGGTC
TGCGGATACGATACCCACTAGAC-GTTGGAGTTCTTACGGGGGGCTCCAGA---GTCGTAGGGAAACCGTTAAGTGGGTC
GCCTGGGGAGTACGGTCGCAAGGCTGAAACTTAAGGGAATTGGCGGGGGAGCACACA--AGGGGTGGAGGC-TGCGGTTC
AATTCGACTCAACGCGAGGAACCTTACCAGCG-GCGACAGCAGAAAATGTGCATGACGACTTGAACTGGTTGTTCAAATC
GTCCGTCTGCAGTGAAGGTCAGGCTGAAGGTCTTACCAGACA----AGCTGAGAGGTGCTGCATGGCCGTCGTCAGTTCG
TGCCG-TGTAAGC--CTTTCTTTACAGAAGCTGTCGCT------TTCTCA-----------------------------A
AGAAAGCCTTGGGAAA--------GAAATA-------TGCA-----AAAAAGAAAGGCTTCGGTAAAGGTGTCCTGTTAA
GTCAGGAAACG------AACGAGACCCGCACCCTGTGTTACTAC---TGGGTCCTTTGG------GCTCAGGTACTCACA
GGGGACCGC----TGGCGACA-AGCCAGAGGAAGGGGCGGGCGACGGCAGGTCA-GTATGG-TCTGAATCCGCTG-GGCA
ACACGCGGCCTACAATGGCCGGAACAATGAGATG-CGACCCTGAAAAACCGTCTGCTAACACTGAACAAAGATTGCTTGG
ATATTCAAGCAGACTTTTCGCAATGTTAGCAATCGGTA--CAGAAGGGGGAGCTAATCTCTAAATCCGGTCTTGGTTCGG
ATCGAGGACTGCAACTTGTCCTCGTGAAG--GCGAAATCCCTAGTAGTCGCTGGTCATCATCC-AGCGGCGAATAAGTTC
CTGCTCCTTGCACACACTGCCCGTCAAGCCAACCGAGTGGGTTTTTGGTGAGGCTCTGGTTTTTGCC-------------
--------------------------------------------------------------------------------
--------------------------------------------------------------------------------
------------------------------------------------------
>lcl|MEHH01000036.1_rrna_16 [locus_tag=HeimAB125_07880] [product=16S ribosomal RNA] [location=3025..4474]
[gbkey=rRNA] Hei
--------------------------------------------------------------------------------
--------------------------------------------------------------------------------
--------------------------------------------------------------------------------
--------------------------------------------------------------------------------
--------------------------------------------------------------------------------
--------------------------------------------------------------------------------
--------------------------------------------------------------------------------
--------------------------------------------------------------------------------
--------------------------------------------------------CCGGTTGATCATGCCGGACCTAGG
TGCTATCAGATTCGCAATAGACATGC-AAGTTT--------------AATGT--------------------CTCTTCGG
AGGCATGGCGAAAGGCTCAGTAACA-CGTCGCCAATCTGCCCTATGGA--------------------------------
-----------------CG-GGGATAACCTCGGGAAA-CTGAGATTAATACCCGAT-AGGTA--TGGAA-TTCTGGAAGG
AGCCAATACCCAA------------------AGGGGCTTCGGTTCGCCATAGGATGAGGCCGCGGC---C----------

```
------------------GATTAGCTAGTAAGTGATGT-AAAGGATCACTTAGGCGATGATCGGTAGGGGTAATGAGAGT
TAGCGCCCCGAGAAGGACACTTAGACACTGGTCCTAGCACTACGGT--GTGCAGCAGTTAGGAATTGTGCCCAATGCGCG
AAAGCGTGAG---ACCG---C-----------GAATCTGAGTGCCAAGGAGAACC-----------TTGGCTGTG-----
-------------------------------ATGGAGTGTTCGTAGCTCCAACAGCAAGCAGAGGGCAAGGCTGGTGCCA
GCCGCCGCGGTAAAACCAGCTCTGCGAGTGCCTAGGACGATTATTGGGCTTAAAGCATCCGTAGCCGGGTAAGTAGGTCT
CTGATTAAATCTGCAAGCTTAACTT-GTAGGCT----------------------------------GTCAGAGATACC-
-----------------------------------GCTAACCTAGGGAATAGGAGAGGTGAACGGTACTGCGAGAGAAGC
GGTGAAATGCGTTGATTCTCGCAGGACCCACAG-----------------------------------------------
-------------------------------------------------------------------TGGCGAAGGCGGT
TCACTGGAATATCTCC-------------------------------------------------------GACGGTGAT
GGATGAAAGCCAGGGGAGCGAAAGGGATTAGAGACCCCTGTAGTCCTGGCCGTAAA------------------------
--------------------------------------------------------------------------------
--------CGATGAGGATTAGGTGTTGGTCATGGCTAAGGGCCTGGTCAGT---GCCAAAGGGAAACTATTAAATCCTCC
GCCTGGGGAGTACGGTCGCAAGGCTGAAACTTAAATGAATTGACGGGAAAGCGCCACC-AGGCACGGGATG-TGTGGTTT
AATTCGACTCAACGCGAGGAAACTCACCTGGG-GCGACTGTTACATGTGTGTCA--------------------------
---------------------GGCTGAAGACCTTACACGAAT--AAAACAGAGAGGTAGTGCATGGCCGTCTCAAGCTCG
TGCCG-TGAGGTG--TGCTCTTAAGTGAGTAAAC----------------------------------------------
--------------------------------------------------------------------------------
----------G------AGCGAGACCCGCGCCCCTATTTGCTAA---GAGCAAGCTTCGGCT-TGGCTGAGGACAATAGG
GGGATCGCT----ATCGATGA-AGATAGATGAAAGAGCGGGCCACGGCAGGTCA-GTATGC-TCCAAATCCCCAG-GGCT
ACACACGCATCACAATGGACAGGACAATGAGAAG-CGACTCCGAAAGGAGAAG---------------------------
---------------------------------------------------CGGACCTC-CAAACCTGTTCGCAGTAGGG
ATCGAGGTCTGGAACCG-ACCTCGTGAAC--ATGGAGCGCCTAGTATCCGTGTGTCATCATCG-CACGGAGAATACGTCC
CCGCTTTTTGTACACACCGCCCGTCGTTGCAACGAAATGAGGTTCGGTTGAGGCTGAGGCGAAT----------------
---------------------------CGGTTCAGTCAAAACTGGGCCTCGTGACGATGCAAAAGTCGTAACAAGGTATC
CGTA----------------------------------------------------------------------------
------------------------------------------------------
>lcl|CP019964.1_rrna_35 [locus_tag=Mia14_0796] [db_xref=RFAM:RF01959] [product=16S ribosomal RNA]
[location=complement(join(751297..751732,752255..753304))] [gbkey=rRNA] Man
--------------------------------------------------------------------------------
--------------------------------------------------------------------------------
--------------------------------------------------------------------------------
--------------------------------------------------------------------------------
--------------------------------------------------------------------------------
--------------------------------------------------------------------------------
--------------------------------------------------------------------------------
--------------------------------------------------------------------------------
-----------------------------------------------------ATTCCAGTCGATGCTGCTGGAGGACAC
TGCTATCAGATTTCGACAGCCATGCAAG--TTAG-------------CTCAA--------------------AGGCATCT
TTGGGCAGCGTACGGCTCAGTAACA-CGTAGTTAATCTAACGTAAAGA--------------------------------
-----------------AA-GGGATAACCTTGGGAAA-CTGAGGCTAATACCTTAT-AGATT--AAGAC-ATCTGGAAGG
AGTTTTAATCGAAAGGA-----GC-AT--GAATTGGAGTGTTTCCGCTTTACGATGAGACTGCGGC---G----------
------------------GATCAGGCAGATGGCGGGGT-AATGGCCCACCATACCTATAACCCGTAGGGGATGTGGGAGC
ATAAGCCCCGAGAAGGGCACTGAGGCAAGGGCCCTAGCGCTACGGC--GTGCAGCAGGCGCGCAAACTCCACAATGTGCG
AAAGCATGAT---GGGG---G-----------GAATCTGAGTGGTTTACGAAAGT-----------AAATCTTTT-----
-------------------------------GCCAAGTATAGCAAGCTTGGCGAATAAGTGCTGGGCAAGACGAGTGGCA
GCCGCCACGGTAATACTCGCAGCACAAGTGGTGTCCACGATTATTGGGCTTAAAGCGCTCGTAGCCGGTCAATGCCGTCT
TATGTGAAATATTGGTGCTCAACAT-CAATGCGT---------------------------------GCATGAGATACC-
-----------------------------------CATTGACTAGGGAGCGGGTGAGGTCAAGAGTACTTATAGGGGAGG
GGTAAAATCCTGTAATCCTATAAGGACTAACGG-----------------------------------------------
-------------------------------------------------------------------TGGCGAAGGCGCT
TGACCAGAACGCATCC-------------------------------------------------------GACGGTGAG
GAGCGAAGGCTAGGAGAACGAATCGGATCAGATACCCGAGTAGCCCTAGCAGTAAA------------------------
--------------------------------------------------------------------------------
--------TTATGCAGACTCTGGTGTTGCAAGTATCTTGAATGCTTGCAGT---ACCGTAGCAAAAGCGTTAAGTCTGCC
GCCTGGGGAGTACGGTCGCAAGATTAAAACTTAAAGCAATTGGCGGGGGAGCACA-CA-AGGGGTGGATGC-TACGGTTT
AATTGAATCCAACGTCGGAAATATTACCGGGA-GCGACGGCAGTGTGAAGGCCA--------------------------
---------------------GACTAAAGATCTTACCTGACA----AGCCGAGAGGTAGTGCATGGCGACCGTCAGCTCG
TGGTG-TGAACTG--TCCGGTTAAGTCCGGTAAC----------------------------------------------
--------------------------------------------------------------------------------
----------G------AGCGAGACTTTTGTCAACAGTTGCCAC---CAGCTTAGAGATAGG----TTGAGCACTCTGTT
GAGACCGCC----TCTGTTTA-AAGAGGAGGAAGGAGAGAGCGACGATACGTCA-GTATGC-TCCGAATCTCCCG-GGCT
ACACGCGGCATACAAAGATCAGCACAATGGGACG-CAATGCCGAAAGGCAAAG---------------------------
---------------------------------------------------CAAATCCCCTAAAACTGACCTAAGTTCGG
ATTGAGGGCTGCAACTCGCCCTCATGAAG--ATGGAATCCCTAGTAATCGCTTGTCACTATCG-AGCGGTGAATTTGTCC
CTGCTCCTTGCACACACTGCCTACCAAGTCACCCAAGTGAGGTTTAACGAGGCGATGCCATT------------------
---------------------------AGGCATTTTCGAGTTAGGGTTTCGCGAGGAGGGCTAAGGTATAACAAGGTATC
CGTAGGGGAACCTGCGGATAGATCACCTCAA-------------------------------------------------
------------------------------------------------------
>lcl|NZBU01000007.1_rrna_15 [locus_tag=CL943_02160] [product=16S ribosomal RNA] [location=14512..>15997]
[gbkey=rRNA] Dia
--------------------------------------------------------------------------------
--------------------------------------------------------------------------------
--------------------------------------------------------------------------------
--------------------------------------------------------------------------------
--------------------------------------------------------------------------------
--------------------------------------------------------------------------------
--------------------------------------------------------------------------------
--------------------------------------------------------------------------------
-----------------------------------------------TGCGTAATTCCAGCTGATCCTGCTGGCGGATAC
TGCTATCGAAATCCGATTTAGACATGCAAGTCTGAAAGACGT-------------------------------ATACGGC
GTCTTTGGCGAACGGCTCAGTAACA-CGTGGGTAATCTACCTTTGGGA--------------------------------
-----------------CA-AGTTAATC-CCGGGAAA-CTGGGGATAATGCTTGAT-AGGCA--ATTGC-AACTGGAAGG
TGCTTTTGCTAAAATGC-----TA-CC--ACACCATGTGGATCGCGCCCAAGGATGAGCCTGCGGC---G----------
------------------GATTAGGTTGTTGGTGGGGT-AAATGCCCACCAAGCCGTTGATCCGTACGGGCCTTGAGAGA
GGTAGCCCGGAGAGGGACACTGAGATAAGGGTCCCAGCCCTACGGG--GTGCAGCAGTCGCGGAACCTTTACAATGCACG
AAAGTGTGAT---AAGG---G-----------GATTTCGAGTGCAGTCTTTGA--------------CTGCTTTT-----
```

-------------------------------GCCAAACCTAAATAGTTTGGCGAATAAGTGGGGGGCAAGATACGTGCCA
GCCGCCGCGGTAACACGTACTCCACAAGTGGTATCCACTAATATTGGGCCTAAAACGTCTGTAGCCGGTTCGATAAGTCC
TTGGTGAAATCATTGAGCTTAACTT-AATGAATG---------------------------------GCCAAGGATACT-
-----------------------------------GTTGGACTCGAGGCCGGGAGACGCCAAGGGAATTTCTGGGGTAGC
GGTGAAATGCTATGATCCCGGAAAGACCACCAG-----------------------------------------------
-------------------------------------------------------------------TAGCGAAGGCGCT
TGGCGAGAACGGGCCT-------------------------------------------------------GACGGTGAG
AGACGAAAGCTGGGGGAGCGACCCGGATTAGATACCCGGTTAGTCCCAGCCGTAAA------------------------
--------------------------------------------------------------------------------
--------CGATGCGGACTAGGT-GTGCTGTCACCCACGTGGTGCTAGCGT---GCCGAAGGGAAGCCATTAAGTCCGCT
GCCAGGGAAGTACGCTCGCAAGGGTGAAACTTAATGGAATTGGCGGGGGCGCACTACA-AGGGGTGGATGC-TGCGGTTT
AATCGGATAAAACCCCTAAGAATCTTACCAGGGCCGACAGCAGAATGATGGACA--------------------------
---------------------GTCTGAAGGGCTTTCCTGACG----CGCTGAGAGGTGTTGTATGGTCATCGTCAGCTCG
TACTA-TGAAGCG--TCCTGTTAACTCAGGCAAC----------------------------------------------
--------------------------------------------------------------------------------
----------G------GGCGAGACCCATGTCCCTAGTTGCTAC---TTTCACTTTTGTGGG-GGGC-----ACTCT-A-
GGGAGACTG----CAGTAGCA-ATACTGAGGAAGGAGTGGGCAACGGTAGATCT-GTATGG-CCCGAATGCCCTG-GGCA
ACACGCGGCATACAATGGCTGAGCCAATGGGTTC-CGACTCCGAGAGGAGAAG---------------------------
---------------------------------------------------GTAATCTCCTAAACTTAGCCTCAGTTCGG
ATCGAGGGTTGTAACCTACCCTCGTGAATGCCTGGAATCCCCAGTAATCGCGTGTCACTATCG-CGCGATGAATAAGTCC
CTGCGCCTTGCACACACCGCCCATCAAGCCAGTCGAGTAGAGCTTGGATGAGCTCTTTCCTTC-----------------
---------------------------GGGAAATATCGATTCTGAGTTCAGCGAGACGGGCTAAGTTGTAACAAGGTGTC
CGTAGGGGAACCTGCGGACGGATCAC------------------------------------------------------
------------------------------------------------------
>lcl|APJZ01000002.1_rrna_15 [locus_tag=Nst1_0248] [product=small subunit ribosomal RNA]
[location=complement(58109..59609)] [gbkey=rRNA] Nst
--------------------------------------------------------------------------------
--------------------------------------------------------------------------------
--------------------------------------------------------------------------------
--------------------------------------------------------------------------------
--------------------------------------------------------------------------------
--------------------------------------------------------------------------------
--------------------------------------------------------------------------------
--------------------------------------------------------------------------------
-------------------------------------------AATCCGCCGAATTCCCGTTGATCCTGCGGGGGGCCAT
CGCTATCTCCTTCCGACTAACCCATGGGAGGCTAGGGATACC------------------------------GGGTAAGG
TATCCCACCGCACGGCTGAGTAACA-CGTCGATAACTCTGTCTCGGGA--------------------------------
-----------------GG-AGAATATCCTCGGGAAA-CTGAGGGTAATTCTCCAT-AGGAA--ATGGG-GTGTGGAATT
CCCCCATTTCCGAAAAGGACTGGG-AT--AATCCCTCCCAGTTCCGCCCGAGAGGAGGTCGGCGGC---C----------
------------------TATCAGGTAGTTGGCGGGGT-AATGGCCCGCCAAGCCTAAGACGGGTAGGGGCCGTGAGAGC
GGGAGCCCCCAGATCGGCACTGAGACAAGGGCCGAGGTCCTACGGG--ATGCACCAGGGGCGAAACCTCCGCAATGGGTG
AAAGCCTGAC---GGGG---G----------GAGGGAGAGTGCCTGGGGTCTTCCCC----------AGGCTTTT-----
-------------------------------TGGGAGTGTAAAAAGCTCCCAGAATAAGCAGTGGGCCAGAGGGGTGGCA
GTCGCCACGGGAATACCCCCGCTGCGAGCGATGGCCTCGATTACTGGGCCTAAAGGGGCCGTAGCCGGGCTGGTGTGACC
CCTGTGAAATCCACGGCAATAATTCCGTGGGCGT---------------------------------GCAGGGGTTACT-
-----------------------------------ACTAGTCTAGGGGACGGAAGAAGCCAAGGGTATTCCCGGGGGAGT
AGCGAAATGCAGCAATCCCGGGAAGACCACCCG-----------------------------------------------
-------------------------------------------------------------------AGGCGAAGGCGCT
TGGCTATTCCGATCCC-------------------------------------------------------GACGGTGAT
GGCCGAAAGCTGGGGGATCAAAAGGGATTAGAGACCCCTGTAGTCCCAGCTGTCAA------------------------
--------------------------------------------------------------------------------
--------CTCTGTGGGCTACCT-GCTGGGTGGACTTAGAGTCCACTCAGT---GGGGTAGGGAAGCCGTTAAGCCCACC
GCCTGGGGAGTACGGCCGCAAGGCTGAAACTTAAAGGAATAGGCGGGGGGGCACA-CA-AGGGGTGGGGTG-TGCGGGTT
AATGGGACCCAACACCGGAAACCTTACCGGGA-GCGACAGCAGGATGATGGTCA--------------------------
---------------------GGCTGAAGACCTTACCGGACA----CGCTGAGAGGAGGTGCATGGCTGCCGTCAGCTCG
TGCCG-TGAGGTA--TCCTGTTAAGTCAGGAAAC----------------------------------------------
--------------------------------------------------------------------------------
----------G------AGCGCGACCCGCACCCGTAGTTACGAC---GGAGAGGATTTCCT----CTCCGGTACACTA--
CGGGGACTG----CCGGGGAA-ACCCGGAGGAAGGCGCGGGCGACGGCAAGTCC-GCATGC-CCCTAATCTCCCG-GGCT
ACACGCGCGCAACAATGGGCGGGACAATGGGAAC-CGACCCCGAAAGGGGAAG---------------------------
---------------------------------------------------GAAATCCCAGAAACCCGCTCTCAGTCCAG
ATTGAGGGCTGTAACTCGCCCTCATGACG--GTGGAATCCCTAGTAATCGGGGGTCACTATTCCCCCGGTGAATACGTCC
CTGCCCCTTGCACTCACCGCCCGTCAAGCCAACCGAGTCGTTTCCTGGTGAGGCCCGTTA--------------------
----------------------------AAGCGGGTCGAACCGGGAAACGGCGAGGGGGGCTAAGTCGTCACAAGGTAGC
CGTAGGGGAACCTGCGGCTGGATCTAGTT---------------------------------------------------
------------------------------------------------------
>lcl|RFJO01000060.1_rrna_29 [locus_tag=D6777_02840] [product=16S ribosomal RNA]
[location=complement(24819..26298)] [gbkey=rRNA] Woa
--------------------------------------------------------------------------------
--------------------------------------------------------------------------------
--------------------------------------------------------------------------------
--------------------------------------------------------------------------------
--------------------------------------------------------------------------------
--------------------------------------------------------------------------------
--------------------------------------------------------------------------------
--------------------------------------------------------------------------------
------------------------------------------------CTTTAATTCCCGTTGATCCTGCGGGAGGTCGC
TGCTATTTGGGTTCCACTTAGCCATGCAAGTCTAGGGGGCTC--------------------------------TTCGGA
GTCACCGGCGAAATGCTCAGTAACA-CGTCGATAATTTACCCTTAAGT--------------------------------
-----------------CT-GGGATAACCACGGGAAA-CTGTGGCTAATACCGGAT-AGAAA--ATTGA-GGCTGGAATG
CGCTTT-TTTCGAAAGC----------------------TCCGGCGCTTAAGGATAAGTCGGCGGC---G----------
------------------GATTAGGCTGTTGGTGAGGT-AACGGCTCACCAAACCGAAGATCCGTAGGGGGGATGAGAGT
CCTAGCCCCGAGAAGGGCACTGAGATAAGGGCCCTAGCCCTAAGGG--GTGCAGCAGGCGGGAAACCTTTACAATACACG
CAAGTGTGAT---AAGG---G----------AACCCAAAGTGCTTGCCTTCG------------GGTGAGCTTTT-----
-------------------------------GTCAAGCGTAAAAACCTTGACGAATAAGTGGTGGGTAAGACCGGTGCCA
GCCGCCGCGGTAACACCGGCGCCACGAGTGGTGACCACGTTTATTGGGCCTAAAGCGTCCGTAGCCGATCTAATAAGTTT
CTTGTGAAATCATTAAGCTTAACTT-AATGAATG---------------------------------GCAAGAAATACT-

-----------------------------------GTTAGGTTAGGGACCGGGAGGAGGCAGAAGTATTCCTTGGGAAGC
GGTAAAATGTTTTAATCCTTGGAAGACTACCTG-----------------------------------------------
-------------------------------------------------------------------TGGCGAAGGCGTC
TGCCTAGAACGGATCC-------------------------------------------------------GACGGTGAG
GGACGAAAGCTGGGGGAGCGAACCGGATTAGATACCCGAGTAGTCCTAGCTGTAAA------------------------
--------------------------------------------------------------------------------
--------CTCTGCGAACTATGT-GTTGCATATCTTCAGTGGATATGCAGT---GCCGAAGCGAAGGTGTTAAGTTCGCC
GCCTGGGAAGTATGGTCGCAAGACTGAAACTTAAAGGAATTGGCGGGGGCGCACTACA-AGGAGTGGAGCG-TGCGGTTT
AATCAGATTCTACACCGTGAACCTCACCAGGA-ACGACGGCAGGATGAAGGTCA--------------------------
---------------------GATTGAAGGTCTTACCTAACA----CGCCGAGAGGAGGTGCATGGCCGCCGTCAGTTCG
TGCCG-TAAGGTG--TCCAGTTAAGTCTGGCAAC----------------------------------------------
--------------------------------------------------------------------------------
----------G------AACAAGACCTATATCTACAGTTGCTAA---CGAGTCCTCCGGGAT-G-ATTGAGCACACTGT-
AGAGACTGC----C--TTGGA-AACAAGGAGGAAGGCGTAGGCAACGGTAGGTG-GTATGC-CCCGAATTTCCTG-GGCT
ACACGCGCGCAACAATGGTAGGGACAATGGGCTA-CAACTCCGAAAGGAGAAG---------------------------
---------------------------------------------------TTAATCCTCGAAACCCTACCTTAGTTCGG
ATTGAGGGCTGCAATTCGCCCTCATGAAG--CTGGAATTCCTAGTAATCGCTTGTCATCAGCG-AGCGGTGAATACGTCC
CGGTGCCTTGCACACACCGCCCGTCAAACCATTCGAGCAGGGTCTAGGTGAGGCTTAGGTTTTT----------------
----------------------------GCCTAAGACGAATCTAGGTTCAGTAAGAAGGGTTAAGTCGTCACAAGGTAGC
CGTAGGGGAACCTGCGGCTGGATCACCTCCTAT-----------------------------------------------
------------------------------------------------------
>lcl|RXIF01000002.1_rrna_27 [locus_tag=EF806_00470] [product=16S ribosomal RNA] [location=complement(41950..43436)] [gbkey=rRNA] Mlt
--------------------------------------------------------------------------------
--------------------------------------------------------------------------------
--------------------------------------------------------------------------------
--------------------------------------------------------------------------------
--------------------------------------------------------------------------------
--------------------------------------------------------------------------------
--------------------------------------------------------------------------------
--------------------------------------------------------------------------------
--------------------------------------------------GCAATTCCGGTTGATCCTGCCGGAGGCCAC
TGCTATGGGAATTCGACTAAGCCATGCAAGTTGAGGGCACCTT-------------------------------TTTAGG
AGACCCAGCGAACTGCTCAGTAACA-CGTGGATAATCTGCCCACAGGA--------------------------------
-----------------TG-GGGATAATCCTGGGAAA-CTGGGAATAATACCCAAT-AGATC--ATTGG-CACTGGAATG
TCCTTT-GATCCA------------------AATGCTTTTCTTACGCCTGTGGATGAGTCTGCGTC---C----------
------------------GATTAGGCTGTTGTTGGGGT-AACGGCCCAACAAACCTATGATCGGTACGGGTCGTGAGAGC
GATCGCCCGGAGATTGGATCTGAGATATGATCCTAGGCCCTACGGG--GTGCAGCAGGCGCGAAAACTTTACAATGTGGG
AAACCATGAT---AAGG---G-----------AATTCCCAGTGTTCCTACATAGTAGG---------AACTGTTT-----
-------------------------------AGGTGCGTAAAAAACACCTAATAGAAAGGGCCGGGTAAGACCGGTGCCA
GCCGCCGCGGTAATACCGGTGGCTCGAGTGGTGGCCGCTATTATTGGGTCTAAAGGGTCCGTAGCCGGCTTGATAAGTTA
CCTGGAAAATCCGGTGGCCTAACCA-TTGGGCGG---------------------------------CCAGGTGATACT-
-----------------------------------ATCAGGCTTGGGACTAGGAGAGACCAGAGGTACTCATGAGGTAGG
GGTGAAATCCTGTAATCTTATGAGGACCACCAG-----------------------------------------------
-------------------------------------------------------------------TGGCGAAGGCGTC
TGGTTAGAATAGGTCC-------------------------------------------------------GACGGTGAT
GGACGAAGGCTGGGGGCGCAAACCGGATTAGATACCCGGGTAGTCCCAGCAGTAAA------------------------
--------------------------------------------------------------------------------
--------CGATGCCAGCTATGT-GTTGCTGAAACCATGGGTTTCAGCAGT---GCCGAAGGGAAGCCGTGAAGCTGGCC
ACCTGGGAAGTACGGCCGCAAGGCTGAAACTTAAAGGAATTGGCGGGGGAGTACCACA-ACCGGTGGAGCC-TGCGGTTT
AATTGGATACAACGCCGGGAATCTTACCGGAG-GGGACAGCAGTATGAAGGTCA--------------------------
---------------------GGCTAAAGACCTTACCAGATC----AGCTGAGAGGAGGTGCATGGCCGTCGCCAGTTCG
TACCG-TGAGGCA--TCCTGTTTAGTCAGGCAAC----------------------------------------------
--------------------------------------------------------------------------------
----------G------ATCGAGACCCGTAGTCTTATTTGCCAG---CATGCCCTCTGG-GG-TGTATGGGGACAATAA-
GACGACCGT----CAGCGCTA-AGCTGAAGGAAGGAGCGGGCTACGGTAGGTCA-GCATGC-CCTGAATCCTCCG-GGAT
ACACGCGGGCTACAATGGTCAGGACAAAAGGTAT-CGACCTCGAAAGAGTGAG---------------------------
---------------------------------------------------ATAATCCCATAAACCTGATCTCAGTCCGG
ATCGAAGGCTGCAACTCGCCTTCGTGAAG--ATGGAATCGGTAGTAATCGTGACTCAAAATGT-CACGGTGAATATGTCC
CTGCTCCTTGCACACACCGCCCGTCAAATCACCCGAGTGGGATCTGGAGGAGGTCTTTCTCGCA----------------
---------------------------GAGGAAGATCGAATCTGGATTCTGCAAGGGGGGTTAAGTCGTAACAAGGTAGC
CGTAGGGGAACCTGCGGCTGGATCACCTCCTA------------------------------------------------
------------------------------------------------------
>lcl|AE004437.1_rrna_42 [gene=rrs] [locus_tag=VNG_r02] [product=16S ribosomal RNA] [location=1875505..1876977] [gbkey=rRNA] Hal
--------------------------------------------------------------------------------
--------------------------------------------------------------------------------
--------------------------------------------------------------------------------
--------------------------------------------------------------------------------
--------------------------------------------------------------------------------
--------------------------------------------------------------------------------
--------------------------------------------------------------------------------
--------------------------------------------------------------------------------
-----------------------------------------------------ATTCCGGTTGATCCTGCCGGAGGCCAT
TGCTATCGGAGTCCGATTTAGCCATGCTAGTTGTGCGGG------------------------------------TTTAG
ACCCGCAGCGGAAAGCTCAGTAACA-CGTGGCCAAGCTACCCTGTGGA--------------------------------
-----------------CG-GGAATACTCTCGGGAAA-CTGAGGCTAATCCCCGAT-AACGC--TTTGC-TCCTGGAAGG
GGCAAA-GCCGGA---------------------AACGCTCCGGCGCCACAGGATGCGGCTGCGGT---C----------
------------------GATTAGGTAGACGGTGGGGT-AACGGCCCACCGTGCCCATAATCGGTACGGGTTGTGAGAGC
AAGAGCCCGGAGACGGAATCTGAGACAAGATTCCGGGCCCTACGGG--GCGCAGCAGGCGCGAAACCTTTACACTGTACG
AAAGTGCGAT---AAGG---G-----------GACTCCGAGTGTGAAGGCATAGAGCCT--------TCACTTTT-----
-------------------------------GTACACCGTAAGGTGGTGCACGAATAAGGACTGGGCAAGACCGGTGCCA
GCCGCCGCGGTAATACCGGCAGTCCGAGTGATGGCCGATCTTATTGGGCCTAAAGCGTCCGTAGCTGGCTGAACAAGTCC
GTTGGGAAATCTGTCCGCTTAACGG-GCAGGCGT---------------------------------CCAGCGGAAACT-
-----------------------------------GTTCAGCTTGGGACCGGAAGACCTGAGGGGTACGTCTGGGGTAGG
AGTGAAATCCTGTAATCCTGGACGGACCGCCGG-----------------------------------------------
-------------------------------------------------------------------TGGCGAAAGCGCC

TCAGGAGAACGGATCC-------------------------------------------------------GACAGTGAG
GGACGAAAGCTAGGGTCTCGAACCGGATTAGATACCCGGGTAGTCCTAGCTGTAAA------------------------
--------------------------------------------------------------------------------
--------CGATGTCCGCTAGGT-GTGGCGCAGGCTACGAGCCTGCGCTGT---GCCGTAGGGAAGCCGAGAAGCGGACC
GCCTGGGAAGTACGTCTGCAAGGATGAAACTTAAAGGAATTGGCGGGGGAGCACTACA-ACCGGAGGAGCC-TGCGGTTT
AATTGGACTCAACGCCGGACATCTCACCAGCC-CCGACAGTAGTAATGACGGTCA-------------------------
---------------------GGTTGATGACCTTAC---CCGAGGCTACTGAGAGGAGGTGCATGGCCGCCGTCAGCTCG
TACCG-TGAGGCG--TCCTGTTAAGTCAGGCAAC----------------------------------------------
--------------------------------------------------------------------------------
----------G------AGCGAGACCCGCACTCCTAATTGCCAG---CAGTACCCTTTG-GG-TAGCTGGGTACATTAG-
GTGGACTGC----CGCTGCCA-AAGCGGAGGAAGGAACGGGCAACGGTAGGTCA-GTATGC-CCCGAATGGGCTG-GGCA
ACACGCGGGCTACAATGGTCGAGACAATGGGAAG-CCACTCCGAGAGGAGGCG---------------------------
---------------------------------------------------CTAATCTCCTAAACTCGATCGTAGTTCGG
ATTGAGGGCTGAAACTCGCCCTCATGAAG--CTGGATTCGGTAGTAATCGCGTGTCAGCAGCG-CGCGGTGAATACGTCC
CTGCTCCTTGCACACACCGCCCGTCAAATCACCCGAGTGGGGTTCGGATGAGGCCGGCAT--------------------
----------------------------GCGCTGGTCAAATCTGGGCTCCGCAAGGGGGATTAAGTCGTAACAAGGTAGC
CGTAGGGGAATCTGCGGCTGGATCACCTCCT-------------------------------------------------
------------------------------------------------------
>lcl|AE010299.1_rrna_1 [locus_tag=MA_0896] [product=16S ribosomal RNA] [location=1073041..1074470] [gbkey=rRNA]
Mac
--------------------------------------------------------------------------------
--------------------------------------------------------------------------------
--------------------------------------------------------------------------------
--------------------------------------------------------------------------------
--------------------------------------------------------------------------------
--------------------------------------------------------------------------------
--------------------------------------------------------------------------------
--------------------------------------------------------------------------------
-----------------------------------------------------ATTCTGGTTGATCCTGCCAGAGGTTAC
TGCTATCGGTGTTCGCCTAAGCCATGCGAGTCATATGTT-----------------------------------CTTTGT
GAACATGGCGTACTGCTCAGTAACA-CGTGGATAACCTGCCCTTGGGT--------------------------------
-----------------CC-GGCATAACCCCGGGAAA-CTGGGGATAATACCGGAT-AACGC--ACATA-TGCTGGAATG
CTTTCT-GCGT-------------------AAAACGGATTCGTCTGCCCAAGGATGGGTCTGCGGC---C----------
------------------TATCAGGTAGTAGTGGGTGT-AACGTACCTACTAGCCGACAACGGGTACGGGTTGTGAGAGC
AAGAGCCCGGAGATGGATTCTGAGACATGAATCCAGGCCCTACGGG--GCGCAGCAGGCGCGAAAACTTTACAATGCGGG
AAACCGTGAT---AAGG---G-----------GACACCGAGTGCCAGCATCATATGCT---------GGCTGTCC-----
-------------------------------GGATGTGTAAAATACATCTGTTAGCAAGGGCCGGGCAAGACCGGTGCCA
GCCGCCGCGGTAACACCGGCGGCCCGAGTGGTGATCGTGATTATTGGGTCTAAAGGGTCCGTAGCCGGTTTGGTCAGTCC
TCCGGGAAATCTGACAGCTCAACTG-TTAGGCTT----------------------------------TCGGGGATACT-
-----------------------------------GCCAGACTTGGAACCGGGAGAGGTAAGAGGTACTACAGGGGTAGG
AGTGAAATCTTGTAATCCCTGTGGGACCACCTG-----------------------------------------------
-------------------------------------------------------------------TGGCGAAGGCGTC
TTACCAGAACGGGTTC-------------------------------------------------------GACGGTGAG
GGACGAAAGCTGGGGGCACGAACCGGATTAGATACCCGGGTAGTCCCAGCCGTAAA------------------------
--------------------------------------------------------------------------------
--------CGATGCTCGCTAGGT-GTCAGGCATGGCGCGACCGTGTCTGGT---GCCGCAGGGAAGCCGTGAAGCGAGCC
ACCTGGGAAGTACGGCCGCAAGGCTGAAACTTAAAGGAATTGGCGGGGGAGCACAACA-ACGGGTGGAGCC-TGCGGTTT
AATTGGACTCAACGCCGGACAACTCACCGGG-ACGACAGCAATATGTAGGCCA---------------------------
---------------------GGCCGAAGACCTTGCCTGAAT----CGCTGAGAGGAGGTGCATGGCCGTCGCCAGTTCG
TACTG-TGAAGCA--TCCTGTTAAGTCAGGCAAC----------------------------------------------
--------------------------------------------------------------------------------
----------G------AGCGAGACCCGTGCCCACTGTTACCAG---CATGTCCTCCG--GG-ACGATGGGTACTCTGT-
GGGGACCGC----CGGTGTTA-AATCGGAGGAAGGTGCGGGCCACGGTAGGTCA-GTATGC-CCCGAATTTCCCG-GGCT
ACACGCGGGCTACAATGAATGGGACAATGGGTCC-CTCCCCTGAAAAGGGCTG---------------------------
---------------------------------------------------GTAATCTCACAAACCCATCCGTAGTTCGG
ATCGAGGGCTGTAACTCGCCCTCGTGAAG--CTGGAATCCGTAGTAATCGCGTTTCAATATAG-CGCGGTGAATACGTCC
CTGCTCCTTGCACACACCGCCCGTCAAACCACCCGAGTGAGGTATGGGTGAGGGCACGGACTTCG---------------
---------------------------TGCCGTGTTCGAACCTGAATTTTGCAAGGGGGGTTA-----------------
--------------------------------------------------------------------------------
------------------------------------------------------
>lcl|AE009439.1_rrna_15 [locus_tag=MKr02] [product=16S ribosomal RNA] [location=complement(516778..518289)]
[gbkey=rRNA] Mka
--------------------------------------------------------------------------------
--------------------------------------------------------------------------------
--------------------------------------------------------------------------------
--------------------------------------------------------------------------------
--------------------------------------------------------------------------------
--------------------------------------------------------------------------------
--------------------------------------------------------------------------------
--------------------------------------------------------------------------------
-----------------------------------------------------ACTCCGGTTGATCCTGCCGGAGGCCAC
CGCTATCGGGGTCCGACTAAGCCATGCAAGTCGAGGGCCGCCCGGCAA--------------------------TGGGGG
CGGCCCGGCGGACGGCTCAGTAACA-CGTGGGTAACCTACCCTCGGGA--------------------------------
-----------------CG-GGGATAACCCCGGCGAAAGTGGGGCTAATCCCCGAT-AGGCG--GGGCG-GCCTGGAACG
GTCCTC-CGCCGAAAGGGCCCGGG-CC--CATGCCGCCCGGGTCCGCCCGAGGATGGGCCTGCGGC---C----------
------------------GATTAGGTAGTTGGCGGGGT-AACGGCCCGCCAAGCCGATAATCGGTACGGGCGGTGAGAGC
CGGAGCCCGGAGACGGGGACTGAGACAAGGCCCCGGGCCCTACGGG--GCGCAGCAGGCGCGAAACCTCCGCAATGCGGG
CAACCGCGAC---GGGG---G-----------GACCCCGAGTGCCGTCGGGGCAAAGCCCC---GGCGGCTGTAC-----
-------------------------------CGGGGTGTAAAAAGCCCCGGGTAGAAAGCGGCGGGCAAGACCGCTGCCA
GCCGCCGCGGTAATAGCGGCGCCGCAAGTGGTGGCCGCTTTTATTGGGCCTAAAGGGGCCGTAGCCGGTCCCGTGGGTCC
CCGCCGAAAGCCCGCGGCTTAACCG-CGGGAGTC----------------------------------GGCGGGAAACT-
-----------------------------------GCGGGACTTGGGACCGGGAGAGGCCGGAGGTACCCCCGGGGTAGG
GGTGAAATCCTGTCATCCCGGGGGGACCGCCAG-----------------------------------------------
-------------------------------------------------------------------TGGCGAAGGCGTC
CGGCTGGAACGGGTCC-------------------------------------------------------GACGGTGAG
GGCCGAAAGCCGGGGGAGCAAACCGGATTAGATACCCGGGTAGTCCCGGCTGTAAA------------------------
--------------------------------------------------------------------------------

```
--------CGATGCGGACTAGGT-GTTGGGGCGGCCACGAGCCGCCCCAGT---GCCGTAGGGAAGCCGTTAAGTCCGCC
GCCTGGGGAGTACGGCCGCAAGGCTGAAACTTAAAGGAATTGGCGGGGGAGCACCACA-ACCGGTGGAGCC-TGCGGTTT
AATTGGATTCAACGCCGGAAACCTTACCGGGG-GCGACAGCAGGATGAAGGCCA--------------------------
---------------------GGTTGACGACCTTGCCGGACG----AGCTGAGAGGAGGTGCATGGCCGCCGTCAGCTCG
TGCCG-TGAGGTG--TCCTGTTAAGTCAGGTAAC----------------------------------------------
--------------------------------------------------------------------------------
----------G------AGCGAGACCCCCGCCCGCAGTTGCCAG---CGGGCCCCGTAAGGG-GCGCCGGGCACTCTGC-
GGGGATCGC----CGCCGTTA-AGGCGGATGAAAGTGGGGGCGACGGCAGGTCC-GTATGC-CCCGAAACCCCCG-GGCT
ACACGCGGGCTACAATGGCGGGGACAATGGGATC-CGACCCCGAAAGGGGGAG---------------------------
---------------------------------------------------GAAATCCCCTAAACCCCGTCGTAGTTCGG
ATTGCGGGCTGCAACTCGCCCGCATGAAG--GTGGAATCGGTAGTAACCGTGCCTCAGAATGG-CACGGTGAATACGTCC
CTGCTCCTTGCACACACCGCCCGTCACGCCACCCGAGCCCCCCGGGGGCAAGCCCCCGGTCCGCA---------------
-------------------------AGGGCTGGGGGCGAGCCCCCGGGGGGTGAGGGGGGCGAAGTCGTAACAAGGTAGC
CGTAGGGGAACCTGCGGCTGGATCACCTC---------------------------------------------------
------------------------------------------------------
>lcl|CP010070.1_rrna_37 [gene=rrfA] [locus_tag=Mpt1_c10460] [product=16S ribosomal RNA]
[location=complement(1070956..1072425)] [gbkey=rRNA] Mte
--------------------------------------------------------------------------------
--------------------------------------------------------------------------------
--------------------------------------------------------------------------------
--------------------------------------------------------------------------------
--------------------------------------------------------------------------------
--------------------------------------------------------------------------------
--------------------------------------------------------------------------------
--------------------------------------------------------------------------------
-----------------------------------------------------ACTCCGGTTGATCCTGCCGGCGGCCAC
CGCTATAGGAATTCGATTAAGACATGCGAGTCGAGAGTC------------------------------------GTTAT
GGACTCGGCGGACTGCTCAGTAACA-CGTGGATAACGTGCCCTTTGGT--------------------------------
-----------------GG-AGGATAATCTCGGGAAA-TTGAGAATAATACTCCAT-AGATC--ATGAG-ATACTGAATG
ACCCAT-GGTCCAAAGT----------------------TCCGGCGCCAAAGGATCGGTCTGCGGC---C----------
------------------TATCAGGTAGTAGTGGGTGT-AACGTACCTACTAGCCTATGACGGGTATGGGCCTTGAGAGA
GGGAGCCCAGAGTTGGATTCTGAGACACGAATCCAGGCCCTACGGG--GCGCAGCAGTCGCGAAAACTTCACACTGGGGG
CAACCCCGAT---GAGG---G-----------AATTCCTAGTGCTAGCACATTAG-TGT--------TAGCTTTT-----
-------------------------------CTTCAGCATAGATAACTGGAGGAATAAGGGCTGGGTAAGACGGGTGCCA
GCCGCCGCGGTAATACCTGCAGCCCAAGTGGTGGTCGATATTATTGAGTCTAAAACGTTCGTAGCCGGTTTAATAAATCC
TTGGGTAAATCGGGGGGCTTAACCT-TCCGAAT----------------------------------TCCGAGGAGACT-
-----------------------------------GTTAAACTTGGGACCGGGAGAGGCAAGAGGTACTTCTGGGGTAGG
GGTAAAATCCTGTAATCCTAGAAGGACCACCGG-----------------------------------------------
-------------------------------------------------------------------TGGCGAAGGCGTC
TTGCTAGAACGGATCC-------------------------------------------------------GACGGTGAG
GGACGAAGCCCTGGGGCGCAAACGGGATTAGATACCCCGGTAGTCCAGGGTGTAAA------------------------
--------------------------------------------------------------------------------
--------CGCTGCAGACTTGGT-GTTGGAGGCCCTTCGGGGGCATTCAGT---GCCGGAGAGAAGTTGTTAAGTCTGCT
ACTTGGGGAGTACGTCCGCAAGGATGAAACTTAAAGGAATTGGCGGGGGAGCACCGCA-ACGGGAGGAGCG-TGCGGTTT
AATTGGATTCAACACCGGAAAACTCACCAGGG-GAGACTGTTACATGAAAGCCA--------------------------
---------------------GGCTAATGACTTTGCTAGATT----TTCAGAGAGGTGGTGCATGGCCGTCGTCAGTTCG
TACCG-TAAGGCG--TTCTCTTAAGTGAGATAAC----------------------------------------------
--------------------------------------------------------------------------------
----------G------AACGAGACCCTCACCAATAGTTGCTAC---TTTACCCTCC---GG-GGTGGAGGCACACTAT-
TGGGACCGC----TGGCGCTA-AGTCAGAGGAAGGAGAGGTCAACGGTAGGTCA-GTATGC-CCCGAATCTCCTG-GGCT
ACACGCGCGCTACAAAGGGCGGGACAATGGGCTC-CGACACCGAAAGGTGAAG---------------------------
---------------------------------------------------GCAATCT-CGAAACCCGTCCGTAGTTCGG
ATTGAGGGTTGTAACTCACCCTCATGAAG--CTGGATTCCGTAGTAATCGCGAATCAACAACT-CGCGGTGAATATGCCC
CTGCTCCTTGCACACACCGCCCGTCAAACCATCCGAGTTGGGTTTCAGTGAGGCTGCCTTTAAT----------------
---------------------------TGGGGTTGTCGAACTGAGATTTAGCAAGGAAGGTTAAGTCGTAACAAGGTATC
TGTAGGGGAACCTGCAGATGGATCACCTCCT-------------------------------------------------
------------------------------------------------------
>lcl|CP015363.1_rrna_55 [gene=rrs] [locus_tag=FAD_r1610] [db_xref=RFAM:RF01959] [product=16S ribosomal RNA]
[location=1594125..1595594] [gbkey=rRNA] Fac
--------------------------------------------------------------------------------
--------------------------------------------------------------------------------
--------------------------------------------------------------------------------
--------------------------------------------------------------------------------
--------------------------------------------------------------------------------
--------------------------------------------------------------------------------
--------------------------------------------------------------------------------
--------------------------------------------------------------------------------
-----------------------------------------------------ACTCCGGTTGATCCTGCCGGCGGCCAC
TGCTATCAAGTTCCGACTAAGCCATGCGAGTCAAGGTAT------------------------------------CGTAA
GATGCCGGCAAACTGCTCAGTAACA-CGTGGATAATCTAACCTTGAGT--------------------------------
-----------------AA-GGGATAACTTCGGGAAA-CTGAAGGTAATACCTTAT-AATTG--CTTAA-AACTGGAATG
TTTTTG-CAATAAAAGT----------------------TACGACGCTCAAGGATGAGTCTGCGAC---C----------
------------------TATCAGGTAGTAGGTGGTGT-AATGGACCACCTAGCCTCAGACGGGTACGGGCCCTGGGAGG
GGTAGCCCGGAGATGGACTCTGAGACATAAGTCCAGGCCCTACGGG--GCGCAGCAGGCGCGAACACTGTGCAATGCGCG
AAAGCGCGAC---ACGG---G-----------GAGCTTGAGTGTCTTGGCAT--AGCCA--------AGACTTTT-----
-------------------------------CTCATTCCTAAAAAGCATGAGGAATAAGTGCTGGGTAAGACGGGTGCCA
GCCGCCGCGGTAACACCCGCAGCACGAGTAGTGGTCACTTTTATTGAGCCTAAAGCGTTCGTAGCCGGTTTTGTAAATCT
TCAGATAAAGCCTGAAGCTTAACTC-CAGAAAG----------------------------------TCTGAAGAGACT-
-----------------------------------GCAAGACTTGAGATCGGGTGAGGTTAAACGTACTTTCAGGGTAGG
GGTAAAATCCTGTAATCCCGGAAGGACGACCAG-----------------------------------------------
-------------------------------------------------------------------TGGCGAAAGCGTT
TAACTAGAACGAATCT-------------------------------------------------------GACGGTAAG
GAACGAAGGCTAGGGTAGCAAACCGGATTAGATACCCGGGTAGTCCTAGCTGTAAA------------------------
--------------------------------------------------------------------------------
--------CATTGCCCATTTGAT-GTTGCTTTTCCGTTGAGGGAAGGCAGT---GTCGGAGCGAAGGTGTTAAATGGGCC
GCTTGGGAAGTATGGTCGCAAGACTGAAACTTAAAGGAATTGGCGGGGGAGCACCGCA-ACGGGAGGAATG-TGCGGTTT
AATTGGATTCAACGCCGGAAAACTCACCGGGA-ACGACCTGTGCATGAGAGTCA--------------------------
```

---------------------ACCTGACGAGCTTACTCGATA----GCAGGAGAGGTGGTGCATGGCCGTCGTCAGCTCG
TACCG-TAGGGCG--TTCACTTAAGTGTGATAAC----------------------------------------------
--------------------------------------------------------------------------------
----------G------AGCGAGACCCACATCTTTAATTGCAAA---TGTATATGAG---AA-TATGCATGCACTTTAG-
AGAAACCGC----CAGCGCTA-AGCTGGAGGAAGGAGTGGTCGACGGCAGGTCA-GTACGC-CCCGAATTTCCCG-GGCT
ACACGCGCATTACAAAGAACGGGACAATACGTTG-CAACCTCGAAAGAGGAAG---------------------------
---------------------------------------------------CTAATCG-CGAAACCCGTCCATAGTTAGG
ATTGAGGGCTGTAACTCGCCCTCATGAAT--CTGGATTCCGTAGTAATCGCGGGTCAACAACC-CGCGGTGAACATGCCC
CTGCTCCTTGCACACACCGCCCGTCAAACCATCCGAGTTGGTGTTGGATGAGGTTTAATTCGAGA---------------
---------------------------GGGTTAAATCAAATCTGATGTCGGTGAGGAGGGTTAAGTCGTAACAAGGTATC
CGTAGGGGAACCTGCGGATGGATCACCTCCA-------------------------------------------------
------------------------------------------------------
>lcl|BA000011.4_rrna_48 [gene=TVG1518472] [product=16S ribosomal RNA] [location=complement(1518773..1520246)] [gbkey=rRNA] Tvo
--------------------------------------------------------------------------------
--------------------------------------------------------------------------------
--------------------------------------------------------------------------------
--------------------------------------------------------------------------------
--------------------------------------------------------------------------------
--------------------------------------------------------------------------------
--------------------------------------------------------------------------------
--------------------------------------------------------------------------------
--------------------------------------------------TTAACTCCGGTCGATCCTGCCGGCGGTCAC
TGCTATCAGGTTCCGACTAAGCCATGCAAGTCACGGGGC------------------------------------CGTAA
GGCACCGGCGAACAGCTCAGTAACA-CGTGGATAATTTACCCTCAGGT--------------------------------
-----------------GG-GGTATAACCTCGGGAAA-CTGAGGCTAATCCCCCAT-AGTCA--TTACA-AACTGGAACG
TTGTAA-TGATGAAAGC----------------------TCCGGCGCCTGAGGATAAGTCTGCGGC---C----------
------------------TATCAGGTAGTAGGTGGTGT-AAAGGACCACCTAGCCTAAGACGGGTACGGGCCCTGAAAGG
GGGAGCCCGGAGATGGACTCTGAGACAACAGTCCAGGCCCTACGGG--GCGCAGCAGGCGCGAAAACTGTGCAATGCGCG
AAAGCGCGAC---ACGG---G-----------GAACCTGAGTGCCTTGACTTTTCGTCA--------AGGCTTTT-----
-------------------------------CTGATGCCTAAAAAGCATCAGGAATAAGGGCTGGGCAAGACGGGTGCCA
GCCGCCGCGGTAACACCCGCAGCTCGAGTGGTGATCACTTTTATTGAGTCTAAAGCGTCCGTAACCGGTCTTATAAATCT
TCAGATAAATTCTCCCGCTTAACGG-AAGAACT----------------------------------TCTGAAGAGACT-
-----------------------------------GTAAGACTTGGGACCGGGTGAGGTTGAATGTACTTTCAGGGTAGG
GGTAAAATCCTGTAATCCTGAAAGGACGACCGG-----------------------------------------------
-------------------------------------------------------------------TGGCGAAAGCGTT
CAACTAGAACGGATCC-------------------------------------------------------GACGGTGAG
GGACGAAGGCTAGGGGAGCAAACCGGATTAGATACCCGGGTAGTCCTAGCTGTAAA------------------------
--------------------------------------------------------------------------------
--------CGCTGCCCACTTGGT-GTTGCTTCTCCGTTGAGGGGGAGCAGT---GCCGTAGCGAAGGTGTTAAGTGGGTC
ACTTGGGGAGTACGGCCGCAAGGCTGAAACTTAAAGGAATTGGCGGGGGAGCACCGCA-ACGGGAGGAGCG-TGCGGTTT
AATTGGATTCAACGCCGGAAAACTCACCGGGA-GCGACCTTCGGATGAGGGTCA--------------------------
---------------------ACCTGACGAATTTACCCGATA----GAAGGAGAGGTGGTGCATGGCCGTCGTCAGCTCG
TACCG-TAGGGCG--TTCACTTAAGTGTGATAAC----------------------------------------------
--------------------------------------------------------------------------------
----------G------AGCAAGACCCCCATCTCTAATTGCTAA---GCTTTCTTAG---CG-GGAAGCTGCACTTTAG-
AGGGACCGC----CAGCGCTA-AGCTGGAGGAAGGAGGGGTCGACGGCAGGTCA-GTACGC-CCCGAATCTCCCG-GGCT
ACACGCGCGCTACAAAGGACGGGACAATGAGTTG-CAACCTCGAAAGGGGAAG---------------------------
---------------------------------------------------CTAACCT-CGAAACCCGTTCGTAGTCAGG
ACTGAGGGCTGTAACTCGCCCTCACGAAT--GTGGATTCCGTAGTAATCGTAGGTCAACAGCC-TACGGTGAATATGCCC
CTGCTCCTTGCACACACCGCCCGTCAAACCATCCGAGCTGGTGTTGGATGAGGGTCTGTC-CATT---------------
---------------------------GGATGGATTCGAATCTGATGTCAGTGAGGAGGGTTAAGTCGTAACAAGGTATC
CGTAGGGGAACCTGCGGATGGATCACCTCCA-------------------------------------------------
------------------------------------------------------
>lcl|AE000782.1_rrna_39 [locus_tag=AF_r03] [product=16S ribosomal RNA] [location=complement(1788987..1790478)] [gbkey=rRNA] Afu
--------------------------------------------------------------------------------
--------------------------------------------------------------------------------
--------------------------------------------------------------------------------
--------------------------------------------------------------------------------
--------------------------------------------------------------------------------
--------------------------------------------------------------------------------
--------------------------------------------------------------------------------
--------------------------------------------------------------------------------
-----------------------------------------------------ATTCTGGTTGATCCTGCCAGAGGCCGC
TGCTATCCGGCTGGGACTAAGCCATGCGAGTCAAGGGGCTTGTATCCCTTCG--------------------GGGATGCA
AGCACCGGCGGACGGCTCAGTAACA-CGTGGACAACCTGCCCTCGGGT--------------------------------
-----------------GG-GGGATAACCCCGGGAAA-CTGGGGCTAATCCCCCAT-AGGGG--ATGGG-TACTGGAATG
TCCCAT-CTCCGAAAGC-------------G--------CTTAGCGCCCGAGGATGGGTCTGCGGC---G----------
------------------GATTAGGTTGTTGGTGGGGT-AACGGCCCACCAAGCCGAAGATCCGTACGGGCCATGAGAGT
GGGAGCCCGGAGATGGACCCTGAGACACGGGTCCAGGCCCTACGGG--GCGCAGCAGGCGCGAAACCTCCGCAATGCGGG
AAACCGCGAC---GGGG---T-----------CAGCCGGAGTGCTCGCGCATCGC-GCG--------GGCTGTCG-----
-------------------------------GGGTGCCTAAAAAGCACCCCACAGCAAGGGCCGGGCAAGGCCGGTGGCA
GCCGCCGCGGTAATACCGGCGGCCCGAGTGGCGGCCACTTTTATTGGGCCTAAAGCGTCCGTAGCCGGGCTGGTAAGTCC
TCCGGGAAATCTGGCGGCTTAACCG-TCAGACTG---------------------------------CCGGAGGATACT-
-----------------------------------GCCAGCCTAGGGACCGGGAGAGGCCGGGGGTATTCCCGGAGTAGG
GGTGAAATCCTGTAATCCCGGGAGGACCACCTG-----------------------------------------------
-------------------------------------------------------------------TGGCGAAGGCGCC
CGGCTGGAACGGGTCC-------------------------------------------------------GACGGTGAG
GGACGAAGGCCAGGGGAGCGAACCGGATTAGATACCCGGGTAGTCCTGGCTGTAAA------------------------
--------------------------------------------------------------------------------
--------CGATGCGGACTAGGT-GTCACCGAAGCTACGAGCTTCGGTGGT---GCCGGAGGGAAGCCGTTAAGTCCGCC
GCCTGGGGAGTACGGCCGCAAGGCTGAAACTTAAAGGAATTGGCGGGGGAGCACTACA-ACGGGTGGAGCC-TGCGGTTT
AATTGGATTCAACGCCGGGAAGCTTACCGGGG-GAGACAGCGGGATGAAGGTCG--------------------------
---------------------GGCTGAAGACCTTACCAGACT----AGCTGAGAGGTGGTGCATGGCCGCCGTCAGTTCG
TACTG-TGAAGCA--TCCTGTTAAGTCAGGCAAC----------------------------------------------
--------------------------------------------------------------------------------

----------G------AGCGAGACCCGCGCCCCCAGTTGCCAG---CGGTTCCCTTCGGGG-AAGCCGGGCACACTGG-
GGGGACTGC----CGGCGCTA-AGCCGGAGGAAGGTGCGGGCAACGGCAGGTCC-GTATGC-CCCGAATCCCCCG-GGCT
ACACGCGGGCTACAATGGCCGGGACAATGGGTAC-CGACCCCGAAAGGGGTAG---------------------------
---------------------------------------------------GTAATCCCCTAAACCCGGTCTAACCTGGG
ATCGAGGGCTGCAACTCGCCCTCGTGAAC--CTGGAATCCGTAGTAATCGCGCCTCAAAATGG-CGCGGTGAATACGTCC
CTGCTCCTTGCACACACCGCCCGTCAAGCCACCCGAGTGGGCCAGGGGCGAGGGGGTGGCCCTA----------------
---------------------------GGCCACCTTCGAGCCCAGGGTCCGCGAGGGGGGCTAAGTCGTAACAAGGTAGC
CGTAGGGGAATCTGCGGCTGGATCACCTCCT-------------------------------------------------
------------------------------------------------------
>lcl|AE009950.1_rrna_7 [locus_tag=PF_r001] [product=16S ribosomal RNA] [location=136930..138426] [gbkey=rRNA]
Pfu
--------------------------------------------------------------------------------
--------------------------------------------------------------------------------
--------------------------------------------------------------------------------
--------------------------------------------------------------------------------
--------------------------------------------------------------------------------
--------------------------------------------------------------------------------
--------------------------------------------------------------------------------
--------------------------------------------------------------------------------
-----------------------------------------------------ATTCCGGTTGATCCTGCCGGAGGCCAC
TGCTATGGGGGTCCGACTAAGCCATGCGAGTCAAGGGGGCGTCCCTTC--------------------------TGGGAC
GCCACCGGCGGACGGCTCAGTAACA-CGTCGGTAACCTACCCTCGGGA--------------------------------
-----------------GG-GGGATAACCCCGGGAAA-CTGGGGCTAATCCCCCAT-AGGCC--TGGGG-TACTGGAAGG
TCCCCA-GGCCGAAAGG-------------GAGCCGTAAGGCTCCGCCCGAGGATGGGCCGGCGGC---C----------
------------------GATTAGGTAGTTGGTGGGGT-AACGGCCCACCAAGCCGAAGATCGGTACGGGCCGTGAGAGC
GGGAGCCCGGAGATGGACACTGAGACACGGGTCCAGGCCCTACGGG--GCGCAGCAGGCGCGAAACCTCCGCAATGCGGG
AAACCGCGAC---GGGG---G-----------GACCCCCAGTGCCGTGCC-TCTG-GCA--------CGGCTTTT-----
-------------------------------CCGGAGTGTAAAAAGCTCCGGGAATAAGGGCTGGGCAAGGCCGGTGGCA
GCCGCCGCGGTAATACCGGCGGCCCGAGTGGTGGCCACTATTATTGGGCCTAAAGCGGCCGTAGCCGGGCCCGTAAGTCC
CTGGCGAAATCCCACGGCTCAACCG-TGGGGCTC---------------------------------GCTGGGGATACT-
-----------------------------------GCGGGCCTTGGGACCGGGAGAGGCCGGGGGTACCCCCGGGGTAGG
GGTGAAATCCTATAATCCCGGGGGGACCGCCAG-----------------------------------------------
-------------------------------------------------------------------TGGCGAAGGCGCC
CGGCTGGAACGGGTCC-------------------------------------------------------GACGGTGAG
GGCCGAAGGCCAGGGGAGCGAACCGGATTAGATACCCGGGTAGTCCTGGCTGTAAA------------------------
--------------------------------------------------------------------------------
--------GGATGCGGGCTAGGT-GTCGGGCGAGCTTCGAGCTCGCCCGGT---GCCGTAGGGAAGCCGTTAAGCCCGCC
GCCTGGGGAGTACGGCCGCAAGGCTGAAACTTAAAGGAATTGGCGGGGGAGCACTACA-AGGGGTGGAGCG-TGCGGTTT
AATTGGATTCAACGCCGGGAACCTCACCGGGG-GCGACGGCAGGATGAAGGCCA--------------------------
---------------------GGCTGAAGGTCTTGCCGGACG----CGCCGAGAGGAGGTGCATGGCCGCCGTCAGCTCG
TACCG-TGAGGCG--TCCACTTAAGTGTGGTAAC-----------------------------------------------
--------------------------------------------------------------------------------
----------G------AGCGAGACCCGCGCCCCCAGTTGCCAG---TCCCTCCCGCTC-GG-GAGGGAGGCACTCTGG-
GGGGACTGC----CGGCGATA-AGCCGGAGGAAGGGGCGGGCGACGGTAGGTCA-GTATGC-CCCGAAACCCCCG-GGCT
ACACGCGCGCTACAATGGGCGGGACAATGGGACC-CGACCCCGAAAGGGGAAG---------------------------
---------------------------------------------------GGAATCCCCTAAACCCGCCCTCAGTTCGG
ATCGCGGGCTGCAACTCGCCCGCGTGAAG--CTGGAATCCCTAGTACCCGCGTGTCATCATCG-CGCGGCGAATACGTCC
CTGCTCCTTGCACACACCGCCCGTCACTCCACCCGAGCGGAGTCCGGGTGAGGCCTGATCTCCTTC--------------
------------------------GGGAGGTCAGGTCGAGCCCGGGCTCCGTGAGGGGGGAGAAGTCGTAACAAGGTAGC
CGTAGGGGAACCTACGGCTCGATCACCTCCT-------------------------------------------------
------------------------------------------------------
>lcl|CP001941.1_rrna_35 [locus_tag=Aboo_R0037] [product=16S ribosomal RNA] [location=1422348..1423820]
[gbkey=rRNA] Abo
--------------------------------------------------------------------------------
--------------------------------------------------------------------------------
--------------------------------------------------------------------------------
--------------------------------------------------------------------------------
--------------------------------------------------------------------------------
--------------------------------------------------------------------------------
--------------------------------------------------------------------------------
--------------------------------------------------------------------------------
-------------------------------------------------------TCCGGTTGATCCTGCCGGAGGCCAC
TGCTATCGGGCTCCGACTAAGCCATGCGAGTTTAGGGGCTCC----TT----------------------------CGGG
GGCACCGGCGGACGGCTCAGTAACA-CGTGGACAACCTACCCTCGGGT--------------------------------
-----------------GG-GGGATAACCTCGGGAAA-CTGAGGCTAATACCCCAT-AGGGG--AAAGG-TACTGGAAGG
TCCTTT-CCCTGAAAGC----------------------TCCGGCGCCCGAGGATGGGTCTGCGGC---C----------
------------------TATCAGGTAGTTGGTGGTGT-AAAGGACCACCAAGCCTGCGACGGGTACGGGCCCTGAGAGG
GGTAGCCCGGAGATGGACTCTGAGACACGAGTCCAGGCCCTACGGG--GCGCAGCAGGCGCGAAACCTTCGCAATGCGCG
AAAGCGCGAC---GAGG---G-----------GAGCCCGAGTGCCCTTCCTTTTG-GGA--------GGGCTTTT-----
-------------------------------CCGGAGTCTAAAAAGCTCCGGGAATAAGGGCTGGGCAAGACGGGTGCCA
GCCGCCGCGGTAATACCCGCAGCTCGAGTGGTGGCCACTATTATTGAGCCTAAAGCGTCCGTAGCCGGACCCGCAAATCC
CTGGGTAAATCCTGGTGCTCAACAT-CAGGAAT----------------------------------TCCGGGGAGACT-
-----------------------------------GCGGGTCTTGGGACCGGGAGAGGCCGGAGGTACTCCCGGGGTAGG
GGTAAAATCCTGTAATCCCGGGGGGACCACCTG-----------------------------------------------
-------------------------------------------------------------------TGGCGAAAGCGTC
CGGCTTGAACGGCTCC-------------------------------------------------------GACGGTGAG
GGACGAAGGCCAGGGGAGCAAACCGGATTAGATACCCGGGTAGTCCTGGCTGTAAA------------------------
--------------------------------------------------------------------------------
--------CGCTGCCCGCTTGGT-GTTGTCGGTCTCTCGGGGGCCGGCAGT---GCCGGAGCGAAGGTGTTAAGCGGGCC
GCTTGGGGAGTACGGCCGCAAGGCTGAAACTTAAAGGAATTGGCGGGAGAGCACTGCA-ACGGGAGGAGCG-TGCGGTTT
AATTGGATTCAACGCCGGAAAACTCACCGGGG-GCGACCGGCACATGTGGGCCA--------------------------
---------------------GGCTGACGACCTTGCCCGAATAAGAGCCGGAGAGGTGGTGCATGGCCATCGTCAGCTCG
TACCG-TAGGGCG--TTCACTTAAGTGTGATAAC-----------------------------------------------
--------------------------------------------------------------------------------
----------G------AGCGAGACCCCCGTCCCCAGTTGCCAC---CTGTCCCTCC---GG-GGGCAGAGCACACTGG-
GGAGACCGC----CAGCGCTA-AGCTGGAGGAAGGTGGGGTCAACGGTAGGTCA-GTACGC-CCCGAATCCCCCG-GGCT
ACACGCGCGCTACAAAGGGCGGGACAATGGGTTG-CGACCCCGAAAGGGGGAG---------------------------

---------------------------------------------------CTAATCC-TGAAACCCGTCCGTAGTTCAG
ATCGAGGGCTGTAACTCGCCCTCGTGAAG--CTGGATTCCGTAGTAATCGCGGGTCAACATCC-CGCGGTGAATGCGCCC
CTGCTCTTTGCACACACCGCCCGTCAAACCATCCGAGTTGGCTCTAGGTGAGGCCCTGTTCTAT----------------
---------------------------GGGCAGGGTCGAACCTGGGGTCAGCGAGGAGGGTTAAGTCGTAACAAGGTATC
CGTAGGGGAACCTGCGGATGGATCACCT----------------------------------------------------
------------------------------------------------------
>lcl|CP003168.1_rrna_42 [locus_tag=AciM339_1371] [product=16S ribosomal RNA] [location=1246328..1247697]
[gbkey=rRNA] Asp
--------------------------------------------------------------------------------
--------------------------------------------------------------------------------
--------------------------------------------------------------------------------
--------------------------------------------------------------------------------
--------------------------------------------------------------------------------
--------------------------------------------------------------------------------
--------------------------------------------------------------------------------
--------------------------------------------------------------------------------
-----------------------------------------------------------------------GGAGGCCAC
TGCTATCGGGCTCCGACTAAGCCATGCGAGTTTAGGGGCTCC----TT----------------------------CGGG
AGCACCGGCGGACTGCTCAGTAACA-CGTGGACAACCTACCCTCGGGT--------------------------------
-----------------GG-GGGATAACCTCGGGAAA-CTGAGGCTAATACCCCAT-AGGCC--TCGGA-TACTGGAAGG
TTCCGA-GACCGAAAGC-----------------------TCCGGCGCCCGAGGATGGGTCTGCGGC---C----------
------------------TATCAGGTAGTTGGTGGTGT-AAAGGACCACCAAGCCTACGACGGGTACGGGCCCTGAGAGG
GGAAGCCCGGAGATGGACTCTGAGACACGAGTCCAGGCCCTACGGG--GCGCAGCAGGCGCGAAACCTTCGCAATGCGCG
AAAGCGCGAC---GAGG---G-----------GAGCCCGAGTGCCCTTCCTTTCG-GGA--------GGGCTTTT-----
-------------------------------CTGGAGCCTAAAAAGCTCCAGGAATAAGGGCTGGGCAAGACGGGTGCCA
GCCGCCGCGGTAATACCCGCAGCTCGAGTGGTGGCCACTATTATTGAGCCTAAAGCGTCCGTAGCCGGACCCGTAAATCC
CTGGGTAAATCCTGGCGCTCAACGT-CAGGAAT----------------------------------TCCGGGGAGACT-
-----------------------------------GCGGGTCTTGGGACCGGGAGAGGCCGGAGGTACTCCCGGGGTAGG
GGTAAAATCCTGTAATCCCGGGGGGACCACCTG-----------------------------------------------
-------------------------------------------------------------------TGGCGAAAGCGTC
CGGCTTGAACGGCTCC-------------------------------------------------------GACGGTGAG
GGACGAAGGCCAGGGGAGCAAACCGGATTAGATACCCGGGTAGTCCTGGCTGTAAA------------------------
--------------------------------------------------------------------------------
--------CGCTGCCCGCTTGGT-GTTGTCGGTCTTTCGGGGGCCGGCAGT---GCCGGAGCGAAGGTGTTAAGCGGGCC
GCTTGGGGAGTACGGCCGCAAGGCTGAAACTTAAAGGAATTGGCGGGAGAGCACTGCA-ACGGGAGGAGCG-TGCGGTTT
AATTGGATTCAACGCCGGAAAACTCACCGGGG-GCGACCGGCACATGTGGGCCA--------------------------
---------------------GGCTGACGACCTTGCCCGAATAAGAGCCGGAGAGGTGGTGCATGGCCATCGTCAGCTCG
TACCG-TAGGGCG--TTCACTTAAGTGTGATAAC----------------------------------------------
--------------------------------------------------------------------------------
----------G------AGCGAGACCCCCGTCCCCAGTTGCTAC---CTGTCCCTCC---GG-GGGCAGAGCACACTGG-
GGAGACCGC----TGGCGCTA-AGCCAGAGGAAGGTGGGGTCAACGGTAGGTCA-GTACGC-CCCGAATCCCCCG-GGCT
ACACGCGCGCTACAAAGGGCGGGACAATGGGTTG-CGACCCCGAAAGGGGGAG---------------------------
---------------------------------------------------CTAATCC-TGAAACCCGTCCGTAGTTCAG
ATCGAGGGCTGTAACTCGCCCTCGTGAAG--CTGGATTCCGTAGTAATCGCGCCTCAACATGG-CGCGGTGAATGCGCCC
CTGCTCTTTGCACACACCGCCCGTCAAACCATCCGAGTTGGCTCTAGGTGAGGCTCTG----------------------
--------------------------------------------------------------------------------
--------------------------------------------------------------------------------
------------------------------------------------------
>lcl|CP002278.1_rrna_4 [locus_tag=Mfer_R0004] [product=16S ribosomal RNA] [location=complement(26992..28471)]
[gbkey=rRNA] Mfe
--------------------------------------------------------------------------------
--------------------------------------------------------------------------------
--------------------------------------------------------------------------------
--------------------------------------------------------------------------------
--------------------------------------------------------------------------------
--------------------------------------------------------------------------------
--------------------------------------------------------------------------------
--------------------------------------------------------------------------------
-----------------------------------------------------ACTCCGTTTGATCCTGGCGGAGGCCAC
TGCTATGGGGGTCCGACTAAGCCATGCAAGTCGAACGGGCCT-----------------------------------TGT
GCCCGTGGCGAACGGCTCAGTAACA-CGTGGACAACCTACCCTGGGGT--------------------------------
-----------------CC-GGGATAACCCCGGGAAA-CTGGGGCTAATCCCGGAT-AGGCG--AGGTC-TCCTGGAATG
GGGCCT-CGCCGAAAGG--------------TC-----TTTTGACCGCCCCAGGATGGGTCTGCGGC---C----------
------------------GATTAGGTAGTTGGTAGGGT-AACGGCCTACCAAGCCTACGATCGGTACGGGTTGTGAGAGC
AAGAGCCCGGAGACGGGGCCTGAGACAAGGCCCCGGGCCCTACGGG--GCGCAGCAGGCGCGAAAACTCCGCAATGCGCG
AAAGCGCGAC---GGGG---G-----------GACCCCCAGTGCCACTCCGTAAG-GAG--------TGGCTTTT-----
-------------------------------CCGGAGTGTAAAAAGCTCCGGGAATAAGGGCTGGGCAAGACCGGTGCCA
GCCGCCGCGGTAACACCGGCAGCCCGAGTGGTGGCCGCGTTTATTGGGCCTAAAGCGTCCGTAGCCGGTCCGGTAAGTCT
CCGGTGAAAGCCCGCAGCTCAACTG-CGGGAGTA---------------------------------GCCGGAGATACT-
-----------------------------------GCCGGACTTGGGGCCGGGAGAGGCCGGAGGTACCCCCGGGGTAGG
GGTGAAATCCTGTAATCCCGGGGGGACCACCTG-----------------------------------------------
-------------------------------------------------------------------TGGCGAAGGCGTC
CGGCTGGAACGGGCCC-------------------------------------------------------GACGGTGAG
GGACGAAAGCCAGGGGAGCGAACCGGATTAGATACCCGGGTAGTCCTGGCCGTAAA------------------------
--------------------------------------------------------------------------------
--------CGATGCGGACTTGGT-GTTGGGGCAACCTCGAGTTGCCCCAGT---GCCGAAGGGAAGCCGTTAAGTCCGCC
GCCTGGGGAGTACGGCCGCAAGGCTGAAACTTAAAGGAATTGGCGGGGGAGCACCACA-ACGCGTGGAGCC-TGCGGTTT
AATTGGATTCAACGCCGGACACCTCACCGGGG-GCGACGGCAGGATGATGGCCA--------------------------
---------------------GGTTGATGACCTTGCCTGACG----AGCCGAGAGGAGGTGCATGGCCGCCGTCAGCTCG
TACCG-TGAGGCG--TCCTGTTAAGTCAGGCAAC----------------------------------------------
--------------------------------------------------------------------------------
----------G------AGCGAGACCCGCGCCCCTAGTTGCCAG---CGGGTCCCGTAAGGG-ACGCCGGGCACACTAG-
GGGGACCGC----CAGCGATA-AGCTGGAGGAAGGTGCGGGCGACGGTAGGTCC-GTATGC-CCCGAAACCCCCG-GGCT
ACACGCGGGCTACAATGGCCGGGACAATGGGTAC-CGACCCCGAAAGGGGGAG---------------------------
---------------------------------------------------GTAATCCCATAAACCCGGCCGTAGTTCGG
ATCGAGGGCTGCAACTCGCCCTCGTGAAG--CTGGAATGCGTAGTAATCGCGGGTCACTATCC-CGCGGTGAATACGTCC
CTGCTCCTTGCACACACCGCCCGTCACGCCACCCAAACGGGGTTCGGATGAGGCCATGGCCTCT----------------

```
---------------------------GGCCATGGTCGAATCCGGGCCCCGTGAGGAGGGCGAAGTCGTAACAAGGTAGC
CGTAGGGGAACCTGCGGCTGGATCACCTCCT-------------------------------------------------
------------------------------------------------------
>lcl|L77117.1_rrna_24 [locus_tag=MJ_r03] [product=Ribosomal RNA] [location=638452..639929] [gbkey=rRNA] Mja
--------------------------------------------------------------------------------
--------------------------------------------------------------------------------
--------------------------------------------------------------------------------
--------------------------------------------------------------------------------
--------------------------------------------------------------------------------
--------------------------------------------------------------------------------
--------------------------------------------------------------------------------
--------------------------------------------------------------------------------
-----------------------------------------------------ATTCCGGTTGATCCTGCCGGAGGCCAC
TGCTATCGGGGTCCGACTAAGCCATGCGAGTCAAGGGGCTCC----CT--------------------------TCGGGG
AGCACCGGCGCACGGCTCAGTAACA-CGTGGCTAACCTACCCTCGGGT--------------------------------
-----------------GG-GGGATAACCTCGGGAAA-CTGAGGCTAATCCCCCAT-AGGGG--AGGAG-GTCTGGAATG
ATCCCT-CCCCGAAAGG-------------CG-------TAAGCCGCCCGAGGATGGGGCTGCGGC---G----------
------------------GATTAGGTAGTTGGTGGGGT-AACGGCCCACCAAGCCTACGATCCGTACGGGCCCTGAGAGG
GGGAGCCCGGAGATGGACACTGAGACACGGGTCCAGGCCCTACGGG--GCGCAGCAGGCGCGAAACCTCCGCAATGCGCG
AAAGCGCGAC---GGGG---G-----------GACCCCGAGTGCCCACGC-CCTG-CGT--------GGGCTTTT-----
-------------------------------CCGGAGTGTAAACAGCTCCGGGAATAAGGGCTGGGCAAGTCCGGTGCCA
GCAGCCGCGGTAATACCGGCGGCCCAAGTGGTGGCCACTGTTATTGGGCCTAAAGCGTCCGTAGCCGGCCCGGTAAGTCT
CTGCTTAAATCCTGCGGCTCAACCG-CAGGGCT----------------------------------GGCAGAGATACT-
------------------------------------GCCGGGCTTGGACCGGGAGAGGCCGGGGGTACCCCAGGGGTAGC
GGTGAAATGCGTTGATCCCTGGGGGACCACCTG-----------------------------------------------
--------------------------------------------------------------------TGGCGAAGGCGCC
CGGCTGGAACGGGTCC-------------------------------------------------------GACGGTGAG
GGACGAAGGCCAGGGGAGCAAACCGGATTAGATACCCGGGTAGTCCTGGCTGTAAA------------------------
--------------------------------------------------------------------------------
--------CTCTGCGGACTAGGT-GTCGCGTCGGCTTCGGGCCGACGCGGT---GCCGAAGGGAAGCCGTTAAGTCCGCC
GCCTGGGGAGTACGGTCGCAAGACTGAAACTTAAAGGAATTGGCGGGGGAGCACTACA-ACGGGTGGAGCC-TGCGGTTT
AATTGGATTCAACGCCGGGCATCTTACCAGGG-GCGACGGCAGGATGAAGGCCA--------------------------
---------------------GGTTGACGACCTTGCCAGACG----CGCCGAGAGGTGGTGCATGGCCGTCGTCAGCTCG
TACCG-TGAGGCG--TCCTGTTAAGTCAGGTAAC----------------------------------------------
--------------------------------------------------------------------------------
----------G------AGCGAGACCCGTGCCCCATGTTGCTAC---CTCCTCCTCC---GG-GAGGAGGGCACTCATG-
GGGGACCGC----CGGCGCTA-AGCCGGAGGAAGGTGCGGGCAACGACAGGTCC-GCATGC-CCCGAATCCCCTG-GGCT
ACACGCGGGCTACAATGGCCGGGACAATGGGACG-CGACCCCGAAAGGGGGAG---------------------------
---------------------------------------------------CGAATCCCCTAAACCCGGTCGTAGTCCGG
ATCGAGGGCTGTAACTCGCCCTCGTGAAG--CCGGAATCCGTAGTAATCGCGCCTCACCATGG-CGCGGTGAATGCGTCC
CTGCTCCTTGCACACACCGCCCGTCACGCCACCCGAGTTGAGCCCAAGTGAGGCCCTGTCCGCAA---------------
---------------------------GGGCAGGGTCGAACTTGGGTTCAGCGAGGGGGGCGAAGTCGTAACAAGGTAGC
CGTAGGGGAACCTGCGGCTGGATCACCTCC--------------------------------------------------
------------------------------------------------------
>lcl|PFPH01000326.1_rrna_11 [locus_tag=COX84_05885] [product=16S ribosomal RNA] [location=15..1774] [gbkey=rRNA]
Mic
--------------------------------------------------------------------------------
--------------------------------------------------------------------------------
--------------------------------------------------------------------------------
--------------------------------------------------------------------------------
----------------------------------------------------------------------CCGTAACTCC
AGTTGATCGTGCTGGAGGGT---ACTGCTATCGAAGTCA-----------------------------------------
---------------------------GATTAAGCCATGCAAGTCCATACGTGCCCATGTACGTATGG---CGAACGGCT
GA-----------------GTAACGCGTGGTCAATCTACCCTGG------------------GGTCGGGCATAACCGCGG
GAAACTGCGGGTAATTC-----------C-CGATAGGTGAGCGATGCTG---GAATGCTTGTTCTCCTAAAGGATGTGCA
AATTGGAGCATAATCGCCCTAGGATGAGACTGCGTCGGATTATGGTAGTTG----------------------------G
TGG----GGTAACGGCCCACCAAGCTGATAATCCGTAGGGGCAATGGGAGTTGTAGCCCCCAGAAGGACACTGAGACAAG
GGTCCTAGCCCTACG--GG-GTGCAGCAGGCGCGAAA-CCTTCTCAATGCGCGCAA-GCGTGAAGAGGTGACTTCGAGTG
ATAGTT-GCTTCATAGTGATTATC--TTTTGCC-AAGTCCAAACAGCTTGGAGA------------------AT------
---------------------------------------AAGATCTGGGTAAGACTAGTGCCAGCCGCCGCGGTAACACT
AGCAGATCAAGT-------GGTACCCACGATTATTGGGTCTAAAGCGCTCGTAGCCGGTTAGATAAATT-----------
----------------------------------------------CACTGTGAAAGAGCC---TCTTTTCTTTT-----
-------------------------------GTAAAAGAAAGGACCTTCGGGAGAAAAGAAAACAATTAGCATATTTTCC
-AATGGTCTTTCCTTCAAAAATAAAAAGAGGTTCCAGGAAATCCTCAGGCTTA---------------------------
-------------------ACCTAG-AGGGCGT----------------------------------GCAGTGAACACT-
-----------------------------------GTCTAGCTTGAGAGCGGGGGAGGTCAGAAGTACTACCGGGGTAGG
GGTAAAATCCTGTGAGCCTGGTAGGACTAACGG-----------------------------------------------
-------------------------------------------------------------------TGGCGAAAGCGTC
TGACCAAAACGCGTCTGACGGTGAGGAGCGAAGGCCGGGGGATCAAATCAGTCTTCAGTTTTCAGTTTCGGGTTTCGAAG
CTGGTTGACGGCTGAGATCAAAAGGGATTAGATACCCCTGTAGTCCCGGCAGTAAA------------------------
--------------------------------------------------------------------------------
--------TTATGCGGACTTGGGTGCTGCAGTACTCTAGAGGTACTGCAGT---GCTATAGCGAAGGCGTTAAGTCCGCC
ACCTGGGGAGTACGGCCGCAAGGTTGAAACTTAAACGAATTGGCGGGGGAGCACCACA-AGGGGTGGATGC-TGCGGTTC
AATTGGATACAACGCCAGGAATAT-TACCAGGAGCGACGGCAGAATGAAGGCCA--------------------------
---------------------GGCTGAAGACCTTGCTAGACA----AGCCGAGTGGTGCTGCATGGCCATCGTCAGCTCG
TGGCG-TGACAAC--ATCAGCCTAATCAGTTTTGAGTTACGAGTTTCGAA-----------------------------A
CCCGAAACCCAGAACC--------CGAAAC-------TATC-----ATGGGTTGGTGTTGCGCTAAAGCTGTCTGGTTAA
GTCCAGCAACG------AGCGAGACCCCTATGTTTAGTTGCCAA---CTACC--TTTT---C-GAAGGTGGTGCACTCTA
GACAGACTG----CCTGCGTA-AGCAGGAGGAAGGAAGGGGCGACGGTAGGTCA-GTATAG-CCCTAATCTCCTG-GGCC
ACACGTGGCATACAATGGATGGGACAATTGGTTG-CAACTCCGAAAGGAGGAG---------------------------
---------------------------------------------------CCAATCCCATAAACCCATCCTCAGTTCGG
ATTGAGGGCTGAAACTCGCCCTCATGAAG--ATGGAATCCCTAGTAATCGCTTGTCAACATCG-AGCGGTGAATATGTCC
CCGCTCCTTGCACACACCGCCCGTCAAGTCACCCAAGCGAAATTATGGTGATGCATAATCCTCTGGATTC----------
----------------------CAGAAGGGTCGTGTTGAACCACAATTTTATGAGGGGGGCTAAGTCGTAACAAGGTATC
CGTAGGGGAACCTGCGGATAGATCACCTCCA-------------------------------------------------
------------------------------------------------------
>lcl|PRDH01000081.1_rrna_32 [locus_tag=C4K48_12820] [product=16S ribosomal RNA] [location=66869..68362]
```

[gbkey=rRNA] Tho
--------------------------------------------------------------------------------
--------------------------------------------------------------------------------
--------------------------------------------------------------------------------
--------------------------------------------------------------------------------
--------------------------------------------------------------------------------
--------------------------------------------------------------------------------
--------------------------------------------------------------------------------
--------------------------------------------------------------------------------
---------------------------------------------------CAACTCTAGTTGATCCTGCTAGAGGACAC
TGCTATCGGCTTGGGGATAAGACATGCAAGTGGAACGGGCTCGCATCCGAG----------------------------T
CCGTC--GCGAACGGCTCAGTAGTG-T--CGCTAACCTACCCTGTGGAGGT-----------------------------
---------------------GGACAACGCCGGGAAA-CTGGCGTCAATCCATCAT-AGGAG--CATCGTTCTGGAAATG
ATGTTG-CTCTCAAAAGGATGTTCCAGGCATGT--GGGACACTCCGCCGCAGGA--------------------------
----CGGGACGGCACCGGATCATACTTGTAAGTGGGGT-AACTGCCCACTTAGGTTATGACCCGAACGGGCGATGTAAGT
CGTGGCCCGGAGAAGGGCACTGAGACAAGGGCCCTAGCCCCACGGG--GCGCAGCAGTTACGAGACCTCACCAATGCGCG
AAAGCGCGAG---TGGG---C-----------TAAGCCGAGTGATGGGGGATAACTCCATC---TGTGGCGGATC-----
-------------------------------CGCAGGGTGGAC------CGCTAGAAAGGAGAGGGCAAGGCTGGTGCCA
GCCGCCGCGGTAAAACCAGCTCTTCGAGTGGTGTCCACAATTATTGAGCTTAAAGCGTTCGTAGCCTGCTTGGTAGGTCC
CTGCTTAAATCCGGTCGCTCAACGA-TTGGTCT----------------------------------GGTAGGGATACC-
-----------------------------------ACCTTGCTTGGGGACGGGAGACGGTGACGGTATTCCGTAGGTAAG
GGTGAAATCTAATGATCTACGGAGGACCACCAG-----------------------------------------------
-------------------------------------------------------------------TGGCGAAGGCGGT
CGTCGAGAACGTGCCC-------------------------------------------------------GACGGTGAG
GAACGAAACCTAGGGGAGCAAATCGGATTAGATACCCGAGTAGTCCTAGGTGTAAA------------------------
--------------------------------------------------------------------------------
--------CGATGTCCGTTAAGTGTTGCATTGG-CCACGAGCCACTGCAGT---GCTGTAGCTAAAGCGATAAACGGACC
GCTTGGGAAGTACGGTCGCAAGACTGAAACTTAAAGGAATTGGCGGGGGAGCACCACA-AGGGGTGGAGCT-TGCGGTTC
AATTGGACTCAACGCCGGAAAACT-TACCCGCCCAGACAGCCGAATGAGAGCCA--------------------------
---------------------GACTGATGATCTTGCTCGACG----AGCTGAGAGCTGGTGCATGGCCATCGTAAATTCG
TGCCG-TGAGGTA--TCAGCTTAAGTGCTGCAAC----------------------------------------------
--------------------------------------------------------------------------------
----------G------AATGAGATCCTCGCCACCTAGTTGCCA---GCCGGGAGTTTAGGCTCTGACGGGGCACACTAG
GGGGACCGC----CAGCGAAA-AGCTGGAGGAAGGTGGGGGCCACGGTAGGTCA-GTATGC-CGCGATATGGGCGGGGCC
ACACGCGTGCTACAATGGCTGGGACAATACGTAA-CAACGCCGAAAGGTGAAG---------------------------
---------------------------------------------------TCA-ATCGCGAAACCCAGTCGTAGTCGGG
ATCGCGGGTTGGAACTCACCCGCGTGAAC--CTGGAATCCCTAGTATCCGCGTGTCACTATCG-CGCGGAGAATACGTCC
CTGCAGGGGACGTGAATCTCACGTACCCGGGGAATTCTTGCACACACCGCCCGTCGCTCCACGCGAGCTGTTCTTGGG--
-----------TGAGGCGTCATCCACTGGGTGATTACGAATCTTGGGACGGCAAGTGGGGAGAAGTCGTAA---------
--------------------------------------------------------------------------------
------------------------------------------------------
>lcl|CP000968.1_rrna_17 [locus_tag=Kcr_R0018] [product=16S ribosomal RNA] [location=1326582..1328013]
[gbkey=rRNA] Kcr
--------------------------------------------------------------------------------
--------------------------------------------------------------------------------
--------------------------------------------------------------------------------
--------------------------------------------------------------------------------
--------------------------------------------------------------------------------
--------------------------------------------------------------------------------
--------------------------------------------------------------------------------
--------------------------------------------------------------------------------
------------------------------------------------------------------------GAGGGAAC
CCCTATCGGGCTCGCACTAAGCCATGCGAGTCTGCTGGGGGCCCCTGC-------------------------------C
CCTGGCGGCGCACGGCTCCGTAATA-CACGGTCAACCTGTCCTGGGGACCGGGA--------------------------
------------------------TAACCTCGGGAAA-CTGAGGTTAATACCGGAT-AGGGG--TGGAT-TCCTGGAATG
GGTCCA-CCCCTAAAGTAGGCGGGGGGACGG----CCCCGCTGAGGCCCCAGGGTGGGACCGTGGC---CT-ATCAG---
------------------------GTAGTAGGTGGGGT-AACGGCCCACCTAGCCTAAGACGGGTACGGGCTCTGAGAGG
AGGAGCCCGGAGATGGGCACTGAGACAAGGGCCCAGGCCCTACGGG--GCGCAGCAGGCGGGAAACTTCCCCAATGCGCG
CAAGCGTGAG---GGAG---T-----------GAGCCCGAGTGCCGCCCGCTGAGGGCGGC---TGTTCCCCTGT-----
-------------------------------GTAAAAAGCAGG------GGGTAGGAAGGGGAGGGTAAGGCTGGTGCCA
GCCGCCGCGGTAAAACCAGCTCCCCGAGGGGTTCCCACGCATACTGGGCCTAAAGCGTCCGTAGCCGGCCCCGTAAGTCC
TCGGTTAAATCCGCCTGAAGA-CAG-GCGGACC----------------------------------GCCGAGGATACT-
-----------------------------------GCGGGGCTAGGGAGCGGGAGGGGCCGAGGGTATTCCGGGGGGAGC
GGTAAAATGCGTAGATCCCCGGAGGACCACCAG-----------------------------------------------
-------------------------------------------------------------------TGGCGAAGGCGCT
CGGCTGGAACGCGTCC-------------------------------------------------------GACGGTGAG
GGACGAAAGCTGGGGGAGCAAACCGGATTAGATACCCGGGTAGTCCCAGCCGTAAA------------------------
--------------------------------------------------------------------------------
--------CGATGCCGGCTAGGTGCCGGCTGAGGTTTCGGCCTCAGCCGGT---GTCGAAGCGAAGGCATTAAGCCGGCC
GCCTGAGGAGTACAGCCGCAAGGCCGAAACTTAAAGGAATTGACGGGGGGGCACCACA-AGGGGTGAATGCCTGCGGCTC
AATTGGACTCAACGCCGGGAATCT-TACCGGGGGCGACAGCAGGATGAAGGTCA--------------------------
---------------------GGCTGAAGACCTTACCTGACG----CGCTGAGGGGTGGTGCATGGCCGTCGCCAGCTCG
TGCCG-TGAGGTG--TCCTGTTAAGTCAGGCAAC----------------------------------------------
--------------------------------------------------------------------------------
----------G------AGCGAGACCCCCGCCCTCAGTTGCCAG---CGGGG--CCTTACGG-CTGGCCGGGCAAACTGG
GGGGACTGC----CGGCGAAG-AGCCGGAGGAAGGAGGGGGCTACGGCAGGTCA-GTATGC-CCCTAATCCCCCG-GGCC
GCACGCGGGCTGCAATGGGCGGGACAGCGGGATG-CGACCCCGAGAGGGGGAG---------------------------
---------------------------------------------------CAAATCCCTGAAACCCGCCCGTGGTTGGG
ATCGAGGGTTGCAACTCGCCCTCGTGAAC--CCGGAATCCCTAGTAACCGCGGTTCTCCATAC-CGCGGTGAATACGTCC
CTGCCCCTTGTACACACCGCCCGTCAACCCACCCGAGTGGACTTGGGGCGAGGCCCAGCTCAATGG--------------
------------------------------CTGGGTCGAGCTTTGGGTCCGCGAGGGGGGGTAAGT--------------
--------------------------------------------------------------------------------
------------------------------------------------------
>lcl|LT981265.1_rrna_18 [locus_tag=NCAV_RRNA1] [product=ribosomal RNA 16S ribosomal RNA]
[location=1002313..1003780] [gbkey=rRNA] Nca
--------------------------------------------------------------------------------
--------------------------------------------------------------------------------

--------------------------------------------------------------------------------
--------------------------------------------------------------------------------
--------------------------------------------------------------------------------
--------------------------------------------------------------------------------
--------------------------------------------------------------------------------
--------------------------------------------------------------------------------
-----------------------------------------------------ACTCCGGTTGATCCTGCCGGACCCGAC
TGCTATCGGATTGGACTAAGCCATGCGAGTCAACTCCGCAAGGGGTG---------------------------------
-------GCGCACGGCTCAGTAACA-CGTAGTCAACCTACCCTGGGGACGGGGATAACC---------------------
-----------------------------TCGGGAAA-CTGAGGCTAAACCCCGAT-GGGCC--ATGGC-TCCTGGAATG
GGCTAT-GG---------------------CTG-AAATCTACATGGCCCCAGGATGGGACTGCGGC---CG-ATCAGGT-
--------------------------TGTTGGTGGGGT-AATGGCCCACCAAGCCTATAACCGGTGCGGGCTCTGAGAGG
AGGAGCCCGGAGATGGGCACTGAGACAAGGGCCCAGGTCCTATGGG--GCGCAGCAGGCGCGAAACCTCCGCAATGGGCG
AAAGCCCGAC---GGGG---T-----------CAGTCCGAGTGGTTCCCGCTGAGGGAACC---TTTTGCCAGCT-----
-------------------------------GTAAACAGGCTG------GCGAATAAGGGGTGGGCAAG-TTTGGTGTCA
GCCGCCGCGGTAATACCAGCACCCCGAGTGGTCGGGACGATTATTGGGCCTAAAGCATCCGTAGCCGGCCCAGTAAGTCC
TCCGTTAAATCCAACGGCTCAACCG-TTGGGCT----------------------------------GCGGGGGATACT-
-----------------------------------GCTGGGCTAGGGGACGGGAGAGGCGAGGGGTATTCCCGGGGTAGG
GGTAAAATCCACTGATCCCGGGAGGACCACCAG-----------------------------------------------
-------------------------------------------------------------------TGGCGAAGGCGCC
TCGCTAGAACGTGCCC-------------------------------------------------------GACGGTGAG
GGATGAAAGCTGGGGGAGCGAACCGGATTAGATACCCGGGTAGTCCCAGCCGTAAA------------------------
--------------------------------------------------------------------------------
--------CGATGTAGGCTCGGTGTTGGGCTGC-CTCAGTGGCAGCCCAGT---GCCGCAGGGAAGCCGTTAAGCCTACC
GCCTGGGGAGTACGGCCGCAAGGCTGAAACTTAAAGGAATTGGCGGGGGAGCACCACA-AGGGGTGAAGCC-TGCGGTTT
AATTGGAGTCAACGCCGGGAACCT-TACCAGGGGTGACAGCAGGATGAAGGTCA--------------------------
---------------------AGCTAAAGACTTTACCAGACA----AGCTGAGAGGAGGTGCATGGCCGTCGCCAGCTCG
TGCCG-TGAGGTG--TCCTGTTAAGTCAGGTAAC----------------------------------------------
--------------------------------------------------------------------------------
----------G------AGCGAGACCCTCGCATCCAGTTGCTAC---CATCA--CTCTTTGGAGCGGTGGGGCTAACTGG
ATGGACTGC----CGCCGATA-AGGCGGAGGAAGGAGGGGGCTACGGCAGGTCA-GTATGC-CCCGAAACCCCTG-GGCC
ACACGCGGGCTGCAATGGTAGGGACAATGGGTTC-CGACCCCGAAAGGGGGAG---------------------------
---------------------------------------------------GCAA-TCCCCAAACCCTACCCCAGTTATG
ATTGAGGGCTGCAACCCGCCCTCATGAAT--CTGGAATCCCTAGTAATCGCGTGTCACCATCG-CGCGGTGAATACGTCC
CTGCTCCTTGCACACACCGCCCGTCGCTTCACCCGAGGTGGGCTTGCGTGAGATGGTGCC-TCT----------------
---------------------------TGGTATCATCGAACGCAAGCTCGCTGAGGGGGGAGAAGTCGTAACAAGGTGGC
CGTAGGGGAACCTGCGGCCGGATCACCTCCT-------------------------------------------------
------------------------------------------------------
>lcl|CP003842.1_rrna_25 [locus_tag=NKOR_05661] [product=16S ribosomal RNA] [location=945953..947418]
[gbkey=rRNA] Nko
--------------------------------------------------------------------------------
--------------------------------------------------------------------------------
--------------------------------------------------------------------------------
--------------------------------------------------------------------------------
--------------------------------------------------------------------------------
--------------------------------------------------------------------------------
--------------------------------------------------------------------------------
--------------------------------------------------------------------------------
-------------------------------------------------------TCCGGTTGATCCTGCCGGACCTGAC
TGCTATCGGATTGATACTAAGCCATGCGAGTCATTGTAGCAATACAAG--------------------------------
-------GCAGACGGCTCAGTAACG-CGTAGTCAACCTAACCTATGGACGGGAATAACC---------------------
-----------------------------TCGGGAAA-CTGAGAATAATGCCCGAT-AGAAC--ACTAT-ATCTGGAATG
GTTTGT-GT---------------------TCCAAATGATTTATCGCCGTAGGATGGGACTGCGGC---CT-ATCAGTT-
--------------------------TGTTGGTGAGGT-AATGGCCCACCAAGACTATTACAGGTACGGGCTCTGAGAGG
AGTAGCCCGGAGATGGGTACTGAGACACGGACCCAGGCCCTATGGG--GCGCAGCAGGCGAGAAAACTTTGCAATGTGCG
AAAGCACGAC---AAGG---T-----------TAATCCGAGTGGTTTCTGCTAAAGGAACC---TTTTGTCAGTC-----
-------------------------------CTAGAAACACTG------ACGAATAAGGGGTGGGCAAGTTCTGGTGTCA
GCCGCCGCGGTAAAACCAGCACCTCAAGTGGTCAGGATGATTATTGGGCCTAAAGCATCCGTAGCCGGCTCTGTAAGTTT
TCGGTTAAATCTGTACGCTCAACGT-ACAGGCT----------------------------------GCCGGGAATACT-
-----------------------------------GCAGAGCTAGGGAGTGGGAGAGGTAGACGGTACTCGGTAGGAAGG
GGTAAAATCCTTTGATCTATTGATGACCACCTG-----------------------------------------------
-------------------------------------------------------------------TGGCGAAGGCGGT
CTACCAGAACACGTCC-------------------------------------------------------GACGGTGAG
GGATGAAAGCTGGGGGAGCAAACCGGATTAGATACCCGGGTAGTCCCAGCTGTAAA------------------------
--------------------------------------------------------------------------------
--------CTATGCAAACTCAGTGATGCATTGG-CTTGTGGCCAATGCAGT---GCTGCAGGGAAGCCGTTAAGTTTGCC
GCCTGGGAAGTACGTACGCAAGTATGAAACTTAAAGGAATTGGCGGGGGAGCACCACA-AGGGGTGAAGCC-TGCGGTTC
AATTGGAGTCAACGCCAGAAATCT-TACCCGGAGAGACAGCAGAATGAAGGTCA--------------------------
---------------------AGCTGAAGACTTTACCAGACA----AGCTGAGAGGTGGTGCATGGCCGTCGCCAGCTCG
TGCCG-TGAGATG--TCCTGTTAAGTCAGGTAAC----------------------------------------------
--------------------------------------------------------------------------------
----------G------AGCGAGATCCCTGCCTCTAGTTGCCAC---CATTA--CTCTCAGGAGTAGTGGGGCGAATTAG
CGGGACCGC----CGCAGTTA-ATGCGGAGGAAGGAAGGGGCCACGGCAGGTCA-GTATGC-CCCGAAACTCTGG-GGCC
ACACGCGGGCTGCAATGGTAACGACAATGGGTTT-CAAATCCGAAAGGATGAG---------------------------
---------------------------------------------------GTAA-TCCTCAAACGTTACCACAGTTATG
ACTGAGGGCTGCAACTCGCCCTCACGAAT--ATGGAATCCCTAGTAACTGCGTGTCATTATCG-CGCGGTGAATACGTCC
CTGCTCCTTGCACACACCGCCCGTCGTTTCATTGAAGTGAGCTTTTAGCGAGGTGACGTCTGAT----------------
---------------------------TGGCGTTATCGAACTTGAGGTTCGTGACAAGGGAAAAGTCGTAACAAGGTGAC
CGTAGGGGAACCTGCGGTCGGATCACCT----------------------------------------------------
------------------------------------------------------
>lcl|BA000048.1_rrna_46 [locus_tag=CSUB_R02] [product=16S ribosomal RNA] [location=complement(1662154..1663645)]
[gbkey=rRNA] Csu
--------------------------------------------------------------------------------
--------------------------------------------------------------------------------
--------------------------------------------------------------------------------
--------------------------------------------------------------------------------
--------------------------------------------------------------------------------

--------------------------------------------------------------------------------
--------------------------------------------------------------------------------
--------------------------------------------------------------------------------
-----------------------------------------------------ACTCCGGTTGATCCTGCCGGACCTCAC
TGCTATCGGGGGGAGACTAAGCCATGCAAGTCTAGGCCCATAGCCACCTG-----------------------------T
GGGCCTGGCGTACGGCTCAGTAACA-CGTGGCCAACCTACCCTGAGGAGGCGGATAACACCGG-----------------
--------------------------------GAAAC-TGGTGCATAAACCGCCAT-AGGTG--AGAGG-ATCTGGAACG
ACCTCT-CGCCGAAAGATGTACACGGGGATGC---CTGTGTACATGCCTCAGGATGGGGCTGCGGC---CC-ATCAGGTA
GT--TGGTGGGGT--------------------------AACGGCCCACCAAGCCTATAACGGGTACGGGCCCTGAGAGG
GGGAGCCCGGAGATGGACTCTGAGACAAGAGTCCAGGCCCTACGGG--GCGCAGCAGGCGCGAAACCTCCGCAATGCGCT
CACGCGTGAC---GGGG---T-----------CACCCCGAGTGCC--ACCCTCGTGGTGGC---TTTTGCGCAGT-----
-------------------------------GTAAAAAGCTGC------GCGAATAAGCGGGGGGCAAG-ACGGGTGTCA
GCCGCCGCGGTAATACCCGCTCCGCGAGTGGTGAGGACTATTACTGGGCTTAAAGCGTCCGTAGCCGGCTCGGCGTGCTC
CGCGTTAAATCCAGCAGCCTAACTG-CTGGGCC----------------------------------GCGCGGATGACT-
-----------------------------------GCCGGGCTTGGGGGTGGGAGAGGGCAGCGGTATTCCCGGGGTAGG
GGTAAAATCCACTGATCCCGGGAGGACCACCAG-----------------------------------------------
-------------------------------------------------------------------TGGCGAAGGCGGC
TGCCCAGAACACGCCC-------------------------------------------------------GACGGTGAG
GGACGAAAGCTGGGGGAGCGATCGGGATTAGATACCCCGGTAGTCCCAGCCGTAAA------------------------
--------------------------------------------------------------------------------
--------CGATGCGGGCTTGGCGTCGGAGCGG-CTACGAGCCGTTCCGGT---GCAGAAGGGAAACCATTAAGCCCGCC
GCCTGGGGAGTACGGCCGCAAGGCTGAAACTTAAAGGAATTGGCGGGGGAGCACCACA-AGGGGTGGACGT-TGCGGCTT
AATTGGAGTCAACGCCGGAAACCT-TACCGGGGGCGACAGCAGGATGATGGCCA--------------------------
---------------------GGTTAAAGGCCTTGCCGGACG----AGCTGAGAGGAGGTGCATGGCCGTCGCCAGTGCG
TGCCG-TGAGGTG--TCCTGTTAATTCAGGCAAC----------------------------------------------
--------------------------------------------------------------------------------
----------G------GACGAGACCCGCGCCTCCAGTTGCGAC---CGGGC--CGAAAGGCGCC-----GAGCACTCTG
GAGGGACTG----CCCCGGAA-ACGGGGAGGAAGGAGCGGGCTACGGCAGGTCA-GTATGC-CCCGAATCCCCCG-GGCC
GCACGCGACGTTCAATGGTGGGGACAGTGGGCTG-CGACCCCGAAAGGGGAAG---------------------------
---------------------------------------------------CCAA-TCCCTAAACCCCACCTCAGTAGGG
ATCGAGGGTTGCAACTCGCCCTCGTGAAC--GCGGAATCCCTAGTAACCGCCCGTTACCACCG-GGCGGTGAATACGTCC
CCGCTCCTTGCACACACTGCCCGTCGCTGCACCCGAGCTGGAACTGGGCGAGGCCCGTCTCGT-----------------
---------------------------TGGCGGGTTCGAGCCCGGTTCTGGTGAGGGGGCAGAAGTCGTAACAAGGTGGC
CGCACGGGAACGTGTGGCCGGATCACCTCCT-------------------------------------------------
------------------------------------------------------
>lcl|NEXG01000006.1_rrna_39 [locus_tag=B9Q05_04580] [product=16S ribosomal RNA]
[location=complement(41335..42824)] [gbkey=rRNA] Mar
--------------------------------------------------------------------------------
--------------------------------------------------------------------------------
--------------------------------------------------------------------------------
--------------------------------------------------------------------------------
--------------------------------------------------------------------------------
--------------------------------------------------------------------------------
--------------------------------------------------------------------------------
--------------------------------------------------------------------------------
-----------------------------------------------------AACCCGGTTGATCCTGCCGGACCCGAC
TGCTATCGGGATAATGCTAAGCCATGGGAGTCGAACGCCTCCAGCCACGTG----------------------------G
GGGCGTGGCGTACGGCTCAGTAGCA--------CACGGCTAA----------TCTACCCTTGGGAGGGG-----------
--------------------TATAACCCCG--GGAAA-CTGGGGCTAATATCCCAT-AGACG--AGCAACC-CTGGAATG
GGTGCT-CATCGAAAGGGCGT-GGGAGCATGCT----TCCGCGTCGCCCAAGGATGAGGCTGTGTT---CC-ATCAGGTA
GT--TGGCGAGGTAACGGCTCGCC-----AAGCCGA-----TAACGGATGCGGGC----------------AGTGAGAGC
TGGAGCCCGGAGATGGGCACTGAGACAAGGGCCCAGGCCCTACGGG--GCGCACCAGGCGAGAAACCTTAGCAATGCGGG
AAACCGTGAC---TGGG---C-----------TATCCCGAGTGCCAGCCGCTAAGGCTGGC---TTTTCCCCAGA-----
-------------------------------GTAAAAACCTGG------GGGAATAAGCGGGGGGCAAG-ACTGGTGTCA
GCCGCCGCGGTAATACCAGCTCCGCGAGTGGTCGGGACGATTATTGAGCCTAAAGCGCTCGTAGCCGGTCCGGTAAGTCC
TGTGCTAAATCCAACGGCTCAACCG-TTGGGCC----------------------------------GCATGGGATACT-
-----------------------------------GCCGGGCTAGGGAGCGGAAGAGGCCAGGGGTACTCCCGGGGTAGG
GGCGAAATCCTATAATCCCGGGAGGACGACCAG-----------------------------------------------
-------------------------------------------------------------------TGGCGAAGGCGCC
TGGCTAGGACGCGTCC-------------------------------------------------------GACGGTGAG
GGGCGAAAGCTGGGGGAGCGAACCGGATTAGATACCCGGGTAGTCCCAGCTGTAAA------------------------
--------------------------------------------------------------------------------
--------CGATGCGGATTGGGTGCCGGTACGG-CTTCGGGCCGTATCGGT---GCCTCAGGGAAGCCGTTAAATCCGCC
GCTTGGGGAGTACGGTCGCAAGACTGAAACTTAAAGGAATTGGCGGGGGAGCACCACA-AGGGGTGAAGCC-TGCGGCTC
AATTGGAGTCAGCGCCGGGAACTT-TACCGGGGGCGACAGCAGTATGAGGGCCA--------------------------
---------------------AGCTGATGACTTTGCCCGACG----CGCTGAGAGGTGGCGCATGGCCGTCGCCAGCTCG
TGCCG-TGAGGTG--TCTGGTTAAGTCCAGGAAC----------------------------------------------
--------------------------------------------------------------------------------
----------G------AGCGAGACCCTCACCCCTAGTTGCCAC---CGCGC--AAGCG-----------GGCACACTAG
GGGGACCGC----CACCGCTA-AGGTGGAGGAGGGAGGGGGCAACGGCAGGTCA-GCATGC-CCCGAATCCCCCG-GGCC
GCACGCGGGCTGCAATGGTGGAGACAATGGGTTC-CGAACCCGAAAGGGGGAG---------------------------
---------------------------------------------------GTAATCCTTGAAACTCCACCGTAGTTGGG
ATCGAGGGCTGCAACCCGCCCTCGTGAAC--CTGGAATCCCCAGTAATCGCGTGTCATCATCG-CGCGGTGAACACGTCC
CCGCTCCTTGCACACACCGCCCGTCGCTCCACCCGAGCTGGTTTCAGATGAGGCGGGATCGGTTCG-C------------
---------------------------CGGTTCCGTCGAATCTGAGACCGGTGAGGGGGGAGAAGTCGTAACAAGGTGGC
CGTAGGGGAACCTGCGGCCGGATCACCTCCT-------------------------------------------------
------------------------------------------------------
>lcl|AE009441.1_rrna_37 [locus_tag=PAErR16S] [product=small subunit 16S ribosomal RNA]
[location=1089640..1091850] [gbkey=rRNA] Pae
-AAACCGGTTGATCCTGCCGGACCCGACCGCTATCGGGGTAGGGCTAAGCCATGCGAGTCGCGCGCCCGGGGGCGCCCGG
GAGCGGCGCACGGCTCAGTAACACGTGCCTAAC--------CTAACCTCGGGAGGGGGACACCCCCGGGAAACTGGGGTC
AATCCCCCATAGGGGAAGGGCGCTGGAAGGCCCCTTCCCCGAAAGGGATGGCGGCCGATCCGCCGCCGTCCGCCCGAGGG
TGGGGGCACGGCCCATCAGGTAGTTGGCGGGTTAAAGGCCCGCCAAGCCGAAGACGGGTAGGGGCGGTGAGAGCCGCGAG
CCCCGAGATCGGCACTGAGACAAGGGCCGAGGCCCTACGGGGTGCAGCAGGCGCGAATACTCCGCAAATGGGGGAAGGCC
GGCTCCTCATGACGGAGGAATTTAACCTAAATGAAGTCAAATATTCGAAATTTGACAAGAAACGTAACATCAGAGTGCCG
GACAAGCCAACAGAACTTCTAGCAGAGGAGATAGCAATACACTTAGGAGACGGCTACCTTTTTTACGA---CGAAGAGGA
TAATAGCTATAGATACGGCGTTGGGCTTAATCCAAAAACCGAGGTTGAATACGCATACGCCGTAGCGGAGTTAATA-GGG

```
GAAGTATATGGTTATAA-----------A-CCTCCGGTAAGAAAGGCGCGTATTGAGATAATGTCCCTAGCGATAGGGAC
GTTTAAACACAAAGTGCTCGGTTTTCCTATAGGCACCAGGACGGGAAACG-----------------------------A
GAAGA--TGCCTCGGATAGGTTGGA-TATTACAAGATGAGAAATTCGCCACGGCGATCATAAGAGGGCT-----------
--------------------TTTAGATACCGAAGGAT-CGGTG-AAAAAGATATCT-AGGAC--TTTGGGGATTGTT-GT
GAAGCA-GAGAAACAAGGATATCGCTATGTTCTACGCCCAGT-GTATTTCTCTATTGGGTTTCACT---CCACGAATATA
CCAATGGAACGAAAAGAACAAGCCTATCTACGCCGTCG-TGGTACTGGGCAAGGATGTGGTAGAAGACTTCCTCCGGACT
ATAAAACCCAGAAATCCCTCTAAGACC-CCCTCTTTTTAGTCTTCC--TCTCACGCCGGCCTCCTCCAGCGGAATGCGGG
CAACCGCGAC---GGGG---C-----------TACCCCGAGTGCCGGGCGAAGAGCCCGGC---TTTTGCCCGGT-----
-------------------------------CTAAAACGCCGG------GCGAATAAGCGGGGGGCAAG-TCTGGTGTCA
GCCGCCGCGGTAATACCAGCTCCGCGAGTGGTCGGGGTGTTTACTGGGCTTAAAGCGCCCGTAGCCGGCCCGGCAAGTCG
CTCCTGAAATCCCCGGGCTCAACCC-GGGGGCG----------------------------------GGGGGCGATACT-
-----------------------------------GCCGGGCTAGGGGGCGGGAGAGGCCGCCGGTACTCCGGGGGTAGG
GGCGAAATCCGTTAATCCCCGGAGGACCACCAG-----------------------------------------------
-------------------------------------------------------------------TGGCGAAAGCGGG
CGGCCAGAACGCGCCC-------------------------------------------------------GACGGTGAG
GGGCGAAAGCCGGGGGAGCAAAGGGGATTAGATACCCCTGTAGTCCCGGCCGTAAA------------------------
--------------------------------------------------------------------------------
--------CGATGCGGGCTAGCTGTCGGTCGGG-CTTAGGGCCCGGCCGGT---GGCGAAGGGAAACCGTTAAGCCCGCC
GCCTGGGGAGTACGGCCGCAAGGCTGAAACTTAAAGGAATTGGCGGGGGGGCACCACA-AGGGGTGAAGCT-TGCGGCTT
AATTGGAGTCAACGCCGGAAACCT-CACCCGGGGCGACAGCAGGATGAAGGCCA--------------------------
---------------------GGCTAACGACCTTGCCGGACG----AGCTGAGAGGAGGTGCATGGCCGTCGTCAGCTCG
TGCCG-TGAGGTG--TCCGGTTAAGTCCGGCAAC----------------------------------------------
--------------------------------------------------------------------------------
----------G------AGCGAGACCCCCGCCCCTAGTTGCTAC---CCCGT--CCTTCGGG---ACGGGGGGCACACTA
GGGGGACTG----CCGGCGTA-AGCCGGAGGAAGGAGGGGGCCACGGCAGGTCA-GTATGC-CCCGAAACCCCGG-GGCT
GCACGCGAGCTGCAATGGCGGGGACAGCGGGATC-CGACCCCGAAAGGGGGAG---------------------------
---------------------------------------------------GCAATCCCGTAAACCCCGCCCCAGTAGGG
ATCGAGGGTTGCAACTCGCCCTCGTGAAC--GTGGAATCCCTAGTAACCGCGTGTCACCAACG-CGCGGTGAATACGTCC
CTGCCCCTTGCACACACCGCCCGTCGCACCACCCGAGGGAGCCCTCTGCGAGGCCCCTCGCCGAAGAG------------
---------------------------GTGGGGGGACGAGCAGGGGGCTCCCAAGGGGGGTGAAGTCGTAACAAGGTAGC
CGTAGGGGAACCTGCGGCTGGATCACCTC---------------------------------------------------
------------------------------------------------------
>lcl|LT549890.1_rrna_47 [locus_tag=SSOP1_r08] [product=16S ribosomal RNA] [location=891113..892713] [gbkey=rRNA]
Sso
--------------------------------------------------------------------------------
--------------------------------------------------------------------------------
--------------------------------------------------------------------------------
--------------------------------------------------------------------------------
--------------------------------------------------------------------------------
--------------------------------------------------------------------------------
--------------------------------------------------------------------------------
--------------------------------------------------------------------------------
----------------------------------------------------AAATCCGGTTGATCCTGCCGGACCCGAC
CGCTATCGGGGTGGGGCTAAGCCATGGGAGTCGTACGCTCCCGGGCAAGGG----------------------------A
GCGTG--GCGGACGGCTGAGTAACA--------CGTGGCTAACCT----------ACCCTGAGGAGGGA-----------
--------------------GATAACCCCGGGA--AA-CTGGGGATAATCTCCCAT-AGGCG--AGGAG-TCCTGGAACG
GTTCCT-CGCTGAAAGGCTCA-TGGGCTATTCCCCGCTCATGAGCGCCTCAGGATGGGGCTGCGGC---CC-ATCAGGTA
GT--TGGGGGGGTAAGGGCCCCCC-----AAGCCTA-----TAACGGGTAGGGGCCGTGAGAG---------------CG
GGAGCCCCCAGTTG-GGCACTGAGACAAGGGCCCAGGCCCTACGGG--GCGCACCAGGCGCGAAACGTCCCCAATGCGCG
GAAGCGTGAG---GGCG---C-----------CACCCCGAGTGC--TCCCGTAA-GGGAGC---TTTTCCCCGCT-----
-------------------------------CTACAAAGGCGG------GGGAATAAGCGGGGGGCAAG-TCTGGTGTCA
GCCGCCGCGGTAATACCAGCCCCGCGAGTGGTCGGGACTCTTACTGGGCCTAAAGCGCCCGTAGCCGGCCCGACAAGTCA
CTCCTTAAAGACCCCGGCTCAACCG-GGGGAAT----------------------------------GGGGGTGATACT-
-----------------------------------GTCGGGCTAGGGGGCGGGAGAGGCCAGCGGTACTCCCGGAGTAGG
GGCGAAATCCTCAGATCTCGGGAGGACCACCAG-----------------------------------------------
-------------------------------------------------------------------TGGCGAAAGCGGC
TGGCTAGAACGCGCCC-------------------------------------------------------GACGGTGAG
GGGCGAAAGCCGGGGCAGCAAAAGGGATTAGATACCCCTGTAGTCCCGGCTGTAAA------------------------
--------------------------------------------------------------------------------
--------CAATGCAGGCTAGGTGTCACATGGG-CTTAGAGCCCATGTGGT---GCCGCAGGGAAGCCGTTAAGCCTGCC
GCCTGGGGAGTACGGTCGCAAGACTGAAACTTAAAGGAATTGGCGGGGGAGCACCACA-AGGGGTGGAACC-TGCGGCTC
AATTGGAGTCAACGCCTGGAATCTTAC-TAGGGGAGACCGCAGGATGACGGCCA--------------------------
---------------------GGCTAACGACCTTGCCTGACT----CGCGGAGAGGAGGTGCATGGCCGTCGCCAGCTCG
TGTTG-TGAAATG--TCCTGTTAAGTCAGGCAAC----------------------------------------------
--------------------------------------------------------------------------------
----------G------AGCGAGACCCCCACCACTAGTTGGTAT---CCTGG--TCTCCGGG---CCGGGACCACACTAG
TGGGACTGC----CGGC-GTA-AGCCGGAGGAGGGAGGGGGCCACGGCAGGTCA-GCATGC-CCCGAAACCCCTG-GGCC
GCACGCGGGTTACAATGGCAGGGACAGCGGGATTCCGACCCCGAAAGGGGAAG---------------------------
---------------------------------------------------GTAATCCCTTAAACCCTGCCGCAGTTGGG
ATCGAGGGCTGAAACTCGCCCTCGTGAAC--GAGGAATCCCTAGTAACCGCAGATCAACAATC-TGCGGTGAATACGTCC
CTGCTCCTTGCACACACCGCCCGTCGCTCCACCCGAGTAGGAGAGGGGTGAGGCCCCTTGCCTTTG-G------------
---------------------------GTGGGGGTCGAGCTTCTCTCCTGCAAGGGGGGAGAAGTCGTAACAAGGTAGC
CGTAGGGGAACCTGCGGCTGGATCACCTCATAGTTACAAACTCCCCCGTCAATTTCCATAGGGGTAGGAAGTCCATAAAA
CCCTTGGTTCCCTTTCACAGATGCGATTCTCCTCTTGGCCGGAGGAGAATTAAG
>lcl|CP010515.1_rrna_6 [locus_tag=SE86_01610] [product=16S ribosomal RNA] [location=complement(329921..331426)]
[gbkey=rRNA] Aci
--------------------------------------------------------------------------------
--------------------------------------------------------------------------------
--------------------------------------------------------------------------------
--------------------------------------------------------------------------------
--------------------------------------------------------------------------------
--------------------------------------------------------------------------------
--------------------------------------------------------------------------------
--------------------------------------------------------------------------------
--------------------------------------------TGCGGCTGAACTCCGGTTGATCCTGCCGGACCCGAC
TGCTATCGGGGTGAGGCTAAGCCATGGGAGTCGCGCGCCCAGCCGCCGCTG----------------------------G
GCGCG--GCGCACGGCTGAGTAACA--------TGTAGCTAACCT----------ACCCTCGGGAGGGG-----------
```

--------------------GACAACCCCG--GGAAA-CTGGGGCTAATCCCCCAT-AGGCA---AGGGGGCCTGGAAGG
GTCTCT-TGCCGAAAGGGACCTCAGGGGGTTATCGCCTGGGGTCCGCCCGAGGATGGGGCTGCACC---CCATCATGGTA
GT--TGGCGGGGTAATGACCCGCC-----AAGCCGA-----CGACGGGTAGGGGCCGTGGGAG---------------CG
GGAGCCCCCAGATG-GGCCCTGAGACAAGGGCCCAGGCCCTACGGG--GCGCACCAGGCGCGAAACCTCCGCAATGCGGG
AAACCGTGAC---GGGG---T-----------CACCCCGAGTGC--CCCCGT-AAGGGGGC---TTTTCCCCGCT-----
-------------------------------GCAAGGAGGCGG------GGGAATAAGCGGGGGGCAAG-TCTGGTGTCA
GCCGCCGCGGTAATACCAGCCCCGCGAGTGGTCGGGACGTCTACTGGGCCTAAAGCGCCCGTAGCCGGCCCCGTAAGTCC
CTCCTGAAAGCCCTGGGCTCAACCC-AGGGAGT----------------------------------GGGGGGGATACT-
-----------------------------------GCGGGGCTAGGGGGCGGGAAAGGCCGGGGGTACCCCAGGGGTAGG
GGCGAAATCCGATAATCCCTGGGGGACCACCAG-----------------------------------------------
-------------------------------------------------------------------TGGCGAAAGCGCC
CGGCTGGAACGCGCCC-------------------------------------------------------GACGGTGAG
GGGCGAAAGCCGGGGGAGCGAACCGGATTAGATACCCGGGTAGTCCCGGCTGTAAA------------------------
--------------------------------------------------------------------------------
--------CGATGCGGGCTAGGTGTCGGGCGGGCGTTAGAGCCCGCCCGGT---GCCGCAGGGAAGCCGTTAAGCCCGCC
GCCTGGGGAGTACGGCCGCAAGGCTGAAACTTTAAGGAATTGGCGGGGGGGCA-CACA-AGGGGTGGAGCC-TGCGGCTC
AATTGGAGTCAACGCCGGGAACCT-CACCGGGGGCGACAGCAGGATGACGGCCA--------------------------
---------------------GGCTAATGACCTTGCCTGACG----CGCTGAGGGGAGGTGCATGGCCGTCGCCAGCTCG
TGCTG-TGAAGTG--TCCTGTTAAGTCAGGCAAC----------------------------------------------
--------------------------------------------------------------------------------
----------G------AGCGAGACCCCCGCCCCTAGTTGCGAC---CCTGC--GGGCGACC---GTGGGGGCACACTAG
GGGGACTGC----CGCCGCTA-AGGCGGAGGAAGGAGGGGGCCACGGCAGGTCA-GCATGC-CCCTAAACCCCCG-GGCT
GCACGCGGGCTACAATGGCGGGGACAGCGGGATC-CGACCCCGAAAGGGGGAG---------------------------
---------------------------------------------------GCAATCCCTGAAACCCCGCCGTAGTCGGG
ATTGGGGGCTGAAACTCGCCCCCATGAAC--CTGGAATCCCTAGTAACCGCGCGTCAACATCG-CGCGGTGAATACGTCC
CTGCCCCTTGTACACACTGCCCGTCGCTCCACCCGAGAGGGGGAGGGGTGAGGCCCTCTCCCTC---G------------
---------------------------GGAGGGGGTCGAACCTCTCCCCCTCGAGGGGGGAGAAGTCGTAACAAGGTAGC
CGTAGGGGAACCTGCGGCTGGATCACCTCTC-------------------------------------------------
------------------------------------------------------
>lcl|BA000002.3_rrna_42 [locus_tag=APE_rRNA05] [product=16S ribosomal RNA]
[location=complement(join(1218714..1219336,1220036..1220913))] [gbkey=rRNA] Ape
--------------------------------------------------------------------------------
--------------------------------------------------------------------------------
--------------------------------------------------------------------------------
--------------------------------------------------------------------------------
--------------------------------------------------------------------------------
--------------------------------------------------------------------------------
--------------------------------------------------------------------------------
--------------------------------------------------------------------------------
-----------------------------------------------------ACTCCGGTTGATCCTGCCGGACCCGAC
CGCTATCGGGGTAGGGCTAAGCCATGGGAGTCGAACGCCCCGCCGCTGCGG----------------------------G
GCGTG--GCGGACGGCTGAGTAACA--------CGTGGCTAACCT----------ACCCTCGGGACGGG-----------
--------------------GATAACCCCTCGGGAAA-CTGGGGCTAATCCCCGAT-AGGCG--GGGGGTGCCTGGAAGG
GTCCTC-CGCCGAAAGGGGCCGCGGGGGGTTATCGCCCGCGGTCCGCCCGAGGATGGGGCTGCGGC---CCATCACGGTA
GT--TGGCGGGGTAATGGCCCGCC-----AAGCCTA-----CGACGGGTAGGGGCCGTGAGAG---------------CG
GGAGCCCCCAGATG-GGCACTGAGACAAGGGCCCAGGCCCTACGGG--GCGCACCAGGCGCGAAAACTCCGCAATGCGGG
CAACCGTGAC---GGGG---C-----------TACCCCGAGTGC--CCCCCTTGGGGGGGC---TTTTCCCCGCT-----
-------------------------------GTAAGTAGGCGG------GGGAATAAGCGGGGGGCAAG-ACTGGTGTCA
GCCGCCGCGGTAATACCAGCCCCGCGAGCGGTCGGGGTGCTTACTGGGCCTAAAGCGCCCGTAGCCGGCTCGACAAGTCC
CCGCCTAAAGCCCCGGGCTCAACCC-GGGGAGT----------------------------------GGCGGGGATACT-
-----------------------------------GTCGGGCTAGGGGGCGGGAGAGGCCGAGGGTACTCCCGGGGTAGG
GGCGAAATCCATTAATCCCGGGAGGACCACCAG-----------------------------------------------
-------------------------------------------------------------------TGGCGAAGGCGCT
CGGCTGGAACGCGCCC-------------------------------------------------------GACGGTGAG
GGGCGAAAGCCGGGGGAGCGAACCGGATTAGATACCCGGGTAGTCCCGGCTGTAAA------------------------
--------------------------------------------------------------------------------
--------CGATGCGGGCTAGGTGTTGGGCGGGCGTGCGAGCCCGCCCAGT---GCCGCAGGGAAGCCGTTAAGCCCGCC
GCCTGGGGAGTACGGCCGCAAGGCTGAAACTTAAAGGAATTGGCGGGGGAGCACCACA-AGGGGTGGAGCC-TGCGGCTT
AATTGGAGTCAACGCCGGGAACCTCCACCGGGGGCGACAGCAGGATGACGGCCA--------------------------
---------------------GGCTGAAGGCCTTGCCCGACG----CGCTGAGGGGAGATGCATGGCCGTCGCCAGCTCG
TGCCG-TGAGGTG--TCCTGTTAAGTCAGGCAAC----------------------------------------------
--------------------------------------------------------------------------------
----------G------AGCGAGACCCCCGCCCCTAGTTGCCAT---CCCCG--GGGCGACC---CGGGGGGCACACTAG
GGGGACTGC----CGCCGCTA-AGGCGGAGGAAGGAGGGGGCAACGGCAGGTCA-GCATGC-CCCTAAACCCCCG-GGCT
GCACGCGGGCTACAATGGCGGGGACAGCGGGTGC-CGACCCCGAAAGGGGGAG---------------------------
---------------------------------------------------GCAATCCCTCAAACCCCGCCTCAGTTGGG
ATCGAGGGCTGCAACTCGCCCTCGTGAAC--GTGGAATCCCTAGTAACCGCGCGTCATCATCG-CGCGGTGAATACGTCC
CTGCTCCTTGCACACACTGCCCGTCGCTCCACCCGAGGGGGGAGGGGGTGAGGCCCTCTCCGCC---G------------
---------------------------GGAGGGGGTCGAACCCCCTCCCCCCGAGGGGGGAGAAGTCGTAACAAGGTAGC
CGTAGGGGAACCTGCGGCTGGATCACCTC---------------------------------------------------
------------------------------------------------------
>lcl|RXGA01000004.1_rrna_36 [locus_tag=Metus_R0038] [product=16S ribosomal RNA] [location=3780..5281]
[gbkey=rRNA] Met
--------------------------------------------------------------------------------
--------------------------------------------------------------------------------
--------------------------------------------------------------------------------
--------------------------------------------------------------------------------
--------------------------------------------------------------------------------
--------------------------------------------------------------------------------
--------------------------------------------------------------------------------
--------------------------------------------------------------------------------
--------------------------------------------------TTAATCCCGGTTGATCCTGCCGGACCCGAC
TGCTATTGGGGTGAGGCTAAGCCATGCGAGTTAGCTCTCCTAGCCACGATG----------------------------G
-AGGGCGGCGTACGGCTCAGTAACA-TGTGGTCAATCTACCCTCAGGACGGAGATAACCTCGGGAAACT-----------
----------------------------------------GAGGCTAATATCCGAT-AGATA--ATGGA-TCCTGGAATG
GTTCAT-TATTCAAACGGCACATT-GCTCATGC--GCAGTGTGCCGCCTGAGGATGAGACTGCACC---CT-ATCAGGTA
GT--TGGTGGGGTAACGGC--------------------------CCACCAAGCCTATAACGGGTACGGGCCCTGAGAGG

GGGAGCCCGGAGATGGGCACTGAGACAAGGGCCCAGGCCCTACGGG--GCGCAGCAGGCGCGAAACCTCTGCAATGCGCG
AAAGCGCGAC---AGGG---C-----------CACCCCGAGTGCTAGCCGCTGAGGTTAGC---TTTTCCTTAGT-----
-------------------------------GTAGTAAGCTAA------GGGAATAAGCGGGGGGCAAG-CCTGGTGTCA
GCCGCCGCGGTAATACCAGCTCCGCGAGTGGTCGGGACGATTATTGGGCCTAAAGCGTCCGTAGCCGGCCTATCAAGTCT
CTGGTTAAACCTCAAGGCTCAACCT-TGAGATT----------------------------------GCTGGAGATACT-
------------------------------------GTTAGGCTAGGGGGCGGGAGAGGTTGAGGGTACTTCGCGGTAGG
GGCGAAATCCTATAATCCGCGAAGGACCACCAG-----------------------------------------------
-------------------------------------------------------------------TGGCGAAGGCGCT
CAACTGGAACGCGCCC-------------------------------------------------------GACGGTGAG
GGACGAAAGCCAGGGGAGCAAAGGGGATTAGATACCCCCGTAGTCCTGGCTGTAAA------------------------
--------------------------------------------------------------------------------
--------CGATGCGGGCTAGGTGTTGGGTTGG-CTTCGTGCCAACCCGGT---GCCGCAGTGAAGACGATAAGCCCGCC
GCCTGGGAAGTACGGCTGCAAGGCTGAAACTTAAAGGAATTGGCGGGGGAGCACCACA-AGGGGTGAAGCC-TGCGGTTT
AATTGGAGTCAACGCCGGGAATCT-TACCGGGAGCGACAGCAAGATGAAGGCCA--------------------------
---------------------AGCTGACGACTTTGCTAGATG----GGCTGAGAGGAGGTGCATGGCCGTCGCCAGTTCG
TGCCG-TGAGGTG--TCCGGTTAAGTCCGGCAAC----------------------------------------------
--------------------------------------------------------------------------------
----------G------AACGAGACCCCCATCCTCTGTTGCTAG---CCTGC--TTCTACGG-AGGTAGGCGCACACGGA
GGAGACTGC----CAGTGTTA-AACTGGAGGAGGGAGGGGGCTACGGCAGGTCA-GCATGC-CCCGAATCTCCCG-GGCC
ACACGCGGGCTGCAATGGTGAGGACAGCGGGTTC-CGACCCCGAAAGGGGAAG---------------------------
---------------------------------------------------GTAATCCCTTAAACCTCGCCGTAGTTGGG
ATTGAGGGCTGTAACCCGCCCTCATGAAC--ATGGAATCCCTAGTAACCGCATGTCACCATCG-TGCGGTGAATACGTCC
CTGCTCCTTGCACACACCGCCCGTCGCTCCACCCGAGTCAAGCTTGGGCGAGGCCCAGACCTGT----------------
---------------------------TGTTTGGGTCGAACCCATGTTTGGCAAGGGGGGAGAAGTCGTAACAAGGTGGC
CGCAGGGGAACCTGCGGCCGGATCACCTCCTA------------------------------------------------
------------------------------------------------------
>lcl|JYIM01000321.1_rrna_16 [locus_tag=Lokiarch_33830] [product=16S ribosomal RNA] [location=1..1179]
[gbkey=rRNA] Lok
--------------------------------------------------------------------------------
--------------------------------------------------------------------------------
--------------------------------------------------------------------------------
--------------------------------------------------------------------------------
--------------------------------------------------------------------------------
--------------------------------------------------------------------------------
--------------------------------------------------------------------------------
--------------------------------------------------------------------------------
--------------------------------------------------------------------------------
--------------------------------------------------------------------------------
--------------------------------------------------------------------------------
--------------------------------------------------------------------------------
--------------------------------------------------------------------------------
--------------------------------------------------------------------------------
---------GAGATGGGTACTGAGACAACGACCCAGGCCTTACGAG--GCGCAGCAGGCGCGAAACCTCCGCAATACACG
AAAGTGTGAC---GGGG---T-----------TACCCAAAGTGTTCAATTATGAACTGTGG---TA----GGTGA-----
-------------------------------GTAATGTTCCCT------ACTAGAAAGGAGAGGGCAAG-GCTGGTGCCA
GCCGCCGCGGTAAAACCAGCTCTTCAAGTGGTCGGGATAATTATTGGGCTTAAAGTGTCCGTAGCCGGTTTAGTAAGTTC
CTGGTAAAATCGGGTAGCTTAACTA-TCTGTAT----------------------------------GCTAGGAATACT-
-----------------------------------GCTATACTAGAGGACGGGAGAGGTCTGAGGTACTACAGGGGTAGG
GGTGAAATCTTATAATCCTTGTAGGACCACCAG-----------------------------------------------
-------------------------------------------------------------------TGGCGAAGGCGTC
AGACTGGAACGTGCCT-------------------------------------------------------GACGGTGAG
GGACGAAAGCCAGGGGAGCGAACCGGATTAGATACCCGGGTAGTCCTGGCCGTAAA------------------------
--------------------------------------------------------------------------------
--------CGATGCATACTAGGTGATGGCATGG-CCATGAGCCATGTCAGT---GCCGTAGGGAAACCGTTAAGTGTGCC
GCCTGGGAAGTACGGTCGCAAGGCTAAAACTTAAAGGAATTGGCGGGGGAGCACCACA-AGGGGTGAAGCC-TGCGGTTC
AATTGGACTCAACGCCGGGAAACT-TACCAGGGGAGACAGCAGAATGATGGTCA--------------------------
---------------------GGTTGACGACCTTACCTGACA----AGCTGAGAGGAGGTGCATGGCCGTCGCCAGTTCG
TGCTG-TGAGGTA--TCCTGTTAAGTCAGGCAAC----------------------------------------------
--------------------------------------------------------------------------------
----------G------AACGAGATCCGCACCTTTATTTGCCAG---CAAGA--AGTCACGACTTCGTTGGGAACACTAA
AGGGACCGC----CGTCGATA-AGACGGAGGAAGGAGCGGGCAAAGGCAGGTCA-GTATGC-CCCGAAACCCCTG-GGCT
ACACGCGGGCTGCAATGGTATGAACAATGGGCTG-TAACTCCGAAAGGAGAAA---------------------------
---------------------------------------------------CCAATCC-CGAAATCATATCTCAGTGGGA
ATTGTCGGCTGTAACCCGCCGACATGAAC--GTGGAATCCCTAGTAATCGTGTGTCATCATCG-CACGGTGAATACGTCT
CTGCTCCTTGCACACACCGCCCGTCGCTCCATCCGAGTGTGCTAAAAATGAGGTATGGTCAGTC----------------
---------------------------TGGTCGTATCGAATTTCTAGTATGCGAGGGGGGAGAAGTCGTAACAAGGTAGC
CGTAGGGGAACCTGCGGCTGGATCACCTCCT-------------------------------------------------
------------------------------------------------------
>lcl|MDVT01000007.1_rrna_28 [locus_tag=OdinLCB4_10090] [product=16S ribosomal RNA] [location=652378..653923]
[gbkey=rRNA] Odi
--------------------------------------------------------------------------------
--------------------------------------------------------------------------------
--------------------------------------------------------------------------------
--------------------------------------------------------------------------------
--------------------------------------------------------------------------------
--------------------------------------------------------------------------------
--------------------------------------------------------------------------------
--------------------------------------------------------------------------------
-----------------------------------------------------ACACTAGTTGATCCTGCTAGACCCGAC
TGCTATCAGGATGAGGCTAAGCCATGCGAGTCGCGCGTCCCAAGCCATGGT----------------------------G
-GGAGCGGCAGACGGCTCAGTAACA-CGTAGTTAACCTGCCCTTAGGACGGGGATAACCGTGGGAAAGCAATGTAACATA
ATTAATTAACAGAGT--AG-TGTTACATGCCCCAAAA-CTGCGGCTAATACCCGAT-AGGGG--AAATG-GCCTGGAATG
GTATTT-CCCTCAAAGGGTTTTGGCGCCATGCT--CGCCAAAATCGCCTAAGGATGGGACTGCGGC---CT-ATTAGGTA
GT--CGGCGGTGTAACGGA--------------------------CCACCGAGCCTATAATAGGTACGGGCCATGTGAGT
GGGAGCCCGGAGATGGGAACTGAGACAAGGTCCCAGGCCCTACGGG--GCGCAGCAGGCGCGAAACCTCCACAATACACG
AAAGTGTGAT---GGGG---T-----------CATCCTGAGTGCCATCTGATGAAGATGGC---TTTTCATCGGC-----
-------------------------------TTAAGGAGCCGG------TGGAATAAGGGGAGGGCAAG-GCTGGTGCCA

```
GCCGCCGCGGTAAAACCAGCTCCTCGAGTGGTCGGGGTGATTATTGGGCCTAAAGCGTCCGTAGCCGGTCTAGTAAGTCC
TCGGTTAAATCCGGCAGCCTAACTG-TCGGCTT----------------------------------GCTGAGGATACT-
-----------------------------------GCTAGACTTGGGGGTGGGAGAGGCTAAAGGTACTCCCAGGGTAGG
GGTGAAATCCTATAATCCTGGGGGGACCACCAG-----------------------------------------------
-------------------------------------------------------------------TGGCGAAGGCGTT
TAGCTGGAACACGCCC-------------------------------------------------------GACGGTGAG
GGACGAAAGCTAGGGGAGCGAACCGGATTAGATACCCGGGTAGTCCTAGCTGTAAA------------------------
--------------------------------------------------------------------------------
--------CGATGCGGGCTAGGTGTTGGTCCGG-CTACGAGCCGGATCAGT---GCCGAAGAGAAGTTGTTAAGCCCGCC
GCCTGGGGAGTACGGCCGCAAGGCTGAAACTTAAAGGAATTGGCGGGGGAGCACCACA-AGGGGTGAAGCC-TGCGGTTT
AATTGGACTCAACGCCGGGAAGCT-TACCGGAGAGACAGCTGGATGATAGCCA---------------------------
---------------------AGCTAAAGACTTTGCTAGACT----AGCTGAGAGGTGGTGCATGGCCGTCGCCAGTTCG
TGCCG-TGAGGTG--TCCTGTTAAGTCAGGCAAC----------------------------------------------
--------------------------------------------------------------------------------
----------G------AACGAGACCCGTACCCTTATTTACCAG---CGGAT--CCCGATGG-GATGCCGGGTACAATAA
GGGGACTGC----CCTCGATA-AGATGGAGGAAGGAGCGGGCCACGGCAGGTCA-GTATGC-CCCGAATCTCCCG-GGCC
ACACGCGGGCTGCAATGGCTAGGACAATGGGTTT-CGACACCGAAAGGTGGAG---------------------------
---------------------------------------------------ATAATCCCCTAAACCTAGTCTCGGTTGGG
ATCGAGGGTTGCAACCCACCCTCGTGAAC--ATGGAATCCCTAGTAACCGCGTGTCACCATCG-CGCGGTGAATACGTCC
CTGCTCCTTGCACACACCGCCCGTCGCTCCATCCGAGTTGGGTCTAGATGAGGTTCACTCCTCA----------------
---------------------------GGGGTGAATCGAATCTAGGCTCGGCAAGGAGGGAGAAGTCGTAACAAGGTAGC
CGTAGGGGAACCTGCGGCTGGATCACCTCCT-------------------------------------------------
------------------------------------------------------
>AAHC01014171.47.1617 Eukaryota;Excavata;Metamonada;Parabasalia;Trichomonadea;Trichomonas;Trichomonas
vaginalis G3 Tva
---CTTGGTTGATCCTGC--------------------------------------------------------------
--------------------------------------------------------------------------------
--------------------------------------------------------------------------------
---------------------------------------------------------------------------CAAGG
AAGCACACTTAGGTCAT-------AGATTAAGCCATGCAAGTGTTAGTTCA-----------GGTAACGA---AACTGCG
AATAGCTCA----------------------TTAATACGCTCAGAATCTATTTGGCG-----------------------
-------------------------------------------------------------------------------G
CGACCAACAGGTCTTAAATGGATAGCAGC------------------------AGCA-ACTCTGGT---GC----TAATA
C-ATGCGATTGTTTCTC--CAGATGTGAATTA-TGGAGGAAAAGTTGAC-CTCATCAGAGGCACGCCATTCGACTGAGTG
ACCTATCAGCTTGTA----------CTTAGGGTCTTTACCTAGGTAGGCTATCACGGGTAACGGGCGGTTACCGTCGGAC
TGCCGGAGAAGGCGCCTGAGAGATA-GCGACTATATC-----CACGGG----T--AGCAGCAGGCGC--------GAAAC
TTTCCCACTCGAGACTTTCGGAGGAGGTAATGACCAG-TTCCATTGGTGCCT------------TTTGGTACTGTGGATA
GG----GGTACGGTTTT-----CC-AC--CGT---ACCGAAACCTAGCAGAGGGCCAGTCTGGTGCCAGCAGCTGCGGTA
ATTCCAGCTCTGCGAGTTTGCTCCCATATTGTTGCAGTTAAAACGCCCGTAGTCTGA---------------ATTGGCCA
GCAATGGTCGTACGTATTTTTACGTTCACT--------------------------------------------------
--------------------------------------------------------------------------------
--------------------------------------------------------------------------------
--------------------------------------------------------------------------------
---------GTGAACAAATCAG-------------GACG-----CTTAGAGTATGGCCACA---------TGAATGACTC
AGCGCAGTATGAAGTCTTTGTTTTCTTCCGAAAACAAGCTCAATGAGAGCCATCGGGGGTAGATCTATCTCATGACGAGT
GGTGGAATACTTTGACTCATGAGA------------------------------------GAGAAGCTG-----------
--------------------------------------------------------------------------------
------------------------------------------AGGCGAAGGCGTCTACCTAGAGGTTTTCTGTCGATCAA
GGGCGAGAGTAGGAGTATCCAACAGGATTAGAGACCCTGGTAGTTCCTACCTTAAACGATGCCGACAGGA----------
--------------------------------------------------------------------------------
---------------------------GTTT-GTCATTTGTTAATGGCAGAATCTTTGGAGAAATCATAGTTCTTGGGCT
CTGGGGGAACTACGACCGCAAGGCTGAAACTTGAAGGAATTGACGGAAGGGCACACC--AGGGGTGGAGCC-TGTGGCTT
AATTTGAATCAACACGGGGAAACTTACCAGGACCAGATGTTTTTTATGACTGACAGGCTTCGGGTCTTTCA---GGATA-
----------------------------------------------TTGTTTTTGGTGGTGCATGGCCGTTGGTGGTGCG
TGGGT-TGACCTGTCTAGCGTTGATTCAGCTAACGAGCGAGATTATCGCCAATTATTTACTTTGCCGAAGTCCTTCGGTT
AAAGTTCT------------------------------------------------------------------------
----------------------------------------------------------------------------AAT-
TGGGACTC---CCTGCGATTTTAGCAGGTGGAAGAGGGTAGCAATAACAGGTCCGTGATGCCCTTTAGATGCTCTGGGCT
GCACGCGTGCTACAATGTTAGGATCAATAGGACT-GCGAGCCT-------------------------------------
---------------------------------------GAGAGGGTGCGCTACTCTT-ATAATCCCTAACGTAGTTGGG
AT-TGACGTTTGTAATCAGCGTCATGAAC--CAGGAATCCCTTGTAAATGTGTGTCAACAACG-CACGTTGAATACGTCC
CTGCCCTTTGTACACACCGCCCGTCGCTCCTACCGATTGGATGACTCGGTGAAATCACCGGATGCTTACGAGCAGA----
-----------------------------AAGTGATTAAATCACGTTATCTAGAGGAAGGAGAAGTCGTAACAAGGTAAC
GGTAGGTGAACCTGCCGTTG------------------------------------------------------------
------------------------------------------------------
>lcl|NC_007268.2_rrna_XR_002460813.1_5427 [locus_tag=LMJF_27_rRNA_06] [db_xref=GeneID:12982905] [product=18S
ribosomal RNA (SSU) RNA] [transcript_id=XR_002460813.1] [location=1052349..1054552] [gbkey=rRNA] Lma
-GATCTGGTTGATTCTGCC-------------------------------------------------------------
--------------------------------------------------------------------------------
--------------------------------------------------------------------------------
---------------------------------------------------------------------------AGTAG
TCATATGCTTGTTTCAA-------GGACTTAGCCATGCATGCCTCAGAATCACTGCA----TTTGCAGGA---ATCTGCG
CATGGCTCA----------------------TTACATCAGACGTAATCTGCCGCAAAAATCTTGCGGTTTCCGCAAAA--
--TTGGATAACTTGGCGAAAC------GCCAAGCTAATACATGAACCAACCGGGTGTTCTCCACTCCAGACGGTG----G
GCAACCATCGTCGTGAGACGCCCAGCGAAT--GAATG--ACAGTAAAACCAATGCCT-TCACTGGC---AG----TAACA
CCCAGCAGTGTTGACTC-------A--ATTCATTCCGTGCGAAAGCCGG-CTTGTTCCGGCGTCTTTTGACGAACAACTG
CCCTATCAGCTGGTG--------ATGGCCGTGTAGTGGACTGCCATGGCGTTGACG-GGAGCGGGGG-ATTAGGGTTCGA
TTCCGGAGAGGGAGCCTGAGAAATA-GCTACCACTTC-----TACGGA----G--GGCAGCAGGCGC--------GCAAA
TTGCCCAATGTCAAAACAAAACGATGAGGCAGCGAAA-AGAAATAGAGTTGTCAGT-CCATTTGGATTGTCATTTCAATG
GGGGATATTTAAACCCA-----TC-CA--ATA---TCGAGTAACAATTGGAGGACAAGTCTGGTGCCAGCACCCGCGGTA
ATTCCAGCTCCAAAAGCGTATATTAATGCTGTTGCTGTTAAAGGGTTCGTAGTTGAA---------------CTGTGGGC
TGTGCAGGTTTGTTCCTGGTCGTCCCGTCCATGTCGGATTTGGTGA--CCCAGGCCCTTGCAGCCCGTGAACATTCAAAG
AAACAAGAAACACGGGAGTGGTTCCTTTCCTGATTTACGCATGTCATGCATGCCAGGGGGCGTCCGTGATTTTTTACTGT
GACTAAAGAAGC------GTGACT-------AAAGCAGTCATTTGACTTGAATTAGAAAGCATGGGATAACAAAGGAGCA
GCCTCTAGGCTACCGTTTCGGCTTTTGTTGGTTTTAAAGGTC-----TATTGGAGATTATGGAGCTGTGCGACAAGT---
-GCTTTCCCATCGCAACTTCGGTTC-GG-TGTGTGGCGCCTTTGAGGGGTTTAGTGCGTCCGGTGCGAGCTCCGGTTCGT
CCGGCCGTAACGCCTTTTCAACTCACGGCCTCTAGGAATGAAGGAGGGTAGTTCGGGGGAGAACGTACTGGGGCGTCAGA
```

GGTGAAATTCTTAGACCGCACCAA------------------------------------GACGAACTA-----------
--------------------------------------------------------------------------------
------------------------------------------CAGCGAAGGCATTCTTCAAGGATACCTTCCTCAATCAA
GAACCAAAGTGTGGAGATCGAAGATGATTAGAGACCATTGTAGTCCACACTGCAAACGATGACACCCATGAATTGGGGAT
CTTATGGGCCGGCCTGCGGCAGGGTTTACCCTGTGTCAGCACCGCGCCCGCTTTTACCAACTTACGTATCTTTTCTATTC
GGCCTTTACCGGCCA-CCCACGGGAATATCC-TCAGCACGTTTTCTGTTTTTTCACGCGAAAGCTTTGAGGTTACAGTCT
CAGGGGGGAGTACGTTCGCAAGAGTGAAACTTAAAGAAATTGACGGAATGGCACCACA-AGACGTGGAGCG-TGCGGTTT
AATTTGACTCAACACGGGGAACTTTACCAGATCCGGACAGGATGAGGATTGACAGATTGAGTGTTCTTTCT---CGATT-
----------------------------------------------CCCTGAATGGTGGTGCATGGCCGCTTTTGGTCGG
TGGAG-TGATTTG--TTTGGTTGATTCCGTCAACGGACGAGATCCAAGCTGCCCAGTAGAATTCAGAATTGCCCATAGAA
TAGCAAACTCATCGGCGGGTTTTACCCAACGGTGGGCCGCA---------------------------------------
----------------------------------------------------------TTCG-GTCGAA---TTCTTCT-
CTGCGGGATTCCTTTGTAATTGCACAAGGTGAAATTTTGGGCAACAGCAGGTCTGTGATGC-TCCTCAATGTTCTGGGCG
ACACGCGCACTACAATGTCAGTGAGAACAAGAAA-AACGACTTTTGTCGAACC----------------------TAC--
-------------------------T-----TG-ATCAAAAGAGTGGGAAACCCCGGAATCACATAGACCCACTTGGGA
CCGAGGATTGCAATTATTGGTCGCGCAAC--GAGGAATGTCTCGTAGGCGCAGCTCATCAAAC-TGTGCCGATTACGTCC
CTGCCATTTGTACACACCGCCCGTCGTTGTTTCCGATGATGGTGCAATACAGGTGATCGGACAGG---------------
---------CGGTGTTTTATCCGCCCGAAAGTTCACCGATATTTCTTCAATAGAGGAAGCAAAAGTCGTAACAAGGTAGC
TGTAGGTGAACCTGCAGCTGGATCATTTT---------------------------------------------------
------------------------------------------------------
>lcl|NC_003279.8_rrna_NR_000053.1_5462 [gene=rrn-1.1] [locus_tag=CELE_F31C3.7] [db_xref=GeneID:13183153]
[product=18S ribosomal RNA] [transcript_id=NR_000053.1] [location=15062083..15063836] [gbkey=rRNA] Cel
ATACCTGATTGATTCTGT--------------------------------------------------------------
--------------------------------------------------------------------------------
--------------------------------------------------------------------------------
---------------------------------------------------------------------------CAGCG
CGATATGCTCAAGTAAA-------AGATTAAGCCATGCATGCTTTGAT------------TCATCAATGA---AATTGCG
TACGGCTCA----------------------TTAGAGCAGATATCACCTTATCCGG---------GA----TCCTCAT--
--ATGGATAACTGCGGAAATA------CTGGAGCTAATACATGCAACTATACCCCAACGCAAGGC---------------
-GGGGTGCAATTATTAGAACAGACCAAAC-----------------------------------------G--TTTTCGG
ACGTTGTTTGTTGACTCTGAATAAA-------GCAGTT--TACTGTCAG-TTTCGACTGACTCTATCCGGAAAGGGTGTC
TGCCCTTTCAACTAG--------ATGGTAGTTTATTGGACTACCATGGTTGTTACGGGTAACGGAGA-ATAAGGGTTCGA
CTCCGGAGAGGGAGCCTTAGAAACG-GCTACCACGTC-----CAAGGA----A--GGCAGCAGGCGC--------GAAAC
TTATCCACTGTTGAGTATGAGATAGTGACTAAAAATA-TAAAGACT--CATCCTTT-----TGGATGAGTTATTTCAATG
AG---TTGAAT-ACAAA-----TG-AT--TCT---TCGAGTAGCAAGGAGAGGGCAAGTCTGGTGCCAGCAGCCGCGGTA
ATTCCAGCTCTCCTAGTGTATCTCGTTATTGCTGCGGTTAAAAAGCTCGTAGTTGGA---------------TCTAGGTT
ACGTGCCGCAGTTCGCAATTTGCGTCAACTGTGGTCGTGACTTCTA--ATT-----------------------------
--------------------------------------TG---------------CTGG---TT----------------
----------T------------------------GAGGTTGGGTTCGCCCTTCAA------------------------
---------------CTGCCAGCAGGTTTACCTTGAATAAA--------TCAGAGTGCTCAATACAAGC-----------
---GCTTGCTTGAATAGCTCAT----------------------CATGGAATAATGAAACA----GGACTTCGGTTCTT-
----------TTTGTTGGTTCTAGAACTGATTTAATGGTTAAGAGGGACAAACCGGGGGCATTCGTATCATTACGCGAGA
GGTGAAATTCGTGGACCGTAGTGA------------------------------------GACGCCCAA-----------
--------------------------------------------------------------------------------
------------------------------------------CAGCGAAAGCATTTGCCAAGAATGTCTTCATTAATCAA
GAACGAAAGTCAGAGGTTCGAAGGCGATTAGATACCGCCCTAGTTCTGACCGTAAACGATGCCATCTCGCGATTCGGAGG
GTTTTT--------------------------------------------------------------------------
---------------------------------------GCCCTGCCGAGGAGCTATCCGGAAACGAAAGTCTTTCGGTT
CCGGGGGTAGTATGGTTGCAAAGCTGAAACTTAAAGAAATTGACGGAAGGGCACCACA-AGGCGTGGAGCT-TGCGGCTT
AATTTGACTCAACACGGGAAAACTCACCCGGTCCGGACACCATTAGGACTGACAGATTGAAAGCTCTTTCT---CGATT-
----------------------------------------------TGGTGGTTGGTGGTGCATGGCCGTTCTTAGTTGG
TGGAG-TGATTTG--TCTGGTTTATTCCGATAACGAGCGAGACTCTAGCCTGCTAAATAGTTGGCGAATCTTCGGGTTCG
TATAACTTCTT---------------------------------------------------------------------
-----------------------------------------------------------------------------AG-
AGGGATAAGC-------GGTGTTTAGCCGCACGAGATTGAGCGATAACAGGTCTGTGATGC-CCTTAGATGTCCGGGGCT
GCACGCGTGCTACACTGGTGGAGTCAGCGGGTTT-TTCCTA---------------------------------------
----------------------------------TGCCGAAAGGTATCGGTAAACCGT-TGAAATTCTTCCATGTCCGGG
ATAGGGTATTGTAATTATTGCCCTTAAAC--GAGGAATGCCTAGTAAGTGTGAGTCATCAGCT-CACGTTGATTACGTCC
CTGCCCTTTGTACACACCGCCCGTCGCTATCCGGGACTGAACTGATTCGAGAAGAGTGGGGACTGTCGCTTCGAGGTTTA
ACGA---------CTTCGTTGTTGCGGAAACCATTTTTATCGCATTGGTTTGAACCGGGTAAAAGTCGTAACAAGGTAGC
TGTAGGTGAACCTGCAGCTGGATCATCG----------------------------------------------------
------------------------------------------------------
>lcl|NW_007931121.1_rrna_NR_133555.1_35008 [gene=18SrRNA:CR45838] [locus_tag=Dmel_CR45838]
[db_xref=FLYBASE:FBtr0346878,GeneID:26067166] [product=18S ribosomal RNA] [transcript_id=NR_133555.1]
[location=55965..57959] [gbkey=rRNA] Dme
-ATTCTGGTTGATCCTGCC-------------------------------------------------------------
--------------------------------------------------------------------------------
--------------------------------------------------------------------------------
---------------------------------------------------------------------------AGTAG
TTATATGCTTGTCTCAA-------AGATTAAGCCATGCATGTCTAAGTACACACGAA---TTAAAAGTGA---AACCGCA
AAAGGCTCA----------------------TTATATCAGTTATGGTTCCTTAGATCGTTAACAGT---------TAC--
--TTGGATAACTGTGGTAATT------CTAGAGCTAATACATGCAATTAAAACATGAACCTTATGGGA------------
-CATGTGCTTTTATTAGGCTAAAACCAAGC---GATC------------------------------------GCAAGAT
CGTTATATTGGTTGAA---CTCTAGATAACAT-GCAGATCGTATGGTCT-TGTACCGACGACAGATCTTTCAAATGTCTG
CCCTATCAACTTTTG--------ATGGTAGTATCTAGGACTACCATGGTTGCAACGGGTAACGGGGA-ATCAGGGTTCGA
TTCCGGAGAGGGAGCCTGAGAAACG-GCTACCACATC-----TAAGGA----A--GGCAGCAGGCGC--------GTAAA
TTACCCACTCCCAGCTCGGGGAGGTAGTGACGAAAAA-TAACAATACAGGACTCATATCCGAGGCCCTGTAATTGGAATG
AG---TACACT-TTAAA-----TC-CT--TTA---ACAAGGACCAATTGGAGGGCAAGTCTGGTGCCAGCAGCCGCGGTA
ATTCCAGCTCCAATAGCGTATATTAAAGTTGTTGCGGTTAAAACGTTCGTAGTTGAA---------------CTTGTGCT
TCATACGGGTAGTACAACTTACAATTGTGGTTAGTACTATACCTTT--ATGTATGTAAGCGTATT---------ACCGG-
------------TGGAGTTCTTATATGTGATTAAATACTT---------------GTAT---TTTTTCATATGTTCCTCC
TATTTAAAAACC------TGCATT-------AGTGCTCTTAAACGAGTGTTATTGT------------------------
---------------GGGCCGGTACTATTACTTTGAACAAA--------TTAGAGTGCTTAAAGCAGGCTTCAAATGC--
--------CTGAATATTCTGTG----------------------CATGGGATAATGAAATA----AGACCTCTGTTCTG-
----C--TTTCATTGGTTTTCAGATCAAGAGGTAATGATTAATAGAAGCAGTTTGGGGGCATTAGTATTACGACGCGAGA
GGTGAAATTCTTGGACCGTCGTAA------------------------------------GACTAACTT-----------
--------------------------------------------------------------------------------

------------------------------------------AAGCGAAAGCATTTGCCAAAGATGTTTTCATTAATCAA
GAACGAAAGTTAGAGGTTCGAAGGCGATCAGATACCGCCCTAGTTCTAACCATAAACGATGCCAGCTAGCAATTGGGTGT
AGCTACTT------------------------------------------------------------------------
-------T---------------------------TATGGCTCTCTCAGTCGCTTCCCGGGAAACCAAAGCTTTTGGGCT
CCGGGGGAAGTATGGTTGCAAAGCTGAAACTTAAAGGAATTGACGGAAGGGCACCACC-AGGAGTGGAGCC-TGCGGCTT
AATTTGACTCAACACGGGAAAACTTACCAGGTCCGAACATAAGTGTGTAAGACAGATTGATAGCTCTTTCT---CGAAT-
----------------------------------------------CTATGGGTGGTGGTGCATGGCCGTTCTTAGTTCG
TGGAG-TGATTTG--TCTGGTTAATTCCGATAACGAACGAGACTCAAATATATTAAATAGATATCTTCAGGATTATGGTG
CTGAAGCTTATGTAGC-----CTTCATTCATGTTGGCAGTAAAATGCTTATTGTGT------TTGAATGTGTTTATGTAA
GTGGA------------GCCGTA--CCTGTTGGTTTGTCCC-------------ATTATAAG-GACACTAGCTTCTTAA-
ATGGACAAA------TTGCGTCTAGCAATAATGAGATTGAGCAATAACAGGTCTGTGATGC-CCTTAGATGTCCTGGGCT
GCACGCGCGCTACAATGAAAGTATCAACGTGTAT-TTCCTAGACCG----------------------------------
---------------------------------------AGAGGTCCGGGTAAACCGC-TGAACCACTTTCATGCTTGGG
ATTGTG-AACTGAAACTGTTCACATGAAC--TTGGAATTCCCAGTAAGTGTGAGTCATTAACT-CGCATTGATTACGTCC
CTGCCCTTTGTACACACCGCCCGTCGCTACTACCGATTGAATTATTTAGTGAGGTCTCCGGACGTGATCACTGTGACGCC
TTGC----GTGTTACGGTTGTTTCGCAAAAGTTGACCGAACTTGATTATTTAGAGGAAGTAAAAGTCGTAACAAGGTTTC
CGTAGGTGAACCTGCGGAAGGATCATTA----------------------------------------------------
------------------------------------------------------
>lcl|NC_004326.2_rrna_XR_002273101.1_1180 [locus_tag=PF3D7_0531600] [db_xref=GeneID:9221861] [product=18S ribosomal RNA] [transcript_id=XR_002273101.1] [location=1289601..1291692] [gbkey=rRNA] Pfa
-AACCTGGTTGATCTTGCC-------------------------------------------------------------
--------------------------------------------------------------------------------
--------------------------------------------------------------------------------
---------------------------------------------------------------------------AGTAG
TCATATGCTTGTCTCAA-------AGATTAAGCCATGCAAGTGAAAGTATATATATATTTTATATGTAGA---AACTGCG
AACGGCTCA----------------------TTAAAACAGTTATAGTCTACTTGACATTTTTATTA-----------T--
--AAGGATAACTACGGAAAAG------CTGTAGCTAATACTTGCTTTATTATCCTTTGATTTTTATCTTTGG--------
--ATAAGTATTGTTAGGCCTTATAAGAA-----------------------------------------AA----AAGTT
ATTAACTTAAGGAATTA--TAACAAAGAAGTAACACGTAATAAATTTAT-TTTATTTAGTGTGTATCAATCGAGTTTCTG
ACCTATCAGCTTTTG--------ATGTTAGGGTATTGGCCTAACATGGCTATGACGGGTAACGGGGA-ATTAGAGTTCGA
TTCCGGAGAGGGAGCCTGAGAAATA-GCTACCACATC-----TAAGGA----A--GGCAGCAGGCGC--------GTAAA
TTACCCAATTCTAAAGAAGAGAGGTAGTGACAAGAAA-TAACAATGCAAGGCCAAT-TT-TTGGTTTTGTAATTGGAATG
GT---GGGAAT-TTAAA-----AC-CT--TCC---CAGAGTAACAATTGGAGGGCAAGTCTGGTGCCAGCAGCCGCGGTA
ATTCCAGCTCCAATAGCGTATATTAAAATTGTTGCAGTTAAAACGCTCGTAGTTGAA---------------TTTCAAAG
AATCGATATTTTATTGTAACTATTCTAGGGGAACTATTTAGCTTT--TCGCTTTAATACGCTTC----------CTCTA-
------------TTATTATGT--TCTT----TAAATAACA---------------AAGA---TT---------------C
TTTTTAAAATCC-----------------------CCACTTTTGCTTTTGCTTTTT------------------------
---------------TTGGGGATTTTGTTACTTTGAGTAAA--------TTAGAGTGTTCAAAGCAAACAGTTAAAGCAT
TTACTGTGTTTGAATACTATAG----------------------CATGGAATAACAAAATT----GAACAAGCTAAAATT
TTTTG--TTCTTTTTTCT-TATTTTGGCTTAGTTACGATTAATAGGAGTAGCTTGGGGACATTCGTATTCAGATGTCAGA
GGTGAAATTCTTAGATTTTCTGGA------------------------------------GACGAACAA-----------
--------------------------------------------------------------------------------
------------------------------------------CTGCGAAAGCATTTGTCTAAAATACTTCCATTAATCAA
GAACGAAAGTTAAGGGAGTGAAGACGATCAGATACCGTCGTAATCTTAACCATAAACTATGCCGACTAGGTGTTGGATGA
AAGTGTTAAAAATAAAAGTCATCT------------------------------------------------TTCGAGGT
GACTTTTA---------------------------GATTGCTTCCTTCAGTACCTTATGAGAAATCAAAGTCTTTGGGTT
CTGGGGCGAGTATTCGCGCAAGCGAGAAAGTTAAAAGAATTGACGGAAGGGCACCACC-AGGCGTGGAGCT-TGCGGCTT
AATTTGACTCAACACGGGGAAACTCACTAGTTTAAGACAAGAGTAGGATTGACAGATTAATAGCTCTTTCT---TGATT-
----------------------------------------------TCTTGGATGGTGATGCATGGCCGTTTTTAGTTCG
TGAATATGATTTG--TCTGGTTAATTCCGATAACGAACGAGATCTTAACCTGCTAATTAGCGGCGAGTACACTATATTCT
TATTTGAAATTGAACATAGGTAACTATACATTTATTCAGTAATCAAATTAGGATATTTTTATTAAAATATCCTTTTCCCT
GTTCTACTAATAATTTGTTTTTTACTCTATTTCTCTCTTCTTTTAAGAATGTACTTGCTTGA-TTGAAAAGCTTCTTAG-
AGGAACATT-----GTGTGTCTAACACAAGGAAGTTTAAGGCAACAACAGGTCTGTGATGT-CCTTAGATGAACTAGGCT
GCACGCGTGCTACACTGATATATATAACGAGTTT-TTAAAAATATGCTTATAT----------------------TTGTA
TCTTTGATGCTTATATTTTGCATACTTTTCCTCCGCCGAAAGGCGTAGGTAATCTTTA-TCAATATATATCGTGATGGGG
ATAGATTATTGCAATTATTAATCTTGAAC--GAGGAATGCCTAGTAAGCATGATTCATCAGAT-TGTGCTGACTACGTCC
CTGCCCTTTGTACACACCGCCCGTCGCTCCTACCGATTGAAAGATATGATGAATTGTTTGGACAAGAAAAATTGAATTAT
AT------------TCTTTTTTTTCTGGAAAAACCGTAAATCCTATCTTTTAAAGGAAGGAGAAGTCGTAACAAGGTTTC
CGTAGGTGAACCTGCGGAAGGATCATTA----------------------------------------------------
------------------------------------------------------
>CABZ01002707.943.2826 Eukaryota;Opisthokonta;Holozoa;Metazoa (Animalia);Eumetazoa;Bilateria;Chordata;Vertebrata;Gnathostomata;Euteleostomi;Actinopterygii;Neopterygii;Teleostei;Danio rerio (zebrafish) Dre
---CCTGGTTGATCCTGC--------------------------------------------------------------
--------------------------------------------------------------------------------
--------------------------------------------------------------------------------
---------------------------------------------------------------------------CAGTA
ATATATGCTTGTCTCAA-------AGATTAAGCCATGCAAGTCTAAGTGCACACGG-AC-GGNACAGTGA---AACTGCG
AATGGCTCA----------------------TTAAATCAGTTATGGTTCCTTTGATCGCTCCATCC-------GTTAC--
--TTGGATAACTGTGGCAATT------CCAGAGCTAATACATGCCAACGAGCGCCGACCCGGTCCCCCCGGGGGCCGGGG
ACGCGTGCATTTATCAGATCCAAAACCCAT--CCGGGTGCGGCGTGGGCTCCCCCCT-CGGGGGTCGGGCCTCGTCCCCC
GGCCGCTTTGGTGACTC--TAGATAACCTCGGGCCGATCGCGCGCCCTC-CGCGGCGGCGACGATTCATTCGAATGTCTG
CCCTATCAACTTTCG--------ATGGTACTTTAGGCGCCTACCATGGTGACCACGGGTAACGGGGA-ATCAGGGTTCGA
TTCCGGAGAGGGAGCCTGAGAAACG-GCTACCACATC-----CAAGGA----A--GGCAGCAGGCGC--------GCAAA
TTACCCATTCCCGACACGGGGAGGTAGTGACGAAAAA-ATAACAATACAGGTCTCTTTC-GAGGCCCTGTAATGGGAATG
AG---CGTATCCTAAAC-----CC-AT--GGG---GCGAGGACCCATTGGAGGGCAAGTCTGGTGCCAGCAGCCGCGGTA
ATTCCAGCTCCAATAGCGTATATTAAAGTTGCTGCAGTTAAAAAGCTCGTAGTTGGA---------------TCTCGGGA
GTGGGCTGGCGGTCCGCCGCGAGGCGAGCCACCGCCTGTCCCGGAC--CCTGCCTCCCGGCG--------------CCC-
------------CCCGG---------------------AT---------------GC-----------------------
-------CCTTA-----------------------GCTGGGTGTCCGGTACCTCGG------------------------
---------------GGCCCGGAGCGTTTACTTTGAAAAAA--------TTAGAGTGTTCAAAGCAGGCCGCCC------
-AGCCGCCGCTGAATACCGCAG----------------------CTAGGAATAATGGAATA----GGACTCCGGTTCTA-
-------TTTTGTGGGTT-TCTGGAACCCGGGGCCATGATTGAGAGGGACGGCCGGGGGCATTCGTATTGCGCCGCTAGA
GGTGAAATTCTTGGACCGGCGCAA------------------------------------GACGGACCG-----------
--------------------------------------------------------------------------------
------------------------------------------GAGCGAAAGCGTTTGCCAAGAACGTTTTCATTAATCAA
GAACGAAAGTCGGAGGTTCGAAGACGATCAGATACCGTCGTAGTTCCGACCGTAAACGATGCCGACCCGCGATCCGGCGG

```
CGTTACTACCAC--------------------------------------------------------------------
-------A----------------------------ATGACCCGCCGGGCAGCGTGTGGGAAACCACGAGTCTCTGGGCT
CTGGGGGAGTATGGTTGCAAAGCTGAAACTTAAAGGAATTGACGGAAGGGCACCACC-AGGAGTGGAGCC-TGCGGCTT
AATTTGACTCAACACGGGGAACCTCACCCGGCCCAGACACGGAAAGGATTGACAGATTGATAGCTCTTTCT---CGATT-
----------------------------------------------CTGTGGGTGGTGGTGCATGGCCGTTCGTGGTTGG
TGGAG-TGATTTG--TCTGGTTGATTCCGATAACGAACGAGACTCTGGCATGCTAACTAGTTACGCGGCCCCGCGCGGTC
AGCGT----CTGCAAC----------------------------------------------------------------
------------------------------------------------------------------------TTCTTAG-
AGGGACAAGTGGCGTT-------CAGCCACGCGAGACTGAGCAATAACAGGTCTGTGATGC-CCTTAGATGTCCGGGGCT
GCACGCGCGCCACAATGGGCGGATCAACGTGTGC-CTACCC---------------------------------------
-------------------------------TG-CGCCGAGAGGCGCGGGTAACCCGT-TGAAACCCGCTCGTGATGGGG
ACCGGGGATTGCAACTATTTCCCTTGAAC--GAGGAATTCCCAGTAAGCGCTGGTCATCAGCC-AGCGCTGATTAAGTCC
CTGCCCTTTGTACACACCGCCCGTCGCTACTACCGATTGAGCGGCACAGTGAGGTCCTCGGATCGGCCCCGCCCGGGGCC
CTTTATCGAGCCCTGGTGGTACGCTGAGAAGACGATCGAACTCTGTCGTTTAGAGGAAGTAAAAGTCGTAACAAGGTTTC
CGTAGGTGAACCTGCGGAAG------------------------------------------------------------
------------------------------------------------------
>AAXT01000001.292853.294499 Eukaryota;SAR;Alveolata;Apicomplexa;Aconoidasida;Piroplasmorida;Babesia;Babesia
bovis Bbo
---CCTGGTTGATCCTGCC-------------------------------------------------------------
--------------------------------------------------------------------------------
--------------------------------------------------------------------------------
---------------------------------------------------------------------------AGTAG
TCATATGCTTGTCTTAA-------AGACTAAGCCATGCATGTCTAAGTACCAGCTT----GTTACGGTGA---AACTGCG
AATGGCTCA----------------------TTACAACAGTTATAGTTTCTTTGGAGTTCACTTTG----------CT--
--ATGGATAACCGCGCTAATT------GTGTGGCTAATACACGTTTGAGGGTTTTCCCGCGTTT----------------
------------------------------------------------------------------------------A-
--CTGGTCTTGT---------------GATTTACAGTAACCTGCGACTC-GCTTTTTGCGATATTCCATTCAAGTTTCTG
ACCCATC--AGCTTG--------ACGGTAGGGTATTGGCCTCCCGAGGCATCGACGGGTAACGGGGA-ATTAGGGTTCGA
TTCCGGAGAGGGAGCCTGCGAGACG-GCTACCACATC-----TAAGGA----A--GGCAGCAGGCGC--------GCAAA
TTACCCAATCCTGACACAGGGAGGTAGTGACAAGAAA-TACCAATA-CGGGGCTAC------TGCTCTGTAATTGGCATG
GG---GGCGAC-CTTCA-----CC-CT--CGC---CCGAGTACCCATTGGAGGGCAAGTCTGGTGCCAGCAGCCGCGGTA
ATTCCAGCTCCAATAGCGTATATTAAACTTGTTGCAGTTAAAAAGCTCGTAGTTGAA---------------TCTCACGT
CCCCCGCTTGGTCCTTTCCTCGCCGGGACG--------------------------------------------------
--------------------------------------------------------------------------------
--------------------------------------------------------------------------------
----------------------CCTCGTTACTTTGAGAAAA--------TTAGAGTGTTTCAAGCAGG------------
---TTTCGCCTGTATAATTGAG----------------------CATGGAATAACCTTGTA----TGA--------CCC-
-------TGTCGTACCGT-TGGTTGACTTTGGGTAATGGTTAATAGGAACGGTTGGGGGCATTCGTACTCGACTGTCAGA
GGTGAAATTCTTAGATTTGTCGAT------------------------------------GACGCACGA-----------
--------------------------------------------------------------------------------
------------------------------------------CTGCGAAAGCATTTGCCAAGGACGCTTCCATTAATCAA
GAACGAAAGTTAGGGGATCGAAGACGATCAGATACCGTCGTAGTCCTAACCTTAAACTATGCCGACTAGGGATTGGTGGT
CGTCGCC-------------------------------------------------------------------------
-------------------------------------TCGACTCCGTCAGCACCTTGAGAGAAATCAAAGTCTTTGGGTT
CTGGGGGAGTATGGTCGCAAGTCTGAAACTTAAAGGAATTGACGGAAGGGCACCACC-AGGCGTGGAGCC-TGCGGCTT
AATTTGACTCAACACGGGGAACCTCACCAGGTCCAGACAGCGTAAGGATTGACAGGTTGATTGTCCTTTCT---TGATT-
----------------------------------------------CTCTGGGTAGTGGTGCATGGCCGTTCTTAGTTGG
TGGAG-TGATTTG--TCTGGTTAATTCCGTTAACGAACGAGACCTTAACCTGCTATTTAGTCGCTCGGTCTTTGTCCGTG
CGCACTT-------------------------------------------------------------------------
--------------------------------------------------------------------------CATAG-
AGGGACTC---TGCGGCGTCAAGCTGCGGTGAGGTTTAAGGCAATAACAGGTCTGTGATGC-CCTTAGATGTCCTGGGCT
GCACGCGCGCTACACTGATGCTTGCCTCGTGTTT-ACCCTCGGCCGA---------------------------------
----------------------------------------TAGG--CCCTGGTAACCC-CTAGTCTGCATCGTGTTTGGG
ATTGATCTTTGCAATTCTAGATCATGAAC--GAGGAATGCCTAGTATGCGCAAGTCATCAGCT-TGTGCAGATTACGTCC
CTGCCCTTTGTACACACCGCCCGTCGCTCCTACCGATCGAGTGATCCGGTGAATTATTCGGAACCTGCTTGCGCGATCCG
T------------CGCCGCTCGCGGGCCAAGTTTGTGAACCTTATCACTTAAAGGAAGGAGAAGTCGTAACAAGGTTTC
CGTAGGTGAACCTGCGGAAG------------------------------------------------------------
------------------------------------------------------
>lcl|NW_013657636.1_rrna_XR_001290156.1_8249 [locus_tag=AMSG_20099] [db_xref=GeneID:25570478] [product=18S
ribosomal RNA] [transcript_id=XR_001290156.1] [location=complement(10307..12135)] [gbkey=rRNA] Ttr
--ACCTGGTTGATCCTGCC-------------------------------------------------------------
--------------------------------------------------------------------------------
--------------------------------------------------------------------------------
---------------------------------------------------------------------------AGTGG
TCATATGCTTGTCTCTA-------GGACTAAGCCATGCACGTATAAGTGTAAGCGA----TTTACAGTGA---AACTGCG
AATGGCTCA----------------------TTAAATCAGTAATAGTCTCTTTGATGGAAGCCGCT----------AC--
--ATGGATAACCGTAGAAAAT------CTAGAGCTAATACACGCCAAGGAGTCATGGCCTAGCATTTTTGCTGGG----G
CGTGCGGTATTTATTAGAGGCCAAGACGA------TG-------------------------------GCGCACTTGTGT
GTCTTCTTTGGTGAGTC--ATGATAACTGTACGAATCGAGGGCTTGTTC-GCAAGCGTCGATGGTTCATTCGAATTTCTG
CCCTATCAACTTTCG--------ATGGTAAGATAGAGGCTTACCATGGTGGTTACGGGTGACGGCGA-A-TGGGGTTCTA
TTCCGGAGAGGGGGCCTGAGAAACG-GCCACCACTTC-----CAAGGA----A--GGCAGCAGGCGC--------GCAAA
TTACCCAATGGCGACACGCCGAGGTAGTGACGAACAA-TAACGATG-GGACCCCTT------TGGGGCCCAATTGGAATG
AG---TACAAA-GTACA-----AG-CC--TTA---ACGAGGAACAATTGGAGGGCAAGTCTGGTGCCAGCAGCCGCGGTA
ATTCCAGCTCCAATAGCGTATATTAAAGCTGTTGCGGTTAAAAGCTCGTAGTTGGAT---------------TTGAGTCT
AGTTGCGGCCGGCTCAGGGCTTTGACTCTGGGGCGGACG-----TG--TAG-------------------------ACT-
------------GTATG---------------------CG---------------CG-----------------------
-------CGGGG-----------------------AGGGCAGGGAATTTTCTCTGC------------------------
---------------TTTCTCGTGTGCTTTACTTGAACAAA--------TTAGAGTGTTTAAAGCAGGCATT--------
---TTTGCCTTGAATACATTAG----------------------CATGGAATAACGATTTG----ACTGGAGCAG-GCA-
-------TGATGTTGTCT-TGATGACTGCGCAGTAATGATGGATAGGGACGGTCGGGGACATTCGTATTGCTCCGTCAGA
GGTGAAATTCTTAGATTGGAGGAA------------------------------------GACGAGCTG-----------
--------------------------------------------------------------------------------
------------------------------------------ATGCGAAAGCCTTTGTCAAGGACGTTTTCATTGATCAA
GAACGAAAGTTAG-GGATCGAAGACGATCAGATACCGTCGTAGTCTTAACCATAAACGATGCCGACCAGGGATTGGGGAA
GGCAGTAGACTT--------------------------------------------------------------------
-------C---------------------------TCACGACTTCCCCAGCACCTTATGAGAAATCTGAGTGTTTGG-GT
TCCGGGGGAGTATGGTCGCAAGGCTGAAACTTAAAGGAATTGACGGAAGGGCACCACC-AGGAGTGGAGCC-TGCGGCTT
```

AATTTGACTCAACACGGGA-AACTTACCAGGCCCAGACATAGCGAGGATTGACAGATTGAGAGCTCTTTCT---TGATT-
----------------------------------------------CTATGGATGGTGGTGCATGGCCGTTCTTAGTTCG
TGGAG-TGATTTG--TCTGCTTAATTGCGATAACGAACGAGACCTTGGGGGTTGCTGCGCGAGGGCGCTACTTGTTAGCG
GCCCGTGCT-----------------------------------------------------------------------
-----------------------------------------------------------------------------CC-
TACTCCTACTGCTGCGTGTTGTAACCAGAGGAAGTTTGAGGCTATAACAGGTCTGTGATGC-CCTTAGATGTCCTGGGCC
GCACGCGCTACACTGGAGCAGGCAGCGAGTGG-GGCTGCCGCCGCAAGGCG------------------------GGAGC
TA----------------------------CCT-GTGCCGGAAGGCACGGGTAATCTT-GCAAACTGGTCCGTGATGGGG
ATAGAT-GTTTGTAATTTGGATCTTGAAC--GAGGAATTCCTAGTAAACGCGCGTCAACAACG-CGCATTGACTACGT-C
CTGCCCTTTGTACACACCGCCCGTCGCTCCTACCGATTGAATGTCAAGGTGAGGGTTCTGGACTCTGCGCTCCTCTTGGG
C--------AACTGGGGGGGAAGCGGGGAAGTTGACCAAATCTTGGCATTTAGAGGAAGGAGAAGTCGTAACAAGGTTTC
CGTAGGTGAACCTGCGGAAGGATCATTA----------------------------------------------------
------------------------------------------------------
>CABX01000137.1970.3762 Eukaryota;SAR;Stramenopiles;Incertae Sedis;Blastocystis;Blastocystis hominis Bho
---TCTGGTCGATCCTGCC-------------------------------------------------------------
--------------------------------------------------------------------------------
--------------------------------------------------------------------------------
---------------------------------------------------------------------------AGTAG
TCATACGCTCGTCTCAA-------AGATTAAGCCATGCATGTGTAAGTCTAAATAGT---TTTAGTGTGA---AACTGCG
AATGGCTCA----------------------TTATATCAGTTATAGTTTATTTGGAATAGTTTCT-----------AT--
--ATGGATAACCGTAGTAATT------CTAGGGCTAATACATGTGTTTTCTTTGATGGGGAACCAT-------------T
AAAGAAGTACTTATTAGACATAAAACCAA----------------------------------------------TTTAA
T---GTTGTTGTGAGTA--ATAATA-ATTGTTGACTTTATCGCATGCTT-GTTAGTAGCGATGTTCCTTTCAAGTTTCTG
CCCTATCAGCTTTCG--------ATGGTAGTGTATTGGACTACCATGGCAGTAACGGGTAACGAAGA-ATTTGGGTTCGA
TTTCGGAGAGGGAGCCTGAGAGATG-GCTACCACATC-----CAAGGA----A--GGCAGCAGGCGC--------GTAAA
TTACCCAATCCTGACACAGGGAGGTAGTGACAATAAA-TCTCAATA-CGAAGCCTT-----TGGTTTTGTAATTGAATTG
AG---AACAAT-GTACA-----AT-CG--TTA---TCGATAAACAATTGGAGGGCAAGTCTGGTGCCAGCAGCCGCGGTA
ATTCCAGCTCCAATAGCGTATATTAACGTTGTTGCAGTTAAAAAGCTCGTAGTTGAA---------------GTTTAGAT
AATTTATTGAAAGTGTTTGTATTCGTACATTCTACTTTTATGTAAG--TTATCGACAT------------------CCA-
------------GGTGG---------------------GT---------------GC-----------------------
-------CATGG-----------------------TTCCTTCGGGGGTTAGGGTAC------------------------
---------------TAGACTGGTCATTTACTGTGAGAAAA--------TTAGAGTGTTTAAA-GCGGACTG--------
---TTCAGTTTGAATAGATTAG----------------------CATGGAATAATAAAATT----GGCTTTCATAGT---
--------CGATTTTTTG-GTTTGTTTATGAAAGCAAGATTAATAGGGACAGTTGGGGGTATTCATATTCAATAGCTAGA
GGTGAAATTCTATGATTTATGGAA------------------------------------GATGAACTA-----------
--------------------------------------------------------------------------------
------------------------------------------GTGCGAAAGCATTTACCAAGGATGTTTTCATTAATCAA
GAACGAAAGCTAGGGGATCAAAGAGGATTAGATACCCTCGTAGTCTTAGCTATAAACGATACCGACTAGGAGTTAGTAGA
TGATCCA-------------------------------------------------------------------------
------------------------------------ATGGTGTCTATTAGTACCTTATGAGAAATCAAAGTCTTTGGGTT
CCGGGGGGAGTATGGTCGCAAGGCTGAAACTTAAAGGAATTGACGGAAGGGCACCACC-AGGAGTGGAGCC-TGCGGCTT
AATTTGACTCAACACGGGGAAACTTACCAGGTCCAGACATAGGAAGGATTGACAGATTGATAGCTCTTTCT---TGATT-
----------------------------------------------CTATGGGTGGTGGTGCATGGCCGTTCTTAGTTGG
TGGAG-TGATTTG--TCTGGTTTATTCCGATAACGAACGAGACTTCCGCCTATTAGTTGGATGAAATGGGATTTTTAGCC
CCATTATTTTTTCATC----------------------------------------------------------------
------------------------------------------------------------------------AGCTTAG-
AGGGACACTG----TGCGTTTGAGTACAGGGAAGCTGGAAGCAATAACAGGTCTGTGATGC-CCTTAGATGTTCTGGGCT
GCACGCGCGCGACACTGAACTATTCAACGAGTAT-CTGAGT---------------------------------------
-------------------------------------CGATAGACTTGGGAAATCTTT-TGAAAATAGTTCGTGATGGGG
ATTGATGCTTGTAATTTTTCATCATGAAC--GAGGAATTCCTAGTAAATGCAAGTCATCAACT-TGCGTTGATTACGTCC
CTGCCCTTTGTACACACCGCCCGTCGCACCTACCGATTGGATGATCCGGTGAACACTTTGGATTTTTAATTATTTATCGG
CTT-----GCTGGTAGATAATTGAAGAGAAGTCGTGTAAATCTTATCATCTAGAGGAAGGTGAAGTCGTAACAAGGTTTC
CGTAGGTGAACCTGCGGAAG------------------------------------------------------------
------------------------------------------------------
>lcl|NT_187388.1_rrna_NR_146119.1_154624 [gene=RNA18SN4] [db_xref=GeneID:109864273] [product=RNA, 18S
pre-ribosomal N4] [transcript_id=NR_146119.1] [location=125931..127799] [gbkey=rRNA] Hsa
-TACCTGGTTGATCCTGC--------------------------------------------------------------
--------------------------------------------------------------------------------
--------------------------------------------------------------------------------
---------------------------------------------------------------------------CAGTA
GCATATGCTTGTCTCAA-------AGATTAAGCCATGCATGTCTAAGTACGCACGG-CC-GGTACAGTGA---AACTGCG
AATGGCTCA----------------------TTAAATCAGTTATGGTTCCTTTGGTCGCTCGCTCCT----CTCCTAC--
--TTGGATAACTGTGGTAATT------CTAGAGCTAATACATGCCGACGGGCGCTGACCCCCTTCGCGGGGG--G----G
ATGCGTGCATTTATCAGATCAAAACCAACC--CGGTCAGCCCCT-------CTCCGG-CCCCGGCCGGGGGGCGGGCGCC
GGCGGCTTTGGTGACTCTAGATAAC--CTCGGGCCGATCGCACGCCCCC-CGTGGCGGCGACGACCCATTCGAACGTCTG
CCCTATCAACTTTCG--------ATGGTAGTCGCCGTGCCTACCATGGTGACCACGGGTGACGGGGA-ATCAGGGTTCGA
TTCCGGAGAGGGAGCCTGAGAAACG-GCTACCACATC-----CAAGGA----A--GGCAGCAGGCGC--------GCAAA
TTACCCACTCCCGACCCGGGGAGGTAGTGACGAAAAA-TAACAATA-CAGGACTCTTTC-GAGGCCCTGTAATTGGAATG
AG---TCCACT-TTAAA-----TC-CT--TTA---ACGAGGATCCATTGGAGGGCAAGTCTGGTGCCAGCAGCCGCGGTA
ATTCCAGCTCCAATAGCGTATATTAAAGTTGCTGCAGTTAAAAAGCTCGTAGTTGGA---------------TCTTGGGA
GCGGGCGGGCGGTCCGCCGCGAGGCGAGCCACCGCCCGTCCCCGCC--CCTTGCC---------T---------CTCGG-
------------CGCCC---------------------CC---------------TC-----------------------
-------GATGC-----------------------TCTTAGCTGAGTGTCCCGCGG------------------------
---------------GGCCCGAAGCGTTTACTTTGAAAAAA--------TTAGAGTGTTCAAAGCAGGCCCG--------
---AGCCGCCTGGATACCGCAG----------------------CTAGGAATAATGGAATA----GGACCG-CGGTTCT-
-------ATTTTGTTGGT-TTTCGGAACTGAGGCCATGATTAAGAGGGACGGCCGGGGGCATTCGTATTGCGCCGCTAGA
GGTGAAATTCTTGGACCGGCGCAA------------------------------------GACGGACCA-----------
--------------------------------------------------------------------------------
------------------------------------------GAGCGAAAGCATTTGCCAAGAATGTTTTCATTAATCAA
GAACGAAAGTCGGAGGTTCGAAGACGATCAGATACCGTCGTAGTTCCGACCATAAACGATGCCGACCGGCGATGCGGCGG
CGTTATTC------------------------------------------------------------------------
-------C----------------------------CATGACCCGCCGGGCAGCTTCCGGGAAACCAAAGTCTTTGGGTT
CCGGGGGGAGTATGGTTGCAAAGCTGAAACTTAAAGGAATTGACGGAAGGGCACCACC-AGGAGTGGAGCC-TGCGGCTT
AATTTGACTCAACACGGGAAACCTCACCCGGCCCGGACACGGACAGGATTGACAGATTGATAGCTCTTTCT---CGATT-
----------------------------------------------CCGTGGGTGGTGGTGCATGGCCGTTCTTAGTTGG
TGGAG-CGATTTG--TCTGGTTAATTCCGATAACGAACGAGACTCTGGCATGCTAACTAGTTACGCGACCCCCGAGCGGT
CGGCGTC--CCCCAAC----------------------------------------------------------------

------------------------------------------------------------------------TTCTTAG-
AGGGACAAGTGGCGTT-------CAGCCACCCGAGATTGAGCAATAACAGGTCTGTGATGC-CCTTAGATGTCCGGGGCT
GCACGCGCGCTACACTGACTGGCTCAGCGTGTGC-CTACCC---------------------------------------
-------------------------------TA-CGCCGGCAGGCGCGGGTAACCCGT-TGAACCCCATTCGTGATGGGG
ATCGGGGATTGCAATTATTCCCCATGAAC--GAGGAATTCCCAGTAAGTGCGGGTCATAAGCT-TGCGTTGATTAAGTCC
CTGCCCTTTGTACACACCGCCCGTCGCTACTACCGATTGGATGGTTTAGTGAGGCCCTCGGATCGGCCCCGCCGGGGTCG
GCCCA--CGGCCCTGGCGGAGCGCTGAGAAGACGGTCGAACTTGACTATCTAGAGGAAGTAAAAGTCGTAACAAGGTTTC
CGTAGGTGAACCTGCGGAAGGATCATTA----------------------------------------------------
------------------------------------------------------
>lcl|NW_012156526.1_rrna_XR_001099855.1_15813 [locus_tag=SPRG_30244] [db_xref=GeneID:24143019] [product=18S ribosomal RNA] [transcript_id=XR_001099855.1] [location=complement(44366..45835)] [gbkey=rRNA] Spa
--ACCTGGTTGATCCTGCC-------------------------------------------------------------
--------------------------------------------------------------------------------
--------------------------------------------------------------------------------
---------------------------------------------------------------------------AGTAG
TCATACGCTTGTCTCAA-------AGATTAAGCCATGCATGTCTAAGTATAAACAATTT-TGTACTGTGA---AACTGCG
AATGGCTCA----------------------TTATATCAGTTATAGTCTACTTGGCAGTACCTTAC-------T--AC--
--TTGGATACCCGTAGTAATT------CTAGAGCTAATACATGCGTAAATACCCAACTGCTTGTCGGA------------
CGGGTAGCATTTATTAGATTGAAACCAATG--CGGCC--------------------------------------TCGGT
CGGTATTGTGTTGAATC--ATAATA--ACTGTGCGGAT--------CGC--TTCACAGCGATAAGTCAATTGAGTTTCTG
CCCTATCAGCTTTGG--------ATGGTAGGATATGGGCCTACCATGGCGTTAACGGGTAACGGGGA-ATTAGGGTTTGA
TTCCGGAGAGGGAGCCTTAGAAACG-GCTACCACATC-----CAAGGA----A--GGCAGCAGGCGC--------GTAAA
TTACCCAATCCTGACACAGGGAGGTAGTGACAATAAA-TAACAATG-CCGGGCTTTTCA--AGTCTGGCAATTGGAATGA
GA---ACAATT-TAAAT-----CC-CC--TTA---ACGAGGATCAATTGGAGGGCAAGTCTGGTGCCAGCAGCCGCGGTA
ATTCCAGCTCCAATAGCGTATATTAAAGTTGTTGCAGTTAAAAAGCTCGTAGTTGGA---------------TTTCTGGT
TTGAGCGTCCGGTCGAGTTTATCTCTGTACTATGGATGCTTGGGCC--A--------TATTTTG------------TGA-
------------GGGGG---------------------CG---------------CT-----------------------
-------TCTGC-----------------------CATTCAGTTGGTGGTTGTGTC------------------------
---------------GACTTGCATCGTTTACTGTGAAAAAT--------TAGAGTGTTTAAGC-AGGCGTTT--------
---GCTCATTTGAATACATTAG----------------------CATGGAATAATAAGATA----CGACCTTGGTGGTC-
-------TATTTTGTTGG-TTTGCACACCGAGGTAATGATTAATAGGGACAGTTGGGGGTATTCATATTTCAACGTCAGA
GGTGAAATTCTTGGATCGTTGAAA------------------------------------GATGAGCTT-----------
--------------------------------------------------------------------------------
------------------------------------------AGGCGAAAGCATTTACCAAGGATGTTTTCATTAATCAA
GAACGAAAGTTAGGGGATCGAAGATGATTAGATACCATCGTAGTCTTAACCATAAACTATGCCGACTCGGGATTGGCAGT
CGTTATTT------------------------------------------------------------------------
-------T---------------------------GAATGACCTTGTCAGCACCGTATGAGAAATCAAAGTCTTTGGGTT
CCGGGGGGAGTATGGTCGCAAGGCTGAAACTTAAAGGAATTGACGGAAGGGCACCACC-AGGAGTGGAGCC-TGCGGCTT
AATTTGACTCAACACGGGGAAACTTACCAGGTCCAGACATAGTAAGGATTGACAGATTGAGAGCTCTTTCT---TGATT-
----------------------------------------------CTATGGGTGGTGGTGCATGGCCGTTCTTAGTTGG
TGGAG-TGATTTG--TCTGGTTAATTCCGTTAACGAACGAGACCTCCGCGTGCTAAATAGTTCTGCTTACCAATTTGGTA
GGT-ATG------GAC----------------------------------------------------------------
------------------------------------------------------------------------TTCTTAG-
AGGGACTTTCAGTGACTAACTGAAGGAAGTTGGTTTGGAGGCAATAACAGGTCTGTGATGC-CCTTAGATGTTCTGGGCC
G-------------------------------------------------------------------------------
--------------------------------------------------------------------------------
--------------------------------------------------------------------------------
--------------------------------------------------------------------------------
--------------------------------------------------------------------------------
--------------------------------------------------------------------------------
------------------------------------------------------
>CABU01003659.782.2081
Eukaryota;SAR;Stramenopiles;Ochrophyta;Phaeophyceae;Ectocarpales;Scytosiphon;Ectocarpus siliculosus Esi
---TCTGGTTGATTCTGCC-------------------------------------------------------------
--------------------------------------------------------------------------------
--------------------------------------------------------------------------------
---------------------------------------------------------------------------AGTAG
TCATACGCTTGTCTCAA-------AGATTAAGCCATGCATGTCTAAGTATAAGCGC-TT-TATACTGTGA---AACTGCG
AATGGCTCA----------------------TTATATCAGTCATAGTTTATTTGAAAGTCCCTTAC-------T--AC--
--ATGGATAACCGTAGTAATT------CTAGAGCTAATACATGCACAAAAGCCCAACTGCCTCGGCGG-----------A
CGGGTTGCATTGATTAGACCGAAACCAATG--CGTCT--------------------------------------TCGGA
CGGTTTTGTGGTGAATC--ATAATC--AC----TTGCGGATCGCACGCT--TCGGCGGCGACGTTTCATTCAAGTTTCTG
CCCTATCAGCTTTGG--------ATGGTAGGGTATTGGCCTACCATGGCTTTAACGGGTAACGGGGA-ATTGGGGTTCGA
TTCCGGAGAGGGAGCCTGAGAGACG-GCTACCACATC-----CAAGGA----A--GGCAGCAGGCGC--------GTAAA
TTACCCAATCCTGACACAGGGAGGTAGTGACAATAAA-TAACAATG-CCGGGCTTTTAC--AAGTCTGGCAATTGGAATG
AG---AGCAAT-TTAAA-----TC-CA--TCA---TCGAGGATCAATTGGAGGGCAAGTCTGGTGCCAGCAGCCGCGGTA
ATTCCAGTTCCAATAGCGTATATTAAAGTTGCTGCAGTTAAAAAGCTCGTAGTTGGA---------------TTTGTGGC
GCGGCCGTCCGGCGGGGCTCCTTCATTGGGGCCGTTTGTCTGGTTT--TTCGGCCGCTCCATTCT-----------CGG-
------------GTAGT---------------------GT---------------GT-----------------------
-------CGCTG-----------------------GCATTAGGTTGTCGGCTTCTT------------------------
---------------CACGCCCGTCGTTTACTGTGAAAAAA--------TTAGAGTGTTCAAA-GCAGGCTT--------
---AGGCCATTGGATACATTAG----------------------CATGGAATAATGAGATA----GGGCCACGACGGTC-
-------TATTTTGTTGG-TTTGCACGTTGTGGTAATGATTAACAGGAACGGTTGGGGGTATTCGTATTCAATTGTCAGA
GGTGAAATTCTTGGATTTATGGAA------------------------------------GACGAACTA-----------
--------------------------------------------------------------------------------
------------------------------------------CTGCGAAAGCATTTACCAAGGATGTTTTCATTAATCAA
GAACGAAAGTTAGGGGATCGAAGATGATTAGATACCATCGTAGTCTTAACCATAAACTATGCCGACTAGGGATTGGCGGT
CGTTAATT------------------------------------------------------------------------
-------T----------------------------AAGGACTCCGTCAGCACCTTCCGAGAAATCAAAGTCTTTGGGTT
CCGGGGGGAGTATGGTCGCAAGGCTGAAACTTAAAGAAATTGACGGAAGGGCACCACC-AGGAGTGGAGCC-TGCGGCTT
AATTTGACTCAACACGGGGAAACTTACCAGGTCCGGACATAGTGAGGATTGACAGATTGAGAGCTCTTTCT---TGATT-
----------------------------------------------CTATGGGTGGTGGTGCATGGCCGTTCT-------
--------------------------------------------------------------------------------
--------------------------------------------------------------------------------
--------------------------------------------------------------------------------
--------------------------------------------------------------------------------
--------------------------------------------------------------------------------

```
--------------------------------------------------------------------------------
--------------------------------------------------------------------------------
--------------------------------------------------------------------------------
--------------------------------------------------------------------------------
--------------------------------------------------------------------------------
------------------------------------------------------
>AAFD02000029.651044.652838
Eukaryota;SAR;Stramenopiles;Ochrophyta;Diatomea;Bacillariophytina;Mediophyceae;Thalassiosira;Thalassiosira
pseudonana CCMP1335 Tps
---CCTGGTTGATTCTGCC-------------------------------------------------------------
--------------------------------------------------------------------------------
--------------------------------------------------------------------------------
---------------------------------------------------------------------------AGTAG
TCATACGCTCGTCTCAA-------AGATTAAGCCATGCATGTCTAAGTATAACCAT----TATACAGGAA---AACTGCG
AACGGCTCA----------------------TTATATCAGTTATTGTTTCTTTGATAGTCCCTTAC-------T--AC--
--TTGGATACCTGTAGTAATT------CTAGAGCTAATACATGCATCAATACCCGACTGTTCGCGGAA-----------G
--GGTAGTATTTATTAGGTATAGACCAAC---CGTCT--------------------------------------TCGGA
CGT-GCTTTGGTGATTC--ATAATA--ACTTATCGGATCGCATGGCTCC--ATGCCGGCGATGGATCATTCAAGTTTCTG
CCCTATCAGCTTTGG--------ATGGTAGTGTATTGGACTACCATGGCTTTAACGGGTAACGAATT-GTTAGGGCAAGA
TTTCGGAGAGGGAGCCTGAGAGACG-GCTACCACATC-----CAAGGA----A--GGCAGCAGGCGC--------GTAAA
TTACCCAATACTGAAACAGTGAGGTAGTGACAATAAA-TAACAATG-CCGGGCCTTTAC--AGGTCTGGCAATTGGAATG
AG---AACAAT-TTAAA-----TC-CC--TTA---TCGAGTATCAATTGGAGGGCAAGTCTGGTGCCAGCAGCCGCGGTA
ATTCCAGCTCCAATAGCGTATATTAAAGTTGTTGCAGTTAAAAAGCTCGTAGTTGGA---------------TTTCTGGC
AGGAGCGACCGGTCTCACACTCAGTGCGAGAACTCG--------TG--TTGTCTCTGGCCATCCT-----------TGG-
------------GGATA---------------------TC---------------CT-----------------------
-------GTTTG-----------------------GCATTAAGTTGTCGGCAGGG-------------------------
---------------GATACCCATCGTTTACTGTGAAAAAA--------TTAGAGTGTTTAAA-GCAGGCTT--------
---ATGCCGTTGAATATATTAG----------------------CATGGAATAATAAGATA----GGACCCTGG--TAC-
-------TATTTTGTTGG-TTTGCGCACCGAGGTAATGATTAAAAGAGACAGGCGGGGCTATTCGTATTGCATTGTCAGA
GGTGAAATTCTTGGATTTCTGCAA------------------------------------GACGAACTA-----------
--------------------------------------------------------------------------------
------------------------------------------CTGCGAAAGCATTTAGCAAGGATGTTTTCATTAATCAA
GAACGAAAGTTAGGGGATCGAAGATGATTAGATACCATCGTAGTCTTAACCATAAACTATGCCGACTCGGGATTGGCGGT
TGTTTT--------------------------------------------------------------------------
-------------------------------------TTGACTCCGCCAGCACCGTATGAGAAATCAAAGTCTTTGGGTT
CCGGGGGGAGTATGGTCGCAAGGCTGAAACTTAAAGAAATTGACGGAAGGGCACCACC-AGGAGTGGAGCC-TGCGGCTT
AATTTGACTCAACACGGGAAAACTTACCAGGTCCAGACATAGTGAGGATTGACAGATTGAGAGTTCTTTCT---TGATT-
----------------------------------------------CTATGGGTGGTGGTGCATGGCCGTTCTTAGTTGG
TGGAG-TGATTTG--TCTGGTTAATTCCGTTAACGAACGAGACCGCCGCCTGCTAAATAGTTCTGCGAATGGTTTTTCAT
TGGCAAG------AGC----------------------------------------------------------------
------------------------------------------------------------------------TTCTTAG-
AGGGACGTTCATTCTACAA----GATGAAGGAAGATGGCGGCAATAACAGGTCTGTGATGC-CCTTAGATGTCCTGGGCC
GCACGCGCGCTACACTGATGCACTCAACGAGCAT-ATAACC---------------------------------------
-------------------------------TT-GGCCGAGAGGCCTGGGTAATCTTG-TTAACATGCATCGTGATAGGG
ATAGATTATTGCAATTATTAATCTTGAAC--GAGGAATTCCTAGTAATCGCAGATCATCAATC-TGCAATGATTACGTCC
CTGCCCTTTGTACACACCGCCCGTCGCACCTACCGATTGGATGGTCCGGTGAGGAGTCGAGATTGTGGCCTGGTTCCTTT
A---------TTGGGATTTGGCTACGAGAACTTCTCCAAACCTTATCATCTAGAGGAAGGTGAAGTCGTAACAAGGTTTC
CGTAGGTGAACCTGCGGAAG------------------------------------------------------------
------------------------------------------------------
>lcl|NC_010143.1_rrna_XR_002461579.1_3941 [locus_tag=CYME_CMQ401R] [db_xref=GeneID:16996415] [product=18S
ribosomal RNA] [transcript_id=XR_002461579.1] [location=1042287..1044075] [gbkey=rRNA] Cme
------GGTTGATCCTGCC-------------------------------------------------------------
--------------------------------------------------------------------------------
--------------------------------------------------------------------------------
---------------------------------------------------------------------------AGTAG
TCATATGCTTGTCTCAA-------AGATTAAGCCATGCATGTCTAAGTATAAACCAGTT-TATACGGTGA---AACTGCG
AATGGCTCA----------------------TTAAAACAGTTATCGTTTATTTGATGGTACCTCCT----------AC--
--ATGGATAACCGTAGGAATT------CTACAGCTAATACATGCCCACACACCCGACTTTGGAA---G-----------G
---GTGGGATTTATCAGCTACAAAACCAA----ACCG--------------------------------------CCTCG
GCGGTCTCTGGTGATTC--ATGGTA--ACTGTCCGGATCGCATGTCCTT-CGGGATGGCGACAGATCATATAAATTTCTG
CCCTATCAACTTTCG--------ATGGTAGGATAGAGGCCTACCATGGTGGTAACGGGTAACGGGGA-ATTAGGGTTCGA
TTCCGGAGAGGGAGCCTGAGAAACG-GCTACCACATC-----CAAGGA----A--GGCAGCAGGCGC--------GCAAA
TTACCCAATCCTGACTCAGGGAGGTAGTGACAAGAAA-TAACGAGA-CCGGGCTCT-TC--GAGTCCGGTAATCGGAATG
AG---TACAAC-TTAAA-----TA-AC--TTA---ACGAGGATCCATTGGAGGGCAAGTCTGGTGCCAGCAGCCGCGGTA
ATTCCAGCTCCAATAGCGTATATTAAAGTTGTTGCAGTTAAAACGCTCGTAGTCGCA---------------GCTCGGAC
GCGGCCCACGGGCCGGCCGCAGGCTTGCGCCGGTGGGCCGGGTCTT--TCG------------------------GCTG-
------------GTGAG---------------------CG---------------CG-----------------------
-------CGGGG-----------------------TCCTCGCGGGCTCCGCGCGGC------------------------
---------------GAGCCAGCCCGTTTACTGTGAACAAA--------TTAGAGTGCTCCAGGCAGGC-----------
---GTTCGCATGCATACGTTAG----------------------CATGGAATAATAGAATA----GGACTT-GGGTCCT-
-------GTTTTGTTGGT-TTTGAGGGCCGAAGTAATGATGAATAGGGACAGTCGGGGGCATTCGTATTCCATTGTCAGA
GGTGAAATTCTTGGATTTATGGAA------------------------------------GACGAACAA-----------
--------------------------------------------------------------------------------
------------------------------------------CAGCGAAAGCATCTGCCAAGGATGTTTTCATTGATCAA
GAACGAAAGTTAGGGGATCGAAGACGATTAGATACCGTCGTAGTCTTAACCATAAACGATGCCGACTCGGGATCGGTGGA
GCACAAG-------------------------------------------------------------------------
--------------------------------------ATACTCCATCGGCACCGTAGGAGAAATCAAAGTGTTTGGGTT
CTGGGGGGAGTATGGTCGCAAGGCTGAAACTTAAAGGAATTGACGGAAGGGCACCACC-AGGAGTGGAGCC-TGCGGCTT
AATTTGACTCAACACGGGAAAACTTACCAGGTCCGGACATAGGGAGGATTGACAGATTGAGAGCTCTTTCT---TGATT-
----------------------------------------------CTATGGGTGGTGGTGCATGGCCGTTCTTAGTTGG
TGGAG-TGATTTG--TCTGGTTAATTCCGTTAACGAACGAGACCTTAACCTGCTAACTAGCGGCGCGAAACGC---GGTT
TCGCGGT------CGC----------------------------------------------------------------
------------------------------------------------------------------------TTCTTAG-
AGGGACGATCTGCGTCTA-----GCAGAGGGAAGTTTGAGGCAATAACAGGTCTGTGATGC-CCTTAGATGTCCTGGGCT
GCACGCGCGCTACACTGATGCAGGCAACGAGCGC-CGCTGCG--------------------------------------
-------------------------------------CCGAGAGGCGTGGCGAATCTG-CCAATCTGCATCGTGCTGGGG
ATAGACCCTTGGAATTATGGGTCTTCAACC--GAGGAATTCCTTGTAAGCGCGAGTCATCAGCT-CGCGCTGAATACGTCC
```

CTGCCCTTTGTACACACCGCCCGTCGCTCCTACCGATTGAATGATCCGGTGAGTTGTCCGGACGGGCGCGCCACCGGCCG
GTTC----GCCGGCGCGGGCGCGCCCGAAAGCTCAACAAACCTTATCATTTAGAGGAAGGAGAAGTCGTAACAAGGTTTC
CGTAGGTGAACCTGCGGAAGGATCATTC----------------------------------------------------
------------------------------------------------------
>lcl|NC_023998.1_rrna_XR_002608757.1_5470 [gene=rRNA_EukSSU] [locus_tag=Bathy11g00685]
[db_xref=GeneID:19013074] [product=ribosomal RNA EukSSU (n/a)] [transcript_id=XR_002608757.1]
[location=join(119769..120317,120751..121976)] [gbkey=rRNA] Bpr
--ACCTGGTTGATCCTGCC-------------------------------------------------------------
--------------------------------------------------------------------------------
--------------------------------------------------------------------------------
---------------------------------------------------------------------------AGTAG
TCATATGCTTGTCTCAA-------AGATTAAGCCATGCATGTCTAAGTATAAGCGTT----ATACTGTGA---AACTGCG
AATGGCTCA----------------------TTAAATCAGCAATAGTTTATTTGGTGGTGTTTACT----------AC--
--TCGGATAACCGTAGTAATT------CTAGAGCTAATACGTGCGTAAATCCCGACTTTTGAAG---------------G
---GACGTATTTATTAGATAAAG------------AC--------------------------------------CGACC
TCGTTTTGCGGTGAATC--ATGATA--ACTTTACGGATCGCATGGGCTT--GTCCCGGCGATGTTCCATTCAAATTTCTG
CCCTATCAACTTTCG--------ATGGTAGGATAGAGGCCTACCATGGTGGTAACGGGTGACGGAGA-ATTAGGGTTCGA
TTCCGGAGAGGGAGCCTGAGAAACG-GCTACCACATC-----CAAGGA----A--GGCAGCAGGCGC--------GCAAA
TTACCCAATCCTGACACAGGGAGGTAGTGACAATAAA-TAACAATA-CCGGGCTTTTTC--AAGTCTGGTAATTGGAATG
AG---AACAAT-CTAAA-----TC-CC--TTA---ACGAGGATCCATTGGAGGGCAAGTCTGGTGCCAGCAGCCGCGGTA
ATTCCAGCTCCAATAGCGTATATTTAAGTTGTTGCAGTTAAAAAGCTCGTAGTTGGA---------------TTTTGGTT
AAGAGGGCGCGGTCGGCCGTTTGGTCTGTACTGCGTTGTCTTGACT--TCCTGATGAGGACATGC---------T-----
--------------------------------------------------------------------------------
-------CTTG----------------------------GTTAACGCTGAGACATG------------------------
---------------GAGTCATCGTGGTTACTTTGAAAAAA--------TTAGAGTGTTCAAAGCGGGC-----------
---TTACGCTTGAATATATTAG----------------------CATGGAATAACACTATA----GGACTC-CTGTCCT-
--------ATCTCGTTGG-TCTCGGGATGGGAGTAATGATTAAGAGGAACAGTTGGGGGCATTCGTATTTCATTGTCAGA
GGTGAAATTCTTGGATTTATGAAA------------------------------------GACGAACTT-----------
--------------------------------------------------------------------------------
------------------------------------------CTGCGAAAGCATTTGCCAAGGATGTTTTCATTAATCAA
GAACGAAAGTTGGGGGCTCGAAGATGATTAGATACCATCCTAGTCTCAACCATAAACGATGCCGACTAGGGATTGGTGGA
TGTTAATT------------------------------------------------------------------------
------------------------------------GATGACTTCACCAGCACCTTATGAGAAATCAAAGTTTTTGGGTT
CCGGGGGGAGTATGGTCGCAAGGCTGAAACTTAAAGGAATTGACGGAAGGGCACCACC-AGGCGTGGAGCC-TGCGGCTT
AATTTGACTCAACACGGGGAAACTTACCAGGTCCAGACATAAGTATGATTGACAGATTGAGAGCTCTTTCT---TGATT-
----------------------------------------------CTATGGGTGGTGGTGCATGGCCGTTCTTAGTTGG
TGGAG-TGATTTG--TCTGGTTAATTCCGTTAACGAACGAGACCTCAGCCTGCTAAATAGTACGGCCCTATT----CTTA
GGGTCGC------GAC----------------------------------------------------------------
------------------------------------------------------------------------TTCTTAG-
AGGGACTATGTGCGTTTA-----GCACATGGAAGTTTGAGGCAATAACAGGTCTGTGATGC-CCTTAGATGTTCTGGGCC
GCACGCGCGCTACACTGACGGACTCAACGAGCTT-ATAACC---------------------------------------
-------------------------------TT-GGCCGAAAGGTCTGGGTAA--TCT-CCAAATTCCGTCGTGATGGGG
ATAGATTATTGCAATTATTAATCTTCAAC--GAGGAATGCCTAGTAAGCGCAAGTCATCAGCT-TGCGTTGATTACGTCC
CTGCCCTTTGTACACACCGCCCGTCGCTCCTACCGATTGAATGGTCCGGTGAAGCGTTCGGACTATGACTCTCTGACG-G
TT------CGCCGTCTAAGTGTCGTGGGAAGTTCGTTGAACCTTATCATTTAGAGGAAGGAGAAGTCGTAACAAGGTTTC
CGTAGGTGAACCTGCGGAAGGATCA-------------------------------------------------------
------------------------------------------------------
>AAXJ01016845.1.1085 Eukaryota;SAR;Alveolata;Protalveolata;Perkinsidae;Perkinsus;Perkinsus marinus ATCC 50983
Pma
--------------------------------------------------------------------------------
--------------------------------------------------------------------------------
--------------------------------------------------------------------------------
--------------------------------------------------------------------------------
--------------------------------------------------------------------------------
--------------------------------------------------------------------------------
--------------------------------------------------------------------------------
--------------------------------------------------------------------------------
------------------------------------------CACGGCT--TGTCCGGCGATGGACCATTCAAGTTTCTG
ACCTATCAGCTATGG--------ACGGTAGGGTATTGGCCTACCGTGGCGTTGACGGGTAACGGGGA-ATTAGGGTTCGA
TTCCGGAGAGGGAGCCTGAGAAACG-GCTACCACATC-----TAAGGA----A--GGCAGCAGGCGC--------GCAAA
TTACCCAATCCTGATACAGGGAGGTAGTGACAAGAAA-TAACAATA-CAGGGCAA--TT--CTGTCTTGTAATTGGAATG
AG---TAGATT-TTAAA-----TC-TC--TTT---ACGAGTATCAATTGGAGGGCAAGTCTGGTGCCAGCAGCCGCGGTA
ATTCCAGCTCCAATAGCGTATATTAAAGTTGTTGCGGTTAAAAAGCTCGTAGTTGGA---------------TTTCTGCC
TTGGGCGACCGATCCACCTTTCCTACGGGATTGGTCGGTATCAGGT--TTGACCTTGGCTTTTTC---------TTGGG-
------------ATTCG---------------------TG---------------CT-----------------------
-------CACGT-----------------------ACTTAACTGTGCGTTGACCGT------------------------
---------------GTTCCAAGACTTTTACTTTGAGGAAA--------TTAGAGTGTTTCAAGCAGGC--T--------
---TATGCCATGAATACATTAG----------------------CATGGAATAATAGGATA----TGA-CTTCGGTCAT-
-------ATTTTGTTGGT-TTCTAGGACTGAAGTAATGATTAATAGGGACAGTCGGGGGCATTCGTATTTAACTGTCAGA
GGTGAAATTCTTGGATTTGTTAAA------------------------------------GACGAACTA-----------
--------------------------------------------------------------------------------
------------------------------------------CTGCGAAAGCATTTGCCAAGGATGTTTTCATTGATCAA
GAACGAAAGTTAGGGGATCGAAGACGATCAGATACCGTCCTAGTCTTAACCATAAACTATGCCGACTAGGGATTGGGGGT
CGTTAATT------------------------------------------------------------------------
------------------------------------TTAGACGCCCTCAGCACCTCGTGAGAAATCAAAGTCTTTGGGTT
CCGGGGGGAGTATGGTCGCAAGGCTGAAACTTAAAGGAATTGACGGAAGGGCACCACC-AGGAGTGGAGCC-TGCGGCTT
AATTTGATTCAACACGGGAAAACTCACCAGGTCCAGACATAGGAAGGATTGACAGATTGATAGCTCTTTCT----GATT-
----------------------------------------------CTATGGGTGGTGGTGCATGGCCGTTCTTAGTTGG
TGGAG-TGATTTG--TCTGGTTAATTCCGTTAACGAACGAGACCTTAACCTGCTAA------------------------
--------------------------------------------------------------------------------
--------------------------------------------------------------------------------
--------------------------------------------------------------------------------
--------------------------------------------------------------------------------
--------------------------------------------------------------------------------
--------------------------------------------------------------------------------
--------------------------------------------------------------------------------
--------------------------------------------------------------------------------

```
--------------------------------------------------------------------------------
------------------------------------------------------
>lcl|NC_001144.5_rrna_NR_132222.1_4065 [locus_tag=RDN18-2] [db_xref=SGD:S000006483,GeneID:9164932]
[product=18S ribosomal RNA] [transcript_id=NR_132222.1] [location=complement(465070..466869)] [gbkey=rRNA] Sce
-TATCTGGTTGATCCTGCC-------------------------------------------------------------
--------------------------------------------------------------------------------
--------------------------------------------------------------------------------
---------------------------------------------------------------------------AGTAG
TCATATGCTTGTCTCAA-------AGATTAAGCCATGCATGTCTAAGTATAAGCAATT--TATACAGTGA---AACTGCG
AATGGCTCA----------------------TTAAATCAGTTATCGTTTATTTGATAGTTCCTTTA-------CTACA--
--TGGTATAACTGTGGTAATT------CTAGAGCTAATACATGCTTAAAATCTCGACCCTT-TGGAAG-----------A
---GATGTATTTATTAGATAAAAAATCAA----TGTC--------------------------------------TTCGG
AC--TCTTTGATGATTC--ATAATA--ACTTTTCGAATCGCATGGCCTT--GTGCTGGCGATGGTTCATTCAAATTTCTG
CCCTATCAACTTTCG--------ATGGTAGGATAGTGGCCTACCATGGTTTCAACGGGTAACGGGGA-ATAAGGGTTCGA
TTCCGGAGAGGGAGCCTGAGAAACG-GCTACCACATC-----CAAGGA----A--GGCAGCAGGCGC--------GCAAA
TTACCCAATCCTAATTCAGGGAGGTAGTGACAATAAA-TAACGATA-CAGGGCCCATTC--GGGTCTTGTAATTGGAATG
AG---TACAAT-GTAAA-----TA-CC--TTA---ACGAGGAACAATTGGAGGGCAAGTCTGGTGCCAGCAGCCGCGGTA
ATTCCAGCTCCAATAGCGTATATTAAAGTTGTTGCAGTTAAAAAGCTCGTAGTTGAA---------------CTTTGGGC
CCGGTTGGCCGGTCCGATTTTTTCGTGTACTGGATT---TCCAACG---GGGC-CTTTC----CT---------TCTGG-
------------CTA-----------------------------------------------------------------
-------ACCTT-----------------------GAG-TCCTTGTGGCTCTTGGC------------------------
---------------GAACCAGGACTTTTACTTTGAAAAAA--------TTAGAGTGTTCAAAGCAGGC--G--------
---TATTGCTCGAATATATTAG----------------------CATGGAATAATAGAATA----GGACGTTTGGTTCT-
-------ATTTTGTTGGT-TTCTAGGACCATCGTAATGATTAATAGGGACGGTCGGGGGCATCAGTATTCAATTGTCAGA
GGTGAAATTCTTGGATTTATTGAA------------------------------------GACTAACTA-----------
--------------------------------------------------------------------------------
------------------------------------------CTGCGAAAGCATTTGCCAAGGACGTTTTCATTAATCAA
GAACGAAAGTTAGGGGATCGAAGATGATCAGATACCGTCGTAGTCTTAACCATAAACTATGCCGACTAGGGATCGGGTGG
TGTTTTTT------------------------------------------------------------------------
-----------------------------------TAATGACCCACTCGGCACCTTACGAGAAATCAAAGTCTTTGGGTT
CTGGGGGGAGTATGGTCGCAAGGCTGAAACTTAAAGGAATTGACGGAAGGGCACCACC-AGGAGTGGAGCC-TGCGGCTT
AATTTGACTCAACACGGGGAAACTCACCAGGTCCAGACACAATAAGGATTGACAGATTGAGAGCTCTTTCT---TGATT-
----------------------------------------------TTGTGGGTGGTGGTGCATGGCCGTTCTTAGTTGG
TGGAG-TGATTTG--TCTGCTTAATTGCGATAACGAACGAGACCTTAACCTACTAAATAGTGGTGCTAGCATT---TGCT
GGTTATC------CAC----------------------------------------------------------------
------------------------------------------------------------------------TTCTTAG-
AGGGACTATCGGTTTCAA-----GCCGATGGAAGTTTGAGGCAATAACAGGTCTGTGATGC-CCTTAGACGTTCTGGGCC
GCACGCGCGCTACACTGACGGAGCCAGCGAGTC--------------------------------------------T-A
AC----------------------------CTT-GGCCGAGAGGTCTTGGTAATCTTG-TGAAACTCCGTCGTGCTGGGG
ATAGAGCATTGTAATTATTGCTCTTCAAC--GAGGAATTCCTAGTAAGCGCAAGTCATCAGCT-TGCGTTGATTACGTCC
CTGCCCTTTGTACACACCGCCCGTCGCTAGTACCGATTGAATGGCTTAGTGAGGCCTCAGGATCTGCTTAGAGAAGGGGG
CA------ACT-CCATCTCAGAGCGGAGAATTTGGACAAACTTGGTCATTTAGAGGAACTAAAAGTCGTAACAAGGTTTC
CGTAGGTGAACCTGCGGAAGGATCATTA----------------------------------------------------
------------------------------------------------------
>lcl|NC_003421.2_rrna_NR_151430.1_5627 [locus_tag=SPRRNA.44] [db_xref=GeneID:14217307] [product=18S ribosomal
RNA] [transcript_id=NR_151430.1] [location=complement(21289..23130)] [gbkey=rRNA] Spo
-TACCTGGTTGATCCTGCC-------------------------------------------------------------
--------------------------------------------------------------------------------
--------------------------------------------------------------------------------
---------------------------------------------------------------------------AGTAG
TCATATGCTTGTCTCAA-------AGATTAAGCCATGCATGTCTAAGTATAAGCAATTT-TGTACTGTGA---AACTGCG
AATGGCTCA----------------------TTAAATCAGTTATCGTTTATTTGATAGTACCTCAA-------CTACT--
--TG-GATAACCGTGGTAATT------CTAGAGCTAATACATGCTAAAAATCCCGACTTTTTTGGAAG-----------G
---GATGTATTTATTAGATAAAAAACCAA----TGCC--------------------------------------TTCGG
GCTTTTTTTGGTGAGTC--ATAATA--ACTTTTCGAATCGCATGGCCTT--GCGCCGGCGATGGTTCATTCAAATTTCTG
CCCTATCAACTTTCG--------ATGGTAGGATAGAGGCCTACCATGGTTTTAACGGGTAACGGGGA-ATTAGGGTTCGA
TTCCGGAGAGGGAGCCTGAGAAACG-GCTACCACATC-----CAAGGA----A--GGCAGCAGGCGC--------GCAAA
TTACCCAATCCCGACACGGGGAGGTAGTGACAAGAAA-TAACAATG-CAGGGCCCTTTC--GGGTCTTGTAATTGGAATG
AG---TACAAT-GTAAA-----TA-CC--TTA---ACGAGGAACAATTGGAGGGCAAGTCTGGTGCCAGCAGCCGCGGTA
ATTCCAGCTCCAATAGCGTATATTAAAGTTGTTGCAGTTAAAAAGCTCGTAGTTGAA---------------CTTTGAGC
CTGGTCGACTGGTCCGCCGCAAGGCGTGTTTACTGGTCATGACCGG---GGTCGTTAAC----CT---------TCTGG-
------------CAAAC---------------------TA---------------CT-----------------------
-------CATGT-----------------------TCT-TTATTGAGCGTGGTAGG------------------------
---------------GAACCAGGACTTTTACCTTGAAAAAA--------TTAGAGTGTTCAAAGCAGGCAAG--------
---TTTTGCTCGAATACATTAG----------------------CATGGAATAATAAAATA----GGACGTGTGGTTCT-
-------ATTTTGTTGGT-TTCTAGGACCGCCGTAATGATTAATAGGGATAGTCGGGGGCATTCGTATTCAATTGTCAGA
GGTGAAATTCTTGGATTTATTGAA------------------------------------GACGAACTA-----------
--------------------------------------------------------------------------------
------------------------------------------CTGCGAAAGCATTTGCCAAGGATGTTTTCATTAATCAA
GAACGAAAGTTAGGGGATCGAAGACGATCAGATACCGTCGTAGTCTTAACCATAAACTATGCCGACTAGGGATCGGGCAA
TGTTTCAT------------------------------------------------------------------------
-------T---------------------------TATCGACTTGCTCGGCACCTTACGAGAAATCAAAGTCTTTGGGTT
CCGGGGGGAGTATGGTCGCAAGGCTGAAACTTAAAGGAATTGACGGAAGGGCACCACAATGGAGTGGAGCC-TGCGGCTT
AATTTGACTCAACACGGGGAAACTCACCAGGTCCAGACATAGTAAGGATTGACAGATTGAGAGCTCTTTCT---TGATT-
----------------------------------------------CTATGGGTGGTGGTGCATGGCCGTTCTTAGTTGG
TGGAG-TGATTTG--TCTGCTTAATTGCGATAACGAACGAGACCTTAACCTGCTAAATAGCTGGATCAGCCATTTTGGCT
GATCATT------AGC----------------------------------------------------------------
------------------------------------------------------------------------TTCTTAG-
AGGGACTATTGGCATAAA-----GCCAATGGAAGTTTGAGGCAATAACAGGTCTGTGATGC-CCTTAGATGTTCTGGGCC
GCACGCGCGCTACACTGACGGAGCCAACGAGTTG-AAAAAAATCTTTTGATTT----------------------TTT-A
TC----------------------------CTT-GGCCGGAAGGTCTGGGTAATCTTG-TTAAACTCCGTCGTGCTGGGG
ATAGAGCATTGCAATTATTGCTCTTCAAC--GAGGAATTCCTAGTAAGCGCAAGTCATCAGCT-TGCGTTGAATACGTCC
CTGCCCTTTGTACACACCGCCCGTCGCTACTACCGATTGAATGGCTTAGTGAGGCCTCTGGATTGGCTTGTTTCTGCTGG
CA------ACGGCGGAAACATTGCCGAGAAGTTGGACAAACTTGGTCATTTAGAGGAAGTAAAAGTCGTAACAAGGTTTC
CGTAGGTGAACCTGCGGAAGGATCATTA----------------------------------------------------
------------------------------------------------------
```
